# Transposon-associated genetic structure of a fungal phytopathogen population of wheat

**DOI:** 10.64898/2026.06.22.733729

**Authors:** Huyen T. T. Phan, Manisha Shankar, Darcy A. B. Jones, Eiko Furuki, Kasia Rybak, Fiona Kamphuis, Hossein Golzar, Richard P. Oliver

**Affiliations:** Centre for Crop and Disease Management, School of Molecular and Life Sciences, Curtin University, Perth, WA, Australia; Department of Primary Industries and Regional Development, South Perth WA 6151; The University of Western Australia, Perth, WA, Australia; University of Nottingham, Nottingham, NG7 2RD, United Kingdom

**Keywords:** Adaptation, Septoria nodorum blotch, Population structure, Genetic diversity, Transposable Element (TE), Molly

## Abstract

Septoria nodorum blotch (SNB) is an economically important fungal disease of wheat caused by *Parastagonospora nodorum*. It is primarily controlled by the breeding of resistant wheat cultivars, but experience over the last 50 years shows that new pathogen populations soon evolve that are more virulent on the current popular cultivars. In this study, we assembled a panel of 360 *P. nodorum* isolates. The collection resolved into eight subpopulations. One core and seven transient populations were found possessing contrasting characters in term of spatial and temporal distribution, mating-type, effector haplotypes and patterns of intact and degraded copies of a Tc-1 mariner transposon, called Molly. Molly can proliferate and randomly insert throughout the fungal genome. Its multiplication in sexual population likely triggered RIP which partially explains the extensive genetic diversity and explains the ability to form new adapted lineages and the observed population structure of this important pathogen of wheat. When tested on wheat, the recently emerged groups exhibited greater pathogenicity on modern elite cultivars consistent with the low-amplitude boom-and-bust cycle observed previously. It is possible that active copies of Molly transpose and contribute to both the birth and death of the transient groups. This study identified and characterised a fungal specific transposable element (TE) which plays a vital role in shaping Australian *P. nodorum* population structure and creating extensive genetic diversity which potentially leads to the pathogen’s better adaptation. The study suggests practical measures to improve the efficiency and longevity of resistance breeding for SNB.

## INTRODUCTION

Diseases in agroecosystems are mainly controlled by the provision of pesticides and genetically resistant cultivars. Many pathogen species have demonstrated the ability to repeatedly evolve to defeat both strategies. *Parastagonospora nodorum* causes septoria nodorum blotch (SNB) of wheat (*Triticum spp*.) (Quaedvlieg et al. 2013; Solomon et al. 2006). The fungus causes significant damage to the leaves and glumes lowering yield and grain quality (Oliver et al. 2012). The first genome assembly was developed in 2005 leading directly to the discovery of NEs as well as other features of significance (Hane et al. 2007). Since then, several other NEs have been discovered. These NEs interact directly or indirectly with their corresponding host receptors encoded by sensitivity genes, in an inverse gene-for-gene fashion and subtly exploit defence responses for their proliferation (Faris et al. 2010; Friesen & Faris, 2021; Lorang et al. 2012; Oliver & Solomon, 2010; Shi et al. 2016). The model helps elucidate how different disease components contribute to the complexity of disease expression at the molecular level in the Pleosporales (Faris & Friesen, 2020; Friesen & Faris, 2021; Kariyawasam et al. 2023). However, the level of disease in the field is not well predicted by considering the currently isolated NEs and matching receptor genes and SNB remains a significant threat to wheat production in many parts of the world with sporadic heavy rainfall and in agrosystems where stubble is retained.

Previous population structure analyses of *P. nodorum* using restriction fragment length polymorphisms (RFLP), simple sequence repeats (SSR) or single nucleotide polymorphisms (SNP) revealed high levels of genetic variability but did not uncover significant population differentiation (Blixt et al. 2008; Keller et al. 1997; Lin et al. 2020; Murphy et al. 2000; Stukenbrock et al. 2006). Richards et al. [17] used genome-wide genomic variation and separated two United States populations located in the Upper Midwest spring wheat and Southern/Eastern winter wheat production regions.

Our previous investigation on the population structure of Western Australian (WA) *P. nodorum* used a collection of 155 isolates and 32 neutral SSR markers. We showed that the population clustered into five groups of two core and three transient groups with contrasting properties (Phan et al. 2019). The two core groups were found throughout the collection locations and times and encompassed 80% of the isolates. The other three groups were restricted to specific times and locations. Changes in transient group prevalence coincided with the extensive adoption of a single or a small group of widely planted wheat varieties, to which the newly emerged groups were more virulent, a phenomenon we called the low-amplitude boom-and-bust (LABAB) cycle, in contrast with the classical boom-and-bust cycle characterised by a catastrophic breakdown of resistance and typically found in biotrophic diseases such as rusts and powdery mildews (Oliver, 2024). We hypothesised that the core groups maintain population variation from which emerge sub-groups that are adapted to new cultivar releases (Phan et al. 2019).

A recent *P. nodorum* pangenome study using 156 WA isolates also found a large diverse core and a few small, homogeneous sub-populations. Of more than 1.3 million SNP variants detected across the pangenome, 78% were RIP-like C:G↔T:A mutations. RIP is a fungal-specific genome defence mechanism against transposon and viral propagation first found in *Neurospora crassa* (Selker et al. 1987). It is active in *P. nodorum* and contributes to genome variability (Bertazzoni et al. 2021; Fouché et al. 2022; Hane & Oliver, 2010; Pereira et al. 2021; Rouxel et al. 2011; Testa et al. 2016). RIP is located unevenly across the genome with some long contiguous regions being known as Large RIP Affected Regions (LRAR) (van Wyk et al. 2019). The high-proportion of RIP-like mutations suggested widespread ‘RIP-leakage’ from transposon-rich repetitive sequences into non-repetitive regions. The study also revealed that the variation was balanced by negative selection against nonsynonymous SNPs observed within protein-coding sequences (Jones et al. 2024).

Transposable elements (TEs) are major components of most eukaryotic genomes and play key roles in evolution as drivers for generating adaptive genetic variations (Abraham et al. 2024; Badet et al. 2024; Fouché et al. 2022; Oggenfuss et al. 2021; Wells & Feschotte, 2020). The TE can promote genome instabilities by non-allelic homologous recombination and can lead to chromosomal rearrangements (Argueso et al. 2008; Fouché et al. 2025). Gene regulation or functional coding sequences can be disrupted or altered by TE mobilisation and can affect host fitness (Abraham et al. 2024; Chuong et al. 2017; Dubin et al. 2018). The occurrence of TE bursts can trigger RIP which often inactivates the TE.

Recent advances in long-read genome sequencing technologies have made fungal whole-chromosome assemblies available (Badet et al. 2020; Derbyshire et al. 2017; Koren et al. 2017; Moolhuijzen et al. 2018; Richards et al. 2018) and large-scale TE pangenomic comparative studies possible (Oggenfuss et al. 2021; Pereira et al. 2021). The most recent TE pangenonic study on *P. nodorum* was carried out by Pereira et al. (2021) on a global collection of 366 genomes. Their study found little variance among global populations in TE composition and copy number despite differences in demographic history and recently active, high-copy TEs which were postulated to have escaped from genomic defences. This comprehensive work showed the complicated evolutionary balances influenced by multiple factors such as TE activity, selection pressure and genomic suppression (Pereira et al. 2021).

This study utilised two large WA isolate collections consisting of 248 isolates collected over a 50-year period (1968-2018) and 112 isolates collected in 2019-2021 and all analysed using whole genome sequencing. We confirm the pattern of core and low-amplitude boom-and-bust cycles in transient groups. We also show that newly emerged groups are more pathogenic on modern wheat elite lines and identify potential role of Molly in shaping the pathogen population structure and adaptation. Patterns of Molly content in the pangenome revealed a potential role of RIP in genomic defence response and diversity generation in shaping this pathogen genetic structure. The study will have a direct impact on ongoing wheat breeding programs by directing the choice of isolates to be used when seeking durable SNB disease resistance. It also highlights the potential value of maximizing genetic diversity in wheat cultivars by either rotations or mixtures to limit the damage caused by SNB.

## MATERIALS AND METHODS

### Fungal collections

The study involved three Australian *P. nodorum* collections: 151 isolates used in Jones et al. (2024); 97 isolates provided by the WA Department of Primary Industries and Regional Development (DPIRD); and 112 isolates collected from 2019 to 2021 in an Australian Grain Technologies disease nursery at Northam, WA. The isolates were collected from 38 locations across all wheat growing areas in WA and over 50-year period (SFigure 1). Details of the 360 isolates are listed in STable 1.

### Short-read sequencing and genome assemblies

Illumina short-reads from genomic DNA were obtained for all 360 *P. nodorum* isolates. Genomic DNA of the 97 DPIRD and 112 current isolates were obtained using a Qiagen DNeasy-Plant-Mini-kit (Venlo, Netherlands, Catalogue ID: 69104). The sequencing reads were evaluated with FastQC v0.11.9 (Andrews, 2010) and subsequently trimmed using trimmomatic (Bolger et al. 2014). The quality of the trimmed reads was tested again with FastQC (fastqc/0.11.9--hdfd78af_1, Andrews (2010)) and only values of >= 30 were used for further analysis. Genomes were assembled using Spades version 3.13.0 (Bankevich et al. 2012) (parameters: LJLJcareful LJLJcovLJcutoff auto) with different kmers (parameters: -k 31, 51, 71, 81, 101, 127). Quality control statistics for these genome assemblies are reported in STable 1 which were created by Quast v5.2.0 (Gurevich et al. 2013) and SFigure 2 using BUSCO v6.0.0 (Manni et al. 2021).

### Long-read sequencing and genome assemblies

Three isolates (15FG38, 16FG168 and PN315) were chosen for nanopore sequencing. Their high molecular weight DNA was extracted using the protocol described by John Tyson (https://www.protocols.io/view/phenol-chloroform-genomic-dna-extraction-from-tiss-4r3l28mq4l1y/v1) and cleaned by Jones et al. (2021). The genomes of 15FG38 & 16FG168 and PN315 were sequenced following the manufacturer’s guidelines for the Ligation Sequencing Kit SQK-LSK109 and SQK-LSK114 (Oxford Nanopore Technologies, Cambridge UK) on two R9.4.1 and a R10.4.1 flow cells respectively; using the MinION sequencing device and the ONT MinKNOW Software (Oxford Nanopore Technologies, Cambridge UK). PN315 long-read nanopore assembly was generated following the pipeline developed by Johannes Debler (https://github.com/JWDebler/nanopore_kit14_assembly). Nanopore assemblies of 15FG38 and 16FG168 were created using similar approach where raw reads were cleaned using chopper v0.10.0 (De Coster & Rademakers, 2023) and seqkit_2.5.1 (Shen et al. 2024). De-novo genome assemblies were generated using nextdenovo v2.5.2 (Hu et al. 2024), then later polished with medaka v2.1.0 (https://github.com/nanoporetech/medaka) and ragtag v2.1.0 (Alonge et al. 2022); sorted by seqkit v2.5.1. A final polishing step was done on all three genomes using Polypolish v0.6.0 program with illumina short-reads (Wick & Holt, 2022). BUSCO was used to assess the completeness of the nanopore genomes. Annotation of the genomes were done using Liftoff v1.6 3 (Shumate & Salzberg, 2021) from the annotations created for 15FG38 and 16FG168 in Jones et al. (2024).

### Variant calling and SNP selection

Program BWA v0.7.17 (Li, 2013) was used to map the cleaned reads of each isolate to the *P. nodorum* reference isolate SN15 (Bertazzoni et al. 2021). Outputs from the alignments were used for genome-wide SNP variant calling using a standard pipeline in the Toolkit for Genome Analysis (GATK v4.2.6.1, https://software.broadinstitute.org/gatk/) suite (McKenna et al. 2010; Poplin et al. 2018). We followed the recommendations for calling variants including: 1) alignment to SN15 reference genome, 2) Mark MarkDuplicates, 3) AddOrReplaceReadGroups, 4) Base quality score recalibration (BQSR), 5) Applying the model generated by BQSR to adjust the base quality scores, 6) Haplotype calling using GATK, 7) Consolidation of GVCFs across samples, 8) Joint genotyping on a cohort, 9) Separate SNPs from indels - GATK SelectVariants. Finally, the discovered SNPs were hard-filtered using GATK VariantFiltration with the following parameters: QD < 2.0, QUAL < 30.0, SOR > 3.0, FS > 60.0, MQ < 40.0, MQRankSum < −12.5, ReadPosRankSum < −8.0. These are recommendations for best practices in GATK4 manuals (Caetano-Anolles, 2023. All 360 isolates remained after filtering for missing value per individual of 0.05% and uniqueness.

The identified SNPs was subsequently pruned for population structure analysis using R package SNPRelate v1.38.0 (Zheng et al. 2012) with the following criteria: maximum missing genotype and loci of 0.05, minimum non-major allele frequency of 0.01 and a linkage disequilibrium threshold of 0.2. A vcf file containing selected 18,208 SNP was successively used for population analysis.

### Phylogenetic tree and population structure analysis

Population structure inference of the 360 *P. nodorum* isolate-collection was conducted using the pruned SNP data-set and STRUCTURE v2.3.4 (Pritchard et al. 2000). The most appropriate number of clusters (K) existed in the collection was determined by running the STRUCTURE program with 10000 replicate burn-in period and 10000 MCMC replicates for a range of values of K between 1 and 15. This process was repeated 10 times for each K to account for random starting points. The best value of K was selected using STRUCTURE HARVESTER (Earl & Vonholdt, 2012) using the method described by Evanno et al. (2005). Final assignments of each isolate to the identified optimum subpopulations were carried out by running STRUCTURE with selected K value with 20000 replicate burn-in period and 20000 MCMC replications. The optimum K value was also used to run Discriminant Analysis of Principal Components (DAPC) in R package adegenet v2.1.6, a multivariate method developed by Jombart et al. (2010) to confirm the grouping assignment of each individual. Given a set of isolates and a group number, DAPC finds linear functions of PCs of the genotype data to classify the isolates in the groups in the way that maximise differences between groups and minimise differences within groups. Cross-validation was conducted to determine optimum number of PCs to include in the model using a range of PC retention numbers (90%:10% training: validation stratified splits) so that it minimised the mean squared error of reclassification (Jombart et al. 2010).

An unweighted-pair-group-method-with-arithmetic-mean (UPGMA) hierarchical clustering algorithm in poppr was used to create a phylogenetic tree (Figure 1) from the selected SNPs using functions ‘aboot’ with 100 bootstraps and genetic distance matrix generated from function ‘bitwise.dist’ in ‘poppr’ package v2.9.6 (Kamvar et al. 2015). Phylogenetic tree and population statistics were generated and visualised using R v4.4.0 with packages: adegenet v2.1.6 (Jombart, 2008), treemap v2.4-4 (Tennekes & Ellis, 2017), vcfR v2.1.10 (Knaus & Grünwald, 2017), poppr v2.9.6 (Kamvar et al. 2015), ggplot2 v3.5.1 (Wickham, 2016), ape v5.8 (Paradis & Schliep, 2019), pegas v1.3 (Paradis, 2010) and iTOL v6.9 (Letunic & Bork, 2024).

**Figure 1:**
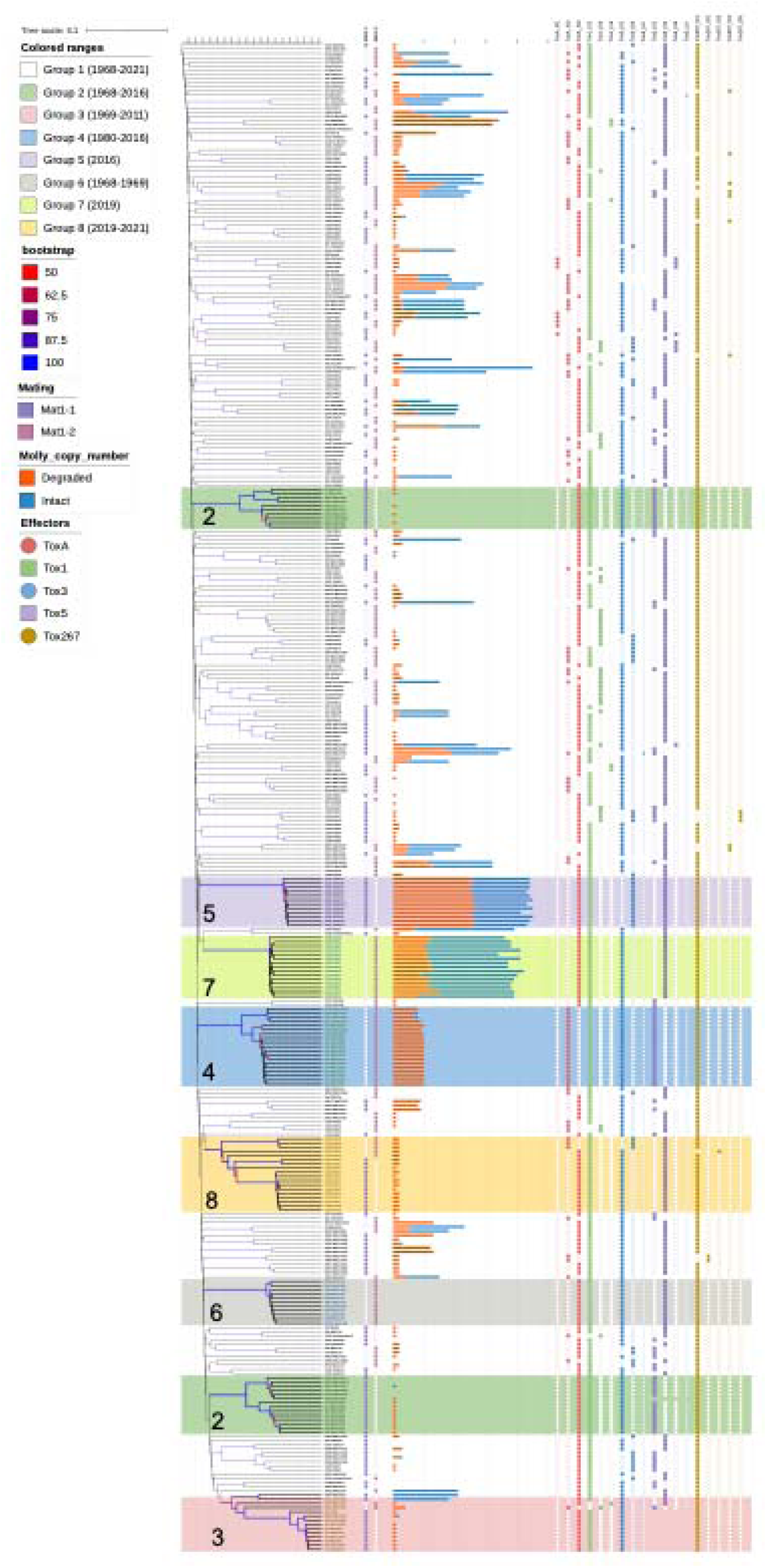
Phylogenetic tree of 360 Australian *P. nodorum* isolates with genetic distance scale indicated, purple squares denotes mating types of each isolate; bar-charts represent the number of Molly copy numbers and their conditions (Intact &Degraded); red circles, green squares, blue circles & stars, dark blue squares & stars, brown circles indicates different effector isoforms of *SnToxA, SnTox1, SnTox3, SnTox5 and SnTox267*, respectively. Isolate identities denoted by “collection_isolate name”; where “collection” include: C = Current collection, O = Old collection, DIP = DPIRD collection.

### Determination of the mating genotype and standardised Index-of-Association (rbarD)

Mating types of these *P. nodorum* isolates were determined by the presence of MAT1-1 or MAT1-2 obtained from MetaEuk v5-34c21f2 package (Levy Karin et al. 2020) using the reference mating-type gene sequences from NCBI database (AF322008 and AY212019). A Chi square test (λ^2^) was used to determine if the observed MAT1-1:MAT1-2 ratio departed from the standard 1:1. Standardised index of association (’rbarD’) from (2001) was calculated for each of the sub-populations using the poppr package v2.9.6 in R (Kamvar et al. 2014). This standardised index of association helps to determine if these tested population are in modes of sexual or clonal reproduction based on linkage disequilibrium. The rbarD values and a *P* values, obtained using a 999-permutation test, were used to determine if a population is sexually reproduced (SFigure 3). These estimates of linkage disequilibrium from random genomic fractions were summarised using a histogram with a vertical value of each population’s rbarD (https://grunwaldlab.github.io/poppr/reference/samp.ia.html).

### Identification of effector isoforms

Effectors protein sequences (SnTox1: SNOG200078; SnTox3: SNOG08981; SnToxA: SNOG16571; SnTox267: CJJ_13380 [74]; SnTox5: MW716005) were used as reference sequences for ‘easy-predict’ function in MetaEuk v5-34c21f2 package (Levy Karin et al. 2020) with default settings (https://github.com/soedinglab/metaeuk?tab=readme-ov-file#easy-predict-workflow). Nucleotide sequences of those effectors were accurately obtained and used for haplotype/isoform analyses.

### Genomic variation between the core and transient groups

Sixteen genome assemblies obtained from the *Sequencing-and-genome-assembly* section were used for genomic variation analysis using Mauve v2.4.0 (Darling et al. 2004) in Geneious-Prime-2025 (www.geneious.com). Since variation exists in the core group, two genomes of the core group (15FG38 and Northam_WGT) were selected as the references for this alignment (Table 1). To identify common pattern for the differences in genomic contents, two to three genomes from each transient group were aligned to either one or both of the reference genomes (15FG38 and Northam_WGT). Mauve aligns syntenic blocks by aligning orthologous and xenologous regions from one genome to the other to produce global alignment of each locally collinear block that has sequence elements conserved between the genome pairs (Darling et al. 2004). The un-aligned regions within these syntenic blocks and unique sequences were searched for homologs via Blastn search function in NCBI as of March 2024 (https://blast.ncbi.nlm.nih.gov/Blast.cgi).

**Table 1:** Differences in gene contents between the core and transient groups.

| Comparison<br>Group 1<br>versus | Isolate | Total<br>loci<br>tested | Unique<br>loci<br>tested | Size range<br>(kb) | *Presence in the P.*<br>*nodorum* reference<br>gene pool (%) | Sources |  |  |  |  |  |  |
| --- | --- | --- | --- | --- | --- | --- | --- | --- | --- | --- | --- | --- |
|  |  |  |  |  |  | SN15 | USA/S<br>N15 | USA | Other - HGT<br>sequences | Note* | no-hit | Molly |
| Group 2 | 15FG38 vs WAC4648 | 168 | 32 | 0.5 - 701 | 93 | 7 | 89 | 58 | 2 | *Alternaria alternata* | 12 | 0 |
|  | 15FG38 vs WAC13529 | 120 | 26 | 0.5 - 418 | 97 | 19 | 73 | 23 | 1 | *Alternaria alternata* | 4 | 0 |
|  | Northam_WGT vs WAC4648 | 187 | 24 | 0.5 - 616 | 94 | 50 | 70 | 56 | 2 | *Alternaria alternata* | 9 | 0 |
|  | Northam_WGT vs WAC13529 | 166 | 17 | 0.5 - 200 | 95 | 59 | 54 | 44 | 2 | *Alternaria alternata* | 7 | 0 |
| Group 3 | 15FG38 vs SN15 | 177 | 28 | 0.5 - 107 | 100 | 83 | 94 | 0 | 0 |  | 0 | 0 |
|  | 15FG38 vs WAC9178 | 170 | 31 | 0.5 - 1,107 | 99 | 86 | 81 | 0 | 0 |  | 2 | 1 |
|  | Northam_WGT vs Sn15 | 146 | 30 | 0.5 - 108 | 100 | 124 | 22 | 0 | 0 |  | 0 | 0 |
|  | Northam_WGT vs WAC2810 | 160 | 32 | 0.5 - 987 | 99 | 121 | 36 | 2 | 0 |  | 1 | 0 |
| Group 4 | 15FG38 vs WAC13526 | 125 | 17 | 0.5 - 380 | 89 | 0 | 72 | 37 | 1 |  | 14 | 1 |
|  | 15FG38 vs WAC13074 | 124 | 19 | 0.5 - 801 | 95 | 21 | 61 | 34 | 1 |  | 6 | 1 |
|  | Northam_WGT vs wac13526 | 143 | 12 | 0.5 - 423 | 94 | 23 | 75 | 33 | 1 |  | 8 | 3 |
|  | Northam_WGT vs wac13070 | 166 | 21 | 0.5 - 470 | 95 | 22 | 89 | 44 | 1 |  | 7 | 3 |
| Group 5 | 15FG38 vs PN155 | 178 | 24 | 0.5 - 72 | 94 | 53 | 65 | 34 | 2 | *Alternaria alternata* | 10 | 14 |
|  | 15FG38 vs PN152 | 169 | 17 | 0.5 - 72 | 95 | 46 | 71 | 33 | 3 | *Alternaria alternata* | 9 | 7 |
|  | Northam_WGT vs Pn155 | 195 | 26 | 0.5 - 127 | 95 | 70 | 53 | 50 | 2 | *Alternaria alternata* | 6 | 14 |
|  | Northam_WGT vs 16FG166 | 179 | 24 | 0.5 - 587 | 93 | 55 | 73 | 34 | 2 | *Alternaria alternata* | 11 | 4 |
| Group 6 | 15FG38 vs DPIRD7 | 154 | 18 | 0.5 - 1,473 | 95 | 39 | 53 | 46 | 8 |  | 8 | 0 |
|  | 15FG38 vs DPIRD8 | 154 | 37 | 0.5 - 1,551 | 93 | 37 | 57 | 47 | 3 |  | 10 | 0 |
|  | Northam_WGT vs DPIRD7 | 153 | 20 | 0.5 - 1,388 | 93 | 23 | 81 | 38 | 8 |  | 3 | 0 |
|  | Northam_WGT vs DPIRD9 | 160 | 24 | 0.5 - 1,444 | 93 | 27 | 90 | 32 | 2 |  | 9 | 0 |
| Group 7 | 15FG38 vs PN343 | 181 | 33 | 0.5 - 755 | 90 | 15 | 83 | 57 | 1 | *Ptt\*\**, *Bipolaris sorokiniana* | 18 | 7 |
|  | 15FG38 vs PN315 | 162 | 18 | 0.5 - 934 | 93 | 15 | 92 | 38 | 2 | *Ptt\*\**, *Bipolaris sorokiniana* | 12 | 3 |
|  | Northam_WGT vs PN343 | 177 | 27 | 0.5 - 810 | 92 | 20 | 104 | 35 | 1 | *Ptt\*\**, *Bipolaris sorokiniana* | 14 | 3 |
|  | Northam_WGT vs PN282 | 171 | 26 | 0.5 - 836 | 93 | 20 | 94 | 43 | 2 | *Ptt\*\**, *Bipolaris sorokiniana* | 10 | 2 |
| Group 8 | 15FG38 vs PN378 | 195 | 36 | 0.5 - 384 | 93 | 54 | 77 | 50 | 0 |  | 14 | 0 |
|  | 15FG38 vs PN379 | 144 | 20 | 0.5 - 640 | 96 | 35 | 69 | 34 | 0 |  | 6 | 0 |
|  | Northam_WGT vs PN378 | 168 | 37 | 0.5 - 375 | 95 | 29 | 84 | 46 | 0 |  | 9 | 0 |
|  | Northam_WGT vs PN322 | 132 | 21 | 0.5 - 922 | 96 | 21 | 76 | 29 | 1 |  | 5 | 0 |
| Total/Mean |  | 4524 | 697 |  | 94 | 1174 | 2038 | 977 | 48 |  | 224 | 63 |

\*Foreign DNAs which hit interesting species
\*\**Ptt* is *Pyrenophora teres f. ter*

### Molly copy numbers

A method to identify Molly copy numbers was initially established using four reference genomes using both short/long-read genomes of SN15 (Bertazzoni et al. 2021) and three telomere-to-telomere nanopore assemblies created in this study. This was later applied to all 360 short-read genomes. For short-read data, all trimmed reads were mapped to Molly reference sequence in NCBI (AJ488502) and their genome assemblies using two programs: BWA-MEM2 v2.2.1 (Li, 2013) and BBMap v39.19 (https://github.com/BioInfoTools/BBMap). Variation in read-coverage of the Molly and their corresponding genome assemblies are used as a proxy for Molly copy number variation among these isolates. Molly copy numbers of long-read genomes were determined by blasting the reference Molly sequence to these genomes using Custom Blast function in Geneious-Prime-2025. Classification of either “Intact/Weak” or “Strong/Degraded” modifications of Molly for short-read genomes were obtained by blasting the Molly sequence to each genome. Strongly-degraded copies were readily identified by their unique-sequence nature. The intact copies were defined by the difference between the total copies, identified from read-coverage method, and the strongly-degraded ones. The intact/degraded copy numbers were validated again by coverage of the contig/scaffolds which hit Molly sequence. All molly copy numbers and classifications of 360 isolates are presented in STable 2.

### Genomic compartment identification

AT-rich regions are a signature of RIP. To determine if Molly and AT-rich regions are co-located and identity affects Molly might have on genome compartmentalisation, GC-content analysis was caried out with 3 molly-containing reference genomes using OcculterCut v1.1 (Testa et al. 2016). In addition, regions impacted by RIP, LRAR, were identified with RIPper (van Wyk et al. 2019). GC-contents and RIP-like SNP of Molly sequences were analysed using Geneious-Prime-2025.

### Whole plant infection assay

Three current popular Australian wheat cultivars (Calibre, Mace, and Scepter; from Era III in Phan et al. (2020), one once popular cultivar (Wyalkatchem; Era II) and three old (EraI, Calingiri, Eradu, Halberd) were used in seedling infection assays as described in Solomon et al. (2003). Their effector sensitivity profiles are reported in STable 3. Significant differences in disease scores for each group, in each Era and each wheat line was determined based on analysis of variance (ANOVA) and the Least Significant Difference test using statistical functions in core R and agricolae v1.3 R packages.

## RESULTS

### Short read genome assemblies and variant calling

Raw reads of the 209 newly sequenced isolates were obtained using short-read Illumina sequencing. The quality was high (SFigure 4) and the average size for the generated genome assemblies was 37.4±0.095Mb which is consistent with the SN15 reference isolate (Bertazzoni et al. 2021) and other *P. nodorum* pan-genome studies (Jones et al. 2024; Richards et al. 2019). The mean completeness estimated via BUSCO scores was high (93.96%) (SFigure 2). General properties of these genomes were summarised in STable 4.

Variant calling was carried out for all 360 isolates and yielded 1.38 million high quality short variants which include SNPs and small indels (insertions and deletions). A subset of 18,208 SNPs was selected for phylogenetic and population diversity analysis which will be presented in the following sections.

### Population structure

PCA analysis, built from PC1 and PC2 revealed five distinct clusters and one conglomerate in the scatterplot (Figure 2A). A population structure of eight groups (SFigure 5A) was found to be the optimum for this collection using STRUCTURE (Pritchard et al. 2000). The model of eight groups was used for DAPC analysis using the adegenet package (Jombart et al. 2010). This analysis deployed 150 PCA as the optimum number determined by the cross-validation test (SFigure 5B). The level of agreement between STRUCTURE and DAPC on the group assignment was high with only 9 out of 360 assignments being inconsistent. These 9 isolates were in the closely related groups 3, 4 and 8 (Figure 2A and 2B) and were assigned according to results from the STRUCTURE program. The group assignments and closely related individuals are consistent with results from Phan et al. (2019) and Jones et al. (2024) as shown in Table 2. In this study, group 1 comprises the core population and corresponds to group 2 in Phan et al. (2019) and 4 in Jones et al. (2024). Phan et al. (2019) identified another core group (group 1 in Phan et al. (2019)) whose members are split between groups 1 and 2 in this study (Table 1). This could be due to collection time of isolates used in that study (up to 2016) and/or the limited number of genetic loci used (32 SSR markers). All three studies agree that the population is characterized by a persistent population (called the core group 1 here) and a number of transient populations, here called groups 2-8 (Tables 2 and 3).

**Figure 2:**
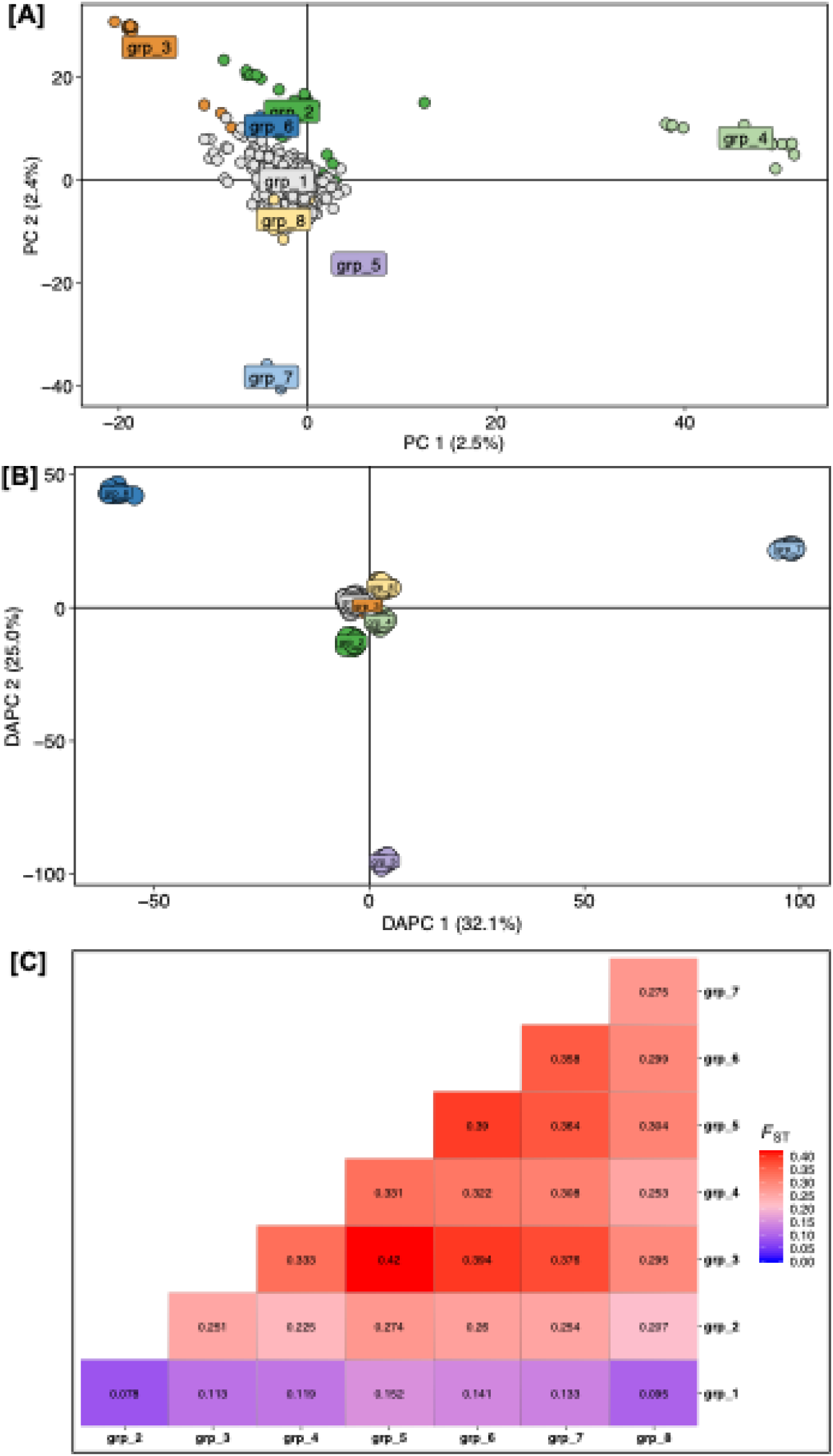
Population structure and genetic diversity analysis of 360 Australian *P. nodorum* isolates. [A] PCA; [B] DAPC; and pairwise Fst of eight groups identified in the collection [C].

**Table 2:**
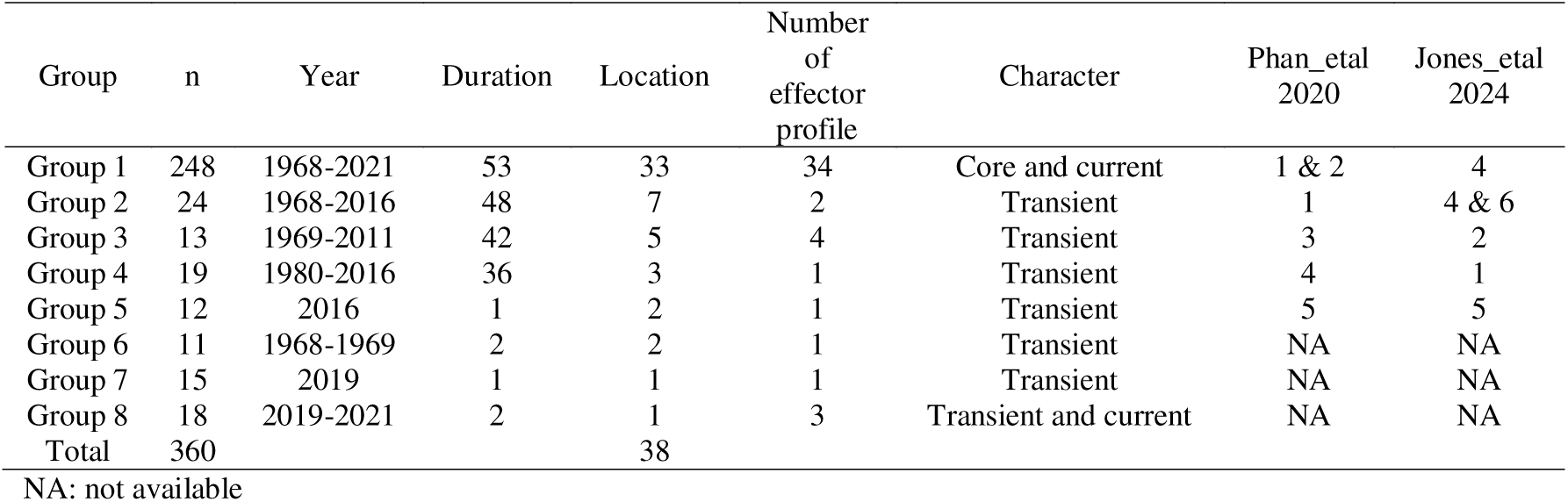
Characteristics of subpopulations found in a collection of 360 Australian *P. nodorum* isolates in current, Phan et al (2020) and Jones et al (2024) studies.

| Group | n | Year | Duration | Location | Number<br>of<br>effector<br>profile | Character | Phan_etal<br>2020 | Jones_etal<br>2024 |
| --- | --- | --- | --- | --- | --- | --- | --- | --- |
| Group 1 | 248 | 1968-2021 | 53 | 33 | 34 | Core and current | 1 & 2 | 4 |
| Group 2 | 24 | 1968-2016 | 48 | 7 | 2 | Transient | 1 | 4 & 6 |
| Group 3 | 13 | 1969-2011 | 42 | 5 | 4 | Transient | 3 | 2 |
| Group 4 | 19 | 1980-2016 | 36 | 3 | 1 | Transient | 4 | 1 |
| Group 5 | 12 | 2016 | 1 | 2 | 1 | Transient | 5 | 5 |
| Group 6 | 11 | 1968-1969 | 2 | 2 | 1 | Transient | NA | NA |
| Group 7 | 15 | 2019 | 1 | 1 | 1 | Transient | NA | NA |
| Group 8 | 18 | 2019-2021 | 2 | 1 | 3 | Transient and current | NA | NA |
| Total | 360 |  |  | 38 |  |  |  |  |

NA: not available

**Table 3:** Basic population statistics for eight groups identified in Australian *P. nodorum* population structure.

|  | Group 1 | Group 2 | Group 3 | Group 4 | Group 5 | Group 6 | Group 7 | Group 8 |
| --- | --- | --- | --- | --- | --- | --- | --- | --- |
| Sample size | 248 | 24 | 13 | 19 | 12 | 11 | 15 | 18 |
| Private alleles | 14712 | 63 | 29 | 38 | 48 | 84 | 218 | 10 |
| Average private alleles | 59.32 | 2.63 | 2.23 | 2.00 | 4.00 | 7.64 | 14.53 | 0.56 |
| Allelic richness | 1.83 | 1.58 | 1.36 | 1.50 | 1.38 | 1.40 | 1.43 | 1.59 |
| rbarD | 0.0012 | 0.0598 | 0.2096 | 0.0125 | 0.0007 | 0.0016 | 0.0003 | 0.0559 |

F_ST_ showed a high genetic differentiation between the seven transient groups which were all more closely related to group 1 (Figure 2C). The genetic distances indicated limited gene flow between transient groups and suggested that each transient group was derived from the core group. Allele richness, also referred to as the mean number of alleles per locus, is another measure of genetic variation. The highest allele richness was found in core group 1, moderate in groups 2, 4, 8 and low in groups 3, 5, 6, 7 which is consistent with their group sizes (Table 3). The average private allele frequency of group 1 is significantly higher than other groups, followed by that of group 7 and lowest is in group 8 (Table 3). This also reflected in the phylogenetic tree, where long branches are commonly found among group 1 isolates (Figure 1).

An UPGMA phylogenetic tree was built using the 18,208 SNP from the 360 genomes (Figure 1) (Kamvar et al. 2015). The tree branches are generally long, indicating a high level of genetic diversification. However, eight lineages with significantly shorter branches are observed comprising seven transient group’s isolates (Figure 1). Group 1 accounted for 69% of the tested isolates. Its members were collected over the entire duration from 1968 to 2021, and at 33 out of 41 sampling locations (Table 2) and were embedded throughout the phylogenetic tree (Figure 1). The transient groups (2-8) contained between 11 to 24 individuals, with collection times ranging from 1-2 years (groups 5, 6, 7, and 8) or 36-48 years (groups 2, 3, and 4) and located in a maximum of 7 wheat growing sites (Table 2). DAPC identified several admixture isolates between group 1 and group 3 (SFigure 6 and STable 5)

### Characteristics of the groups

Mating type was assigned to each isolate based on the presence of either the MAT1-1 or MAT1-2 idiomorphs (Figure 1 and STable 1). The MAT1-1: MAT1-2 ratio was varied among the subpopulations with some comprised only isolates of either MAT1-1 or MAT1-2 (groups 2, 5, 4, 6, 7). In groups 3 (11 MAT1-1: 2 MAT1-2) and 8 (13 MAT1-1 : 5 MAT1-2), the ratio was highly deviated from 1:1. A χ2 test demonstrated that only the core group 1 (121 MAT1-1 : 127 MAT1-2) has balanced mating-type proportions (*P=* 0.703). In addition to the MAT loci, we also used 18,208 other loci to calculate the Index of association values (I_A_) and rbarD (Standardized Index of association) to determine if a population is in linkage equilibrium. This test helps to determine if populations are clonal or sexual. The null hypothesis of the test is alleles observed at different loci are not linked, i.e. the tested population is sexual (Brown, 1980; Kamvar et al. 2014). Using a permutation approach, the analysis showed that no sexual reproduction occurred in any of the 8 sub-populations studied (SFigure 3).

Effector (NE) profiles for each isolate was obtained from their genome sequences using MetaEuk program (Levy Karin et al. 2020) and is presented in Figure 1. Overall, 92.8% of isolates carried a likely active allele of all 5 effectors. Null alleles were detected in 24 isolates and non-functional alleles were detected in only 2 isolates. There are a limited number of isoforms in this large collection with only 2 for *SnTox3*, 3 for *SnToxA*, *SnTox1, SnTox5* and 5 for *SnTox267*. In addition, non-functional versions and the null alleles of *SnTox3* and *SnTox5* were also found (Figure 1, STable 1). There was striking conservation of effector haplotypes for most of the transient groups; they are monomorphic for all members of groups 2, 4, 5, 6 and 7 (Figure 1, Table 2). Group 3 has the highest number of effector profiles per isolate (Table 2). Novel isoforms of *SnTox5* and *SnTox267* were identified and are detailed in STable 1.

### Differences in genetic contents revealed by genome alignments

To shed light on the mechanisms behind the formation of the groups, information on the identity of the DNA sequences which make up the differences between core and transient groups was sought. Genome alignments between two group 1 isolates, with two or three isolates from each transient group were carried out using Mauve v 2.4.0 (Darling et al. 2004). Two types of sequence differences were detected: (1) ubiquitous sequences scattered within syntenic blocks; (2) sequences unique to either the reference or queried genomes. Table 1 summarises 4,524 such loci. They can be grouped into 6 categories depending on whether similar sequences are found in;

1. the reference Australian isolate (SN15; Bertazzoni et al. (2021)),
2. the reference USA isolates (Sn4, SN79 and SN2000; Richards et al. (2018)),
3. all 4 reference isolates,
4. the transposon Molly,
5. other organisms,
6. or no-hit/unknown.

Table 1 shows a clear pattern in the sporadic presence of the transposon Molly in the 8 groups. Molly is a Tc-1 Mariner element discovered by a transposon-trapping process using UK isolates (Hane & Oliver, 2010; Rawson, 2000) (NCBI AJ488502). This transposon sequence was predominantly found in conserved syntenic blocks in the 28 genome alignments, except on two occasions with a unique fragment in isolates 16FG165 and 15FG168 of Group 5. The multiple copy numbers of Molly which were inserted into otherwise perfect syntenic blocks between isolates are evidence of its ability to jump (SFigure 7).

### Long-read genomes as references for Molly characterisation

To facilitate Molly copy-number identification and characterisation, three telomere-to-telomere Molly-carrying genome assemblies from group 1, 5 and 7 isolates (15FG38, 16FG168, PN315) were generated and their statistics are shown in STable 6 and Figure 3. These genomes are of high quality with all scaffolds/chromosomes having one or both telomere-ends. They were built with 50-100 times long-read coverages and a final polishing step with Illumina highly accurate short-reads. BUSCO scores were high (99%, STable 6) making them suitable for TE analysis. In addition, long/short-read reference genomes of SN15 and Sn2000 obtained from NCBI (project: PRJNA632579, PRJNA398070) were also used as part of reference isolates. Identical Molly copy-numbers were detected using either short or long reads of these reference genomes; consistently they are 19, 44 and 36 for 15FG38, 16FG168 and PN315 respectively which validated the methods used (STable 2). To facilitate comparison among these genomes, all chromosome names and their orientation are consistent with the reference genome of SN15, scaffold names are isolate-specific and their numbers are based on length (Figure 3). Identified Molly sites and their characters are depicted in Figure 3 and reported in STable 7 showing 71% of them fell into LRAR (low GC-content regions). The Molly-insertion patterns presented in Figure 3 indicated random and genome-wide propagation of the TE. Application of this read-mapped coverage method to the whole collection uncovered large variation of Molly copy numbers in group 1, but relative conservation in all other groups (Table 4).

**Figure 3:**
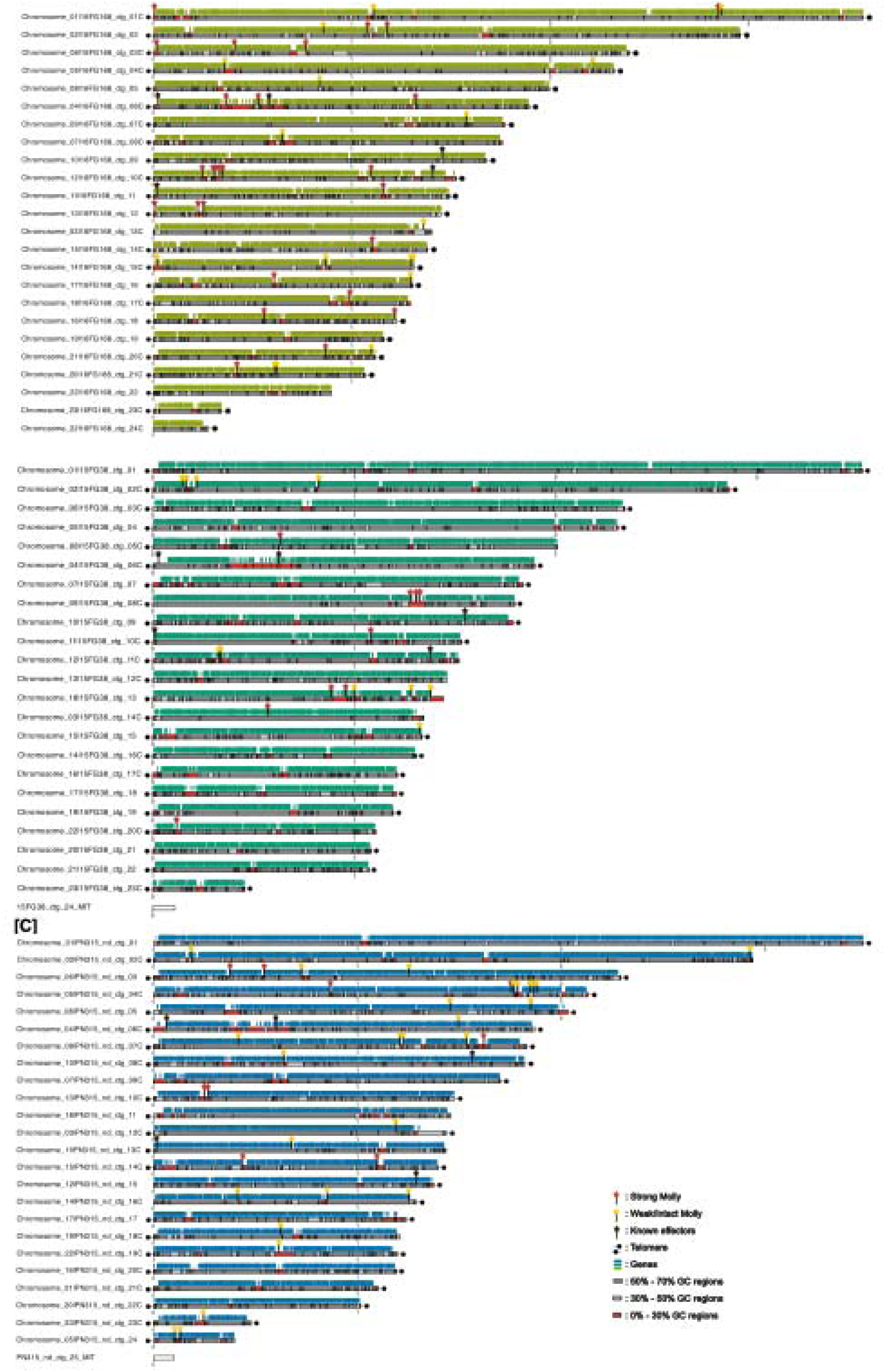
Genomic sites of Molly in three nanopore reference genomes of 15FG38, 16FG168 and PN315. All chromosome names and orientation are consistent with SN15 reference genome. The scaffold names are isolate specific and numbers are based on length. Molly classification and genomic compartmentalisation based on GC contents are indicated in Figure legends.

**Table 4:** Presence of *Stagonospora nodorum* transposon Molly in isolates of eight groups identified in Australian *P. nodorum* population.

| Group | Numbers | Modification |  |
| --- | --- | --- | --- |
|  |  | Degraded*<br>(>25 SNP) | Intact/weak*<br>(0 - 25 SNP) |
| Group 1 | 0 - 45 | 3.08 | 3.65 |
| Group 2 | 0 - 1 | 0.58 | 0.04 |
| Group 3 | 0 - 18 | 1.08 | 1.46 |
| Group 4 | 8 - 10 | 9.47 | 0.11 |
| Group 5 | **41 - 45** | **25.75** | **18** |
| Group 6 | 0 | 0.00 | 0.00 |
| Group 7 | **34 - 42** | **11.40** | **27.07** |
| Group 8 | 1 - 2 | 1.83 | 0.00 |

\*Average Molly copy numbers

The sequence of the intact archetype copy of Molly was also blasted against each of 360 genomes (STable 2). Hits were classified as weak/intact or strong/degraded modifications if there are equal/less or more than 25 SNPs found in a copy, respectively. The separation of intact and degraded Molly help comparison between isolates/groups more meaningful. In addition, as degraded Molly tend to carry mutated and likely misfunctioned transposase gene, this classification indirectly infers active and inactive stages of Molly. From the data set of these 360 genomes, intact copies of Molly were found in some isolates of group 1; when present the copies were more frequently intact than in other groups. Intact and degraded copies were common in all isolates of groups 4, 5 and 7, rare in groups 2, 3 and 8 and absent from group 6 (Table 4, STable 2). The great majority of the alterations in the Molly sequences are characteristic of RIP. RIP-like SNP signatures were found in 91 to 100% of all SNPs identified in Molly sequences of 10 randomly selected Molly-carrying isolates regardless if intact or not with one exception (Figure 4A, STable 8). Together they affect from 14.20% (PN295) to 34.03% (WAC8384) of total Molly sequences (STable 8). AT-rich regions, a signature of RIP, are also evidenced from GC-content plots of the three reference genomes displaying distinct AT-rich and GC-equilibrated region types (Figure 4B & SData 1).

**Figure 4:**
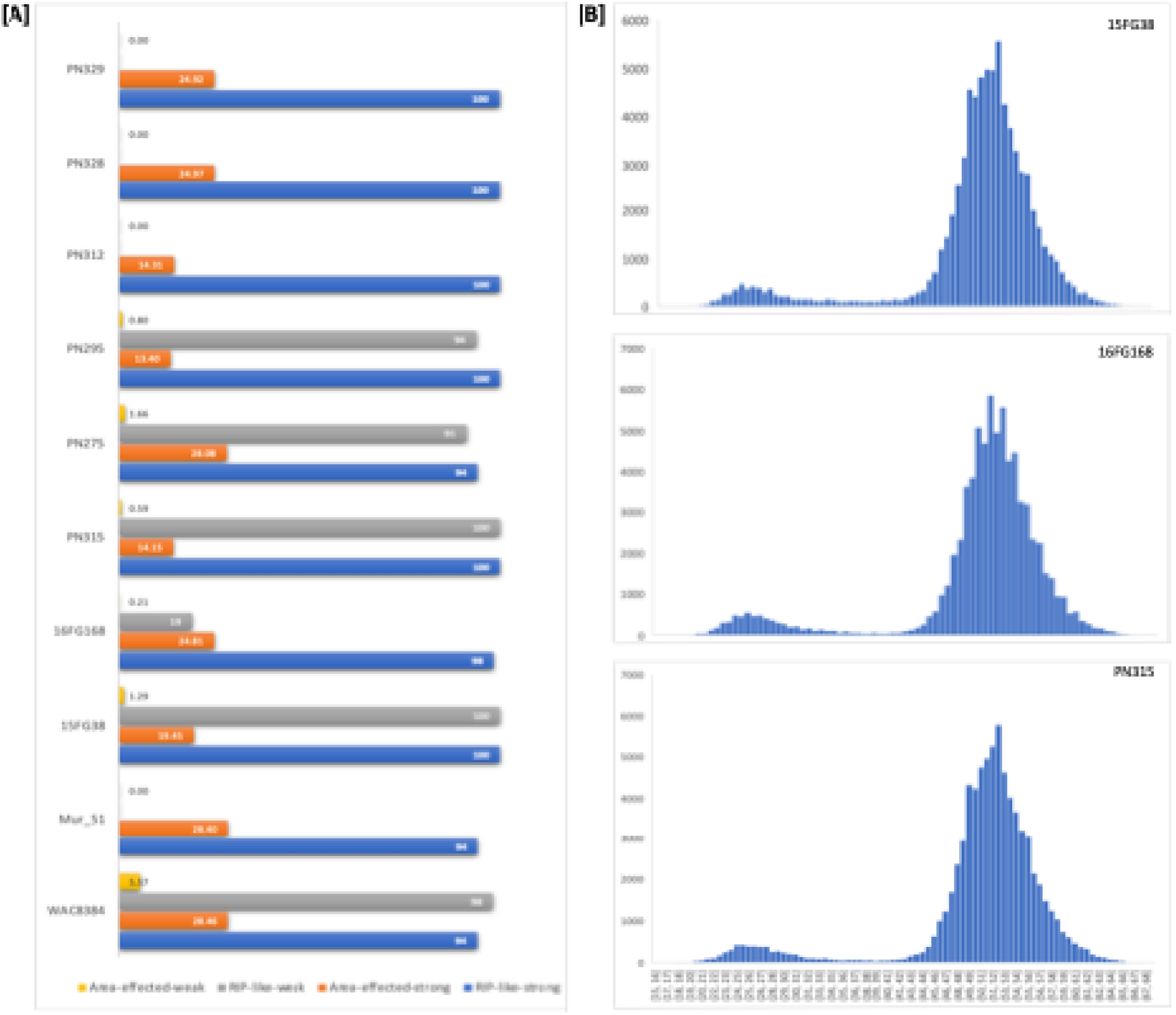
Characterisation of repeat-induced point mutations (RIP) in three Molly-carrying reference genomes. [A]: Percentages of RIP-like SNP and RIP-affected areas in weakly or strongly degraded Molly sequences; [B]: Histogram of GC contents in genomes of three Molly carrying isolates.

The burst of Molly and their abundant signatures of RIP found in the reference genomes suggest they could potentially disturb nearby genes. In 48% of the analysed Molly, the TE have inserted within 1kb distance from a gene including 5 less than 10bp (STable 7). Molly was found to be closed proximity to *SnToxA* in 16FG168 (Figure 3). Of these 99 Molly, putative transposases were only annotated in 54 intact and 26 degraded albeit with different levels of homologies (STable 7).

### Variance in pathogenicity

The previous study identified shifts in the *P. nodorum* population structure which were consistent with adaptations to newly introduced and widely planted wheat cultivars (Phan et al. 2019). To determine if pathogenicity is different among these subpopulations against cultivars released at different time lines/Eras, we selected three isolates from each group (3x8=24) and inoculated them on three modern (Era III) wheat cultivars (Calibre, Mace and Scepter); Wyalkatchem, the dominant cultivar from Era II; and three older cultivars from Era I (Calingiri, Eradu and Halberd). The results are shown in Figure 5 and SFigure 8 where SNB scores were plotted against all transient groups (groups 2-8) in order of their emerging times. Figure 5A showed that Era I cultivars (Halberd, Eradu and Calingiri) (Phan et al. 2019) are significantly different among these pathogen subpopulations and most susceptible to old group 6 isolates. There was also significant difference between these groups against Era II and III cultivars (Figure 5B and 5C) where the highest virulence was exhibited by the more recent transient groups 7 and 8.

**Figure 5:**
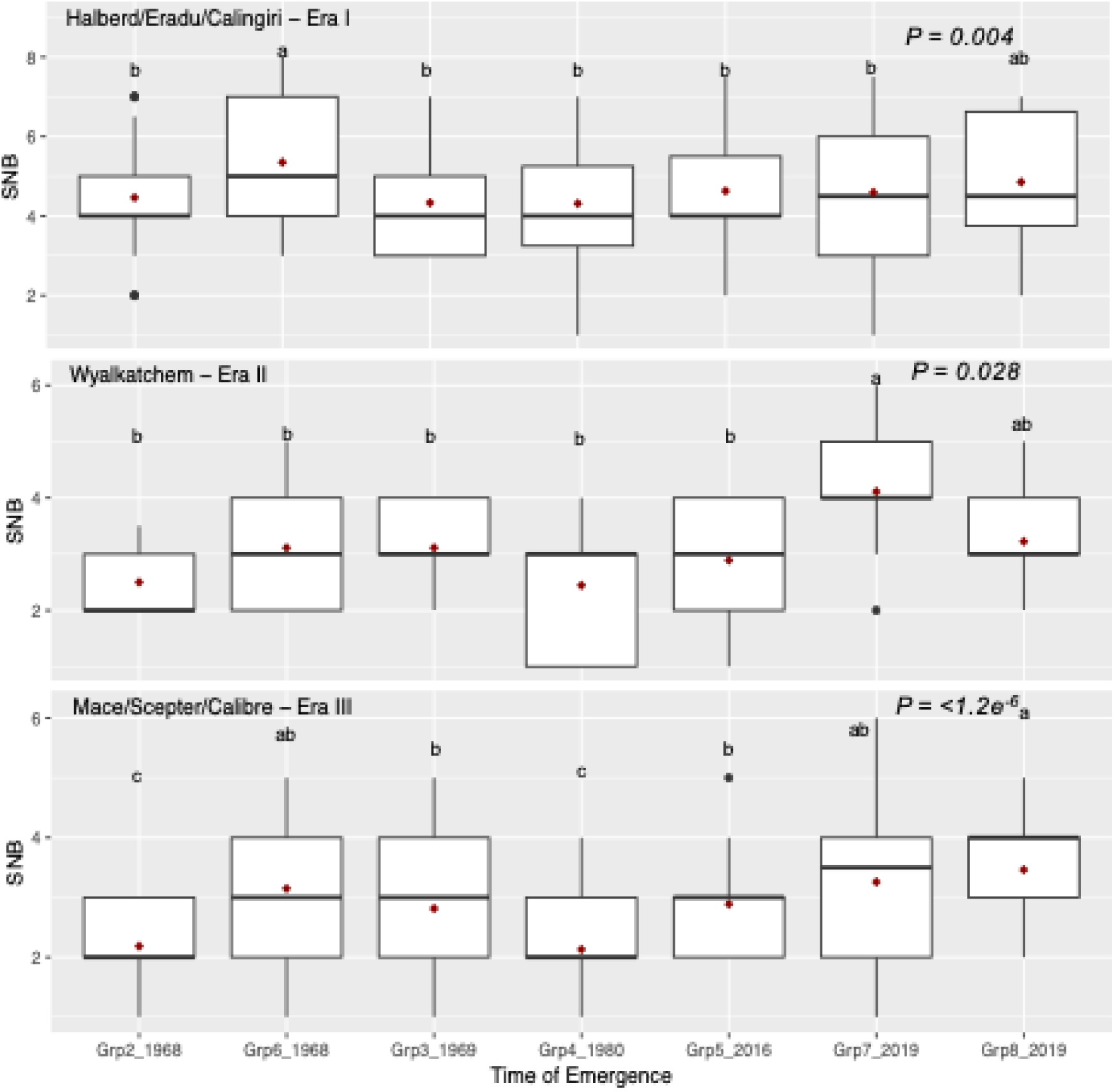
Pathogenicity of WA dominant wheat lines belonging to Era I [A], II [B] and III [C] responding to 7 transient groups identified in WA *P. nodorum* population. These boxplots were based on SNB scores (y axis) and time of emergence (x axis) of the 7 transient groups. Red dots denote means. Different letters represent significant difference.

## Discussion

SNB is a damaging fungal disease of wheat in Australia and parts of USA and Norway (Lin et al. 2020; Richards et al. 2019). In Western Australia, the latest survey estimated that it caused AU$112m p.a. in losses, second amongst all diseases (Murray & Brennan, 2009; Oliver et al. 2009). To effectively control the disease, knowledge about the pathogen’s evolutionary potential is vital. Studies on population genetic diversity help to: 1) understand evolutionary processes at the population and genomic levels; 2) identify important evolutionary factors that shape the pathogen’s population structure; and 3) identify mechanisms which create genomic plasticity enabling adaptation to changing environments (McDonald & Linde, 2002).

In this study, using 360 isolates and 18,208 SNP markers we identified a population structure of eight groups of which one was core, found throughout the collection period and across nearly all collection sites, and seven transient groups. We confirmed the previously identified groups 1 to 5 and found an additional three groups: group 6 was collected in 1968-1969; groups 7 and 8 were recent. The classification of the groups and the differentiation into core and transient was consistent with previous studies as shown in Table 2.

In line with finding from Phan et al. (2019) and Jones et al. (2024), the transient groups appear to be derived from the core group. All transient groups are more closely related to the core group than they are to each others. According to this model, the core population continuously progresses from season to season producing new propagules of diverse genetic composition. Sporadically, a novel population emerges that propagates sufficiently to differentiate itself from the core group but does not spread widely nor continue for more than a few years. Consistent with this model, the transient groups have low intra-group genetic diversity. Most have only a single mating-type idiomorph and a low variance necrotrophic effector profile. Their genic contents are also consistent. Some groups contain large numbers of copies of Molly, some of which are intact and seem to be active (Figure 1 and STable 2).

Our study found the majority (77.2%) of these 360 isolates carrying copies of Molly and a quarter (25.56%) possess active forms. They were identified in isolates collected in 1968 right through to 2021, the entire duration of the collection (STable 2). Evidence from this study showed that Molly can be inserted randomly throughout the pathogen genome, in both genic and non-genic regions (Figure 3 and STable 7). The majority of strongly RIP-modified Molly copies are located in LRAR/AT-rich regions, suggesting the association of this TE and RIP and its effect on host two-speed genome architecture (Figure 3, 4 and STable 7). Given the prevalence of this TE and its ability to activate RIP, we speculate that all genomes without detectable Molly copies previously harboured copies that subsequently were inactivated and hence stabilised but with significantly diversity added. This would explain why each isolate in the main group (group 1) has its own genomic makeup (evidenced by the long branches) which can become their own lineage when opportunity comes as was observed with temporal groups (group 2 to 8).

The presence of the transposon Molly and RIP could potentially explain how some of the transient groups were created. Copies of Molly varied greatly in group 1 isolates but had consistent patterns of low diversity in the other groups. More than 90% of SNP found within the Molly sequences were RIP-like indicating that RIP was active after the bursts of the TE. The evidence of limited genetic diversity and consistent mating, effector and Molly profiles observed within these temporal groups suggests that the emergence of new lineages could come initially from a single isolate which might had differentially proliferated on a dominant cultivar and quickly became a rather diverse sub-population after rounds of invasion and propagations of Molly which in turn triggered RIP.

When these newly emerged groups with active TE reproduce asexually, they could be driven to extinction by an uncontrolled proliferation of TEs as proposed by Dolgin and Charlesworth (2006) which seem to have happened with group 5 and 7 found in this study. No strain belonging to groups 5 or 7 was isolated in the following year from samples collected from the same disease-nursery fields. When these new lineages with active TE reproduced sexually, RIP will be active, mutating intact copies of Molly and thereby stabilising their genomes. Newly created recombinants may not look exactly like either of their original parents but rather a new member in the extensive diverse pool as can be observed with group 1. Although not common, several admixture isolates identified between group 1 and 3 could be such recombinants (SFigure 6 and STable 5). This will add diversity to the genetic pool and explain the closely relatedness between core and temporal groups as shown in Figure 2C. In addition, the emergence/disappearance of other temporal groups could be due to fitness and/or opportunities. The emergence of group 2, 3, 5 and 8 coincides with the mass adoption of one or a few cultivars as can be seen in this and Phan et al. (2019) studies.

The activation of this TE is likely due to environmental stresses. Transposition bursts have been reported to be associated with stresses-induced activation for other TEs in many fungal species (Naito et al 2009; Le et al 2014, Makarevitch et al 2015, Fouché et al 2020). In this study, we have found the two Molly burst events (group 5 in 2016 and group 7 in 2019) that coincided with adoption peaks of two popular wheat cultivars, Mace & Scepter, respectively (SFigure 9).

Forty-eight-percent of the 99 studied Molly was inserted within 1kb distance from a neighbouring-gene, 13% of them have no putative-transposase gene predicted in their sequences, and 6% encode a different hypothetical protein (HBI01_066130, STable 7). These alterations, together with potential RIP leakage into neighbouring regions, showed the physical impact Molly can make to their host genomes which could lead to changes in expression of the near-by genes and ultimately to changes in their phenotypes and possibly in adaptation. Using subset of this collection, Jones et al. (2024) found 33% of SNPs in effector genes are non-synonymous RIP-like. *P. nodorum*’s SnTox1 discovery paper reported a truncated Molly located right next to the effector gene in three out of four isolates tested (Liu et al. 2012). In addition, mounting evidence in other plant pathogens confirms TE regulate effectors, facilitate new effector emergence and diversification leading to variation in virulence (Fouché et al. 2022, Rouxel et al. 2011, Abraham et al. 2024).

To test the pathogenicity of these transient groups, old and modern wheat lines were challenged with groups 2-8. Significant variance in pathogenicity was observed among 7 groups tested on WA wheat cultivars – three old, one intermediate and three modern (Figure 5 and SFigure 8). Halberd, Eradu and Calingiri are old cultivars planted since 1969, 1981 and 1997 respectively and defined Era I (Figure 5). Wyalkatchem was introduced in 2002 and became the dominant cultivar in Era II when it accounted for up to 30% of wheat area. Era III is defined by the massive dominance of Mace (70% in 2016) and its progeny, Scepter and Calibre (Phan et al. 2019). Figure 5 shows the virulence of each transient group against wheats from the three Eras. Wyalkatchem was resistant for the local isolates during 2005 - 2012, since then the pathogen has gradually become more and more aggressive and peaked at 2019 by group 7 on the cultivar. The same pattern was seen with newly dominant wheat Mace, Scepter and Calibre which were resistant to groups 2 to 5, less resistant to group 7 and most susceptible to the current transient group 8 (Figure 5).

This study shows a possible mechanism, Molly proliferation and RIP activation, which can explain *P. nodorum* population structure and temporally formed lineages. The subgroups are positively associated with differential virulence on contemporary cultivars. We do not envisage that individual transposon insertion events caused significant changes in virulence. Rather we imagine that the cumulative effect of transposon movement affected many genes (by ablation and altering expression) that in turn impacted the virulence. Studies have shown that TE and their genomic defence mechanisms can create alteration, deletion or translocation of genes (Oliver, 2012; Richards et al. 2019; Thon et al. 2006; Yoshida et al. 2016); modify expression patterns of adjacent genes (Abraham et al. 2024); provide genetic variation (Badet et al. 2024; Oggenfuss et al. 2021; Raffaele et al. 2010); and contribute to the ‘two-speed’ genome phenomenon (Oliver 2012; Dong et al. 2015; Faino et al. 2016; Testa et al. 2016). Furthermore, mobile elements play a crucial role in diversification of asexually reproducing fungal pathogens, foster their quick adaptation in the arm’s race with their hosts by generating novel and diverse genotypes (Croll & McDonald, 2012; de Jonge et al. 2013; Raffaele & Kamoun, 2012; Rouxel et al. 2011). Directly testing this hypothesis to mimic natural process would be a mammoth task, requiring examination of hundreds of genes interrupted by or near Molly copies. Such a task is well beyond the scope of the current investigation which was focussed on finding mechanism for pathogen diversification and practical solutions for wheat growers and breeders.

### Conclusions

This current project identified potential capacity of Molly in shaping the pathogen population structure and creating genomic diversity. It has revealed that the Australian *P. nodorum* population in any particular field or time consists of isolates from a diverse core and one or two transient more adapted groups. The transient groups follow the ‘boom and bust’ patterns as described before (Phan et al. 2019). This study adds weight to previous recommendations for controlling variable pathogens using genetic rather than chemical means. It is critical to make large annual isolate collections, to characterise them both genotypically and phenotypically, and thereby differentiate new and existing groups. Such studies allow breeders to select the most useful isolates for pathogenicity testing in breeding for both short term resistance (current transient group/s) and sustainable resistance (core). This study also highlights the potential value of cultivar mixes for improving disease control by using cultivars with contrasting reactions to current and core groups which promise a sustainable disease management (Kristoffersen et al. 2020) and the possible benefit of recycling old cultivars which could become resistant to temporal nonadaptive isolates.

## DECLARATIONS

### Ethics approval and consent to participate

Not applicable

### Consent for publication

Not applicable

### Availability of data and materials

Genomic data for DPIRD and current collections are available in NCBI databases, under BioProject ID PRJNA1130627 and PRJNA1128849, respectively. Nanopore data is available under BioProject ID PRJNA1302158.

### Authors’ contributions

HP conceived the experiment. HP and RPO wrote the manuscript. HP, DJ, EF, FK, and KR conducted all experiments. Results were analysed by HP, DJ, MS, HG, EF, FK, KR. All provided intellectual feedback and edited the manuscript. All authors read and approved the manuscript.

## ACKNOWLEDGEMENTS

This study was supported by the Centre for Crop and Disease Management and the Grains Research and Development Corporation (Grant:CUR00023). It was carried out with the assistance of resources and services from the Pawsey-Supercomputing-Centre.

We would like to acknowledge Dr James Hane for his help in obtaining genomic sequences of the DPIRD isolates.

## Supporting Information legends

1. **STable 1**: Metadata of isolates used in this study
2. **STable 2**: *Stagonospora nodorum* transposon Molly profiles of 360 isolates
3. **STable 3**: Effectors sensitivity profiles of 7 wheat lines
4. **STable 4**: QC genome assemblies using Quast
5. **STable 5**: Group asignment of 360 *P. nodorum* isolates by Discriminant Analysis of Principal Components analysis
6. *Stagonospora nodorum* transposon Molly profiles of 360 isolates
7. **STable 6**: Nanopore reference genomes statistics
8. **STable 7**: Molly meta data in three reference genomes
9. **STable 8**: GC-content and SNP found in Molly
10. **SFigure 1**: Locations [A]; numbers and times [B] of isolate collections used in this study.
11. **SFigure 2**: BUSCO outputs for genome assemblies of DPIRD and current isolates. Isolate indexes were used as listed in STable 1
12. **SFigure 3**: Permutation tests for Standardised Index of Association (rbarD) for each of 8 groups found in 360 Australian *P. nodorum* isolate collection
13. **SFigure 4**: QC for sequencing reads of DPIRD and current isolates. Isolate indexes were used as listed in STable 1
14. **SFigure 5**: STRUCTURE ka value [A] and cross validation in DAPC analysis [B] of 360 Australian *P. nodorum* collection
15. **SFigure 6**: Haplotype plot of 360 isolates as identified by Discriminant Analysis of Principal Components. Black arrows indicate admixture strains. Isolate order (from left to right) was the same as in STable 5
16. **SFigure 7**: Sequence alignment of *Stagonospora nodorum* transposon Molly found in three randomly chosen isolates. [A]: Northam-WGT/PN141–Grp1 vs 16GH168/PN155–Grp5; [B]: 15FG38/PN125–Grp1 vs 16FG168/PN155–Grp5; [C]: 15FG38/PN125-Grp1 vs PN343–Grp7.
17. **SFigure 8**: SNB of 7 wheat lines by years and cultivars
18. **SFigure 9**: Top 10 most grown wheat cultivars in Western Australia from 2015 to 2024. Data was obtained from DPIRD (Department of Primary Industries and Regional Development) as “Crop Sowing Guides” (https://grdc.com.au/resources-and-publications/all-publications/nvtcrop-sowing-guides/wa-crop-sowing-guide). Y- axis: percent of total wheat grown area; x-axis: Years; line-colour indicates wheat cultivars as in SFigure9 legend.
19. **SData**: RIPper analysis of 3 reference genomes

