## Supplementary_Figures for "Transposon-associated genetic structure of a fungal phytopathogen population of wheat"

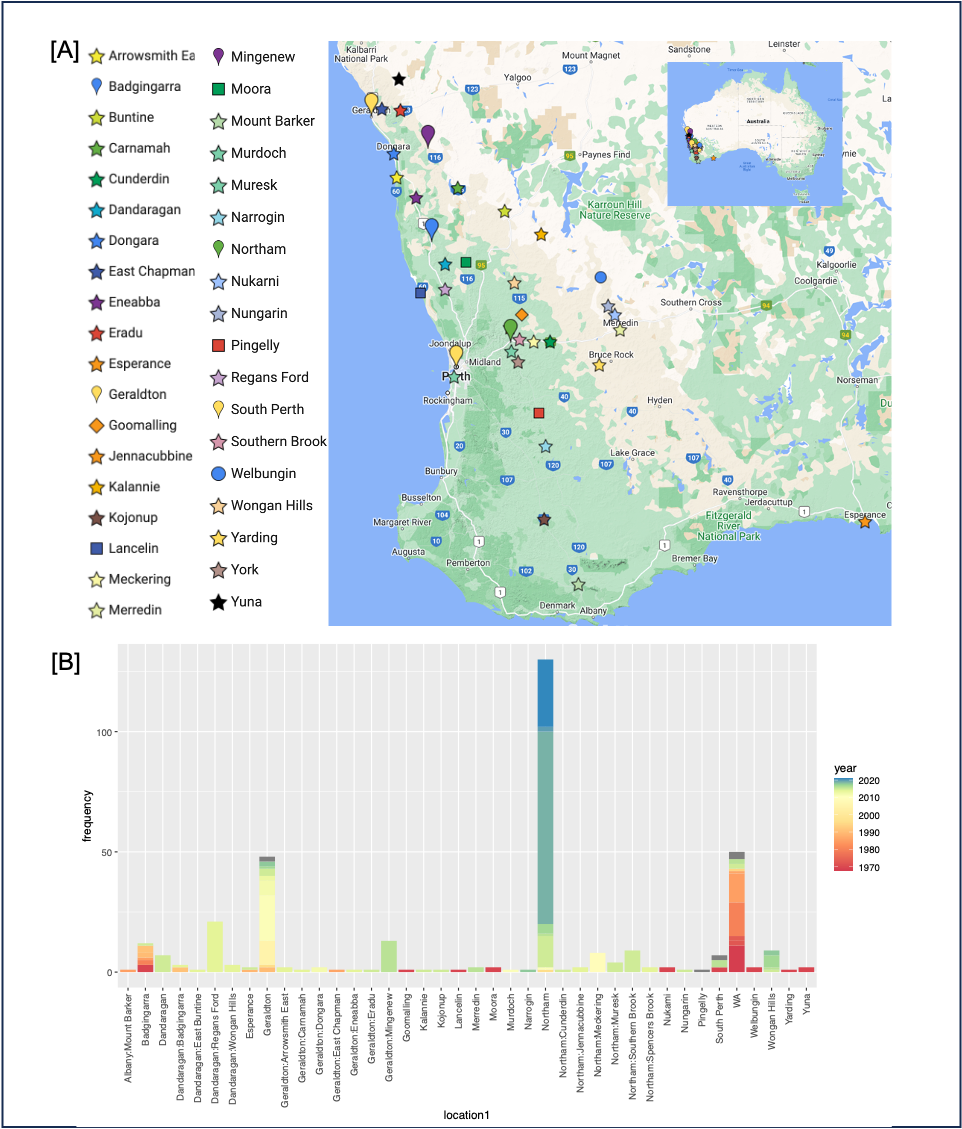

**SFigure 1**: Locations [A]; numbers and times [B] of isolate collections used in this study.

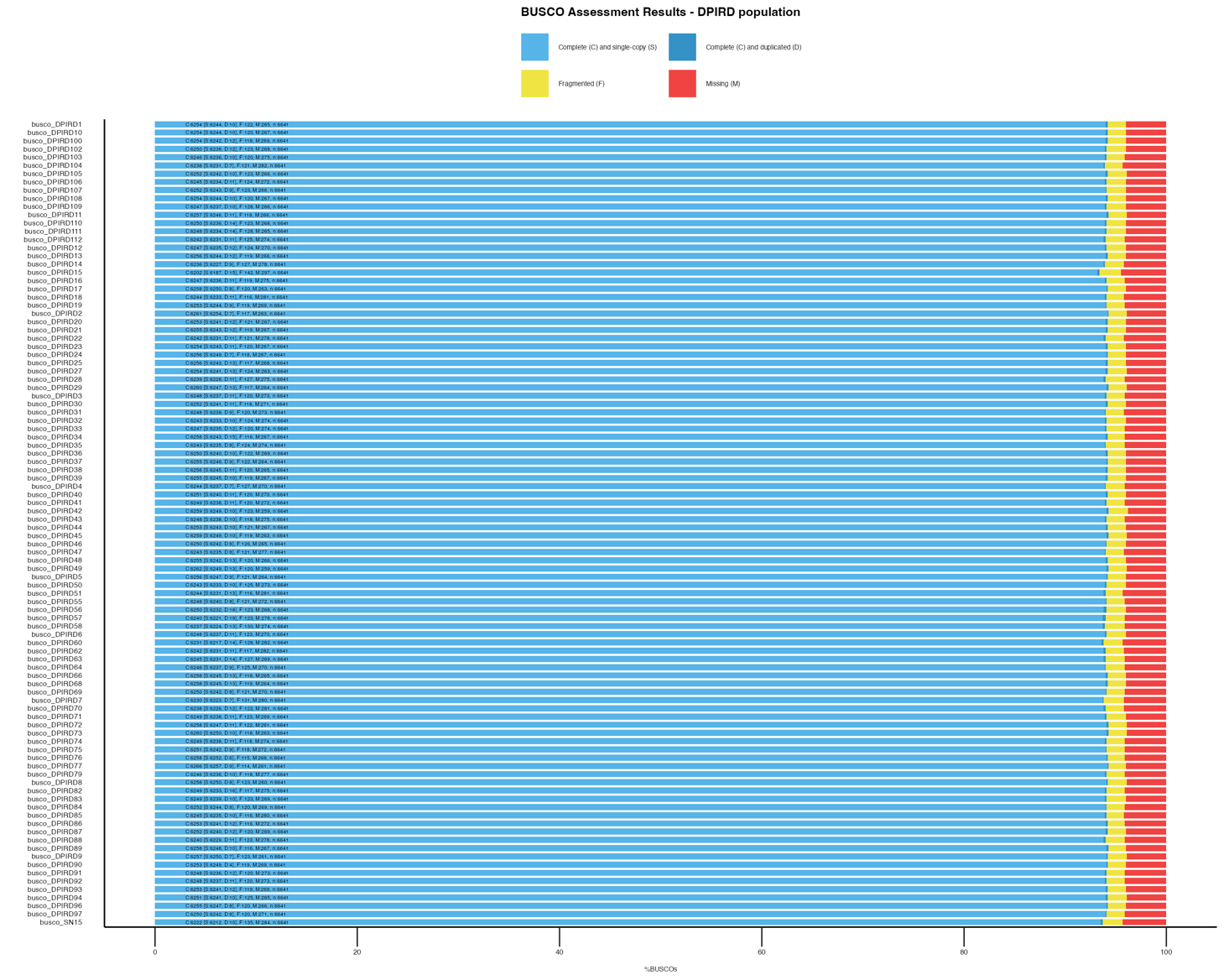

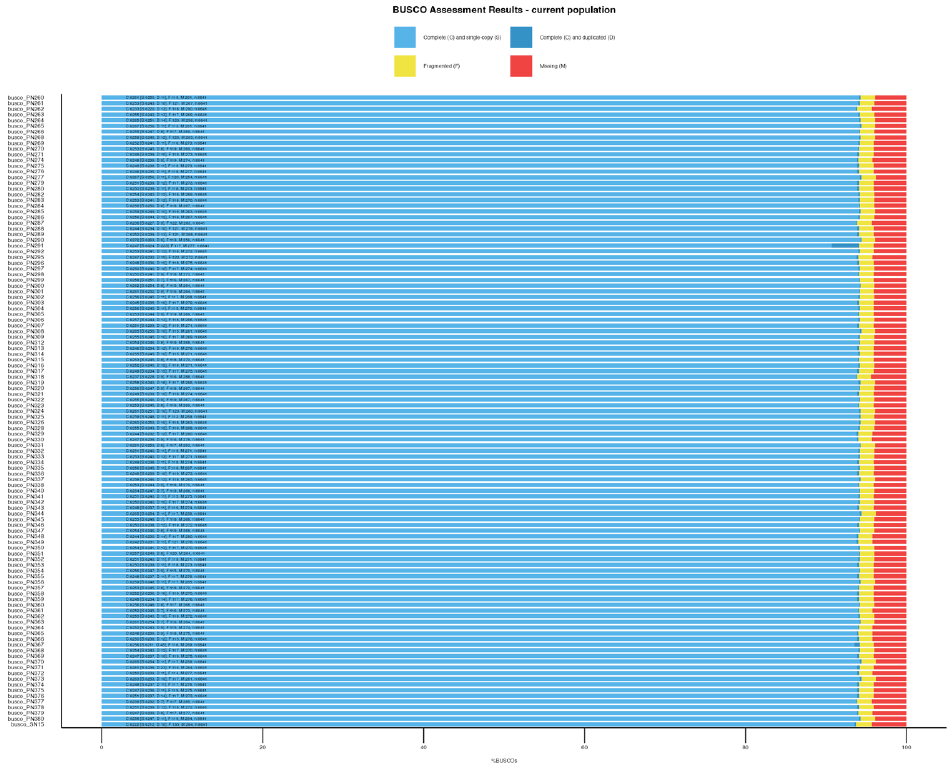

**SFigure 2**: BUSCO outputs for genome assemblies of DPIRD and current isolates. Isolate indexes were used as listed in STable 1

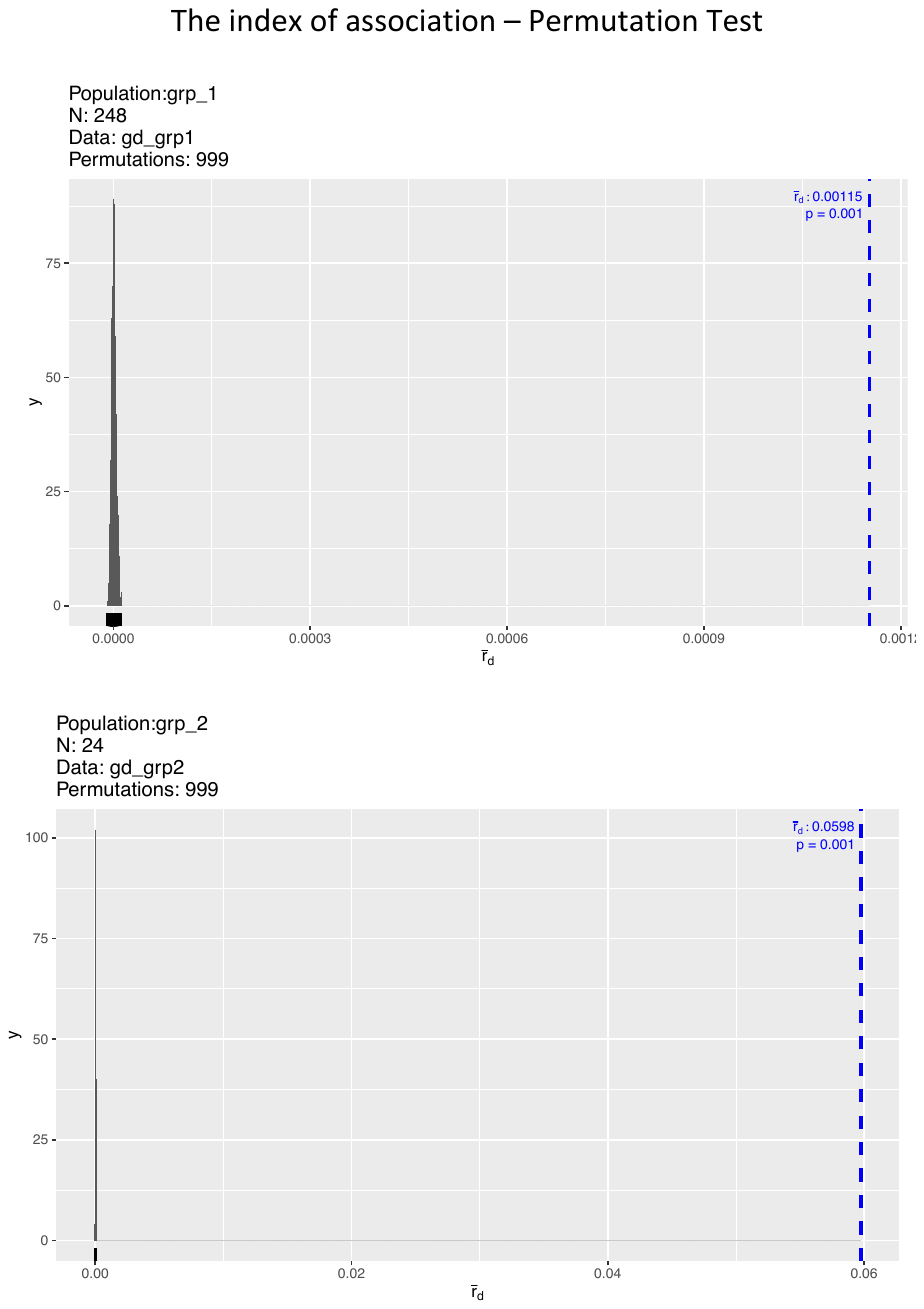

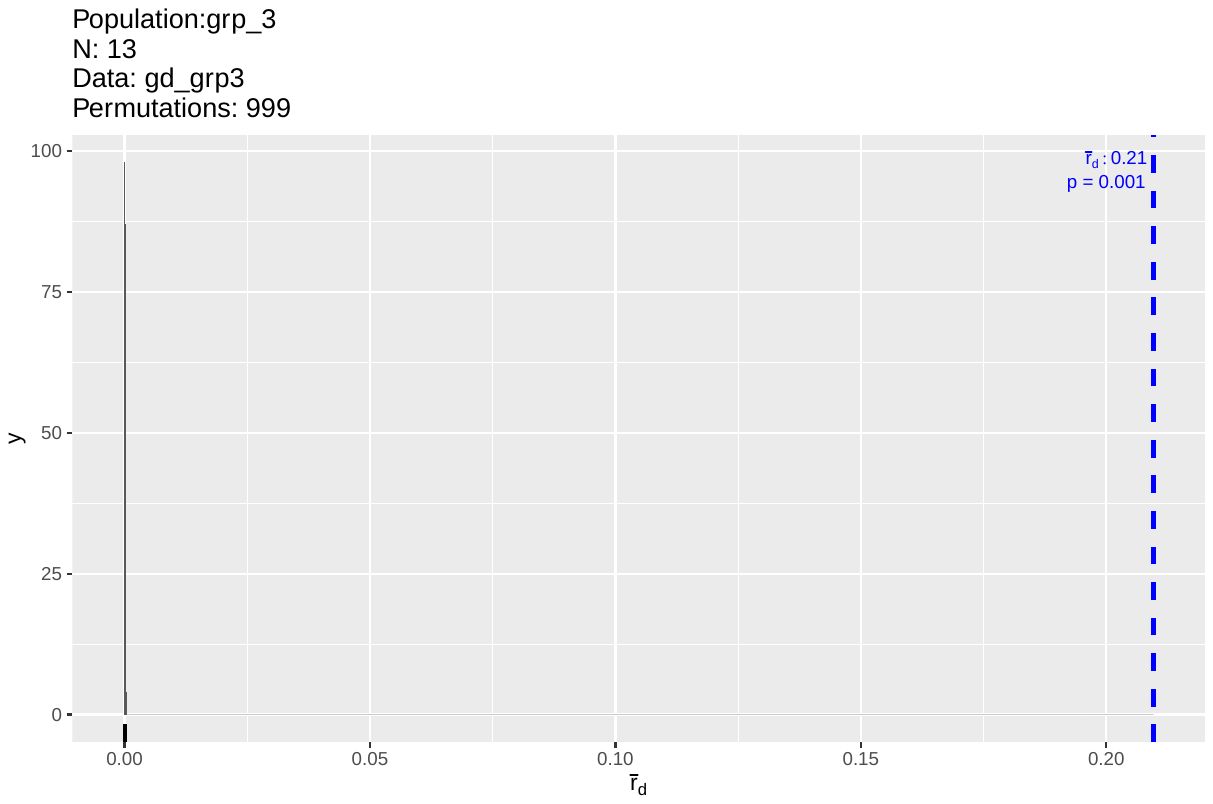

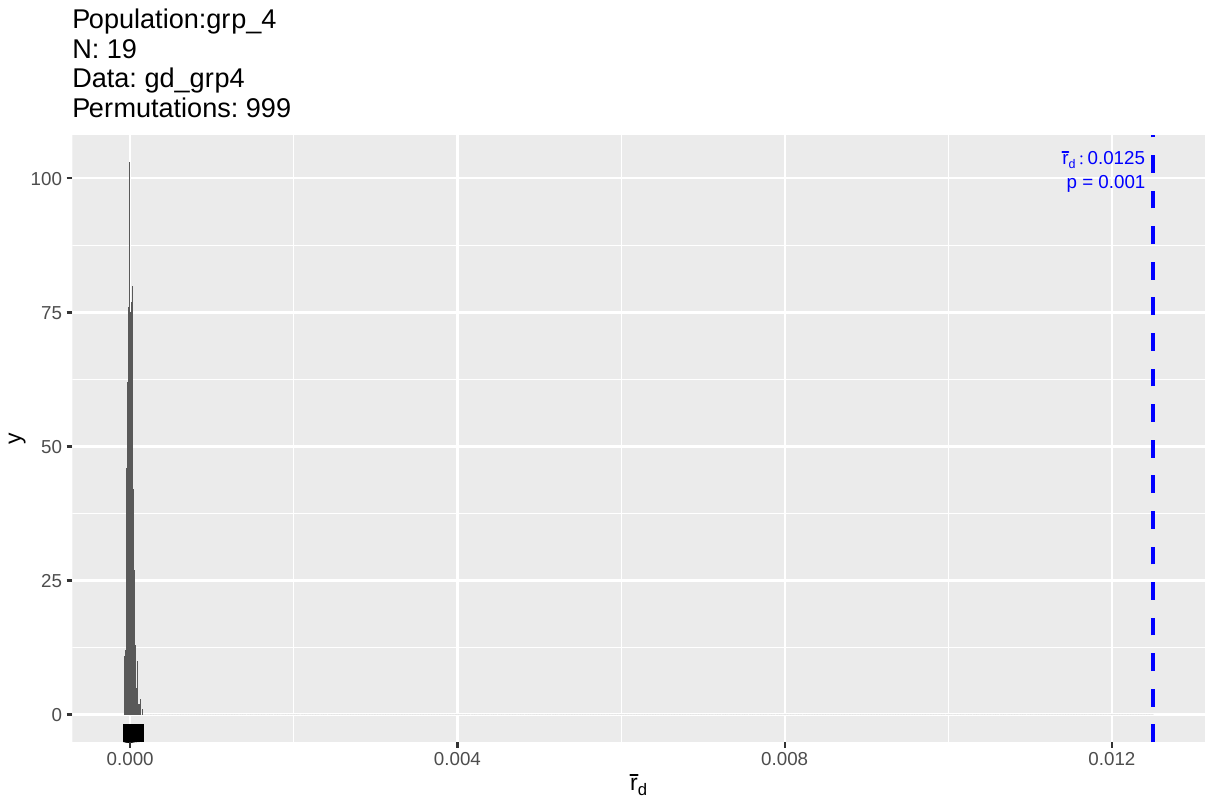

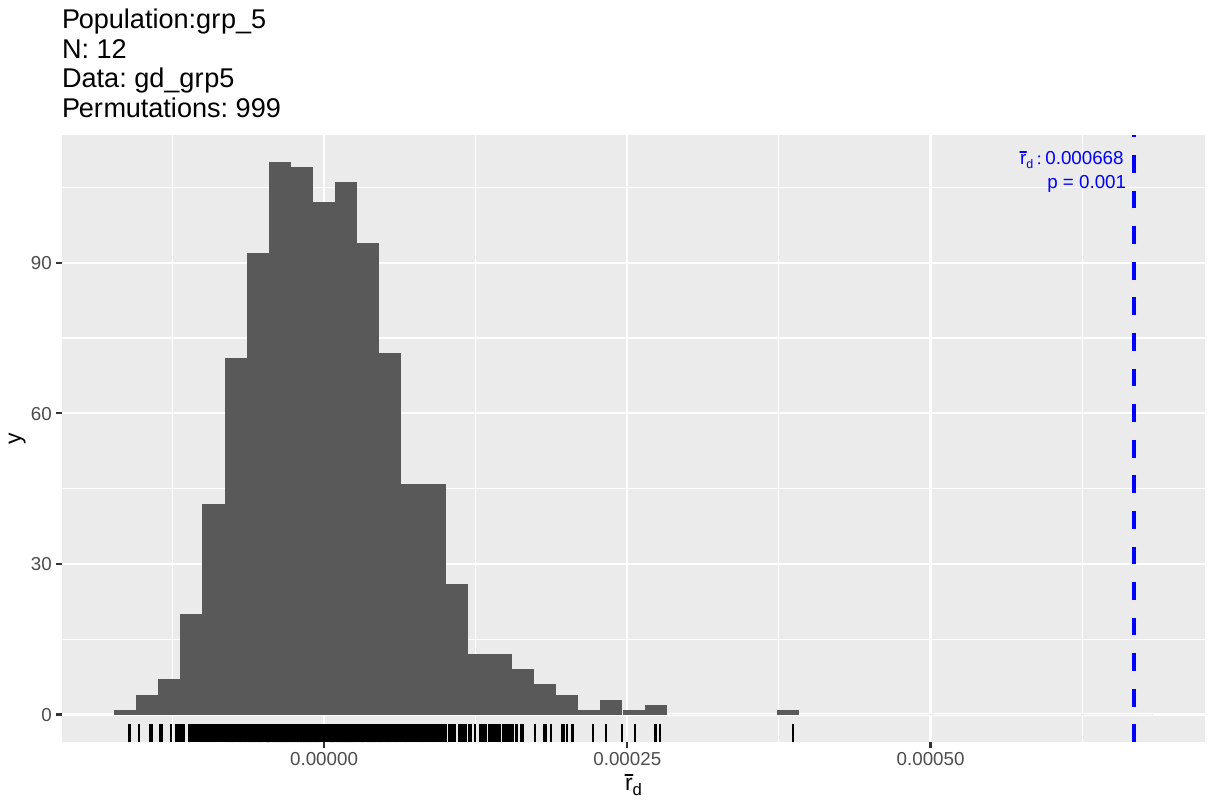

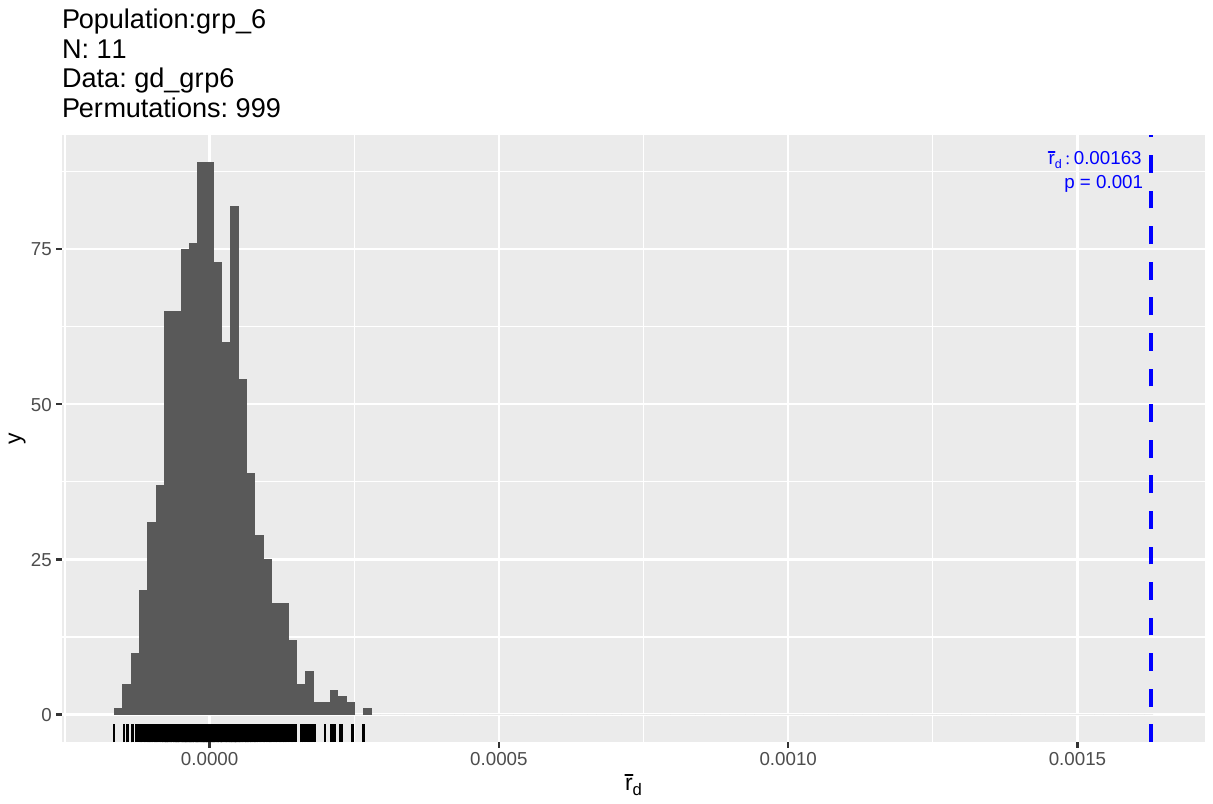

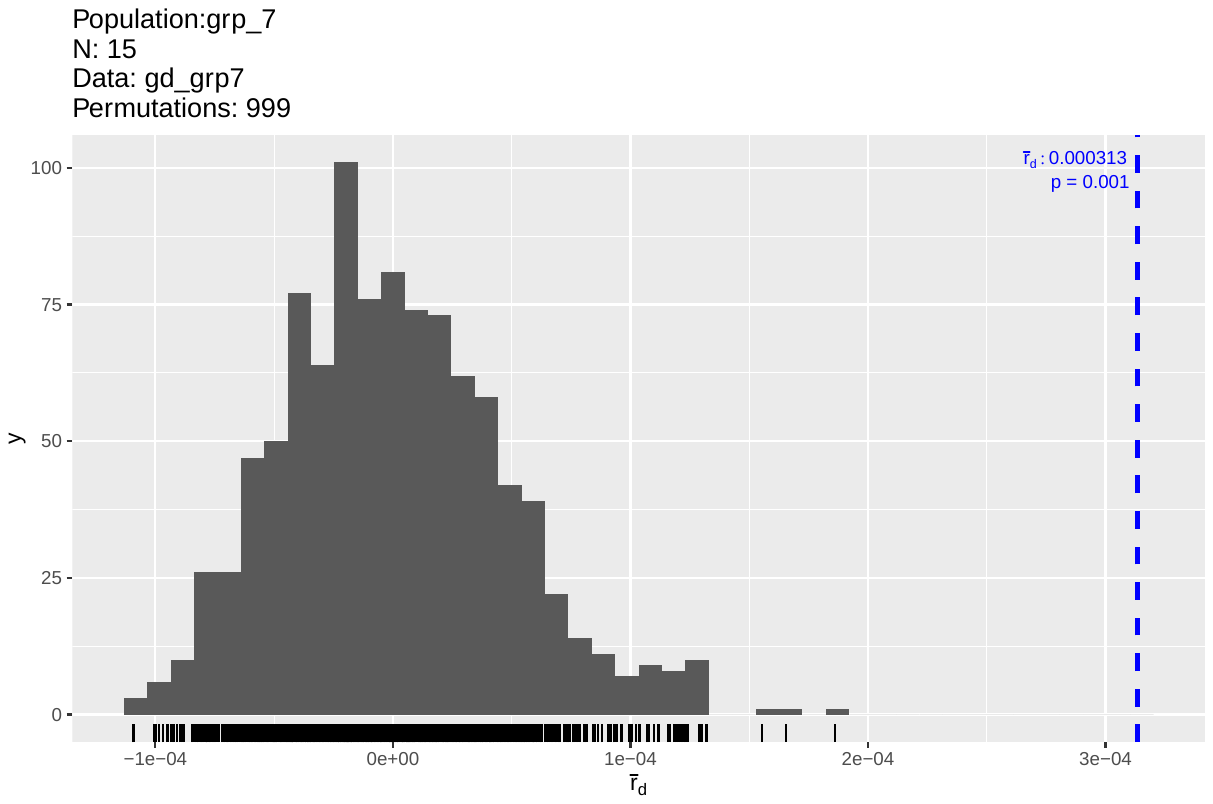

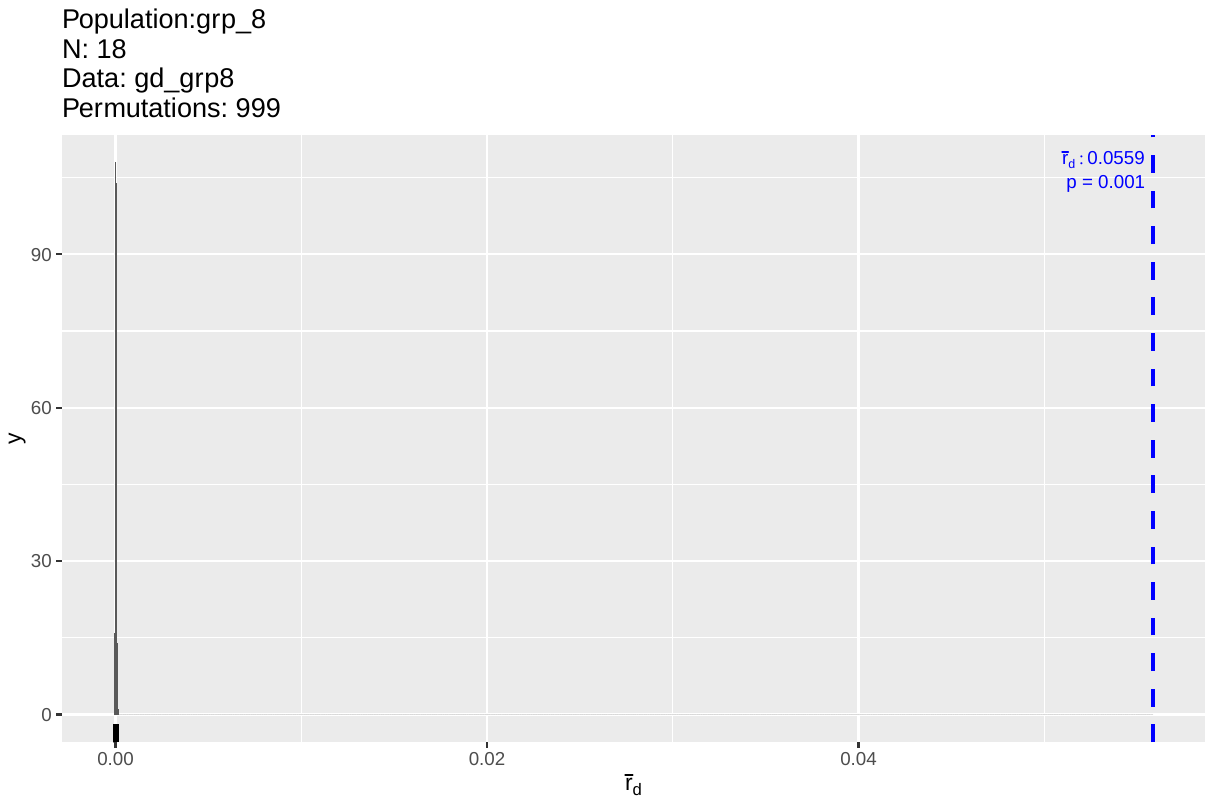

**SFigure 3**: Permutation tests for Standardised Index of Association (rbarD) for each of 8 groups found in 360 Australian *P. nodorum* isolate collection

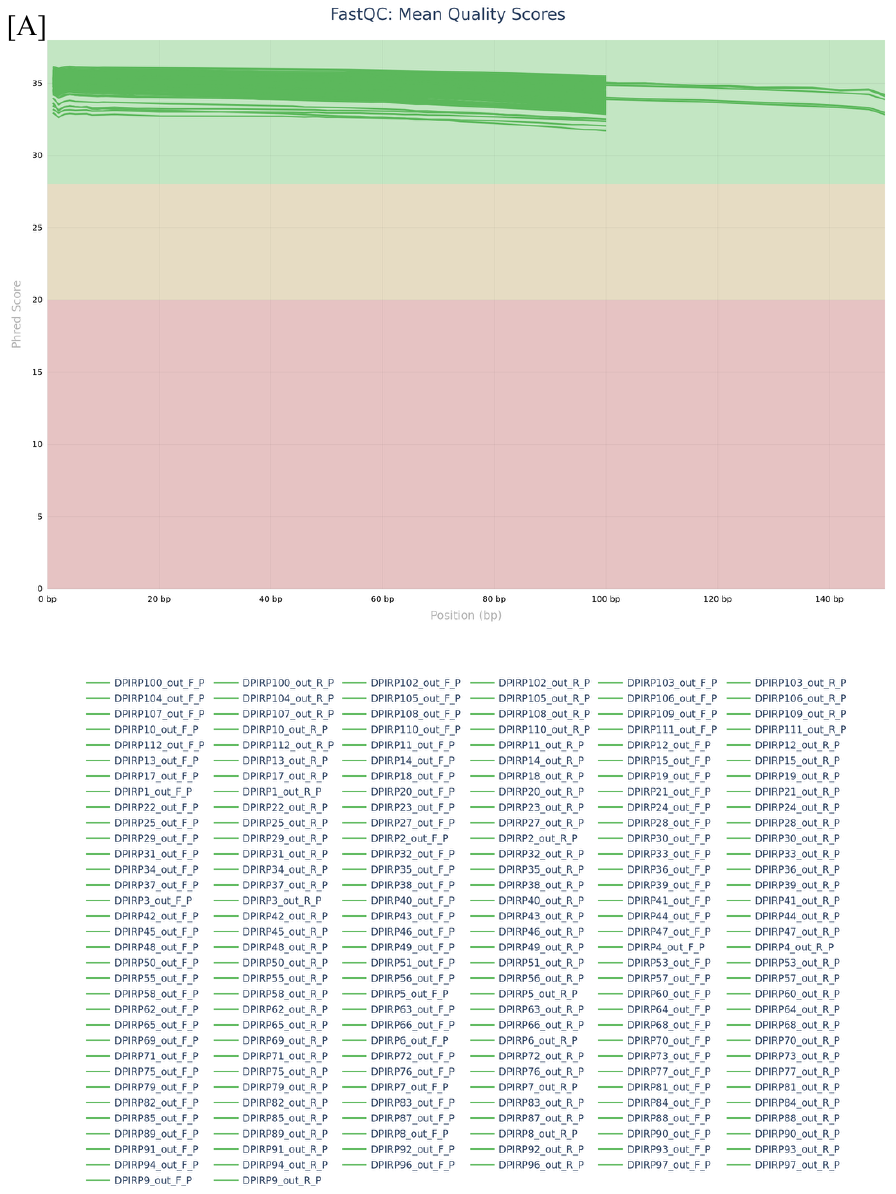

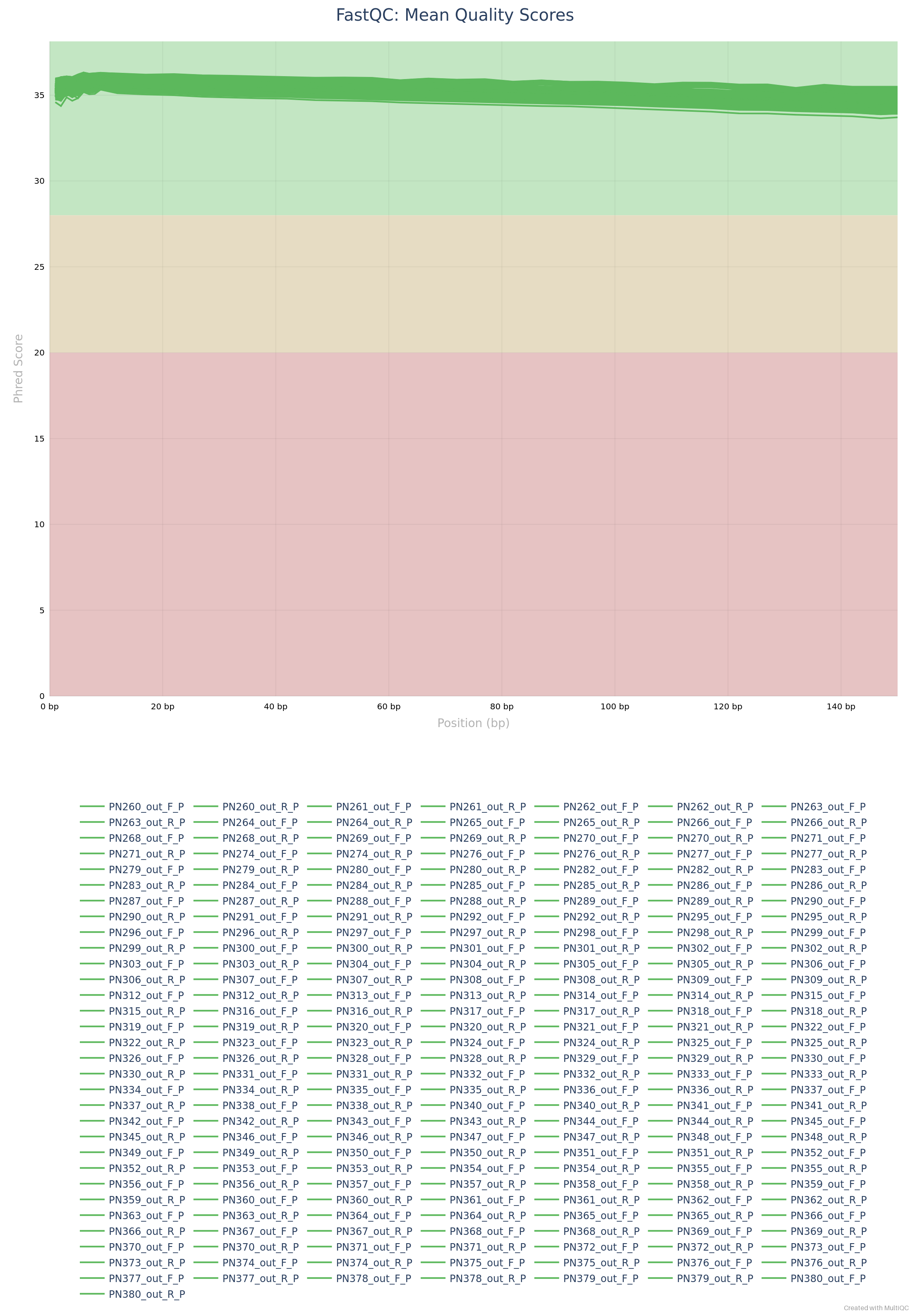

**SFigure 4**: QC for sequencing reads of DPIRD and current isolates. Isolate indexes were used as listed in STable 1

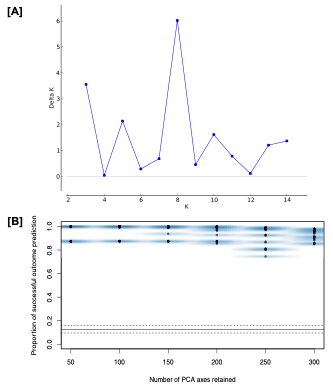

**SFigure 5**: STRUCTURE ka value [A] and cross validation in DAPC analysis [B] of 360 Australian *P. nodorum* collection

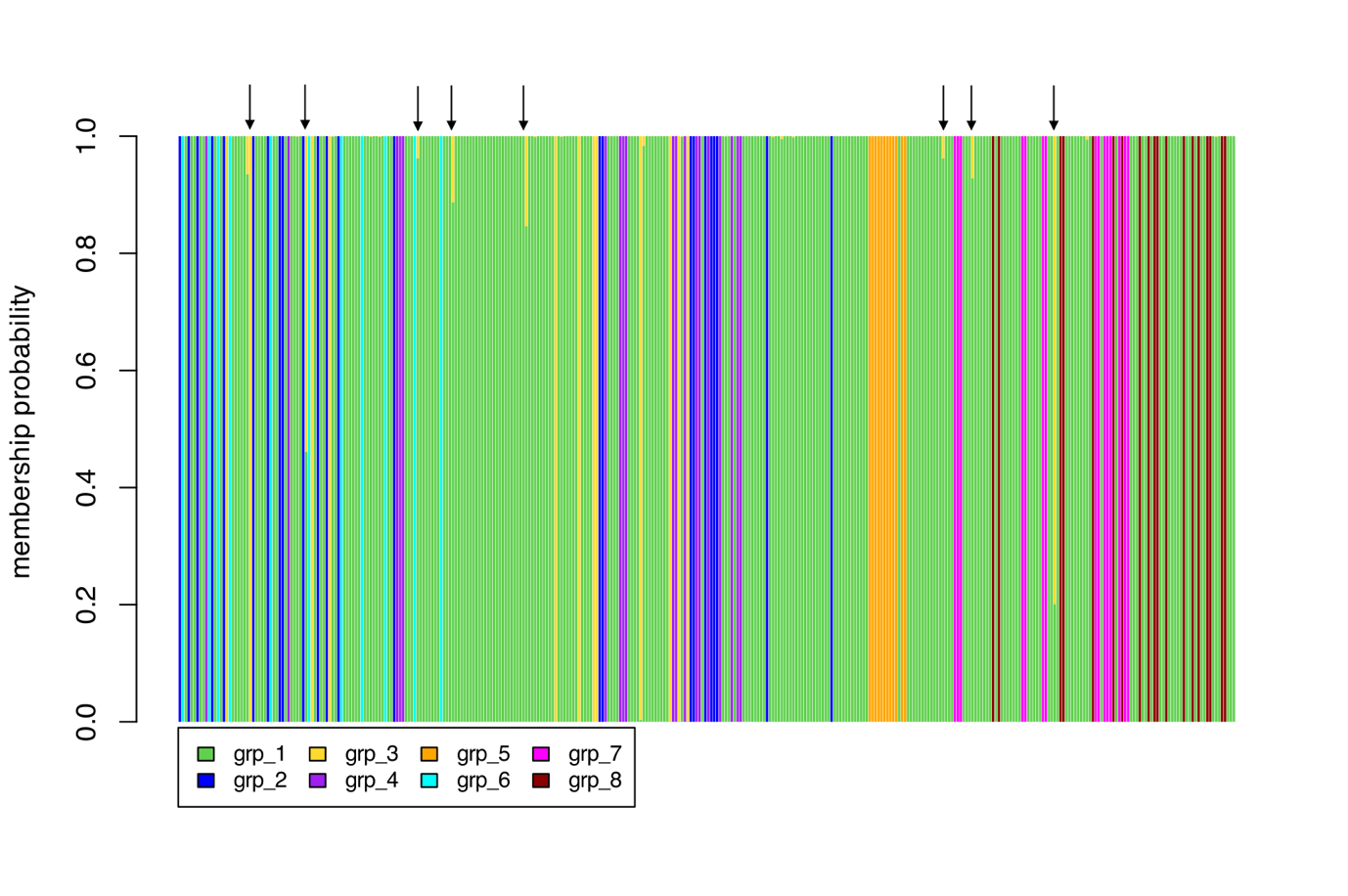

**SFigure 6**: Haplotype plot of 360 isolates as identified by Discriminant Analysis of Principal Components. Black arrows indicate admixture strains. Isolate order (from left to right) was the same as in STable 5

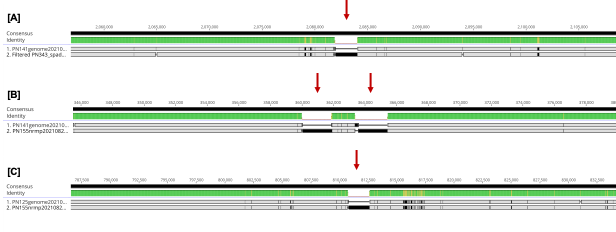

**SFigure 7**: Sequence alignment of *Stagonospora nodorum* transposon Molly found in three randomly chosen isolates. [A]: Northam-WGT/PN141–Grp1 vs 16GH168/PN155–Grp5; [B]: 15FG38/PN125–Grp1 vs 16FG168/PN155–Grp5; [C]: 15FG38/PN125-Grp1 vs PN343–Grp7.

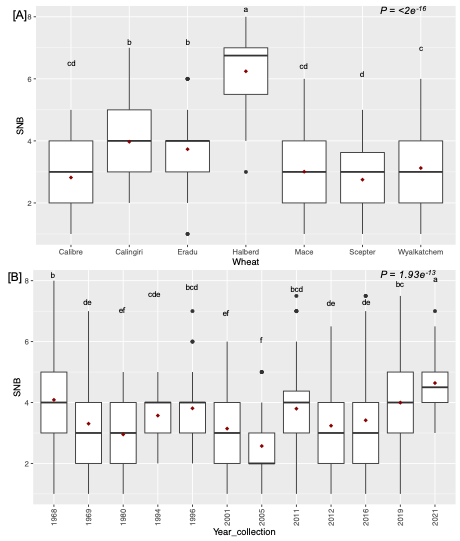

**SFigure 8**: SNB of 7 wheat lines by years and cultivars

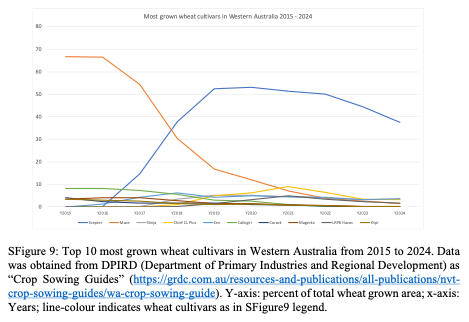

**SFigure 9**: Top 10 most grown wheat cultivars in Western Australia from 2015 to 2024. Data was obtained from DPIRD (Department of Primary Industries and Regional Development) as “Crop Sowing Guides” (https://grdc.com.au/resources-and-publications/all-publications/nvtcrop-sowing-guides/wa-crop-sowing-guide). Y-axis: percent of total wheat grown area; x-axis: Years; line-colour indicates wheat cultivars as in SFigure9 legend.
