## Supplementary_Tables for "Transposon-associated genetic structure of a fungal phytopathogen population of wheat"

**STable 1**: Metadata of isolates used in this study

| # | Original_name | Isolate_index | Phylogenetic_tree_name | Collection | Location | Year | Source | pop_phan20 | op_Jones2 | Group |
|---|---|---|---|---|---|---|---|---|---|---|
| 1 | WAC1141 | DPIRD1 | DIP1_WAC1141 | DPIRD | WA | 1968 | This study | ND | ND | grp_2 |
| 2 | WAC1201 | DPIRD10 | DIP10_WAC1201 | DPIRD | WA | 1968 | This study | ND | ND | grp_6 |
| 3 | WAC14144 | DPIRD100 | DIP100_WAC14144 | DPIRD | Wongan Hills | 2018 | This study | ND | ND | grp_1 |
| 4 | WAC2816 | DPIRD102 | DIP102_WAC2816 | DPIRD | WA | 1980 | This study | ND | ND | grp_2 |
| 5 | WAC2812 | DPIRD103 | DIP103_WAC2812 | DPIRD | WA | 1980 | This study | ND | ND | grp_1 |
| 6 | WAC8397 | DPIRD104 | DIP104_WAC8397 | DPIRD | Badgingarra | 1991 | This study | ND | ND | grp_1 |
| 7 | WAC2804 | DPIRD105 | DIP105_WAC2804 | DPIRD | WA | 1980 | This study | ND | ND | grp_2 |
| 8 | WAC1544 | DPIRD106 | DIP106_WAC1544 | DPIRD | South Perth | 1969 | This study | ND | ND | grp_1 |
| 9 | WAC2799 | DPIRD107 | DIP107_WAC2799 | DPIRD | WA | 1980 | This study | ND | ND | grp_1 |
| 10 | WAC4312 | DPIRD108 | DIP108_WAC4312 | DPIRD | WA | 1985 | This study | ND | ND | grp_4 |
| 11 | WAC1198 | DPIRD109 | DIP109_WAC1198 | DPIRD | WA | 1968 | This study | ND | ND | grp_6 |
| 12 | WAC1494 | DPIRD11 | DIP11_WAC1494 | DPIRD | Pingelly | 1969 | This study | ND | ND | grp_2 |
| 13 | WAC8380 | DPIRD110 | DIP110_WAC8380 | DPIRD | Esperance | 1989 | This study | ND | ND | grp_1 |
| 14 | WAC1194 | DPIRD111 | DIP111_WAC1194 | DPIRD | WA | 1968 | This study | ND | ND | grp_6 |
| 15 | WAC4307 | DPIRD112 | DIP112_WAC4307 | DPIRD | WA | 1985 | This study | ND | ND | grp_1 |
| 16 | WAC1495 | DPIRD12 | DIP12_WAC1495 | DPIRD | Moora | 1969 | This study | ND | ND | grp_2 |
| 17 | WAC1496 | DPIRD13 | DIP13_WAC1496 | DPIRD | Badgingarra | 1969 | This study | ND | ND | grp_3 |
| 18 | WAC1497 | DPIRD14 | DIP14_WAC1497 | DPIRD | Welbungin | 1969 | This study | ND | ND | grp_6 |
| 19 | WAC1498 | DPIRD15 | DIP15_WAC1498 | DPIRD | Yuna | 1969 | This study | ND | ND | grp_1 |
| 20 | WAC1500 | DPIRD16 | DIP16_WAC1500 | DPIRD | South Perth | 1969 | This study | ND | ND | grp_1 |
| 21 | WAC1508 | DPIRD17 | DIP17_WAC1508 | DPIRD | Nukami | 1969 | This study | ND | ND | grp_1 |
| 22 | WAC1545 | DPIRD18 | DIP18_WAC1545 | DPIRD | South Perth | 1969 | This study | ND | ND | grp_1 |
| 23 | WAC1546 | DPIRD19 | DIP19_WAC1546 | DPIRD | Nukami | 1969 | This study | ND | ND | grp_1 |
| 24 | WAC1178 | DPIRD2 | DIP2_WAC1178 | DPIRD | WA | 1968 | This study | ND | ND | grp_1 |
| 25 | WAC1547 | DPIRD20 | DIP20_WAC1547 | DPIRD | Goomalling | 1969 | This study | ND | ND | grp_3 |
| 26 | WAC1548 | DPIRD21 | DIP21_WAC1548 | DPIRD | Lancelin | 1969 | This study | ND | ND | grp_2 |
| 27 | WAC1549 | DPIRD22 | DIP22_WAC1549 | DPIRD | Badgingarra | 1969 | This study | ND | ND | grp_1 |
| 28 | WAC1550 | DPIRD23 | DIP23_WAC1550 | DPIRD | Yarding | 1969 | This study | ND | ND | grp_1 |
| 29 | WAC1551 | DPIRD24 | DIP24_WAC1551 | DPIRD | Yuna | 1969 | This study | ND | ND | grp_1 |
| 30 | WAC1552 | DPIRD25 | DIP25_WAC1552 | DPIRD | Badgingarra | 1969 | This study | ND | ND | grp_1 |
| 31 | WAC1564 | DPIRD27 | DIP27_WAC1564 | DPIRD | Moora | 1969 | This study | ND | ND | grp_2 |
| 32 | WAC1565 | DPIRD28 | DIP28_WAC1565 | DPIRD | Welbungin | 1969 | This study | ND | ND | grp_6 |
| 33 | WAC2217 | DPIRD29 | DIP29_WAC2217 | DPIRD | WA | 1972 | This study | ND | ND | grp_1 |
| 34 | WAC1179 | DPIRD3 | DIP3_WAC1179 | DPIRD | WA | 1968 | This study | ND | ND | grp_1 |
| 35 | WAC2284 | DPIRD30 | DIP30_WAC2284 | DPIRD | WA | 1973 | This study | ND | ND | grp_2 |
| 36 | WAC2285 | DPIRD31 | DIP31_WAC2285 | DPIRD | WA | 1980 | This study | 1 | 4 | grp_2 |
| 37 | WAC2286 | DPIRD32 | DIP32_WAC2286 | DPIRD | WA | 1980 | This study | ND | ND | grp_1 |
| 38 | WAC2796 | DPIRD33 | DIP33_WAC2796 | DPIRD | WA | 1980 | This study | ND | ND | grp_4 |
| 39 | WAC2797 | DPIRD34 | DIP34_WAC2797 | DPIRD | WA | 1980 | This study | ND | ND | grp_1 |
| 40 | WAC2798 | DPIRD35 | DIP35_WAC2798 | DPIRD | WA | 1980 | This study | ND | ND | grp_1 |
| 41 | WAC2802 | DPIRD36 | DIP36_WAC2802 | DPIRD | WA | 1980 | This study | ND | ND | grp_1 |
| 42 | WAC2803 | DPIRD37 | DIP37_WAC2803 | DPIRD | WA | 1980 | This study | ND | ND | grp_1 |
| 43 | WAC2805 | DPIRD38 | DIP38_WAC2805 | DPIRD | WA | 1980 | This study | ND | ND | grp_2 |
| 44 | WAC2806 | DPIRD39 | DIP39_WAC2806 | DPIRD | WA | 1980 | This study | ND | ND | grp_3 |
| 45 | WAC1188 | DPIRD4 | DIP4_WAC1188 | DPIRD | WA | 1968 | This study | ND | ND | grp_6 |
| 46 | WAC2808 | DPIRD40 | DIP40_WAC2808 | DPIRD | Badgingarra | 1980 | This study | ND | ND | grp_3 |
| 47 | WAC2809 | DPIRD41 | DIP41_WAC2809 | DPIRD | WA | 1980 | This study | ND | ND | grp_1 |
| 48 | WAC2817 | DPIRD42 | DIP42_WAC2817 | DPIRD | Badgingarra | 1980 | This study | ND | ND | grp_2 |
| 49 | WAC4292 | DPIRD43 | DIP43_WAC4292 | DPIRD | WA | 1985 | This study | ND | ND | grp_1 |
| 50 | WAC4301 | DPIRD44 | DIP44_WAC4301 | DPIRD | WA | 1985 | This study | ND | ND | grp_1 |
| 51 | WAC4304 | DPIRD45 | DIP45_WAC4304 | DPIRD | WA | 1985 | This study | ND | ND | grp_2 |
| 52 | WAC4308 | DPIRD46 | DIP46_WAC4308 | DPIRD | WA | 1985 | This study | ND | ND | grp_3 |
| 53 | WAC4310 | DPIRD47 | DIP47_WAC4310 | DPIRD | WA | 1985 | This study | ND | ND | grp_1 |
| 54 | WAC4311 | DPIRD48 | DIP48_WAC4311 | DPIRD | Badgingarra | 1985 | This study | ND | ND | grp_1 |

| | | | | | | | | | | |
|---|---|---|---|---|---|---|---|---|---|---|
| 55 | WAC4316 | DPIRD49 | DIP49_WAC4316 | DPIRD | WA | 1985 | This study | ND | ND | grp_2 |
| 56 | WAC1190 | DPIRD5 | DIP5_WAC1190 | DPIRD | WA | 1968 | This study | ND | ND | grp_6 |
| 57 | WAC4317 | DPIRD50 | DIP50_WAC4317 | DPIRD | WA | 1985 | This study | ND | ND | grp_1 |
| 58 | WAC4318 | DPIRD51 | DIP51_WAC4318 | DPIRD | WA | 1985 | This study | ND | ND | grp_1 |
| 59 | WAC8383 | DPIRD55 | DIP55_WAC8383 | DPIRD | Geraldton | 1990 | This study | ND | ND | grp_1 |
| 60 | WAC8387 | DPIRD56 | DIP56_WAC8387 | DPIRD | Badgingarra | 1990 | This study | ND | ND | grp_1 |
| 61 | WAC8391 | DPIRD57 | DIP57_WAC8391 | DPIRD | Badgingarra | 1990 | This study | ND | ND | grp_1 |
| 62 | WAC8399 | DPIRD58 | DIP58_WAC8399 | DPIRD | Badgingarra | 1991 | This study | ND | ND | grp_1 |
| 63 | WAC1193 | DPIRD6 | DIP6_WAC1193 | DPIRD | WA | 1968 | This study | ND | ND | grp_6 |
| 64 | WAC8417 | DPIRD60 | DIP60_WAC8417 | DPIRD | Badgingarra | 1991 | This study | ND | ND | grp_1 |
| 65 | WAC13864 | DPIRD62 | DIP62_WAC13864 | DPIRD | Badgingarra | 2015 | This study | ND | ND | grp_1 |
| 66 | WAC13865 | DPIRD63 | DIP63_WAC13865 | DPIRD | Kalannie | 2015 | This study | ND | ND | grp_1 |
| 67 | WAC13866 | DPIRD64 | DIP64_WAC13866 | DPIRD | Kojonup | 2015 | This study | ND | ND | grp_1 |
| 68 | WAC13868 | DPIRD66 | DIP66_WAC13868 | DPIRD | Wongan Hills | 2015 | This study | ND | ND | grp_1 |
| 69 | WAC13870 | DPIRD68 | DIP68_WAC13870 | DPIRD | Geraldton | 2015 | This study | ND | ND | grp_1 |
| 70 | WAC13871 | DPIRD69 | DIP69_WAC13871 | DPIRD | Geraldton | 2015 | This study | ND | ND | grp_1 |
| 71 | WAC1195 | DPIRD7 | DIP7_WAC1195 | DPIRD | WA | 1968 | This study | ND | ND | grp_6 |
| 72 | WAC13872 | DPIRD70 | DIP70_WAC13872 | DPIRD | Geraldton | 2015 | This study | ND | ND | grp_1 |
| 73 | WAC13873 | DPIRD71 | DIP71_WAC13873 | DPIRD | Nungarin | 2015 | This study | ND | ND | grp_1 |
| 74 | WAC13957 | DPIRD72 | DIP72_WAC13957 | DPIRD | South Perth | 2016 | This study | ND | ND | grp_2 |
| 75 | WAC13958 | DPIRD73 | DIP73_WAC13958 | DPIRD | South Perth | 2016 | This study | ND | ND | grp_4 |
| 76 | WAC13959 | DPIRD74 | DIP74_WAC13959 | DPIRD | South Perth | 2016 | This study | ND | ND | grp_4 |
| 77 | WAC13960 | DPIRD75 | DIP75_WAC13960 | DPIRD | South Perth | 2016 | This study | ND | ND | grp_4 |
| 78 | WAC13966 | DPIRD76 | DIP76_WAC13966 | DPIRD | Merredin | 2016 | This study | ND | ND | grp_1 |
| 79 | WAC13967 | DPIRD77 | DIP77_WAC13967 | DPIRD | Merredin | 2016 | This study | ND | ND | grp_1 |
| 80 | WAC13969 | DPIRD79 | DIP79_WAC13969 | DPIRD | Wongan Hills | 2016 | This study | ND | ND | grp_1 |
| 81 | WAC1197 | DPIRD8 | DIP8_WAC1197 | DPIRD | WA | 1968 | This study | ND | ND | grp_6 |
| 82 | WAC13979 | DPIRD82 | DIP82_WAC13979 | DPIRD | Wongan Hills | 2017 | This study | ND | ND | grp_1 |
| 83 | WAC14056 | DPIRD83 | DIP83_WAC14056 | DPIRD | Wongan Hills | 2017 | This study | ND | ND | grp_1 |
| 84 | WAC14057 | DPIRD84 | DIP84_WAC14057 | DPIRD | Wongan Hills | 2017 | This study | ND | ND | grp_1 |
| 85 | WAC14058 | DPIRD85 | DIP85_WAC14058 | DPIRD | Wongan Hills | 2017 | This study | ND | ND | grp_1 |
| 86 | WAC14059 | DPIRD86 | DIP86_WAC14059 | DPIRD | Geraldton | 2017 | This study | ND | ND | grp_1 |
| 87 | WAC14060 | DPIRD87 | DIP87_WAC14060 | DPIRD | Geraldton | 2017 | This study | ND | ND | grp_1 |
| 88 | WAC14061 | DPIRD88 | DIP88_WAC14061 | DPIRD | Northam | 2017 | This study | ND | ND | grp_1 |
| 89 | WAC14062 | DPIRD89 | DIP89_WAC14062 | DPIRD | Wongan Hills | 2017 | This study | ND | ND | grp_1 |
| 90 | WAC1199 | DPIRD9 | DIP9_WAC1199 | DPIRD | WA | 1968 | This study | ND | ND | grp_6 |
| 91 | WAC14066 | DPIRD90 | DIP90_WAC14066 | DPIRD | Northam | 2017 | This study | ND | ND | grp_1 |
| 92 | WAC14067 | DPIRD91 | DIP91_WAC14067 | DPIRD | Northam | 2017 | This study | ND | ND | grp_1 |
| 93 | WAC14068 | DPIRD92 | DIP92_WAC14068 | DPIRD | Northam | 2017 | This study | ND | ND | grp_1 |
| 94 | WAC14137 | DPIRD93 | DIP93_WAC14137 | DPIRD | Geraldton | 2018 | This study | ND | ND | grp_1 |
| 95 | WAC14138 | DPIRD94 | DIP94_WAC14138 | DPIRD | Geraldton | 2018 | This study | ND | ND | grp_1 |
| 96 | WAC14140 | DPIRD96 | DIP96_WAC14140 | DPIRD | Narrogin | 2018 | This study | ND | ND | grp_1 |
| 97 | WAC14141 | DPIRD97 | DIP97_WAC14141 | DPIRD | Wongan Hills | 2018 | This study | ND | ND | grp_1 |
| 98 | WAC2216 | O1 | O1_WAC2216 | OLD | WA | 1972 | han et al 202 | 1 | 4 | grp_1 |
| 99 | WAC8384 | O10 | O10_WAC8384 | OLD | Geraldton | 1990 | han et al 202 | 1 | 4 | grp_1 |
| 100 | 15FG102 | O100 | O100_15FG102 | OLD | Dandaragan | 2015 | han et al 202 | 2 | ND | grp_1 |
| 101 | 15FG103 | O101 | O101_15FG103 | OLD | Dandaragan | 2015 | han et al 202 | 2 | ND | grp_1 |
| 102 | 15FG104 | O102 | O102_15FG104 | OLD | Dandaragan | 2015 | han et al 202 | 2 | ND | grp_1 |
| 103 | 15FG105 | O103 | O103_15FG105 | OLD | Dandaragan | 2015 | han et al 202 | 2 | ND | grp_1 |
| 104 | FG106 | O104 | O104_FG106 | OLD | Northam | 2015 | han et al 202 | 2 | 4 | grp_1 |
| 105 | FG107 | O105 | O105_FG107 | OLD | Northam | 2015 | han et al 202 | 2 | 4 | grp_1 |
| 106 | FG108 | O106 | O106_FG108 | OLD | Northam | 2015 | han et al 202 | 2 | 4 | grp_1 |
| 107 | 15FG107 | O107 | O107_15FG107 | OLD | Jennacubbine | 2014 | han et al 202 | 2 | ND | grp_1 |
| 108 | 15FG109 | O108 | O108_15FG109 | OLD | Arrowsmith Eas | 2014 | han et al 202 | 2 | ND | grp_1 |
| 109 | 15FG110 | O109 | O109_15FG110 | OLD | Southern Brool | 2015 | han et al 202 | 2 | ND | grp_1 |
| 110 | WAC8390 | O11 | O11_WAC8390 | OLD | Badgingarra | 1991 | han et al 202 | 1 | 4 | grp_1 |

| | | | | | | | | | |
|---|---|---|---|---|---|---|---|---|---|
| 111 | 15FG112 | O110 | O110_15FG112 | OLD | Southern Brool | 2015 | han et al 202 | 2 | ND | grp_1 |
| 112 | 15FG113 | O111 | O111_15FG113 | OLD | Southern Brool | 2015 | han et al 202 | 2 | ND | grp_1 |
| 113 | 15FG114 | O112 | O112_15FG114 | OLD | Southern Brool | 2015 | han et al 202 | 2 | ND | grp_1 |
| 114 | 15FG115 | O113 | O113_15FG115 | OLD | Southern Brool | 2015 | han et al 202 | 2 | ND | grp_1 |
| 115 | 15FG116 | O114 | O114_15FG116 | OLD | Southern Brool | 2015 | han et al 202 | 2 | ND | grp_1 |
| 116 | 15FG117 | O115 | O115_15FG117 | OLD | Southern Brool | 2015 | han et al 202 | 2 | ND | grp_1 |
| 117 | 15FG118 | O116 | O116_15FG118 | OLD | Southern Brool | 2015 | han et al 202 | 2 | ND | grp_1 |
| 118 | 15FG119 | O117 | O117_15FG119 | OLD | Southern Brool | 2015 | han et al 202 | 2 | 4 | grp_1 |
| 119 | 15FG120 | O118 | O118_15FG120 | OLD | Jennacubbine | 2014 | han et al 202 | 2 | 4 | grp_1 |
| 120 | 15FG04 | O119 | O119_15FG04 | OLD | WA | 2015 | han et al 202 | 2 | ND | grp_1 |
| 121 | WAC8410 | O12 | O12_WAC8410 | OLD | Badgingarra | 1991 | han et al 202 | 1 | 4 | grp_1 |
| 122 | 15FG28 | O120 | O120_15FG28 | OLD | Northam | 2015 | han et al 202 | 2 | 4 | grp_1 |
| 123 | 15FG33 | O122 | O122_15FG33 | OLD | Northam | 2015 | han et al 202 | 2 | 4 | grp_1 |
| 124 | 15FG37 | O123 | O123_15FG37 | OLD | Muresk | 2015 | han et al 202 | 2 | 4 | grp_1 |
| 125 | 15FG38 | O124 | O124_15FG38 | OLD | Muresk | 2015 | han et al 202 | 2 | 4 | grp_1 |
| 126 | 15FG47 | O127 | O127_15FG47 | OLD | Muresk | 2015 | han et al 202 | 2 | 4 | grp_1 |
| 127 | 15FG49 | O128 | O128_15FG49 | OLD | Muresk | 2015 | han et al 202 | 2 | 4 | grp_1 |
| 128 | 15FG226 | O129 | O129_15FG226 | OLD | Cunderdin | 2015 | han et al 202 | 2 | 4 | grp_1 |
| 129 | WAC8635 | O13 | O13_WAC8635 | OLD | WA | 1994 | han et al 202 | 3 | 2 | grp_3 |
| 130 | 15FG229 | O130 | O130_15FG229 | OLD | Eradu | 2015 | han et al 202 | 1 | 4 | grp_1 |
| 131 | 15FG237 | O131 | O131_15FG237 | OLD | Esperance | 2015 | han et al 202 | 1 | 4 | grp_1 |
| 132 | Northam_Mace2 | O132 | O132_Northam_Mace2 | OLD | Northam | 2015 | han et al 202 | 1 | 4 | grp_1 |
| 133 | Northam_Magent | O133 | O133_Northam_Magenta | OLD | Northam | 2015 | han et al 202 | 2 | 4 | grp_1 |
| 134 | Northam_Emu1 | O134 | O134_Northam_Emu1 | OLD | Northam | 2015 | han et al 202 | 1 | 4 | grp_1 |
| 135 | Northam_Emu2 | O135 | O135_Northam_Emu2 | OLD | Northam | 2015 | han et al 202 | 2 | 4 | grp_1 |
| 136 | Northam_Mace1 | O136 | O136_Northam_Mace1 | OLD | Northam | 2015 | han et al 202 | 2 | 4 | grp_1 |
| 137 | WAC9178 | O14 | O14_WAC9178 | OLD | South Perth | 1996 | han et al 202 | 3 | 2 | grp_3 |
| 138 | FG_W003_5 | O140 | O140_FG_W003_5 | OLD | WA | 2015 | han et al 202 | 1 | 4 | grp_1 |
| 139 | Nor_RAC2182_1 | O141 | O141_Nor_RAC2182_1 | OLD | Northam | 2015 | han et al 202 | 2 | 4 | grp_1 |
| 140 | Northam_WGT | O142 | O142_Northam_WGT | OLD | Northam | 2015 | han et al 202 | 2 | 4 | grp_1 |
| 141 | Nor_RAC2182_2 | O143 | O143_Nor_RAC2182_2 | OLD | Northam | 2015 | han et al 202 | 2 | 4 | grp_1 |
| 142 | WAC13418 | O15 | O15_WAC13418 | OLD | Geraldton | 1996 | han et al 202 | 3 | 2 | grp_3 |
| 143 | SN15 | O16 | O16_SN15 | OLD | WA | 2001 | han et al 202 | 3 | 2 | grp_3 |
| 144 | WAC13068 | O17 | O17_WAC13068 | OLD | Geraldton | 2005 | han et al 202 | 1 | 6 | grp_2 |
| 145 | WAC13069 | O18 | O18_WAC13069 | OLD | Geraldton | 2005 | han et al 202 | 1 | 6 | grp_2 |
| 146 | WAC13070 | O19 | O19_WAC13070 | OLD | Geraldton | 2005 | han et al 202 | 4 | 1 | grp_4 |
| 147 | WAC2285 | O2 | O2_WAC2285 | OLD | WA | 1973 | han et al 202 | ND | ND | grp_1 |
| 148 | WAC13071 | O20 | O20_WAC13071 | OLD | Geraldton | 2005 | han et al 202 | 1 | 4 | grp_1 |
| 149 | WAC13072 | O21 | O21_WAC13072 | OLD | Geraldton | 2005 | han et al 202 | 1 | 4 | grp_1 |
| 150 | WAC13073 | O22 | O22_WAC13073 | OLD | Geraldton | 2005 | han et al 202 | 1 | 4 | grp_1 |
| 151 | WAC13074 | O23 | O23_WAC13074 | OLD | Geraldton | 2005 | han et al 202 | 4 | 1 | grp_4 |
| 152 | WAC13075 | O24 | O24_WAC13075 | OLD | Geraldton | 2005 | han et al 202 | 4 | 1 | grp_4 |
| 153 | WAC13076 | O25 | O25_WAC13076 | OLD | Geraldton | 2005 | han et al 202 | 4 | 1 | grp_4 |
| 154 | WAC13077 | O26 | O26_WAC13077 | OLD | Geraldton | 2005 | han et al 202 | 1 | 4 | grp_1 |
| 155 | Meck1 | O27 | O27_Meck1 | OLD | Meckering | 2009 | han et al 202 | 1 | 4 | grp_1 |
| 156 | Meck3 | O28 | O28_Meck3 | OLD | Meckering | 2009 | han et al 202 | 1 | 4 | grp_1 |
| 157 | Meck5 | O29 | O29_Meck5 | OLD | Meckering | 2009 | han et al 202 | ND | 4 | grp_1 |
| 158 | WAC2810 | O3 | O3_WAC2810 | OLD | NA | 1980 | han et al 202 | ND | 2 | grp_3 |
| 159 | Meck6 | O30 | O30_Meck6 | OLD | Meckering | 2009 | han et al 202 | 1 | 4 | grp_1 |
| 160 | Meck8 | O31 | O31_Meck8 | OLD | Meckering | 2009 | han et al 202 | 1 | 4 | grp_1 |
| 161 | S1FT3A | O32 | O32_S1FT3A | OLD | Meckering | 2009 | han et al 202 | 1 | 4 | grp_1 |
| 162 | S4FT3A | O33 | O33_S4FT3A | OLD | Meckering | 2009 | han et al 202 | 1 | 4 | grp_1 |
| 163 | S1FT3B | O34 | O34_S1FT3B | OLD | Meckering | 2009 | han et al 202 | 1 | 4 | grp_1 |
| 164 | Gerald1 | O35 | O35_Gerald1 | OLD | Geraldton | 2011 | han et al 202 | ND | 4 | grp_1 |
| 165 | Gerald4 | O36 | O36_Gerald4 | OLD | Geraldton | 2011 | han et al 202 | 1 | 4 | grp_1 |
| 166 | Mur_51 | O37 | O37_Mur_51 | OLD | Murdoch | 2011 | han et al 202 | 1 | 4 | grp_1 |

| | | | | | | | | | |
|---|---|---|---|---|---|---|---|---|---|
| 167 | WAC13402 | O39 | O39_WAC13402 | OLD | Geraldton | 2011 han et al 202 | 1 | 4 | grp_1 |
| 168 | WAC2813 | O4 | O4_WAC2813 | OLD | WA | 1980 han et al 202 | 3 | 2 | grp_3 |
| 169 | WAC13403 | O40 | O40_WAC13403 | OLD | Geraldton | 2011 han et al 202 | 4 | 1 | grp_4 |
| 170 | WAC13404 | O41 | O41_WAC13404 | OLD | Geraldton | 2011 han et al 202 | 4 | 1 | grp_4 |
| 171 | WAC13405 | O42 | O42_WAC13405 | OLD | Geraldton | 2011 han et al 202 | 3 | 2 | grp_3 |
| 172 | WAC13443 | O43 | O43_WAC13443 | OLD | Geraldton | 2011 han et al 202 | 1 | 4 | grp_1 |
| 173 | WAC13446 | O44 | O44_WAC13446 | OLD | Geraldton | 2011 han et al 202 | 4 | 1 | grp_4 |
| 174 | WAC13447 | O45 | O45_WAC13447 | OLD | Geraldton | 2011 han et al 202 | 3 | 2 | grp_3 |
| 175 | WAC13523 | O46 | O46_WAC13523 | OLD | Geraldton | 2011 han et al 202 | 1 | 6 | grp_2 |
| 176 | WAC13524 | O47 | O47_WAC13524 | OLD | Geraldton | 2011 han et al 202 | 1 | 6 | grp_2 |
| 177 | WAC13525 | O48 | O48_WAC13525 | OLD | Geraldton | 2011 han et al 202 | 4 | 1 | grp_4 |
| 178 | WAC13526 | O49 | O49_WAC13526 | OLD | Geraldton | 2011 han et al 202 | 4 | 1 | grp_4 |
| 179 | WAC4303 | O5 | O5_WAC4303 | OLD | WA | 1985 han et al 202 | 1 | 4 | grp_1 |
| 180 | WAC13527 | O50 | O50_WAC13527 | OLD | Geraldton | 2011 han et al 202 | 1 | 6 | grp_2 |
| 181 | WAC13528 | O51 | O51_WAC13528 | OLD | Geraldton | 2011 han et al 202 | 4 | 1 | grp_4 |
| 182 | WAC13529 | O52 | O52_WAC13529 | OLD | Geraldton | 2011 han et al 202 | 1 | 6 | grp_2 |
| 183 | WAC13530 | O53 | O53_WAC13530 | OLD | Geraldton | 2011 han et al 202 | 1 | 6 | grp_2 |
| 184 | WAC13531 | O54 | O54_WAC13531 | OLD | Geraldton | 2011 han et al 202 | 1 | 6 | grp_2 |
| 185 | WAC13532 | O55 | O55_WAC13532 | OLD | Geraldton | 2011 han et al 202 | 4 | 1 | grp_4 |
| 186 | WAC13615 | O56 | O56_WAC13615 | OLD | Geraldton | 2012 han et al 202 | 2 | 4 | grp_1 |
| 187 | WAC13616 | O57 | O57_WAC13616 | OLD | Geraldton | 2012 han et al 202 | 2 | 4 | grp_1 |
| 188 | WAC13617 | O58 | O58_WAC13617 | OLD | Geraldton | 2012 han et al 202 | 2 | 4 | grp_1 |
| 189 | WAC13630 | O59 | O59_WAC13630 | OLD | Geraldton | 2012 han et al 202 | 4 | 1 | grp_4 |
| 190 | WAC4321 | O6 | O6_WAC4321 | OLD | Mount Barker | 1985 han et al 202 | 1 | 4 | grp_1 |
| 191 | WAC13631 | O60 | O60_WAC13631 | OLD | Geraldton | 2012 han et al 202 | 4 | 1 | grp_4 |
| 192 | WAC13632 | O61 | O61_WAC13632 | OLD | Geraldton | 2012 han et al 202 | 4 | 1 | grp_4 |
| 193 | WAC13666 | O62 | O62_WAC13666 | OLD | Geraldton | 2013 han et al 202 | 1 | 4 | grp_1 |
| 194 | WAC13667 | O63 | O63_WAC13667 | OLD | Geraldton | 2013 han et al 202 | 1 | 4 | grp_1 |
| 195 | WAC13690 | O64 | O64_WAC13690 | OLD | Dongara | 2012 han et al 202 | 1 | 4 | grp_1 |
| 196 | WAC13691 | O65 | O65_WAC13691 | OLD | Dongara | 2012 han et al 202 | 1 | 4 | grp_1 |
| 197 | 206FG226 | O66 | O66_206FG226 | OLD | Regans Ford | 2014 han et al 202 | 1 | 4 | grp_1 |
| 198 | 205FG215_1 | O67 | O67_205FG215_1 | OLD | Regans Ford | 2014 han et al 202 | 2 | 4 | grp_1 |
| 199 | 201FG209 | O68 | O68_201FG209 | OLD | Regans Ford | 2014 han et al 202 | 1 | 4 | grp_1 |
| 200 | 201FG211 | O69 | O69_201FG211 | OLD | Regans Ford | 2014 han et al 202 | 1 | 4 | grp_1 |
| 201 | WAC4319 | O7 | O7_WAC4319 | OLD | WA | 1985 han et al 202 | 1 | 4 | grp_2 |
| 202 | 202FG212 | O70 | O70_202FG212 | OLD | Regans Ford | 2014 han et al 202 | 1 | 4 | grp_1 |
| 203 | 204FG221 | O71 | O71_204FG221 | OLD | Regans Ford | 2014 han et al 202 | 1 | 4 | grp_1 |
| 204 | 204FG223 | O72 | O72_204FG223 | OLD | Regans Ford | 2014 han et al 202 | 1 | 4 | grp_1 |
| 205 | 205FG225 | O73 | O73_205FG225 | OLD | Regans Ford | 2014 han et al 202 | 2 | 4 | grp_1 |
| 206 | 206FG227 | O74 | O74_206FG227 | OLD | Regans Ford | 2014 han et al 202 | 1 | 4 | grp_1 |
| 207 | 205FG216 | O75 | O75_205FG216 | OLD | Regans Ford | 2014 han et al 202 | 1 | 4 | grp_1 |
| 208 | 201FG219 | O76 | O76_201FG219 | OLD | Regans Ford | 2014 han et al 202 | 1 | 4 | grp_1 |
| 209 | 201FG218 | O77 | O77_201FG218 | OLD | Regans Ford | 2014 han et al 202 | 1 | 4 | grp_1 |
| 210 | 204FG214 | O78 | O78_204FG214 | OLD | Regans Ford | 2014 han et al 202 | 1 | 4 | grp_1 |
| 211 | 903FG214 | O79 | O79_903FG214 | OLD | Regans Ford | 2014 han et al 202 | ND | 4 | grp_1 |
| 212 | WAC4808 | O8 | O8_WAC4808 | OLD | East Chapman | 1986 han et al 202 | 1 | 4 | grp_1 |
| 213 | 201FG49 | O80 | O80_201FG49 | OLD | Wongan Hills | 2014 han et al 202 | 2 | 4 | grp_1 |
| 214 | 203FG58 | O81 | O81_203FG58 | OLD | Wongan Hills | 2014 han et al 202 | 1 | 4 | grp_1 |
| 215 | 205FG63 | O82 | O82_205FG63 | OLD | Wongan Hills | 2014 han et al 202 | 2 | 4 | grp_1 |
| 216 | 206FG66 | O83 | O83_206FG66 | OLD | Wongan Hills | 2014 han et al 202 | 1 | 4 | grp_1 |
| 217 | 206FG67 | O84 | O84_206FG67 | OLD | Wongan Hills | 2014 han et al 202 | 2 | 4 | grp_1 |
| 218 | 206FG68 | O85 | O85_206FG68 | OLD | Wongan Hills | 2014 han et al 202 | 1 | 4 | grp_1 |
| 219 | 205FG142 | O86 | O86_205FG142 | OLD | York | 2014 han et al 202 | 2 | 4 | grp_1 |
| 220 | 53FG143_1 | O87 | O87_53FG143_1 | OLD | East Buntine | 2014 han et al 202 | 1 | 4 | grp_1 |
| 221 | 201FG208 | O88 | O88_201FG208 | OLD | Regans Ford | 2014 han et al 202 | 1 | 4 | grp_1 |
| 222 | 203FG213 | O89 | O89_203FG213 | OLD | Regans Ford | 2014 han et al 202 | 1 | 4 | grp_1 |

| | | | | | | | | | | |
|---|---|---|---|---|---|---|---|---|---|---|
| 223 | WAC4648 | O9 | O9_WAC4648 | OLD | WA | 1986 | han et al 202 | 1 | 4 | grp_2 |
| 224 | 204FG222 | O91 | O91_204FG222 | OLD | Regans Ford | 2014 | han et al 202 | 1 | 4 | grp_1 |
| 225 | 205FG410 | O92 | O92_205FG410 | OLD | Wongan Hills | 2014 | han et al 202 | 1 | 4 | grp_1 |
| 226 | 202FG414 | O93 | O93_202FG414 | OLD | York | 2014 | han et al 202 | 2 | 4 | grp_1 |
| 227 | WAC739 | O94 | O94_WAC739 | OLD | Eneabba | 2014 | han et al 202 | 2 | 4 | grp_1 |
| 228 | WAC740 | O95 | O95_WAC740 | OLD | Badgingarra | 2014 | han et al 202 | 2 | 4 | grp_1 |
| 229 | WAC741 | O96 | O96_WAC741 | OLD | Carnamah | 2014 | han et al 202 | 2 | 4 | grp_1 |
| 230 | 15FG99 | O97 | O97_15FG99 | OLD | Dandaragan | 2015 | han et al 202 | 2 | 4 | grp_1 |
| 231 | 15FG100 | O98 | O98_15FG100 | OLD | Dandaragan | 2015 | han et al 202 | 2 | ND | grp_1 |
| 232 | 15FG101 | O99 | O99_15FG101 | OLD | Dandaragan | 2015 | han et al 202 | 2 | ND | grp_1 |
| 233 | 16FG06 | O143 | O143_16FG06 | OLD | Mingenew | 2016 | han et al 202 | 1 | 4 | grp_1 |
| 234 | 16FG158 | O144 | O144_16FG158 | OLD | Mingenew | 2016 | han et al 202 | 1 | 4 | grp_1 |
| 235 | 16FG159 | O145 | O145_16FG159 | OLD | Mingenew | 2016 | han et al 202 | 1 | 4 | grp_1 |
| 236 | 16FG160 | O146 | O146_16FG160 | OLD | Mingenew | 2016 | han et al 202 | 5 | 5 | grp_5 |
| 237 | 16FG161 | O147 | O147_16FG161 | OLD | Mingenew | 2016 | han et al 202 | 5 | 5 | grp_5 |
| 238 | 16FG162 | O148 | O148_16FG162 | OLD | Mingenew | 2016 | han et al 202 | 5 | 5 | grp_5 |
| 239 | 16FG163_2 | O150 | O150_16FG163_2 | OLD | Mingenew | 2016 | han et al 202 | 5 | 5 | grp_5 |
| 240 | 16FG164 | O151 | O151_16FG164 | OLD | Mingenew | 2016 | han et al 202 | 5 | 5 | grp_5 |
| 241 | 16FG165 | O152 | O152_16FG165 | OLD | Mingenew | 2016 | han et al 202 | 5 | 5 | grp_5 |
| 242 | 16FG166 | O153 | O153_16FG166 | OLD | Mingenew | 2016 | han et al 202 | 5 | 5 | grp_5 |
| 243 | 16FG167 | O154 | O154_16FG167 | OLD | Mingenew | 2016 | han et al 202 | 5 | 5 | grp_5 |
| 244 | 16FG168 | O155 | O155_16FG168 | OLD | Mingenew | 2016 | han et al 202 | 5 | 5 | grp_5 |
| 245 | 16FG169 | O156 | O156_16FG169 | OLD | Mingenew | 2016 | han et al 202 | 5 | 5 | grp_5 |
| 246 | WAC13955 | O157 | O157_WAC13955 | OLD | Northam | 2016 | han et al 202 | 1 | 4 | grp_1 |
| 247 | 16FG170 | O158 | O158_16FG170 | OLD | WA | 2016 | han et al 202 | 5 | 5 | grp_5 |
| 248 | 16FG171 | O159 | O159_16FG171 | OLD | WA | 2016 | han et al 202 | 5 | 5 | grp_5 |
| 249 | Mur_S3 | O160 | O160_Mur_S3 | OLD | Murdoch | 2011 | han et al 202 | 2 | 4 | grp_1 |
| 250 | 14FG141 | O161 | O161_14FG141 | OLD | Arrowsmith Eas | 2014 | han et al 202 | 2 | 4 | grp_1 |
| 251 | PN260 | C260 | C260_PN260 | Current | Northam | 2019 | This study | ND | ND | grp_1 |
| 252 | PN261 | C261 | C261_PN261 | Current | Northam | 2019 | This study | ND | ND | grp_1 |
| 253 | PN262 | C262 | C262_PN262 | Current | Northam | 2019 | This study | ND | ND | grp_1 |
| 254 | PN263 | C263 | C263_PN263 | Current | Northam | 2019 | This study | ND | ND | grp_1 |
| 255 | PN264 | C264 | C264_PN264 | Current | Northam | 2019 | This study | ND | ND | grp_1 |
| 256 | PN265 | C265 | C265_PN265 | Current | Northam | 2019 | This study | ND | ND | grp_1 |
| 257 | PN266 | C266 | C266_PN266 | Current | Northam | 2019 | This study | ND | ND | grp_1 |
| 258 | PN268 | C268 | C268_PN268 | Current | Northam | 2019 | This study | ND | ND | grp_1 |
| 259 | PN269 | C269 | C269_PN269 | Current | Northam | 2019 | This study | ND | ND | grp_1 |
| 260 | PN270 | C270 | C270_PN270 | Current | Northam | 2019 | This study | ND | ND | grp_1 |
| 261 | PN271 | C271 | C271_PN271 | Current | Northam | 2019 | This study | ND | ND | grp_1 |
| 262 | PN274 | C274 | C274_PN274 | Current | Northam | 2019 | This study | ND | ND | grp_1 |
| 263 | PN275 | C275 | C275_PN275 | Current | Northam | 2019 | This study | ND | ND | grp_1 |
| 264 | PN276 | C276 | C276_PN276 | Current | Northam | 2019 | This study | ND | ND | grp_1 |
| 265 | PN277 | C277 | C277_PN277 | Current | Northam | 2019 | This study | ND | ND | grp_1 |
| 266 | PN279 | C279 | C279_PN279 | Current | Northam | 2019 | This study | ND | ND | grp_1 |
| 267 | PN280 | C280 | C280_PN280 | Current | Northam | 2019 | This study | ND | ND | grp_7 |
| 268 | PN282 | C282 | C282_PN282 | Current | Northam | 2019 | This study | ND | ND | grp_7 |
| 269 | PN283 | C283 | C283_PN283 | Current | Northam | 2019 | This study | ND | ND | grp_7 |
| 270 | PN284 | C284 | C284_PN284 | Current | Northam | 2019 | This study | ND | ND | grp_1 |
| 271 | PN285 | C285 | C285_PN285 | Current | Northam | 2019 | This study | ND | ND | grp_1 |
| 272 | PN286 | C286 | C286_PN286 | Current | Northam | 2019 | This study | ND | ND | grp_1 |
| 273 | PN287 | C287 | C287_PN287 | Current | Northam | 2019 | This study | ND | ND | grp_1 |
| 274 | PN288 | C288 | C288_PN288 | Current | Northam | 2019 | This study | ND | ND | grp_1 |
| 275 | PN289 | C289 | C289_PN289 | Current | Northam | 2019 | This study | ND | ND | grp_1 |
| 276 | PN290 | C290 | C290_PN290 | Current | Northam | 2019 | This study | ND | ND | grp_1 |
| 277 | PN291 | C291 | C291_PN291 | Current | Northam | 2019 | This study | ND | ND | grp_1 |
| 278 | PN292 | C292 | C292_PN292 | Current | Northam | 2019 | This study | ND | ND | grp_1 |

| | | | | | | | | | | |
|---|---|---|---|---|---|---|---|---|---|---|
| 279 | PN295 | C295 | C295_PN295 | Current | Northam | 2019 | This study | ND | ND | grp_1 |
| 280 | PN296 | C296 | C296_PN296 | Current | Northam | 2019 | This study | ND | ND | grp_8 |
| 281 | PN297 | C297 | C297_PN297 | Current | Northam | 2019 | This study | ND | ND | grp_1 |
| 282 | PN298 | C298 | C298_PN298 | Current | Northam | 2019 | This study | ND | ND | grp_8 |
| 283 | PN299 | C299 | C299_PN299 | Current | Northam | 2019 | This study | ND | ND | grp_1 |
| 284 | PN300 | C300 | C300_PN300 | Current | Northam | 2019 | This study | ND | ND | grp_1 |
| 285 | PN301 | C301 | C301_PN301 | Current | Northam | 2019 | This study | ND | ND | grp_1 |
| 286 | PN302 | C302 | C302_PN302 | Current | Northam | 2019 | This study | ND | ND | grp_1 |
| 287 | PN303 | C303 | C303_PN303 | Current | Northam | 2019 | This study | ND | ND | grp_1 |
| 288 | PN304 | C304 | C304_PN304 | Current | Northam | 2019 | This study | ND | ND | grp_1 |
| 289 | PN305 | C305 | C305_PN305 | Current | Northam | 2019 | This study | ND | ND | grp_1 |
| 290 | PN306 | C306 | C306_PN306 | Current | Northam | 2019 | This study | ND | ND | grp_7 |
| 291 | PN307 | C307 | C307_PN307 | Current | Northam | 2019 | This study | ND | ND | grp_7 |
| 292 | PN308 | C308 | C308_PN308 | Current | Northam | 2019 | This study | ND | ND | grp_1 |
| 293 | PN309 | C309 | C309_PN309 | Current | Northam | 2019 | This study | ND | ND | grp_1 |
| 294 | PN312 | C312 | C312_PN312 | Current | Northam | 2019 | This study | ND | ND | grp_1 |
| 295 | PN313 | C313 | C313_PN313 | Current | Northam | 2019 | This study | ND | ND | grp_1 |
| 296 | PN314 | C314 | C314_PN314 | Current | Northam | 2019 | This study | ND | ND | grp_1 |
| 297 | PN315 | C315 | C315_PN315 | Current | Northam | 2019 | This study | ND | ND | grp_7 |
| 298 | PN316 | C316 | C316_PN316 | Current | Northam | 2019 | This study | ND | ND | grp_7 |
| 299 | PN317 | C317 | C317_PN317 | Current | Northam | 2019 | This study | ND | ND | grp_1 |
| 300 | PN318 | C318 | C318_PN318 | Current | Northam | 2019 | This study | ND | ND | grp_1 |
| 301 | PN319 | C319 | C319_PN319 | Current | Northam | 2019 | This study | ND | ND | grp_1 |
| 302 | PN320 | C320 | C320_PN320 | Current | Northam | 2019 | This study | ND | ND | grp_1 |
| 303 | PN321 | C321 | C321_PN321 | Current | Northam | 2019 | This study | ND | ND | grp_8 |
| 304 | PN322 | C322 | C322_PN322 | Current | Northam | 2019 | This study | ND | ND | grp_8 |
| 305 | PN323 | C323 | C323_PN323 | Current | Northam | 2019 | This study | ND | ND | grp_1 |
| 306 | PN324 | C324 | C324_PN324 | Current | Northam | 2019 | This study | ND | ND | grp_1 |
| 307 | PN325 | C325 | C325_PN325 | Current | Northam | 2019 | This study | ND | ND | grp_1 |
| 308 | PN326 | C326 | C326_PN326 | Current | Northam | 2019 | This study | ND | ND | grp_1 |
| 309 | PN328 | C328 | C328_PN328 | Current | Northam | 2019 | This study | ND | ND | grp_1 |
| 310 | PN329 | C329 | C329_PN329 | Current | Northam | 2019 | This study | ND | ND | grp_1 |
| 311 | PN330 | C330 | C330_PN330 | Current | Northam | 2019 | This study | ND | ND | grp_1 |
| 312 | PN331 | C331 | C331_PN331 | Current | Northam | 2019 | This study | ND | ND | grp_1 |
| 313 | PN332 | C332 | C332_PN332 | Current | Northam | 2019 | This study | ND | ND | grp_1 |
| 314 | PN333 | C333 | C333_PN333 | Current | Northam | 2019 | This study | ND | ND | grp_8 |
| 315 | PN334 | C334 | C334_PN334 | Current | Northam | 2019 | This study | ND | ND | grp_7 |
| 316 | PN335 | C335 | C335_PN335 | Current | Northam | 2019 | This study | ND | ND | grp_7 |
| 317 | PN336 | C336 | C336_PN336 | Current | Northam | 2019 | This study | ND | ND | grp_1 |
| 318 | PN337 | C337 | C337_PN337 | Current | Northam | 2019 | This study | ND | ND | grp_7 |
| 319 | PN338 | C338 | C338_PN338 | Current | Northam | 2019 | This study | ND | ND | grp_7 |
| 320 | PN340 | C340 | C340_PN340 | Current | Northam | 2019 | This study | ND | ND | grp_7 |
| 321 | PN341 | C341 | C341_PN341 | Current | Northam | 2019 | This study | ND | ND | grp_8 |
| 322 | PN342 | C342 | C342_PN342 | Current | Northam | 2019 | This study | ND | ND | grp_1 |
| 323 | PN343 | C343 | C343_PN343 | Current | Northam | 2019 | This study | ND | ND | grp_7 |
| 324 | PN344 | C344 | C344_PN344 | Current | Northam | 2019 | This study | ND | ND | grp_8 |
| 325 | PN345 | C345 | C345_PN345 | Current | Northam | 2019 | This study | ND | ND | grp_7 |
| 326 | PN346 | C346 | C346_PN346 | Current | Northam | 2019 | This study | ND | ND | grp_7 |
| 327 | PN347 | C347 | C347_PN347 | Current | Northam | 2019 | This study | ND | ND | grp_1 |
| 328 | PN348 | C348 | C348_PN348 | Current | Northam | 2019 | This study | ND | ND | grp_1 |
| 329 | PN349 | C349 | C349_PN349 | Current | Northam | 2019 | This study | ND | ND | grp_1 |
| 330 | PN350 | C350 | C350_PN350 | Current | Northam | 2019 | This study | ND | ND | grp_8 |
| 331 | PN351 | C351 | C351_PN351 | Current | Northam | 2020 | This study | ND | ND | grp_1 |
| 332 | PN352 | C352 | C352_PN352 | Current | Northam | 2020 | This study | ND | ND | grp_1 |
| 333 | PN353 | C353 | C353_PN353 | Current | Northam | 2021 | This study | ND | ND | grp_1 |
| 334 | PN354 | C354 | C354_PN354 | Current | Northam | 2021 | This study | ND | ND | grp_1 |

| | | | | | | | | | | |
|---|---|---|---|---|---|---|---|---|---|---|
| 335 | PN355 | C355 | C355_PN355 | Current | Northam | 2021 | This study | ND | ND | grp_8 |
| 336 | PN356 | C356 | C356_PN356 | Current | Northam | 2021 | This study | ND | ND | grp_8 |
| 337 | PN357 | C357 | C357_PN357 | Current | Northam | 2021 | This study | ND | ND | grp_1 |
| 338 | PN358 | C358 | C358_PN358 | Current | Northam | 2021 | This study | ND | ND | grp_1 |
| 339 | PN359 | C359 | C359_PN359 | Current | Northam | 2021 | This study | ND | ND | grp_8 |
| 340 | PN360 | C360 | C360_PN360 | Current | Northam | 2021 | This study | ND | ND | grp_1 |
| 341 | PN361 | C361 | C361_PN361 | Current | Northam | 2021 | This study | ND | ND | grp_1 |
| 342 | PN362 | C362 | C362_PN362 | Current | Northam | 2021 | This study | ND | ND | grp_1 |
| 343 | PN363 | C363 | C363_PN363 | Current | Northam | 2021 | This study | ND | ND | grp_1 |
| 344 | PN364 | C364 | C364_PN364 | Current | Northam | 2021 | This study | ND | ND | grp_1 |
| 345 | PN365 | C365 | C365_PN365 | Current | Northam | 2021 | This study | ND | ND | grp_8 |
| 346 | PN366 | C366 | C366_PN366 | Current | Northam | 2021 | This study | ND | ND | grp_1 |
| 347 | PN367 | C367 | C367_PN367 | Current | Northam | 2021 | This study | ND | ND | grp_1 |
| 348 | PN368 | C368 | C368_PN368 | Current | Northam | 2021 | This study | ND | ND | grp_8 |
| 349 | PN369 | C369 | C369_PN369 | Current | Northam | 2021 | This study | ND | ND | grp_1 |
| 350 | PN370 | C370 | C370_PN370 | Current | Northam | 2021 | This study | ND | ND | grp_8 |
| 351 | PN371 | C371 | C371_PN371 | Current | Northam | 2021 | This study | ND | ND | grp_1 |
| 352 | PN372 | C372 | C372_PN372 | Current | Northam | 2021 | This study | ND | ND | grp_1 |
| 353 | PN373 | C373 | C373_PN373 | Current | Northam | 2021 | This study | ND | ND | grp_8 |
| 354 | PN374 | C374 | C374_PN374 | Current | Northam | 2021 | This study | ND | ND | grp_8 |
| 355 | PN375 | C375 | C375_PN375 | Current | Northam | 2021 | This study | ND | ND | grp_1 |
| 356 | PN376 | C376 | C376_PN376 | Current | Northam | 2021 | This study | ND | ND | grp_1 |
| 357 | PN377 | C377 | C377_PN377 | Current | Northam | 2021 | This study | ND | ND | grp_1 |
| 358 | PN378 | C378 | C378_PN378 | Current | Northam | 2021 | This study | ND | ND | grp_8 |
| 359 | PN379 | C379 | C379_PN379 | Current | Northam | 2021 | This study | ND | ND | grp_8 |
| 360 | PN380 | C380 | C380_PN380 | Current | Northam | 2021 | This study | ND | ND | grp_1 |

**STable 1(cont): Metadata of isolates used in this study**

| # | Original_name | ToxA_isoform | ToxA_reference | Tox1_isoform | Tox1_reference | Tox3_isoform | Tox3_reference | Tox5_isoform | Tox5_reference | Tox267_isoform | Mating_type |
|---|---|---|---|---|---|---|---|---|---|---|---|
| 1 | WAC1141 | I03 | H03_EF108456 | I01 | H01_JN791682.1 | I01 | JX997413.1_H1 | I01 | MW715921.1, MW715898.1 | Aus_I01 | 1 |
| 2 | WAC1201 | I03 | H03_EF108456 | I01 | H01_JN791682.1 | I01 | JX997413.1_H1 | I02 | new isoform | Aus_I01 | 2 |
| 3 | WAC14144 | I03 | H03_EF108456 | I01 | H01_JN791682.1 | I01 | JX997413.1_H1 | I02 | new isoform | Aus_I01 | 2 |
| 4 | WAC2816 | I03 | H03_EF108456 | I01 | H01_JN791682.1 | I01 | JX997413.1_H1 | I01 | MW715921.1, MW715898.1 | Aus_I01 | 1 |
| 5 | WAC2812 | I03 | H03_EF108456 | I01 | H01_JN791682.1 | I01 | JX997413.1_H1 | I02 | new isoform | Aus_I01 | 2 |
| 6 | WAC8397 | I02 | H02_EF108458 | I01 | H01_JN791682.1 | Null | NA | I02 | new isoform | Aus_I01 | 1 |
| 7 | WAC2804 | I03 | H03_EF108456 | I01 | H01_JN791682.1 | I01 | JX997413.1_H1 | I01 | MW715921.1, MW715898.1 | Aus_I01 | 1 |
| 8 | WAC1544 | I03 | H03_EF108456 | I01 | H01_JN791682.1 | I01 | JX997413.1_H1 | I02 | new isoform | Aus_I01 | 2 |
| 9 | WAC2799 | I03 | H03_EF108456 | I01 | H01_JN791682.1 | I01 | JX997413.1_H1 | I01 | MW715921.1, MW715898.1 | Aus_I01 | 1 |
| 10 | WAC4312 | I02 | H02_EF108458 | I01 | H01_JN791682.1 | I01 | JX997413.1_H1 | I01 | MW715921.1, MW715898.1 | Aus_I01 | 2 |
| 11 | WAC1198 | I03 | H03_EF108456 | I01 | H01_JN791682.1 | I01 | JX997413.1_H1 | I02 | new isoform | Aus_I01 | 2 |
| 12 | WAC1494 | I03 | H03_EF108456 | I01 | H01_JN791682.1 | I01 | JX997413.1_H1 | I01 | MW715921.1, MW715898.1 | Aus_I01 | 1 |
| 13 | WAC8380 | I03 | H03_EF108456 | I01 | H01_JN791682.1 | I01 | JX997413.1_H1 | I02 | new isoform | Aus_I01 | 2 |
| 14 | WAC1194 | I03 | H03_EF108456 | I01 | H01_JN791682.1 | I01 | JX997413.1_H1 | I02 | new isoform | Aus_I01 | 2 |
| 15 | WAC4307 | I03 | H03_EF108456 | I01 | H01_JN791682.1 | I01 | JX997413.1_H1 | I02 | new isoform | Aus_I01 | 2 |
| 16 | WAC1495 | I03 | H03_EF108456 | I01 | H01_JN791682.1 | I01 | JX997413.1_H1 | I01 | MW715921.1, MW715898.1 | Aus_I01 | 1 |
| 17 | WAC1496 | I02 | H02_EF108458 | I02 | H02_JN791683.1 | I01 | JX997413.1_H1 | I01 | MW715921.1, MW715898.1 | Aus_I01 | 2 |
| 18 | WAC1497 | I03 | H03_EF108456 | I01 | H01_JN791682.1 | I01 | JX997413.1_H1 | I02 | new isoform | Aus_I01 | 2 |
| 19 | WAC1498 | I03 | H03_EF108456 | I01 | H01_JN791682.1 | I01 | JX997413.1_H1 | I02 | new isoform | Aus_I01 | 2 |
| 20 | WAC1500 | I03 | H03_EF108456 | I01 | H01_JN791682.1 | Null | NA | I02 | new isoform | Aus_I01 | 2 |
| 21 | WAC1508 | I03 | H03_EF108456 | I01 | H01_JN791682.1 | I01 | JX997413.1_H1 | I02 | new isoform | Aus_I01 | 1 |
| 22 | WAC1545 | I02 | H02_EF108458 | I01 | H01_JN791682.1 | Null | NA | I02 | new isoform | Aus_I01 | 2 |
| 23 | WAC1546 | I03 | H03_EF108456 | I01 | H01_JN791682.1 | I01 | JX997413.1_H1 | Null | NA | Aus_I01 | 1 |
| 24 | WAC1178 | I03 | H03_EF108456 | I01 | H01_JN791682.1 | I02 | JX997405.1_H2 | I01 | MW715921.1, MW715898.1 | Aus_I01 | 2 |
| 25 | WAC1547 | I03 | H03_EF108456 | I13/I24 | H13_JN791693.1 | I01 | JX997413.1_H1 | I02 | new isoform | Aus_I01 | 2 |
| 26 | WAC1548 | I03 | H03_EF108456 | I01 | H01_JN791682.1 | I01 | JX997413.1_H1 | I01 | MW715921.1, MW715898.1 | Aus_I01 | 1 |
| 27 | WAC1549 | I03 | H03_EF108456 | I01 | H01_JN791682.1 | I01 | JX997413.1_H1 | I02 | new isoform | Aus_I01 | 1 |
| 28 | WAC1550 | I03 | H03_EF108456 | I01 | H01_JN791682.1 | I01 | JX997413.1_H1 | I02 | new isoform | Aus_I01 | 2 |
| 29 | WAC1551 | I03 | H03_EF108456 | I01 | H01_JN791682.1 | I01 | JX997413.1_H1 | I02 | new isoform | Aus_I01 | 2 |
| 30 | WAC1552 | I03 | H03_EF108456 | I01 | H01_JN791682.1 | I01 | JX997413.1_H1 | I01 | MW715921.1, MW715898.1 | Aus_I01 | 2 |
| 31 | WAC1564 | I03 | H03_EF108456 | I01 | H01_JN791682.1 | I01 | JX997413.1_H1 | I01 | MW715921.1, MW715898.1 | Aus_I01 | 1 |
| 32 | WAC1565 | I03 | H03_EF108456 | I01 | H01_JN791682.1 | I01 | JX997413.1_H1 | I02 | new isoform | Aus_I01 | 2 |
| 33 | WAC2217 | I03 | H03_EF108456 | I02 | H12_JN791692.1 | I01 | JX997413.1_H1 | I02 | new isoform | Aus_I01 | 2 |
| 34 | WAC1179 | I03 | H03_EF108456 | I01 | H01_JN791682.1 | I01 | JX997413.1_H1 | I02 | new isoform | Aus_I01 | 2 |
| 35 | WAC2284 | I03 | H03_EF108456 | I01 | H01_JN791682.1 | I01 | JX997413.1_H1 | I01 | MW715921.1, MW715898.1 | Aus_I01 | 1 |
| 36 | WAC2285 | I03 | H03_EF108456 | I01 | H01_JN791682.1 | I01 | JX997413.1_H1 | I01 | MW715921.1, MW715898.1 | Aus_I01 | 1 |
| 37 | WAC2286 | I03 | H03_EF108456 | I01 | H01_JN791682.1 | I01 | JX997413.1_H1 | I02 | new isoform | Aus_I01 | 2 |
| 38 | WAC2796 | I02 | H02_EF108458 | I01 | H01_JN791682.1 | I01 | JX997413.1_H1 | I01 | MW715921.1, MW715898.1 | Aus_I01 | 2 |
| 39 | WAC2797 | Null | NA | I01 | H01_JN791682.1 | I01 | JX997413.1_H1 | I02 | new isoform | Aus_I04 | 2 |
| 40 | WAC2798 | I02 | H02_EF108458 | I01 | H01_JN791682.1 | functior | Null | I01 | MW715921.1, MW715898.1 | Aus_I01 | 1 |
| 41 | WAC2802 | I03 | H03_EF108456 | I01 | H01_JN791682.1 | I01 | JX997413.1_H1 | I02 | new isoform | Aus_I01 | 1 |
| 42 | WAC2803 | I03 | H03_EF108456 | I01 | H01_JN791682.1 | I01 | JX997413.1_H1 | I01 | MW715921.1, MW715898.1 | Aus_I01 | 1 |
| 43 | WAC2805 | I03 | H03_EF108456 | I01 | H01_JN791682.1 | I01 | JX997413.1_H1 | Null | NA | Aus_I01 | 1 |
| 44 | WAC2806 | I03 | H03_EF108456 | I01 | H01_JN791682.1 | I01 | JX997413.1_H1 | I02 | new isoform | Aus_I01 | 1 |
| 45 | WAC1188 | I03 | H03_EF108456 | I01 | H01_JN791682.1 | I01 | JX997413.1_H1 | I02 | new isoform | Aus_I01 | 2 |
| 46 | WAC2808 | I03 | H03_EF108456 | I01 | H01_JN791682.1 | I01 | JX997413.1_H1 | Null | NA | Aus_I01 | 1 |
| 47 | WAC2809 | I03 | H03_EF108456 | I01 | H01_JN791682.1 | I01 | JX997413.1_H1 | I02 | new isoform | Aus_I01 | 1 |
| 48 | WAC2817 | I03 | H03_EF108456 | I01 | H01_JN791682.1 | I01 | JX997413.1_H1 | I01 | MW715921.1, MW715898.1 | Aus_I01 | 1 |
| 49 | WAC4292 | I03 | H03_EF108456 | I02 | H02_JN791683.1 | I01 | JX997413.1_H1 | I02 | new isoform | Aus_I01 | 2 |
| 50 | WAC4301 | I03 | H03_EF108456 | I01 | H01_JN791682.1 | I01 | JX997413.1_H1 | Null | NA | Aus_I01 | 1 |
| 51 | WAC4304 | I03 | H03_EF108456 | I01 | H01_JN791682.1 | I01 | JX997413.1_H1 | I01 | MW715921.1, MW715898.1 | Aus_I01 | 1 |
| 52 | WAC4308 | I03 | H03_EF108456 | I01 | H01_JN791682.1 | I01 | JX997413.1_H1 | Null | NA | Aus_I01 | 1 |
| 53 | WAC4310 | I03 | H03_EF108456 | I01 | H01_JN791682.1 | I01 | JX997413.1_H1 | I02 | new isoform | Aus_I01 | 1 |
| 54 | WAC4311 | I03 | H03_EF108456 | I01 | H01_JN791682.1 | I01 | JX997413.1_H1 | I01 | MW715921.1, MW715898.1 | Aus_I01 | 1 |
| 55 | WAC4316 | I03 | H03_EF108456 | I01 | H01_JN791682.1 | I01 | JX997413.1_H1 | I01 | MW715921.1, MW715898.1 | Aus_I01 | 1 |
| 56 | WAC1190 | I03 | H03_EF108456 | I01 | H01_JN791682.1 | I01 | JX997413.1_H1 | I02 | new isoform | Aus_I01 | 2 |
| 57 | WAC4317 | I02 | H02_EF108458 | I01 | H01_JN791682.1 | I01 | JX997413.1_H1 | Null | NA | Aus_I01 | 2 |
| 58 | WAC4318 | Null | NA | I01 | H01_JN791682.1 | I01 | JX997413.1_H1 | I02 | new isoform | Aus_I04 | 2 |
| 59 | WAC8383 | I03 | H03_EF108456 | I13/I24 | H13_JN791693.1 | I01 | JX997413.1_H1 | I02 | new isoform | Aus_I01 | 2 |
| 60 | WAC8387 | I03 | H03_EF108456 | I01 | H01_JN791682.1 | I01 | JX997413.1_H1 | I02 | new isoform | Aus_I01 | 1 |
| 61 | WAC8391 | I03 | H03_EF108456 | I01 | H01_JN791682.1 | I01 | JX997413.1_H1 | I02 | new isoform | Aus_I01 | 1 |
| 62 | WAC8399 | I02 | H02_EF108458 | I01 | H01_JN791682.1 | I01 | JX997413.1_H1 | I01 | MW715921.1, MW715898.1 | Aus_I01 | 2 |
| 63 | WAC1193 | I03 | H03_EF108456 | I01 | H01_JN791682.1 | I01 | JX997413.1_H1 | I02 | new isoform | Aus_I01 | 2 |
| 64 | WAC8417 | I03 | H03_EF108456 | I01 | H01_JN791682.1 | I01 | JX997413.1_H1 | I01 | MW715921.1, MW715898.1 | Aus_I01 | 1 |
| 65 | WAC13864 | I02 | H02_EF108458 | I01 | H01_JN791682.1 | I02 | JX997405.1_H2 | I02 | new isoform | Aus_I01 | 2 |
| 66 | WAC13865 | I03 | H03_EF108456 | I02 | H02_JN791683.1 | I01 | JX997413.1_H1 | I05 | MW715975.1, MW715966.1 | Aus_I01 | 1 |
| 67 | WAC13866 | I03 | H03_EF108456 | I01 | H01_JN791682.1 | I01 | JX997413.1_H1 | I02 | new isoform | Aus_I01 | 1 |
| 68 | WAC13868 | I02 | H02_EF108458 | I01 | H01_JN791682.1 | I02 | JX997405.1_H2 | I02 | new isoform | Aus_I01 | 2 |
| 69 | WAC13870 | I02 | H02_EF108458 | I01 | H01_JN791682.1 | I01 | JX997413.1_H1 | I02 | new isoform | Aus_I01 | 2 |
| 70 | WAC13871 | I02 | H02_EF108458 | I01 | H01_JN791682.1 | I01 | JX997413.1_H1 | I02 | new isoform | Aus_I01 | 2 |
| 71 | WAC1195 | I03 | H03_EF108456 | I01 | H01_JN791682.1 | I01 | JX997413.1_H1 | I02 | new isoform | Aus_I01 | 2 |
| 72 | WAC13872 | I02 | H02_EF108458 | I01 | H01_JN791682.1 | I01 | JX997413.1_H1 | I02 | new isoform | Aus_I01 | 2 |
| 73 | WAC13873 | I02 | H02_EF108458 | I01 | H01_JN791682.1 | I01 | JX997413.1_H1 | I02 | new isoform | Aus_I01 | 2 |
| 74 | WAC13957 | I03 | H03_EF108456 | I01 | H01_JN791682.1 | I01 | JX997413.1_H1 | I01 | MW715921.1, MW715898.1 | Aus_I01 | 1 |
| 75 | WAC13958 | I02 | H02_EF108458 | I01 | H01_JN791682.1 | I01 | JX997413.1_H1 | I01 | MW715921.1, MW715898.1 | Aus_I01 | 2 |
| 76 | WAC13959 | I02 | H02_EF108458 | I01 | H01_JN791682.1 | I01 | JX997413.1_H1 | I01 | MW715921.1, MW715898.1 | Aus_I01 | 2 |
| 77 | WAC13960 | I02 | H02_EF108458 | I01 | H01_JN791682.1 | I01 | JX997413.1_H1 | I01 | MW715921.1, MW715898.1 | Aus_I01 | 2 |

| | | | | | | | | | | | |
|---|---|---|---|---|---|---|---|---|---|---|---|
| 78 | WAC13966 | I03 | H03_EF108456 | I01 | H01_JN791682.1 | I01 | JX997413.1_H1 | I02 | new isoform | Aus_I01 | 1 |
| 79 | WAC13967 | I03 | H03_EF108456 | I01 | H01_JN791682.1 | I01 | JX997413.1_H1 | I02 | new isoform | Aus_I01 | 1 |
| 80 | WAC13969 | I03 | H03_EF108456 | I01 | H01_JN791682.1 | I01 | JX997413.1_H1 | I02 | new isoform | Aus_I01 | 1 |
| 81 | WAC1197 | I03 | H03_EF108456 | I01 | H01_JN791682.1 | I01 | JX997413.1_H1 | I02 | new isoform | Aus_I01 | 2 |
| 82 | WAC13979 | I03 | H03_EF108456 | I01 | H01_JN791682.1 | I02 | JX997405.1_H2 | I01 | MW715921.1, MW715898.1 | Aus_I01 | 1 |
| 83 | WAC14056 | I03 | H03_EF108456 | I01 | H01_JN791682.1 | I01 | JX997413.1_H1 | I02 | new isoform | Aus_I01 | 1 |
| 84 | WAC14057 | I02 | H02_EF108458 | I01 | H01_JN791682.1 | I01 | JX997413.1_H1 | I02 | new isoform | Aus_I02 | 1 |
| 85 | WAC14058 | I03 | H03_EF108456 | I01 | H01_JN791682.1 | I01 | JX997413.1_H1 | I02 | new isoform | Aus_I01 | 1 |
| 86 | WAC14059 | I03 | H03_EF108456 | I01 | H01_JN791682.1 | I01 | JX997413.1_H1 | I02 | new isoform | Aus_I01 | 1 |
| 87 | WAC14060 | I03 | H03_EF108456 | I01 | H01_JN791682.1 | I01 | JX997413.1_H1 | I02 | new isoform | Aus_I01 | 1 |
| 88 | WAC14061 | I03 | H03_EF108456 | I01 | H01_JN791682.1 | I01 | JX997413.1_H1 | I02 | new isoform | Aus_I01 | 1 |
| 89 | WAC14062 | I03 | H03_EF108456 | I01 | H01_JN791682.1 | I02 | JX997405.1_H2 | I01 | MW715921.1, MW715898.1 | Aus_I01 | 1 |
| 90 | WAC1199 | I03 | H03_EF108456 | I01 | H01_JN791682.1 | I01 | JX997413.1_H1 | I02 | new isoform | Aus_I01 | 2 |
| 91 | WAC14066 | I03 | H03_EF108456 | I01 | H01_JN791682.1 | I01 | JX997413.1_H1 | I02 | new isoform | Aus_I01 | 1 |
| 92 | WAC14067 | I02 | H02_EF108458 | I01 | H01_JN791682.1 | I01 | JX997413.1_H1 | I02 | new isoform | Aus_I02 | 1 |
| 93 | WAC14068 | I02 | H02_EF108458 | I01 | H01_JN791682.1 | I02 | JX997405.1_H2 | I02 | new isoform | Aus_I01 | 2 |
| 94 | WAC14137 | I03 | H03_EF108456 | I01 | H01_JN791682.1 | I02 | JX997405.1_H2 | I02 | new isoform | Aus_I01 | 1 |
| 95 | WAC14138 | I03 | H03_EF108456 | I01 | H01_JN791682.1 | I02 | JX997405.1_H2 | I01 | MW715921.1, MW715898.1 | Aus_I01 | 1 |
| 96 | WAC14140 | I03 | H03_EF108456 | I01 | H01_JN791682.1 | I01 | JX997413.1_H1 | I02 | new isoform | Aus_I01 | 1 |
| 97 | WAC14141 | I03 | H03_EF108456 | I01 | H01_JN791682.1 | I01 | JX997413.1_H1 | I02 | new isoform | Aus_I01 | 1 |
| 98 | WAC2216 | I02 | H02_EF108458 | I01 | H01_JN791682.1 | I01 | JX997413.1_H1 | I01 | MW715921.1, MW715898.1 | Aus_I01 | 1 |
| 99 | WAC8384 | I03 | H03_EF108456 | I13/I24 | H13_JN791693.1 | I01 | JX997413.1_H1 | I02 | new isoform | Aus_I01 | 2 |
| 100 | 15FG102 | I02 | H02_EF108458 | I01 | H01_JN791682.1 | I01 | JX997413.1_H1 | I02 | new isoform | Aus_I01 | 2 |
| 101 | 15FG103 | I02 | H02_EF108458 | I01 | H01_JN791682.1 | I01 | JX997413.1_H1 | I02 | new isoform | Aus_I01 | 2 |
| 102 | 15FG104 | I03 | H03_EF108456 | I01 | H01_JN791682.1 | I01 | JX997413.1_H1 | I02 | new isoform | Aus_I01 | 2 |
| 103 | 15FG105 | I02 | H02_EF108458 | I01 | H01_JN791682.1 | I01 | JX997413.1_H1 | I02 | new isoform | Aus_I01 | 2 |
| 104 | FG106 | I03 | H03_EF108456 | I01 | H01_JN791682.1 | I02 | JX997405.1_H2 | I02 | new isoform | Aus_I01 | 2 |
| 105 | FG107 | I03 | H03_EF108456 | I01 | H01_JN791682.1 | I02 | JX997405.1_H2 | I02 | new isoform | Aus_I01 | 1 |
| 106 | FG108 | I03 | H03_EF108456 | I02 | H02_JN791683.1 | I01 | JX997413.1_H1 | I02 | new isoform | Aus_I01 | 1 |
| 107 | 15FG107 | I03 | H03_EF108456 | I01 | H01_JN791682.1 | I01 | JX997413.1_H1 | I02 | new isoform | Aus_I01 | 1 |
| 108 | 15FG109 | I03 | H03_EF108456 | I01 | H01_JN791682.1 | I01 | JX997413.1_H1 | I02 | new isoform | Aus_I01 | 1 |
| 109 | 15FG110 | I03 | H03_EF108456 | I01 | H01_JN791682.1 | I01 | JX997413.1_H1 | I01 | MW715921.1, MW715898.1 | Aus_I01 | 1 |
| 110 | WAC8390 | I03 | H03_EF108456 | I01 | H01_JN791682.1 | I01 | JX997413.1_H1 | I02 | new isoform | Aus_I01 | 2 |
| 111 | 15FG112 | I02 | H02_EF108458 | I01 | H01_JN791682.1 | I01 | JX997413.1_H1 | I02 | new isoform | Aus_I01 | 1 |
| 112 | 15FG113 | I02 | H02_EF108458 | I01 | H01_JN791682.1 | I01 | JX997413.1_H1 | I02 | new isoform | Aus_I01 | 1 |
| 113 | 15FG114 | I03 | H03_EF108456 | I02 | H12_JN791692.1 | I01 | JX997413.1_H1 | I02 | new isoform | Aus_I01 | 2 |
| 114 | 15FG115 | I03 | H03_EF108456 | I01 | H01_JN791682.1 | I01 | JX997413.1_H1 | I02 | new isoform | Aus_I01 | 1 |
| 115 | 15FG116 | I03 | H03_EF108456 | I02 | H12_JN791692.1 | I01 | JX997413.1_H1 | I02 | new isoform | Aus_I01 | 2 |
| 116 | 15FG117 | I03 | H03_EF108456 | I02 | H12_JN791692.1 | I01 | JX997413.1_H1 | I02 | new isoform | Aus_I01 | 2 |
| 117 | 15FG118 | I03 | H03_EF108456 | I01 | H01_JN791682.1 | I01 | JX997413.1_H1 | I01 | MW715921.1, MW715898.1 | Aus_I01 | 2 |
| 118 | 15FG119 | I03 | H03_EF108456 | I01 | H01_JN791682.1 | I01 | JX997413.1_H1 | I01 | MW715921.1, MW715898.1 | Aus_I01 | 2 |
| 119 | 15FG120 | I02 | H02_EF108458 | I02 | H02_JN791683.1 | I01 | JX997413.1_H1 | I01 | MW715921.1, MW715898.1 | Aus_I01 | 1 |
| 120 | 15FG04 | I03 | H03_EF108456 | I01 | H01_JN791682.1 | I01 | JX997413.1_H1 | I02 | new isoform | Aus_I01 | 2 |
| 121 | WAC8410 | I02 | H02_EF108458 | I01 | H01_JN791682.1 | I02 | JX997405.1_H2 | I02 | new isoform | Aus_I01 | 2 |
| 122 | 15FG28 | I03 | H03_EF108456 | I01 | H01_JN791682.1 | I01 | JX997413.1_H1 | I02 | new isoform | Aus_I01 | 1 |
| 123 | 15FG33 | I03 | H03_EF108456 | I01 | H01_JN791682.1 | I01 | JX997413.1_H1 | I01 | MW715921.1, MW715898.1 | Aus_I01 | 2 |
| 124 | 15FG37 | I02 | H02_EF108458 | I01 | H01_JN791682.1 | I01 | JX997413.1_H1 | I02 | new isoform | Aus_I04 | 1 |
| 125 | 15FG38 | I03 | H03_EF108456 | I01 | H01_JN791682.1 | I01 | JX997413.1_H1 | I02 | new isoform | Aus_I01 | 2 |
| 126 | 15FG47 | I03 | H03_EF108456 | I02 | H12_JN791692.1 | I01 | JX997413.1_H1 | I02 | new isoform | Aus_I01 | 2 |
| 127 | 15FG49 | I03 | H03_EF108456 | I02 | H12_JN791692.1 | I01 | JX997413.1_H1 | I02 | new isoform | Aus_I01 | 2 |
| 128 | 15FG226 | I03 | H03_EF108456 | I01 | H01_JN791682.1 | I01 | JX997413.1_H1 | I01 | MW715921.1, MW715898.1 | Aus_I01 | 1 |
| 129 | WAC8635 | I03 | H03_EF108456 | I01 | H01_JN791682.1 | I01 | JX997413.1_H1 | Null | NA | Aus_I01 | 1 |
| 130 | 15FG229 | I02 | H02_EF108458 | I01 | H01_JN791682.1 | I01 | JX997413.1_H1 | I02 | new isoform | Aus_I01 | 2 |
| 131 | 15FG237 | I02 | H02_EF108458 | I01 | H01_JN791682.1 | I01 | JX997413.1_H1 | I02 | new isoform | Aus_I01 | 2 |
| 132 | Northam_Mace2 | I02 | H02_EF108458 | I02 | H02_JN791683.1 | I01 | JX997413.1_H1 | I02 | new isoform | Aus_I01 | 1 |
| 133 | Northam_Magen1 | I03 | H03_EF108456 | I01 | H01_JN791682.1 | I01 | JX997413.1_H1 | I02 | new isoform | Aus_I01 | 2 |
| 134 | Northam_Emu1 | I03 | H03_EF108456 | I01 | H01_JN791682.1 | I02 | JX997405.1_H2 | I02 | new isoform | Aus_I01 | 2 |
| 135 | Northam_Emu2 | I03 | H03_EF108456 | I02 | H02_JN791683.1 | I01 | JX997413.1_H1 | I01 | MW715921.1, MW715898.1 | Aus_I01 | 2 |
| 136 | Northam_Mace1 | I03 | H03_EF108456 | I01 | H01_JN791682.1 | I01 | JX997413.1_H1 | I02 | new isoform | Aus_I01 | 1 |
| 137 | WAC9178 | I03 | H03_EF108456 | I01 | H01_JN791682.1 | I01 | JX997413.1_H1 | Null | NA | Aus_I01 | 1 |
| 138 | FG_W003_5 | I02 | H02_EF108458 | I01 | H01_JN791682.1 | I01 | JX997413.1_H1 | I02 | new isoform | Aus_I01 | 2 |
| 139 | Nor_RAC2182_1 | I02 | H02_EF108458 | I02 | H12_JN791692.1 | I01 | JX997413.1_H1 | I02 | new isoform | Aus_I01 | 2 |
| 140 | Northam_WGT | I03 | H03_EF108456 | I01 | H01_JN791682.1 | I01 | JX997413.1_H1 | I02 | new isoform | Aus_I01 | 1 |
| 141 | Nor_RAC2182_2 | I03 | H03_EF108456 | I01 | H01_JN791682.1 | I02 | JX997405.1_H2 | I02 | new isoform | Aus_I01 | 2 |
| 142 | WAC13418 | I03 | H03_EF108456 | I01 | H01_JN791682.1 | I01 | JX997413.1_H1 | Null | NA | Aus_I01 | 1 |
| 143 | SN15 | I03 | H03_EF108456 | I01 | H01_JN791682.1 | I01 | JX997413.1_H1 | Null | NA | Aus_I01 | 1 |
| 144 | WAC13068 | I03 | H03_EF108456 | I01 | H01_JN791682.1 | I01 | JX997413.1_H1 | I01 | MW715921.1, MW715898.1 | Aus_I01 | 1 |
| 145 | WAC13069 | I03 | H03_EF108456 | I01 | H01_JN791682.1 | I01 | JX997413.1_H1 | I01 | MW715921.1, MW715898.1 | Aus_I01 | 1 |
| 146 | WAC13070 | I02 | H02_EF108458 | I01 | H01_JN791682.1 | I01 | JX997413.1_H1 | I01 | MW715921.1, MW715898.1 | Aus_I01 | 2 |
| 147 | WAC2285 | I03 | H03_EF108456 | I01 | H01_JN791682.1 | I01 | JX997413.1_H1 | I01 | MW715921.1, MW715898.1 | Aus_I01 | 1 |
| 148 | WAC13071 | I03 | H03_EF108456 | I01 | H01_JN791682.1 | I01 | JX997413.1_H1 | I02 | new isoform | Aus_I01 | 1 |
| 149 | WAC13072 | I03 | H03_EF108456 | I02 | H02_JN791683.1 | I01 | JX997413.1_H1 | I02 | new isoform | Aus_I01 | 2 |
| 150 | WAC13073 | I03 | H03_EF108456 | I02 | H02_JN791683.1 | I01 | JX997413.1_H1 | I02 | new isoform | Aus_I01 | 2 |
| 151 | WAC13074 | I02 | H02_EF108458 | I01 | H01_JN791682.1 | I01 | JX997413.1_H1 | I01 | MW715921.1, MW715898.1 | Aus_I01 | 2 |
| 152 | WAC13075 | I02 | H02_EF108458 | I01 | H01_JN791682.1 | I01 | JX997413.1_H1 | I01 | MW715921.1, MW715898.1 | Aus_I01 | 2 |
| 153 | WAC13076 | I02 | H02_EF108458 | I01 | H01_JN791682.1 | I01 | JX997413.1_H1 | I01 | MW715921.1, MW715898.1 | Aus_I01 | 2 |
| 154 | WAC13077 | I03 | H03_EF108456 | I02 | H02_JN791683.1 | I01 | JX997413.1_H1 | I02 | new isoform | Aus_I01 | 2 |
| 155 | Meck1 | I03 | H03_EF108456 | I01 | H01_JN791682.1 | I01 | JX997413.1_H1 | I02 | new isoform | Aus_I01 | 1 |
| 156 | Meck3 | I03 | H03_EF108456 | I01 | H01_JN791682.1 | I01 | JX997413.1_H1 | I02 | new isoform | Aus_I01 | 2 |
| 157 | Meck5 | I03 | H03_EF108456 | I01 | H01_JN791682.1 | I01 | JX997413.1_H1 | I02 | new isoform | Aus_I01 | 1 |
| 158 | WAC2810 | I03 | H03_EF108456 | I01 | H01_JN791682.1 | I01 | JX997413.1_H1 | Null | NA | Aus_I01 | 1 |

| | | | | | | | | | | | |
|---|---|---|---|---|---|---|---|---|---|---|---|
| 159 | Meck6 | I03 | H03_EF108456 | I01 | H01_JN791682.1 | I02 | JX997405.1_H2 | I02 | new isoform | Aus_I01 | 1 |
| 160 | Meck8 | I03 | H03_EF108456 | I02 | H02_JN791683.1 | I01 | JX997413.1_H1 | I02 | new isoform | Aus_I01 | 1 |
| 161 | S1FT3A | I03 | H03_EF108456 | I01 | H01_JN791682.1 | I01 | JX997413.1_H1 | I02 | new isoform | Aus_I01 | 1 |
| 162 | S4FT3A | I03 | H03_EF108456 | I01 | H01_JN791682.1 | I01 | JX997413.1_H1 | I02 | new isoform | Aus_I01 | 2 |
| 163 | S1FT3B | I03 | H03_EF108456 | I02 | H02_JN791683.1 | I01 | JX997413.1_H1 | I02 | new isoform | Aus_I01 | 1 |
| 164 | Gerald1 | I03 | H03_EF108456 | I01 | H01_JN791682.1 | I01 | JX997413.1_H1 | I02 | new isoform | Aus_I01 | 1 |
| 165 | Gerald4 | I02 | H02_EF108458 | I02 | H02_JN791683.1 | I01 | JX997413.1_H1 | I02 | new isoform | Aus_I01 | 1 |
| 166 | Mur_51 | I02 | H02_EF108458 | I01 | H01_JN791682.1 | I01 | JX997413.1_H1 | I02 | new isoform | Aus_I01 | 1 |
| 167 | WAC13402 | I03 | H03_EF108456 | I02 | H02_JN791683.1 | I01 | JX997413.1_H1 | I02 | new isoform | Aus_I01 | 2 |
| 168 | WAC2813 | I03 | H03_EF108456 | I01 | H01_JN791682.1 | I01 | JX997413.1_H1 | Null | NA | Aus_I01 | 1 |
| 169 | WAC13403 | I02 | H02_EF108458 | I01 | H01_JN791682.1 | I01 | JX997413.1_H1 | I01 | MW715921.1, MW715898.1 | Aus_I01 | 2 |
| 170 | WAC13404 | I02 | H02_EF108458 | I01 | H01_JN791682.1 | I01 | JX997413.1_H1 | I01 | MW715921.1, MW715898.1 | Aus_I01 | 2 |
| 171 | WAC13405 | I03 | H03_EF108456 | I01 | H01_JN791682.1 | I01 | JX997413.1_H1 | Null | NA | Aus_I01 | 1 |
| 172 | WAC13443 | I03 | H03_EF108456 | I02 | H02_JN791683.1 | I01 | JX997413.1_H1 | I02 | new isoform | Aus_I01 | 2 |
| 173 | WAC13446 | I02 | H02_EF108458 | I01 | H01_JN791682.1 | I01 | JX997413.1_H1 | I01 | MW715921.1, MW715898.1 | Aus_I01 | 2 |
| 174 | WAC13447 | I03 | H03_EF108456 | I01 | H01_JN791682.1 | I01 | JX997413.1_H1 | Null | NA | Aus_I01 | 1 |
| 175 | WAC13523 | I03 | H03_EF108456 | I01 | H01_JN791682.1 | I01 | JX997413.1_H1 | I01 | MW715921.1, MW715898.1 | Aus_I01 | 1 |
| 176 | WAC13524 | I03 | H03_EF108456 | I01 | H01_JN791682.1 | I01 | JX997413.1_H1 | I01 | MW715921.1, MW715898.1 | Aus_I01 | 1 |
| 177 | WAC13525 | I02 | H02_EF108458 | I01 | H01_JN791682.1 | I01 | JX997413.1_H1 | I01 | MW715921.1, MW715898.1 | Aus_I01 | 2 |
| 178 | WAC13526 | I02 | H02_EF108458 | I01 | H01_JN791682.1 | I01 | JX997413.1_H1 | I01 | MW715921.1, MW715898.1 | Aus_I01 | 2 |
| 179 | WAC4303 | I03 | H03_EF108456 | I01 | H01_JN791682.1 | I01 | JX997413.1_H1 | I02 | new isoform | Aus_I01 | 2 |
| 180 | WAC13527 | I03 | H03_EF108456 | I01 | H01_JN791682.1 | I01 | JX997413.1_H1 | I01 | MW715921.1, MW715898.1 | Aus_I01 | 1 |
| 181 | WAC13528 | I02 | H02_EF108458 | I01 | H01_JN791682.1 | I01 | JX997413.1_H1 | I01 | MW715921.1, MW715898.1 | Aus_I01 | 2 |
| 182 | WAC13529 | I03 | H03_EF108456 | I01 | H01_JN791682.1 | I01 | JX997413.1_H1 | I01 | MW715921.1, MW715898.1 | Aus_I01 | 1 |
| 183 | WAC13530 | I03 | H03_EF108456 | I01 | H01_JN791682.1 | I01 | JX997413.1_H1 | I01 | MW715921.1, MW715898.1 | Aus_I01 | 1 |
| 184 | WAC13531 | I03 | H03_EF108456 | I01 | H01_JN791682.1 | I01 | JX997413.1_H1 | I01 | MW715921.1, MW715898.1 | Aus_I01 | 1 |
| 185 | WAC13532 | I02 | H02_EF108458 | I01 | H01_JN791682.1 | I01 | JX997413.1_H1 | I01 | MW715921.1, MW715898.1 | Aus_I01 | 2 |
| 186 | WAC13615 | I02 | H02_EF108458 | I01 | H01_JN791682.1 | I01 | JX997413.1_H1 | I01 | MW715921.1, MW715898.1 | Aus_I01 | 2 |
| 187 | WAC13616 | I02 | H02_EF108458 | I01 | H01_JN791682.1 | I01 | JX997413.1_H1 | I01 | MW715921.1, MW715898.1 | Aus_I01 | 2 |
| 188 | WAC13617 | I02 | H02_EF108458 | I01 | H01_JN791682.1 | I01 | JX997413.1_H1 | I01 | MW715921.1, MW715898.1 | Aus_I01 | 2 |
| 189 | WAC13630 | I02 | H02_EF108458 | I01 | H01_JN791682.1 | I01 | JX997413.1_H1 | I01 | MW715921.1, MW715898.1 | Aus_I01 | 2 |
| 190 | WAC4321 | I03 | H03_EF108456 | I01 | H01_JN791682.1 | I01 | JX997413.1_H1 | functio | NA | Aus_I01 | 2 |
| 191 | WAC13631 | I02 | H02_EF108458 | I01 | H01_JN791682.1 | I01 | JX997413.1_H1 | I01 | MW715921.1, MW715898.1 | Aus_I01 | 2 |
| 192 | WAC13632 | I02 | H02_EF108458 | I01 | H01_JN791682.1 | I01 | JX997413.1_H1 | I01 | MW715921.1, MW715898.1 | Aus_I01 | 2 |
| 193 | WAC13666 | I03 | H03_EF108456 | I01 | H01_JN791682.1 | I01 | JX997413.1_H1 | I02 | new isoform | Aus_I01 | 1 |
| 194 | WAC13667 | I03 | H03_EF108456 | I01 | H01_JN791682.1 | I01 | JX997413.1_H1 | I02 | new isoform | Aus_I01 | 1 |
| 195 | WAC13690 | I02 | H02_EF108458 | I01 | H01_JN791682.1 | I02 | JX997405.1_H2 | I02 | new isoform | Aus_I01 | 2 |
| 196 | WAC13691 | I02 | H02_EF108458 | I01 | H01_JN791682.1 | I02 | JX997405.1_H2 | I02 | new isoform | Aus_I01 | 2 |
| 197 | 206FG226 | I03 | H03_EF108456 | I01 | H01_JN791682.1 | I01 | JX997413.1_H1 | I01 | MW715921.1, MW715898.1 | Aus_I01 | 2 |
| 198 | 205FG215_1 | I03 | H03_EF108456 | I01 | H01_JN791682.1 | I01 | JX997413.1_H1 | I02 | new isoform | Aus_I01 | 1 |
| 199 | 201FG209 | Null | NA | I01 | H01_JN791682.1 | I01 | JX997413.1_H1 | I01 | MW715921.1, MW715898.1 | Aus_I01 | 1 |
| 200 | 201FG211 | I02 | H02_EF108458 | I01 | H01_JN791682.1 | I02 | JX997405.1_H2 | I02 | new isoform | Aus_I01 | 2 |
| 201 | WAC4319 | I03 | H03_EF108456 | I01 | H01_JN791682.1 | I01 | JX997413.1_H1 | I01 | MW715921.1, MW715898.1 | Aus_I01 | 1 |
| 202 | 202FG212 | I03 | H03_EF108456 | I01 | H01_JN791682.1 | I01 | JX997413.1_H1 | I02 | new isoform | Aus_I01 | 1 |
| 203 | 204FG221 | I03 | H03_EF108456 | I01 | H01_JN791682.1 | I02 | JX997405.1_H2 | I02 | new isoform | Aus_I01 | 2 |
| 204 | 204FG223 | I03 | H03_EF108456 | I01 | H01_JN791682.1 | I01 | JX997413.1_H1 | I02 | new isoform | Aus_I01 | 2 |
| 205 | 205FG225 | I03 | H03_EF108456 | I01 | H01_JN791682.1 | I01 | JX997413.1_H1 | I02 | new isoform | Aus_I01 | 1 |
| 206 | 206FG227 | I03 | H03_EF108456 | I01 | H01_JN791682.1 | I01 | JX997413.1_H1 | I02 | new isoform | Aus_I01 | 1 |
| 207 | 205FG216 | I03 | H03_EF108456 | I01 | H01_JN791682.1 | I01 | JX997413.1_H1 | I02 | new isoform | Aus_I01 | 1 |
| 208 | 201FG219 | I03 | H03_EF108456 | I01 | H01_JN791682.1 | I01 | JX997413.1_H1 | I02 | new isoform | Aus_I01 | 2 |
| 209 | 201FG218 | I03 | H03_EF108456 | I01 | H01_JN791682.1 | I02 | JX997405.1_H2 | I02 | new isoform | Aus_I01 | 2 |
| 210 | 204FG214 | I02 | H02_EF108458 | I01 | H01_JN791682.1 | I01 | JX997413.1_H1 | I01 | MW715921.1, MW715898.1 | Aus_I04 | 2 |
| 211 | 903FG214 | I02 | H02_EF108458 | I01 | H01_JN791682.1 | Null | NA | I02 | new isoform | Aus_I01 | 2 |
| 212 | WAC4808 | I03 | H03_EF108456 | I01 | H01_JN791682.1 | I01 | JX997413.1_H1 | I02 | new isoform | Aus_I01 | 2 |
| 213 | 201FG49 | I03 | H03_EF108456 | I01 | H01_JN791682.1 | I01 | JX997413.1_H1 | I01 | MW715921.1, MW715898.1 | Aus_I01 | 1 |
| 214 | 203FG58 | I02 | H02_EF108458 | I01 | H01_JN791682.1 | I01 | JX997413.1_H1 | I01 | MW715921.1, MW715898.1 | Aus_I01 | 2 |
| 215 | 205FG63 | I01 | H01_EF108451 | I01 | H01_JN791682.1 | Null | NA | I05 | MW715975.1, MW715966.1 | Aus_I01 | 1 |
| 216 | 206FG66 | I02 | H02_EF108458 | I01 | H01_JN791682.1 | I01 | JX997413.1_H1 | I01 | MW715921.1, MW715898.1 | Aus_I01 | 1 |
| 217 | 206FG67 | I03 | H03_EF108456 | I01 | H01_JN791682.1 | I01 | JX997413.1_H1 | I02 | new isoform | Aus_I01 | 1 |
| 218 | 206FG68 | I03 | H03_EF108456 | I01 | H01_JN791682.1 | I01 | JX997413.1_H1 | I02 | new isoform | Aus_I01 | 2 |
| 219 | 205FG142 | I02 | H02_EF108458 | I01 | H01_JN791682.1 | I01 | JX997413.1_H1 | I02 | new isoform | Aus_I01 | 2 |
| 220 | 53FG143_1 | I03 | H03_EF108456 | I01 | H01_JN791682.1 | I01 | JX997413.1_H1 | I02 | new isoform | Aus_I01 | 1 |
| 221 | 201FG208 | I02 | H02_EF108458 | I01 | H01_JN791682.1 | Null | NA | I02 | new isoform | Aus_I01 | 2 |
| 222 | 203FG213 | I01 | H01_EF108451 | I01 | H01_JN791682.1 | I01 | JX997413.1_H1 | I05 | MW715975.1, MW715966.1 | Aus_I01 | 2 |
| 223 | WAC4648 | I03 | H03_EF108456 | I01 | H01_JN791682.1 | I01 | JX997413.1_H1 | I01 | MW715921.1, MW715898.1 | Aus_I01 | 1 |
| 224 | 204FG222 | I02 | H02_EF108458 | I01 | H01_JN791682.1 | I01 | JX997413.1_H1 | I02 | new isoform | Aus_I01 | 2 |
| 225 | 205FG410 | I02 | H02_EF108458 | I01 | H01_JN791682.1 | I01 | JX997413.1_H1 | I01 | MW715921.1, MW715898.1 | Aus_I01 | 1 |
| 226 | 202FG414 | I03 | H03_EF108456 | I01 | H01_JN791682.1 | I01 | JX997413.1_H1 | I02 | new isoform | Aus_I01 | 1 |
| 227 | WAC739 | I03 | H03_EF108456 | I01 | H01_JN791682.1 | I02 | JX997405.1_H2 | I01 | MW715921.1, MW715898.1 | Aus_I01 | 2 |
| 228 | WAC740 | I03 | H03_EF108456 | I01 | H01_JN791682.1 | I01 | JX997413.1_H1 | I02 | new isoform | Aus_I01 | 1 |
| 229 | WAC741 | I03 | H03_EF108456 | I01 | H01_JN791682.1 | I02 | JX997405.1_H2 | I01 | MW715921.1, MW715898.1 | Aus_I01 | 2 |
| 230 | 15FG99 | I03 | H03_EF108456 | I01 | H01_JN791682.1 | I01 | JX997413.1_H1 | I02 | new isoform | Aus_I01 | 1 |
| 231 | 15FG100 | I02 | H02_EF108458 | I01 | H01_JN791682.1 | I01 | JX997413.1_H1 | I02 | new isoform | Aus_I01 | 2 |
| 232 | 15FG101 | I02 | H02_EF108458 | I02 | H02_JN791683.1 | I01 | JX997413.1_H1 | I02 | new isoform | Aus_I01 | 1 |
| 233 | 16FG06 | I03 | H03_EF108456 | I01 | H01_JN791682.1 | I01 | JX997413.1_H1 | I02 | new isoform | Aus_I01 | 1 |
| 234 | 16FG158 | I03 | H03_EF108456 | I01 | H01_JN791682.1 | I02 | JX997405.1_H2 | I02 | new isoform | Aus_I01 | 1 |
| 235 | 16FG159 | I02 | H02_EF108458 | I01 | H01_JN791682.1 | I01 | JX997413.1_H1 | I02 | new isoform | Aus_I01 | 2 |
| 236 | 16FG160 | I03 | H03_EF108456 | I01 | H01_JN791682.1 | I02 | JX997405.1_H2 | I02 | new isoform | Aus_I01 | 1 |
| 237 | 16FG161 | I03 | H03_EF108456 | I01 | H01_JN791682.1 | I02 | JX997405.1_H2 | I02 | new isoform | Aus_I01 | 1 |
| 238 | 16FG162 | I03 | H03_EF108456 | I01 | H01_JN791682.1 | I02 | JX997405.1_H2 | I02 | new isoform | Aus_I01 | 1 |
| 239 | 16FG163_2 | I03 | H03_EF108456 | I01 | H01_JN791682.1 | I02 | JX997405.1_H2 | I02 | new isoform | Aus_I01 | 1 |

| | | | | | | | | | | | |
|---|---|---|---|---|---|---|---|---|---|---|---|
| 240 | 16FG164 | I03 | H03_EF108456 | I01 | H01_JN791682.1 | I02 | JX997405.1_H2 | I02 | new isoform | Aus_I01 | 1 |
| 241 | 16FG165 | I03 | H03_EF108456 | I01 | H01_JN791682.1 | I02 | JX997405.1_H2 | I02 | new isoform | Aus_I01 | 1 |
| 242 | 16FG166 | I03 | H03_EF108456 | I01 | H01_JN791682.1 | I02 | JX997405.1_H2 | I02 | new isoform | Aus_I01 | 1 |
| 243 | 16FG167 | I03 | H03_EF108456 | I01 | H01_JN791682.1 | I02 | JX997405.1_H2 | I02 | new isoform | Aus_I01 | 1 |
| 244 | 16FG168 | I03 | H03_EF108456 | I01 | H01_JN791682.1 | I02 | JX997405.1_H2 | I02 | new isoform | Aus_I01 | 1 |
| 245 | 16FG169 | I03 | H03_EF108456 | I01 | H01_JN791682.1 | I02 | JX997405.1_H2 | I02 | new isoform | Aus_I01 | 1 |
| 246 | WAC13955 | I03 | H03_EF108456 | I01 | H01_JN791682.1 | I01 | JX997413.1_H1 | I02 | new isoform | Aus_I04 | 2 |
| 247 | 16FG170 | I03 | H03_EF108456 | I01 | H01_JN791682.1 | I02 | JX997405.1_H2 | I02 | new isoform | Aus_I01 | 1 |
| 248 | 16FG171 | I03 | H03_EF108456 | I01 | H01_JN791682.1 | I02 | JX997405.1_H2 | I02 | new isoform | Aus_I01 | 1 |
| 249 | Mur_S3 | I03 | H03_EF108456 | I01 | H01_JN791682.1 | I01 | JX997413.1_H1 | I02 | new isoform | Aus_I01 | 2 |
| 250 | 14FG141 | I02 | H02_EF108458 | I13/I24 | H13_JN791693.1 | I01 | JX997413.1_H1 | I02 | new isoform | Aus_I01 | 2 |
| 251 | PN260 | I03 | H03_EF108456 | I01 | H01_JN791682.1 | I01 | JX997413.1_H1 | I02 | new isoform | Aus_I01 | 2 |
| 252 | PN261 | I03 | H03_EF108456 | I01 | H01_JN791682.1 | I01 | JX997413.1_H1 | I02 | new isoform | Aus_I01 | 1 |
| 253 | PN262 | I01 | H01_EF108451 | I01 | H01_JN791682.1 | I01 | JX997413.1_H1 | I05 | MW715975.1, MW715966.1 | Aus_I01 | 2 |
| 254 | PN263 | I03 | H03_EF108456 | I01 | H01_JN791682.1 | I02 | JX997405.1_H2 | I01 | MW715921.1, MW715898.1 | Aus_I01 | 1 |
| 255 | PN264 | I03 | H03_EF108456 | I01 | H01_JN791682.1 | I01 | JX997413.1_H1 | I02 | new isoform | Aus_I01 | 1 |
| 256 | PN265 | I01 | H01_EF108451 | I01 | H01_JN791682.1 | I01 | JX997413.1_H1 | I02 | new isoform | Aus_I01 | 2 |
| 257 | PN266 | I03 | H03_EF108456 | I13/I24 | H13_JN791693.1 | I01 | JX997413.1_H1 | I02 | new isoform | Aus_I01 | 1 |
| 258 | PN268 | I02 | H02_EF108458 | I01 | H01_JN791682.1 | I01 | JX997413.1_H1 | I01 | MW715921.1, MW715898.1 | Aus_I01 | 1 |
| 259 | PN269 | I02 | H02_EF108458 | I01 | H01_JN791682.1 | I01 | JX997413.1_H1 | I02 | new isoform | Aus_I01 | 2 |
| 260 | PN270 | I03 | H03_EF108456 | I02 | H02_JN791683.1 | I02 | JX997405.1_H2 | I05 | MW715975.1, MW715966.1 | Aus_I01 | 2 |
| 261 | PN271 | I03 | H03_EF108456 | I01 | H01_JN791682.1 | I01 | JX997413.1_H1 | I02 | new isoform | Aus_I01 | 2 |
| 262 | PN274 | I03 | H03_EF108456 | I02 | H02_JN791683.1 | I02 | JX997405.1_H2 | I05 | MW715975.1, MW715966.1 | Aus_I01 | 2 |
| 263 | PN275 | I03 | H03_EF108456 | I01 | H01_JN791682.1 | I01 | JX997413.1_H1 | I02 | new isoform | Aus_I04 | 1 |
| 264 | PN276 | I02 | H02_EF108458 | I01 | H01_JN791682.1 | I02 | JX997405.1_H2 | I02 | new isoform | Aus_I01 | 2 |
| 265 | PN277 | I03 | H03_EF108456 | I01 | H01_JN791682.1 | I01 | JX997413.1_H1 | I01 | MW715921.1, MW715898.1 | Aus_I01 | 2 |
| 266 | PN279 | I02 | H02_EF108458 | I01 | H01_JN791682.1 | I01 | JX997413.1_H1 | I02 | new isoform | Aus_I01 | 1 |
| 267 | PN280 | I03 | H03_EF108456 | I01 | H01_JN791682.1 | I01 | JX997413.1_H1 | I02 | new isoform | Aus_I01 | 2 |
| 268 | PN282 | I03 | H03_EF108456 | I01 | H01_JN791682.1 | I01 | JX997413.1_H1 | I02 | new isoform | Aus_I01 | 2 |
| 269 | PN283 | I03 | H03_EF108456 | I01 | H01_JN791682.1 | I01 | JX997413.1_H1 | I02 | new isoform | Aus_I01 | 2 |
| 270 | PN284 | I03 | H03_EF108456 | I01 | H01_JN791682.1 | I02 | JX997405.1_H2 | I02 | new isoform | Aus_I01 | 1 |
| 271 | PN285 | I02 | H02_EF108458 | I01 | H01_JN791682.1 | I01 | JX997413.1_H1 | I02 | new isoform | Aus_I01 | 2 |
| 272 | PN286 | I01 | H01_EF108451 | I01 | H01_JN791682.1 | I01 | JX997413.1_H1 | I05 | MW715975.1, MW715966.1 | Aus_I01 | 2 |
| 273 | PN287 | I02 | H02_EF108458 | I01 | H01_JN791682.1 | I01 | JX997413.1_H1 | Null | NA | Aus_I01 | 1 |
| 274 | PN288 | I03 | H03_EF108456 | I01 | H01_JN791682.1 | I01 | JX997413.1_H1 | I02 | new isoform | Aus_I01 | 1 |
| 275 | PN289 | I03 | H03_EF108456 | I01 | H01_JN791682.1 | I01 | JX997413.1_H1 | I02 | new isoform | Aus_I01 | 1 |
| 276 | PN290 | I01 | H01_EF108451 | I01 | H01_JN791682.1 | I01 | JX997413.1_H1 | I02 | new isoform | Aus_I01 | 2 |
| 277 | PN291 | I03 | H03_EF108456 | I13/I24 | H13_JN791693.1 | I01 | JX997413.1_H1 | I02 | new isoform | Aus_I01 | 1 |
| 278 | PN292 | I03 | H03_EF108456 | I01 | H01_JN791682.1 | I01 | JX997413.1_H1 | I02 | new isoform | Aus_I01 | 2 |
| 279 | PN295 | I02 | H02_EF108458 | I01 | H01_JN791682.1 | I01 | JX997413.1_H1 | I02 | new isoform | Aus_I01 | 2 |
| 280 | PN296 | I02 | H02_EF108458 | I01 | H01_JN791682.1 | I02 | JX997405.1_H2 | I02 | new isoform | Aus_I01 | 2 |
| 281 | PN297 | I01 | H01_EF108451 | I01 | H01_JN791682.1 | I01 | JX997413.1_H1 | I02 | new isoform | Aus_I01 | 1 |
| 282 | PN298 | I02 | H02_EF108458 | I01 | H01_JN791682.1 | I02 | JX997405.1_H2 | I02 | new isoform | Aus_I01 | 2 |
| 283 | PN299 | I03 | H03_EF108456 | I02 | H02_JN791683.1 | I02 | JX997405.1_H2 | I01 | MW715921.1, MW715898.1 | Aus_I05 | 1 |
| 284 | PN300 | I03 | H03_EF108456 | I02 | H02_JN791683.1 | I02 | JX997405.1_H2 | I01 | MW715921.1, MW715898.1 | Aus_I05 | 1 |
| 285 | PN301 | I03 | H03_EF108456 | I02 | H02_JN791683.1 | I02 | JX997405.1_H2 | I01 | MW715921.1, MW715898.1 | Aus_I05 | 1 |
| 286 | PN302 | I03 | H03_EF108456 | I01 | H01_JN791682.1 | I01 | JX997413.1_H1 | I01 | MW715921.1, MW715898.1 | Aus_I01 | 2 |
| 287 | PN303 | I03 | H03_EF108456 | I01 | H01_JN791682.1 | I01 | JX997413.1_H1 | I02 | new isoform | Aus_I01 | 1 |
| 288 | PN304 | I03 | H03_EF108456 | I01 | H01_JN791682.1 | I01 | JX997413.1_H1 | I02 | new isoform | Aus_I01 | 2 |
| 289 | PN305 | I03 | H03_EF108456 | I02 | H02_JN791683.1 | I02 | JX997405.1_H2 | I05 | MW715975.1, MW715966.1 | Aus_I01 | 2 |
| 290 | PN306 | I03 | H03_EF108456 | I01 | H01_JN791682.1 | I01 | JX997413.1_H1 | I02 | new isoform | Aus_I01 | 2 |
| 291 | PN307 | I03 | H03_EF108456 | I01 | H01_JN791682.1 | I01 | JX997413.1_H1 | I02 | new isoform | Aus_I01 | 2 |
| 292 | PN308 | I03 | H03_EF108456 | I02 | H02_JN791683.1 | I02 | JX997405.1_H2 | I02 | new isoform | Aus_I01 | 1 |
| 293 | PN309 | I01 | H01_EF108451 | I01 | H01_JN791682.1 | I01 | JX997413.1_H1 | I02 | new isoform | Aus_I01 | 1 |
| 294 | PN312 | I03 | H03_EF108456 | I01 | H01_JN791682.1 | I01 | JX997413.1_H1 | I01 | MW715921.1, MW715898.1 | Aus_I04 | 2 |
| 295 | PN313 | I03 | H03_EF108456 | I01 | H01_JN791682.1 | I01 | JX997413.1_H1 | I01 | MW715921.1, MW715898.1 | Aus_I04 | 2 |
| 296 | PN314 | I03 | H03_EF108456 | I02 | H02_JN791683.1 | I02 | JX997405.1_H2 | I02 | new isoform | Aus_I01 | 1 |
| 297 | PN315 | I03 | H03_EF108456 | I01 | H01_JN791682.1 | I01 | JX997413.1_H1 | I02 | new isoform | Aus_I01 | 2 |
| 298 | PN316 | I03 | H03_EF108456 | I01 | H01_JN791682.1 | I01 | JX997413.1_H1 | I02 | new isoform | Aus_I01 | 2 |
| 299 | PN317 | I03 | H03_EF108456 | I01 | H01_JN791682.1 | I02 | JX997405.1_H2 | I01 | MW715921.1, MW715898.1 | Aus_I01 | 1 |
| 300 | PN318 | I03 | H03_EF108456 | I01 | H01_JN791682.1 | I01 | JX997413.1_H1 | I02 | new isoform | Aus_I01 | 2 |
| 301 | PN319 | I03 | H03_EF108456 | I01 | H01_JN791682.1 | I01 | JX997413.1_H1 | I02 | new isoform | Aus_I01 | 1 |
| 302 | PN320 | I03 | H03_EF108456 | I01 | H01_JN791682.1 | I02 | JX997405.1_H2 | I02 | new isoform | Aus_I01 | 1 |
| 303 | PN321 | I03 | H03_EF108456 | I01 | H01_JN791682.1 | I01 | JX997413.1_H1 | I02 | new isoform | Aus_I01 | 1 |
| 304 | PN322 | I03 | H03_EF108456 | I01 | H01_JN791682.1 | I01 | JX997413.1_H1 | I02 | new isoform | Aus_I01 | 1 |
| 305 | PN323 | I03 | H03_EF108456 | I01 | H01_JN791682.1 | I01 | JX997413.1_H1 | I02 | new isoform | Aus_I01 | 1 |
| 306 | PN324 | I03 | H03_EF108456 | I01 | H01_JN791682.1 | I01 | JX997413.1_H1 | I02 | new isoform | Aus_I01 | 1 |
| 307 | PN325 | I03 | H03_EF108456 | I02 | H02_JN791683.1 | I02 | JX997405.1_H2 | I02 | new isoform | Aus_I01 | 2 |
| 308 | PN326 | I03 | H03_EF108456 | I02 | H12_JN791692.1 | I01 | JX997413.1_H1 | I02 | new isoform | Aus_I01 | 1 |
| 309 | PN328 | I03 | H03_EF108456 | I01 | H01_JN791682.1 | I01 | JX997413.1_H1 | I02 | new isoform | Aus_I01 | 1 |
| 310 | PN329 | I03 | H03_EF108456 | I01 | H01_JN791682.1 | I01 | JX997413.1_H1 | I02 | new isoform | Aus_I01 | 1 |
| 311 | PN330 | I03 | H03_EF108456 | I02 | H02_JN791683.1 | I01 | JX997413.1_H1 | I01 | MW715921.1, MW715898.1 | Aus_I01 | 1 |
| 312 | PN331 | I03 | H03_EF108456 | I01 | H01_JN791682.1 | I01 | JX997413.1_H1 | I02 | new isoform | Aus_I01 | 1 |
| 313 | PN332 | I03 | H03_EF108456 | I01 | H01_JN791682.1 | I01 | JX997413.1_H1 | I02 | new isoform | Aus_I01 | 2 |
| 314 | PN333 | I03 | H03_EF108456 | I01 | H01_JN791682.1 | I01 | JX997413.1_H1 | I02 | new isoform | Aus_I01 | 1 |
| 315 | PN334 | I03 | H03_EF108456 | I01 | H01_JN791682.1 | I01 | JX997413.1_H1 | I02 | new isoform | Aus_I01 | 2 |
| 316 | PN335 | I03 | H03_EF108456 | I01 | H01_JN791682.1 | I01 | JX997413.1_H1 | I02 | new isoform | Aus_I01 | 2 |
| 317 | PN336 | I03 | H03_EF108456 | I01 | H01_JN791682.1 | I01 | JX997413.1_H1 | I02 | new isoform | Aus_I01 | 1 |
| 318 | PN337 | I03 | H03_EF108456 | I01 | H01_JN791682.1 | I01 | JX997413.1_H1 | I02 | new isoform | Aus_I01 | 2 |
| 319 | PN338 | I03 | H03_EF108456 | I01 | H01_JN791682.1 | I01 | JX997413.1_H1 | I02 | new isoform | Aus_I01 | 2 |
| 320 | PN340 | I03 | H03_EF108456 | I01 | H01_JN791682.1 | I01 | JX997413.1_H1 | I02 | new isoform | Aus_I01 | 2 |

| | | | | | | | | | | | |
|---|---|---|---|---|---|---|---|---|---|---|---|
| 321 | PN341 | I03 | H03_EF108456 | I01 | H01_JN791682.1 | I01 | JX997413.1_H1 | I02 | new isoform | Aus_I01 | 1 |
| 322 | PN342 | I03 | H03_EF108456 | I01 | H01_JN791682.1 | I01 | JX997413.1_H1 | I02 | new isoform | Aus_I01 | 2 |
| 323 | PN343 | I03 | H03_EF108456 | I01 | H01_JN791682.1 | I01 | JX997413.1_H1 | I02 | new isoform | Aus_I01 | 2 |
| 324 | PN344 | I03 | H03_EF108456 | I01 | H01_JN791682.1 | I01 | JX997413.1_H1 | I02 | new isoform | Aus_I01 | 1 |
| 325 | PN345 | I03 | H03_EF108456 | I01 | H01_JN791682.1 | I01 | JX997413.1_H1 | I02 | new isoform | Aus_I01 | 2 |
| 326 | PN346 | I03 | H03_EF108456 | I01 | H01_JN791682.1 | I01 | JX997413.1_H1 | I02 | new isoform | Aus_I01 | 2 |
| 327 | PN347 | I03 | H03_EF108456 | I01 | H01_JN791682.1 | I01 | JX997413.1_H1 | I01 | MW715921.1, MW715898.1 | Aus_I01 | 1 |
| 328 | PN348 | I03 | H03_EF108456 | I01 | H01_JN791682.1 | I02 | JX997405.1_H2 | I02 | new isoform | Aus_I01 | 2 |
| 329 | PN349 | I02 | H02_EF108458 | I02 | H02_JN791683.1 | I01 | JX997413.1_H1 | I01 | MW715921.1, MW715898.1 | Aus_I01 | 2 |
| 330 | PN350 | I03 | H03_EF108456 | I01 | H01_JN791682.1 | I01 | JX997413.1_H1 | I02 | new isoform | Aus_I01 | 1 |
| 331 | PN351 | I02 | H02_EF108458 | I02 | H12_JN791692.1 | I01 | JX997413.1_H1 | I02 | new isoform | Aus_I01 | 1 |
| 332 | PN352 | I03 | H03_EF108456 | I01 | H01_JN791682.1 | I01 | JX997413.1_H1 | I01 | MW715921.1, MW715898.1 | Aus_I01 | 2 |
| 333 | PN353 | I03 | H03_EF108456 | I01 | H01_JN791682.1 | I01 | JX997413.1_H1 | I02 | new isoform | Aus_I01 | 2 |
| 334 | PN354 | I02 | H02_EF108458 | I02 | H02_JN791683.1 | I01 | JX997413.1_H1 | I02 | new isoform | Aus_I01 | 2 |
| 335 | PN355 | I03 | H03_EF108456 | I01 | H01_JN791682.1 | I01 | JX997413.1_H1 | I02 | new isoform | Aus_I01 | 1 |
| 336 | PN356 | I03 | H03_EF108456 | I01 | H01_JN791682.1 | I01 | JX997413.1_H1 | I02 | new isoform | Aus_I01 | 1 |
| 337 | PN357 | I03 | H03_EF108456 | I01 | H01_JN791682.1 | I01 | JX997413.1_H1 | I01 | MW715921.1, MW715898.1 | Aus_I01 | 2 |
| 338 | PN358 | I03 | H03_EF108456 | I01 | H01_JN791682.1 | I01 | JX997413.1_H1 | I02 | new isoform | Aus_I01 | 2 |
| 339 | PN359 | I03 | H03_EF108456 | I01 | H01_JN791682.1 | I01 | JX997413.1_H1 | I02 | new isoform | Aus_I03 | 2 |
| 340 | PN360 | I03 | H03_EF108456 | I01 | H01_JN791682.1 | I01 | JX997413.1_H1 | I01 | MW715921.1, MW715898.1 | Aus_I01 | 2 |
| 341 | PN361 | I03 | H03_EF108456 | I01 | H01_JN791682.1 | I01 | JX997413.1_H1 | I02 | new isoform | Aus_I01 | 1 |
| 342 | PN362 | I03 | H03_EF108456 | I01 | H01_JN791682.1 | I01 | JX997413.1_H1 | I02 | new isoform | Aus_I01 | 1 |
| 343 | PN363 | I03 | H03_EF108456 | I01 | H01_JN791682.1 | I01 | JX997413.1_H1 | I02 | new isoform | Aus_I01 | 1 |
| 344 | PN364 | I03 | H03_EF108456 | I01 | H01_JN791682.1 | I01 | JX997413.1_H1 | I02 | new isoform | Aus_I01 | 2 |
| 345 | PN365 | I03 | H03_EF108456 | I01 | H01_JN791682.1 | I01 | JX997413.1_H1 | I02 | new isoform | Aus_I01 | 1 |
| 346 | PN366 | I02 | H02_EF108458 | I01 | H01_JN791682.1 | I01 | JX997413.1_H1 | I02 | new isoform | Aus_I01 | 2 |
| 347 | PN367 | I03 | H03_EF108456 | I01 | H01_JN791682.1 | I01 | JX997413.1_H1 | I02 | new isoform | Aus_I01 | 1 |
| 348 | PN368 | I03 | H03_EF108456 | I01 | H01_JN791682.1 | I01 | JX997413.1_H1 | I02 | new isoform | Aus_I01 | 1 |
| 349 | PN369 | I03 | H03_EF108456 | I01 | H01_JN791682.1 | I01 | JX997413.1_H1 | I02 | new isoform | Aus_I04 | 2 |
| 350 | PN370 | I03 | H03_EF108456 | I01 | H01_JN791682.1 | I01 | JX997413.1_H1 | I02 | new isoform | Aus_I01 | 1 |
| 351 | PN371 | I03 | H03_EF108456 | I01 | H01_JN791682.1 | I01 | JX997413.1_H1 | I02 | new isoform | Aus_I01 | 1 |
| 352 | PN372 | I02 | H02_EF108458 | I02 | H02_JN791683.1 | I01 | JX997413.1_H1 | I02 | new isoform | Aus_I01 | 2 |
| 353 | PN373 | I03 | H03_EF108456 | I01 | H01_JN791682.1 | I01 | JX997413.1_H1 | I02 | new isoform | Aus_I01 | 1 |
| 354 | PN374 | I03 | H03_EF108456 | I01 | H01_JN791682.1 | I01 | JX997413.1_H1 | I02 | new isoform | Aus_I01 | 1 |
| 355 | PN375 | I03 | H03_EF108456 | I01 | H01_JN791682.1 | I01 | JX997413.1_H1 | I02 | new isoform | Aus_I01 | 1 |
| 356 | PN376 | I03 | H03_EF108456 | I02 | H12_JN791692.1 | I01 | JX997413.1_H1 | I02 | new isoform | Aus_I01 | 2 |
| 357 | PN377 | I03 | H03_EF108456 | I02 | H12_JN791692.1 | I01 | JX997413.1_H1 | I02 | new isoform | Aus_I01 | 2 |
| 358 | PN378 | I02 | H02_EF108458 | I01 | H01_JN791682.1 | I02 | JX997405.1_H2 | I02 | new isoform | Aus_I01 | 2 |
| 359 | PN379 | I03 | H03_EF108456 | I01 | H01_JN791682.1 | I01 | JX997413.1_H1 | I02 | new isoform | Aus_I01 | 2 |
| 360 | PN380 | I03 | H03_EF108456 | I01 | H01_JN791682.1 | I01 | JX997413.1_H1 | I02 | new isoform | Aus_I01 | 1 |

**STable 2: Stagonospora nodorum transposon Molly profiles of 360 isolates**

| Group | Collection | Isolate | Isolate_tree_name | Year of collection | Copy # | Modification | | Molly_copy_number_bwa | Molly_copy_number_bbmap | molly_coverage_bwa | genome_coverage_bwa | molly_coverage_bbmap | genome_coverage_bbmap |
|---|---|---|---|---|---|---|---|---|---|---|---|---|---|
| | | | | | | Strong (>20 SNP) | Weak/intact (0-20 SNP) | | | | | | |
| Reference | OLD_group 3 | Sn15_NCBI | na | 2000 | 1 | 1 | 0 | 0.12 | 1.00 | 3.94 | 32.71 | 34.70 | 34.77 |
| | OLD_group 3 | Sn15_illumina | na | 2000 | 1 | 1 | 0 | 0.11 | 0.99 | 3.94 | 34.59 | 34.70 | 34.97 |
| | USA | Sn2000_NCBI | na | NA | 0 | 0 | 0 | 0.05 | 0.17 | 1.53 | 29.44 | 4.94 | 29.82 |
| | USA | Sn2000_illumina | na | NA | 0 | 0 | 0 | 0.05 | 0.15 | 1.53 | 32.29 | 4.94 | 32.65 |
| | OLD_group 1 | 15FG38_nanopore | O124_15FG38 | 2015 | 19 | 9 | 10 | 19.20 | 21.41 | 652.16 | 33.97 | 762.80 | 35.62 |
| | OLD_group 1 | 15FG38_illumina | O124_15FG38 | 2015 | 19 | 9 | 10 | 18.74 | 21.29 | 652.16 | 34.79 | 762.80 | 35.83 |
| | OLD_group 5 | 16FG168_nanopore | O155_16FG168 | 2016 | 44 | 27 | 17 | 43.97 | 44.91 | 7060.10 | 160.56 | 7370.69 | 164.12 |
| | OLD_group 5 | 16FG168_illumina | O155_16FG168 | 2016 | 44 | 27 | 17 | 43.33 | 44.75 | 7060.10 | 162.94 | 7370.69 | 164.71 |
| | Current_group 7 | PN315_nanopore | C315_PN315 | 2019 | 36 | 9 | 27 | 36.94 | 37.56 | 2175.55 | 58.89 | 2293.29 | 61.05 |
| | Current_group 7 | PN315_illumina | C315_PN315 | 2019 | 36 | 9 | 27 | 35.50 | 37.07 | 2175.55 | 61.28 | 2293.29 | 61.86 |
| Group 1 | DPIRD | WAC1178 | DIP2_WAC1178 | 1968 | 2 | 2 | 0 | 0.36 | 2.16 | 11.37 | 31.74 | 69.45 | 32.10 |
| | DPIRD | WAC1179 | DIP3_WAC1179 | 1968 | 6 | 6 | 0 | 3.79 | 6.02 | 120.32 | 31.72 | 193.41 | 32.10 |
| | DPIRD | WAC 1544 | DIP106_WAC1544 | 1969 | 2 | 2 | 0 | 1.77 | 3.48 | 65.22 | 36.80 | 130.12 | 37.39 |
| | DPIRD | WAC1498 | DIP15_WAC1498 | 1969 | 1 | 1 | 0 | 0.09 | 2.20 | 2.76 | 29.70 | 66.29 | 30.07 |
| | DPIRD | WAC1500 | DIP16_WAC1500 | 1969 | 1 | 1 | 0 | 1.08 | 1.32 | 347.95 | 323.28 | 433.36 | 328.68 |
| | DPIRD | WAC1508 | DIP17_WAC1508 | 1969 | 0 | 0 | 0 | 0.06 | 2.03 | 1.82 | 31.96 | 65.60 | 32.38 |
| | DPIRD | WAC1545 | DIP18_WAC1545 | 1969 | 0 | 0 | 0 | 0.07 | 2.03 | 2.22 | 32.27 | 66.24 | 32.65 |
| | DPIRD | WAC1546 | DIP19_WAC1546 | 1969 | 4 | 4 | 0 | 2.02 | 5.39 | 63.12 | 31.21 | 170.31 | 31.62 |
| | DPIRD | WAC1549 | DIP22_WAC1549 | 1969 | 0 | 0 | 0 | 0.07 | 1.83 | 2.35 | 31.81 | 59.06 | 32.19 |
| | DPIRD | WAC1550 | DIP23_WAC1550 | 1969 | 21 | 0 | 21 | 19.16 | 21.25 | 559.88 | 29.22 | 627.71 | 29.54 |
| | DPIRD | WAC1551 | DIP24_WAC1551 | 1969 | 1 | 1 | 0 | 0.09 | 1.89 | 2.62 | 30.26 | 57.77 | 30.60 |
| | DPIRD | WAC1552 | DIP25_WAC1552 | 1969 | 1 | 1 | 0 | 0.69 | 1.61 | 22.25 | 32.05 | 52.04 | 32.41 |
| | DPIRD | WAC2217 | DIP29_WAC2217 | 1972 | 1 | 1 | 0 | 0.09 | 2.44 | 2.78 | 32.21 | 79.48 | 32.59 |
| | DPIRD | WAC 2812 | DIP103_WAC2812 | 1980 | 3 | 3 | 0 | 2.76 | 4.16 | 103.29 | 37.38 | 157.45 | 37.89 |
| | DPIRD | WAC 2799 | DIP107_WAC2799 | 1980 | 9 | 9 | 0 | 7.22 | 9.85 | 265.64 | 36.80 | 368.54 | 37.42 |
| | DPIRD | WAC2286 | DIP32_WAC2286 | 1980 | 18 | 5 | 13 | 16.64 | 17.92 | 532.99 | 32.03 | 581.51 | 32.45 |
| | DPIRD | WAC2797 | DIP34_WAC2797 | 1980 | 20 | 5 | 15 | 19.30 | 20.40 | 5030.04 | 260.57 | 5432.03 | 266.31 |
| | DPIRD | WAC2798 | DIP35_WAC2798 | 1980 | 34 | 19 | 15 | 28.21 | 33.82 | 891.68 | 31.61 | 1082.74 | 32.02 |
| | DPIRD | WAC2802 | DIP36_WAC2802 | 1980 | 1 | 1 | 0 | 0.87 | 2.78 | 27.45 | 31.45 | 88.53 | 31.86 |
| | DPIRD | WAC2803 | DIP37_WAC2803 | 1980 | 9 | 9 | 0 | 7.17 | 9.70 | 228.89 | 31.94 | 313.40 | 32.32 |
| | DPIRD | WAC2809 | DIP41_WAC2809 | 1980 | 13 | 4 | 9 | 11.44 | 13.47 | 361.01 | 31.57 | 430.71 | 31.97 |
| | DPIRD | WAC 4307 | DIP112_WAC4307 | 1985 | 3 | 3 | 0 | 2.94 | 4.54 | 92.39 | 31.46 | 144.64 | 31.85 |
| | DPIRD | WAC4292 | DIP43_WAC4292 | 1985 | 0 | 0 | 0 | 0.03 | 0.77 | 0.86 | 33.17 | 25.88 | 33.57 |
| | DPIRD | WAC4301 | DIP44_WAC4301 | 1985 | 8 | 8 | 0 | 7.56 | 10.44 | 243.13 | 32.16 | 339.66 | 32.54 |
| | DPIRD | WAC4310 | DIP47_WAC4310 | 1985 | 21 | 0 | 21 | 20.05 | 21.85 | 643.43 | 32.10 | 710.13 | 32.49 |
| | DPIRD | WAC4311 | DIP48_WAC4311 | 1985 | 0 | 0 | 0 | 0.04 | 0.66 | 1.16 | 33.05 | 21.97 | 33.52 |
| | DPIRD | WAC4317 | DIP50_WAC4317 | 1985 | 15 | 4 | 11 | 13.83 | 16.28 | 439.19 | 31.75 | 523.96 | 32.18 |
| | DPIRD | WAC4318 | DIP51_WAC4318 | 1985 | 22 | 5 | 17 | 19.55 | 22.23 | 612.16 | 31.31 | 704.50 | 31.69 |
| | DPIRD | WAC 8380 | DIP110_WAC8380 | 1989 | 32 | 13 | 19 | 31.84 | 35.35 | 1181.81 | 37.12 | 1328.78 | 37.59 |
| | DPIRD | WAC8383 | DIP55_WAC8383 | 1990 | 32 | 32 | 0 | 26.19 | 32.74 | 811.56 | 30.99 | 1026.85 | 31.36 |
| | DPIRD | WAC8387 | DIP56_WAC8387 | 1990 | 20 | 8 | 12 | 16.69 | 19.98 | 503.36 | 30.15 | 610.45 | 30.55 |

| | | | | | | | | | | | | |
|---|---|---|---|---|---|---|---|---|---|---|---|---|
| DPIRD | WAC8391 | DIP57_WAC8391 | 1990 | 21 | 1 | 20 | 19.69 | 21.42 | 610.58 | 31.01 | 673.63 | 31.45 |
| DPIRD | WAC 8397 | DIP104_WAC8397 | 1991 | 25 | 12 | 13 | 22.42 | 26.38 | 833.43 | 37.17 | 994.70 | 37.71 |
| DPIRD | WAC8399 | DIP58_WAC8399 | 1991 | 32 | 11 | 21 | 30.44 | 33.54 | 931.77 | 30.61 | 1039.87 | 31.01 |
| DPIRD | WAC8417 | DIP60_WAC8417 | 1991 | 38 | 17 | 21 | 33.19 | 38.33 | 974.17 | 29.36 | 1142.44 | 29.80 |
| DPIRD | WAC13864 | DIP62_WAC13864 | 2015 | 0 | 0 | 0 | 0.05 | 1.46 | 1.49 | 32.35 | 47.86 | 32.75 |
| DPIRD | WAC13865 | DIP63_WAC13865 | 2015 | 27 | 3 | 24 | 24.63 | 27.19 | 719.32 | 29.21 | 807.66 | 29.71 |
| DPIRD | WAC13866 | DIP64_WAC13866 | 2015 | 13 | 13 | 0 | 6.08 | 12.67 | 193.08 | 31.76 | 407.40 | 32.15 |
| DPIRD | WAC13868 | DIP66_WAC13868 | 2015 | 0 | 0 | 0 | 0.02 | 1.68 | 0.72 | 32.36 | 55.15 | 32.74 |
| DPIRD | WAC13870 | DIP68_WAC13870 | 2015 | 0 | 0 | 0 | 0.03 | 1.45 | 0.87 | 32.00 | 47.16 | 32.51 |
| DPIRD | WAC13871 | DIP69_WAC13871 | 2015 | 0 | 0 | 0 | 0.03 | 1.54 | 1.06 | 32.10 | 50.11 | 32.49 |
| DPIRD | WAC13872 | DIP70_WAC13872 | 2015 | 0 | 0 | 0 | 0.03 | 1.13 | 0.86 | 31.78 | 36.50 | 32.21 |
| DPIRD | WAC13873 | DIP71_WAC13873 | 2015 | 1 | 1 | 0 | 0.03 | 1.47 | 0.94 | 31.79 | 47.32 | 32.18 |
| DPIRD | WAC13966 | DIP76_WAC13966 | 2016 | 2 | 2 | 0 | 1.11 | 2.88 | 35.44 | 31.82 | 92.71 | 32.20 |
| DPIRD | WAC13967 | DIP77_WAC13967 | 2016 | 2 | 2 | 0 | 0.94 | 2.66 | 29.90 | 31.69 | 85.16 | 32.04 |
| DPIRD | WAC13969 | DIP79_WAC13969 | 2016 | 2 | 2 | 0 | 1.57 | 3.46 | 49.08 | 31.18 | 109.11 | 31.58 |
| DPIRD | WAC13979 | DIP82_WAC13979 | 2017 | 3 | 2 | 1 | 3.21 | 4.79 | 110.24 | 34.35 | 167.90 | 35.06 |
| DPIRD | WAC14056 | DIP83_WAC14056 | 2017 | 3 | 2 | 1 | 2.39 | 4.09 | 64.79 | 27.16 | 113.34 | 27.72 |
| DPIRD | WAC14057 | DIP84_WAC14057 | 2017 | 2 | 2 | 0 | 1.15 | 2.66 | 33.75 | 29.32 | 79.53 | 29.86 |
| DPIRD | WAC14058 | DIP85_WAC14058 | 2017 | 0 | 0 | 0 | 0.03 | 1.70 | 1.18 | 37.69 | 64.85 | 38.24 |
| DPIRD | WAC14059 | DIP86_WAC14059 | 2017 | 1 | 1 | 0 | 0.06 | 0.77 | 19.95 | 314.82 | 247.10 | 320.94 |
| DPIRD | WAC14060 | DIP87_WAC14060 | 2017 | 0 | 0 | 0 | 0.02 | 1.51 | 0.83 | 37.52 | 57.45 | 38.09 |
| DPIRD | WAC14061 | DIP88_WAC14061 | 2017 | 3 | 3 | 0 | 0.42 | 2.96 | 15.61 | 37.52 | 112.76 | 38.09 |
| DPIRD | WAC14062 | DIP89_WAC14062 | 2017 | 0 | 0 | 0 | 0.34 | 1.57 | 10.70 | 31.49 | 49.95 | 31.84 |
| DPIRD | WAC14066 | DIP90_WAC14066 | 2017 | 2 | 2 | 0 | 0.87 | 1.99 | 32.62 | 37.68 | 76.05 | 38.25 |
| DPIRD | WAC14067 | DIP91_WAC14067 | 2017 | 2 | 2 | 0 | 1.24 | 3.01 | 39.62 | 31.95 | 97.27 | 32.33 |
| DPIRD | WAC14068 | DIP92_WAC14068 | 2017 | 0 | 0 | 0 | 0.03 | 1.38 | 1.00 | 37.90 | 53.16 | 38.43 |
| DPIRD | WAC14144 | DIP100_WAC14144 | 2018 | 13 | 13 | 0 | 12.04 | 15.05 | 391.31 | 32.49 | 498.66 | 33.14 |
| DPIRD | WAC14137 | DIP93_WAC14137 | 2018 | 2 | 2 | 0 | 0.31 | 2.98 | 11.79 | 37.71 | 114.15 | 38.25 |
| DPIRD | WAC14138 | DIP94_WAC14138 | 2018 | 0 | 0 | 0 | 0.44 | 1.92 | 16.17 | 37.07 | 72.08 | 37.62 |
| DPIRD | WAC14140 | DIP96_WAC14140 | 2018 | 13 | 13 | 0 | 12.26 | 14.25 | 395.04 | 32.23 | 464.52 | 32.61 |
| DPIRD | WAC14141 | DIP97_WAC14141 | 2018 | 12 | 12 | 0 | 11.03 | 12.93 | 354.77 | 32.17 | 420.63 | 32.53 |
| OLD | WAC2216 | O1_WAC2216 | 1972 | 0 | 0 | 0 | 0.23 | 1.26 | 12.87 | 55.68 | 71.21 | 56.32 |
| OLD | WAC2285 | O2_WAC2285 | 1973 | 0 | 0 | 0 | 0.08 | 0.64 | 4.55 | 54.72 | 35.37 | 55.53 |
| OLD | WAC4303 | O5_WAC4303 | 1985 | 23 | 6 | 17 | 22.87 | 23.85 | 707.85 | 30.95 | 747.47 | 31.34 |
| OLD | WAC4321 | O6_WAC4321 | 1985 | 29 | 9 | 20 | 29.00 | 31.84 | 841.39 | 29.02 | 932.48 | 29.28 |
| OLD | WAC4808 | O8_WAC4808 | 1986 | 3 | 3 | 0 | 2.65 | 4.24 | 153.89 | 57.98 | 249.20 | 58.76 |
| OLD | WAC8384 | O10_WAC8384 | 1990 | 34 | 31 | 3 | 33.26 | 37.83 | 1668.49 | 50.17 | 1922.08 | 50.81 |
| OLD | WAC8390 | O11_WAC8390 | 1991 | 21 | 5 | 16 | 18.86 | 20.61 | 544.86 | 28.89 | 603.98 | 29.30 |
| OLD | WAC8410 | O12_WAC8410 | 1991 | 18 | 8 | 10 | 17.61 | 19.72 | 512.06 | 29.08 | 580.37 | 29.43 |
| OLD | WAC13071 | O20_WAC13071 | 2005 | 26 | 16 | 12 | 27.58 | 29.89 | 895.73 | 32.47 | 981.67 | 32.85 |
| OLD | WAC13072 | O21_WAC13072 | 2005 | 1 | 1 | 0 | 0.34 | 0.98 | 11.85 | 34.34 | 34.16 | 34.77 |
| OLD | WAC13073 | O22_WAC13073 | 2005 | 0 | 0 | 0 | 0.04 | 0.55 | 1.71 | 48.51 | 26.86 | 49.07 |
| OLD | WAC13077 | O26_WAC13077 | 2005 | 0 | 0 | 0 | 0.04 | 0.77 | 1.06 | 28.43 | 22.51 | 29.22 |
| OLD | Meck1 | O27_Meck1 | 2009 | 1 | 1 | 0 | 1.08 | 2.21 | 33.25 | 30.74 | 68.65 | 31.10 |
| OLD | Meck3 | O28_Meck3 | 2009 | 1 | 1 | 0 | 1.10 | 1.47 | 34.39 | 31.24 | 46.42 | 31.63 |
| OLD | Meck5 | O29_Meck5 | 2009 | 1 | 1 | 0 | 1.10 | 1.90 | 37.95 | 34.49 | 66.55 | 35.00 |
| OLD | Meck6 | O30_Meck6 | 2009 | 22 | 0 | 22 | 21.05 | 22.03 | 699.48 | 33.23 | 739.78 | 33.58 |
| OLD | Meck8 | O31_Meck8 | 2009 | 3 | 3 | 0 | 1.02 | 2.76 | 36.24 | 35.44 | 98.94 | 35.82 |
| OLD | S1FT3A | O32_S1FT3A | 2009 | 18 | 2 | 16 | 16.34 | 18.21 | 622.14 | 38.06 | 702.98 | 38.61 |

| | | | | | | | | | | | | |
|---|---|---|---|---|---|---|---|---|---|---|---|---|
| OLD | S4FT3A | O33_S4FT3A | 2009 | 0 | 0 | 0 | 0.02 | 0.36 | 0.62 | 33.78 | 12.26 | 34.15 |
| OLD | S1FT3B | O34_S1FT3B | 2009 | 18 | 2 | 16 | 16.92 | 17.90 | 646.13 | 38.19 | 692.45 | 38.69 |
| OLD | Gerald1 | O35_Gerald1 | 2011 | 0 | 0 | 0 | 0.09 | 1.04 | 2.81 | 31.91 | 33.63 | 32.26 |
| OLD | Gerald4 | O36_Gerald4 | 2011 | 2 | 2 | 0 | 1.40 | 3.27 | 61.50 | 43.91 | 144.97 | 44.36 |
| OLD | Mur_51 | O37_Mur_51 | 2011 | 13 | 13 | 0 | 12.82 | 16.27 | 426.69 | 33.28 | 547.24 | 33.64 |
| OLD | WAC13402 | O39_WAC13402 | 2011 | 0 | 0 | 0 | 0.05 | 0.59 | 1.54 | 33.00 | 19.56 | 33.38 |
| OLD | WAC13443 | O43_WAC13443 | 2011 | 0 | 0 | 0 | 0.02 | 0.72 | 0.76 | 37.57 | 27.66 | 38.17 |
| OLD | WAC13615 | O56_WAC13615 | 2012 | 23 | 3 | 20 | 21.71 | 23.53 | 620.49 | 28.58 | 679.55 | 28.89 |
| OLD | WAC13616 | O57_WAC13616 | 2012 | 23 | 3 | 20 | 21.23 | 23.12 | 978.20 | 46.07 | 1079.65 | 46.70 |
| OLD | WAC13617 | O58_WAC13617 | 2012 | 23 | 3 | 20 | 21.73 | 23.69 | 1100.56 | 50.65 | **1219.02** | 51.46 |
| OLD | WAC13690 | O64_WAC13690 | 2012 | 0 | 0 | 0 | 0.36 | 1.25 | 11.64 | 32.38 | 41.03 | 32.79 |
| OLD | WAC13691 | O65_WAC13691 | 2012 | 0 | 0 | 0 | 0.17 | 1.39 | 6.39 | 38.59 | 54.32 | 39.17 |
| OLD | WAC13666 | O62_WAC13666 | 2013 | 0 | 0 | 0 | 0.18 | 1.50 | 5.40 | 30.81 | 46.83 | 31.13 |
| OLD | WAC13667 | O63_WAC13667 | 2013 | 0 | 0 | 0 | 0.09 | 0.87 | 5.05 | 57.91 | 50.98 | 58.53 |
| OLD | 15FG107 | O107_15FG107 | 2014 | 0 | 0 | 0 | 0.11 | 0.95 | 6.66 | 58.24 | 56.00 | 58.96 |
| OLD | 15FG109 | O108_15FG109 | 2014 | 0 | 0 | 0 | 0.03 | 0.32 | 1.11 | 34.33 | 11.19 | 34.76 |
| OLD | 15FG120 | O118_15FG120 | 2014 | 1 | 1 | 0 | 1.09 | 1.60 | 68.72 | 63.03 | 102.04 | 63.78 |
| OLD | 206FG226 | O66_206FG226 | 2014 | 1 | 1 | 0 | 1.17 | 1.66 | 41.15 | 35.32 | 59.49 | 35.81 |
| OLD | 205FG215_1 | O67_205FG215_1 | 2014 | 16 | 4 | 12 | 15.89 | 16.96 | 548.70 | 34.54 | 593.12 | 34.98 |
| OLD | 201FG209 | O68_201FG209 | 2014 | 26 | 2 | 24 | 25.59 | 26.84 | 896.71 | 35.04 | 953.87 | 35.54 |
| OLD | 201FG211 | O69_201FG211 | 2014 | 19 | 0 | 19 | 17.92 | 19.38 | 640.98 | 35.78 | 702.01 | 36.23 |
| OLD | 202FG212 | O70_202FG212 | 2014 | 18 | 3 | 15 | 17.51 | 18.66 | 618.54 | 35.32 | 667.40 | 35.77 |
| OLD | 204FG221 | O71_204FG221 | 2014 | 2 | 2 | 0 | 2.01 | 2.71 | 59.25 | 29.46 | 80.79 | 29.86 |
| OLD | 204FG223 | O72_204FG223 | 2014 | 0 | 0 | 0 | 0.07 | 0.81 | 2.36 | 32.32 | 26.66 | 32.86 |
| OLD | 205FG225 | O73_205FG225 | 2014 | 12 | 2 | 10 | 11.72 | 12.46 | 401.94 | 34.30 | 432.24 | 34.69 |
| OLD | 206FG227 | O74_206FG227 | 2014 | 3 | 3 | 0 | 1.47 | 3.53 | 60.56 | 41.18 | 147.17 | 41.64 |
| OLD | 205FG216 | O75_205FG216 | 2014 | 2 | 2 | 0 | 1.07 | 1.99 | 60.03 | 56.35 | 114.27 | 57.28 |
| OLD | 201FG219 | O76_201FG219 | 2014 | 1 | 1 | 0 | 0.08 | 0.90 | 3.56 | 44.02 | 39.99 | 44.60 |
| OLD | 201FG218 | O77_201FG218 | 2014 | 19 | 2 | 17 | 18.52 | 19.25 | 798.37 | 43.10 | 838.95 | 43.57 |
| OLD | 204FG214 | O78_204FG214 | 2014 | 2 | 2 | 0 | 2.10 | 2.77 | 92.35 | 43.98 | 123.12 | 44.44 |
| OLD | 903FG214 | O79_903FG214 | 2014 | 19 | 8 | 11 | 17.68 | 19.57 | 595.34 | 33.68 | 666.42 | 34.05 |
| OLD | 201FG49 | O80_201FG49 | 2014 | 1 | 1 | 0 | 0.08 | 0.63 | 3.11 | 40.34 | 25.67 | 40.77 |
| OLD | 203FG58 | O81_203FG58 | 2014 | 2 | 2 | 0 | 0.41 | 2.70 | 13.04 | 31.89 | 86.88 | 32.23 |
| OLD | 205FG63 | O82_205FG63 | 2014 | 1 | 1 | 0 | 0.14 | 0.71 | 4.34 | 30.64 | 22.11 | 30.96 |
| OLD | 206FG66 | O83_206FG66 | 2014 | 1 | 1 | 0 | 1.09 | 1.84 | 42.33 | 38.93 | 72.48 | 39.36 |
| OLD | 206FG67 | O84_206FG67 | 2014 | 0 | 0 | 0 | 0.05 | 0.64 | 1.77 | 36.47 | 23.65 | 36.88 |
| OLD | 206FG68 | O85_206FG68 | 2014 | 1 | 1 | 0 | 0.23 | 1.82 | 8.32 | 35.57 | 65.38 | 35.96 |
| OLD | 205FG142 | O86_205FG142 | 2014 | 0 | 0 | 0 | 0.02 | 0.41 | 0.55 | 30.21 | 12.59 | 30.52 |
| OLD | 53FG143_1 | O87_53FG143_1 | 2014 | 0 | 0 | 0 | 0.06 | 1.06 | 1.91 | 30.21 | 32.29 | 30.55 |
| OLD | 201FG208 | O88_201FG208 | 2014 | 18 | 8 | 10 | 15.69 | 17.82 | 866.12 | 55.19 | 996.32 | 55.90 |
| OLD | 203FG213 | O89_203FG213 | 2014 | 2 | 2 | 0 | 1.43 | 2.45 | 42.82 | 29.94 | 74.30 | 30.29 |
| OLD | 204FG222 | O91_204FG222 | 2014 | 14 | 3 | 11 | 13.62 | 15.25 | 442.45 | 32.48 | 501.54 | 32.89 |
| OLD | 205FG410 | O92_205FG410 | 2014 | 0 | 0 | 0 | 0.09 | 1.37 | 5.34 | 62.30 | 86.27 | 63.00 |
| OLD | 202FG414 | O93_202FG414 | 2014 | 2 | 2 | 0 | 1.33 | 2.63 | 77.98 | 58.57 | 156.48 | 59.39 |
| OLD | WAC739 | O94_WAC739 | 2014 | 2 | 2 | 0 | 1.24 | 2.14 | 42.11 | 33.96 | 73.31 | 34.33 |
| OLD | WAC740 | O95_WAC740 | 2014 | 1 | 1 | 0 | 1.05 | 1.60 | 45.09 | 42.79 | 69.40 | 43.27 |
| OLD | WAC741 | O96_WAC741 | 2014 | 2 | 2 | 0 | 1.19 | 2.04 | 60.90 | 51.24 | 105.91 | 52.00 |
| OLD | 15FG102 | O100_15FG102 | 2015 | 27 | 17 | 10 | 26.26 | 28.01 | 802.09 | 30.54 | 866.61 | 30.94 |
| OLD | 15FG103 | O101_15FG103 | 2015 | 29 | 18 | 11 | 28.85 | 30.85 | 1712.83 | 59.36 | 1853.57 | 60.08 |

| | | | | | | | | | | | | |
|---|---|---|---|---|---|---|---|---|---|---|---|---|
| OLD | 15FG104 | O102_15FG104 | 2015 | 21 | 13 | 8 | 21.19 | 21.89 | 820.56 | 38.73 | 857.72 | 39.18 |
| OLD | 15FG105 | O103_15FG105 | 2015 | 18 | 2 | 16 | 16.46 | 17.91 | 725.98 | 44.11 | 800.79 | 44.70 |
| OLD | FG106 | O104_FG106 | 2015 | 0 | 0 | 0 | 0.03 | 0.69 | 1.68 | 53.77 | 37.26 | 54.40 |
| OLD | FG107 | O105_FG107 | 2015 | 0 | 0 | 0 | 0.06 | 0.58 | 1.98 | 34.29 | 20.03 | 34.70 |
| OLD | FG108 | O106_FG108 | 2015 | 0 | 0 | 0 | 0.09 | 0.95 | 5.32 | 58.09 | 55.87 | 58.80 |
| OLD | 15FG110 | O109_15FG110 | 2015 | 1 | 1 | 0 | 0.21 | 1.62 | 10.58 | 49.51 | 78.64 | 48.44 |
| OLD | 15FG112 | O110_15FG112 | 2015 | 1 | 1 | 0 | 0.04 | 0.44 | 2.71 | 61.34 | 27.20 | 62.20 |
| OLD | 15FG113 | O111_15FG113 | 2015 | 1 | 1 | 0 | 0.06 | 0.87 | 3.50 | 55.23 | 48.58 | 55.97 |
| OLD | 15FG114 | O112_15FG114 | 2015 | 2 | 2 | 0 | 0.19 | 1.48 | 13.52 | 71.12 | 106.93 | 72.04 |
| OLD | 15FG115 | O113_15FG115 | 2015 | 2 | 2 | 0 | 0.50 | 2.81 | 28.66 | 57.77 | 164.21 | 58.51 |
| OLD | 15FG116 | O114_15FG116 | 2015 | 1 | 1 | 0 | 0.16 | 1.12 | 8.64 | 54.87 | 62.08 | 55.55 |
| OLD | 15FG117 | O115_15FG117 | 2015 | 1 | 1 | 0 | 0.19 | 1.25 | 14.12 | 75.62 | 95.59 | 76.56 |
| OLD | 15FG118 | O116_15FG118 | 2015 | 1 | 1 | 0 | 0.18 | 1.15 | 12.10 | 66.33 | 76.90 | 67.11 |
| OLD | 15FG119 | O117_15FG119 | 2015 | 1 | 1 | 0 | 0.17 | 1.17 | 9.35 | 55.14 | 65.51 | 55.79 |
| OLD | 15FG04 | O119_15FG04 | 2015 | 22 | 5 | 17 | 20.18 | 21.80 | 1850.02 | 91.68 | 2030.06 | 93.13 |
| OLD | 15FG28 | O120_15FG28 | 2015 | 1 | 1 | 0 | 0.39 | 1.56 | 16.92 | 43.04 | 67.79 | 43.57 |
| OLD | 15FG33 | O122_15FG33 | 2015 | 1 | 1 | 0 | 1.29 | 2.45 | 74.26 | 57.66 | 142.98 | 58.27 |
| OLD | 15FG37 | O123_15FG37 | 2015 | 3 | 3 | 0 | 1.00 | 4.71 | 48.79 | 48.97 | 233.25 | 49.56 |
| OLD | 15FG38 | O124_15FG38 | 2015 | 20 | 9 | 11 | 18.74 | 21.29 | 652.16 | 34.79 | 762.80 | 35.83 |
| OLD | 15FG47 | O127_15FG47 | 2015 | 0 | 0 | 0 | 0.07 | 0.78 | 2.22 | 33.22 | 26.13 | 33.59 |
| OLD | 15FG49 | O128_15FG49 | 2015 | 0 | 0 | 0 | 0.04 | 0.57 | 2.38 | 54.11 | 31.03 | 54.82 |
| OLD | 15FG226 | O129_15FG226 | 2015 | 0 | 0 | 0 | 0.06 | 1.24 | 2.24 | 34.48 | 43.34 | 34.86 |
| OLD | 15FG229 | O130_15FG229 | 2015 | 3 | 3 | 0 | 3.13 | 3.88 | 124.85 | 39.91 | 157.01 | 40.43 |
| OLD | 15FG237 | O131_15FG237 | 2015 | 3 | 3 | 0 | 3.29 | 4.00 | 189.14 | 57.43 | 232.58 | 58.10 |
| OLD | Northam_Mace2 | O132_Northam_Mace2 | 2015 | 1 | 1 | 0 | 1.13 | 1.82 | 42.14 | 37.31 | 68.53 | 37.72 |
| OLD | Northam_Magenta | O133_Northam_Mager | 2015 | 45 | 4 | 41 | 43.71 | 45.12 | 1359.24 | 31.09 | 1418.85 | 31.45 |
| OLD | Northam_Emu1 | O134_Northam_Emu1 | 2015 | 0 | 0 | 0 | 0.03 | 0.35 | 1.16 | 34.70 | 12.46 | 35.16 |
| OLD | Northam_Emu2 | O135_Northam_Emu2 | 2015 | 0 | 0 | 0 | 0.08 | 0.98 | 2.59 | 32.32 | 31.95 | 32.68 |
| OLD | Northam_Mace1 | O136_Northam_Mace1 | 2015 | 2 | 2 | 0 | 1.20 | 2.39 | 43.05 | 35.82 | 86.35 | 36.18 |
| OLD | FG_W003_5 | O140_FG_W003_5 | 2015 | 3 | 3 | 0 | 3.28 | 3.97 | 184.73 | 56.31 | 228.16 | 57.47 |
| OLD | Nor_RAC2182_1 | O141_Nor_RAC2182_ | 2015 | 15 | 1 | 14 | 14.23 | 15.30 | 418.80 | 29.43 | 455.42 | 29.77 |
| OLD | Northam_WGT | O142_Northam_WGT | 2015 | 2 | 2 | 0 | 1.15 | 1.91 | 42.24 | 36.62 | 70.80 | 37.06 |
| OLD | Nor_RAC2182_2 | O143_Nor_RAC2182_ | 2015 | 0 | 0 | 0 | 0.21 | 2.05 | 10.35 | 49.03 | 101.82 | 49.70 |
| OLD | 15FG99 | O97_15FG99 | 2015 | 1 | 1 | 0 | 0.06 | 0.97 | 3.70 | 60.07 | 58.84 | 60.76 |
| OLD | 15FG100 | O98_15FG100 | 2015 | 32 | 2 | 30 | 31.96 | 33.31 | 1820.12 | 56.95 | 1925.86 | 57.81 |
| OLD | 15FG101 | O99_15FG101 | 2015 | 2 | 2 | 0 | 1.23 | 2.18 | 68.05 | 55.12 | 121.51 | 55.87 |
| OLD | 16FG06 | O143_16FG06 | 2016 | 1 | 1 | 0 | 1.01 | 1.43 | 144.77 | 143.78 | 209.64 | 146.66 |
| OLD | 16FG158 | O144_16FG158 | 2016 | 1 | 1 | 0 | 1.41 | 2.33 | 222.75 | 158.28 | 375.74 | 161.04 |
| OLD | 16FG159 | O145_16FG159 | 2016 | 1 | 1 | 0 | 0.78 | 2.19 | 129.18 | 166.45 | 369.28 | 168.82 |
| OLD | WAC13955 | O157_WAC13955 | 2016 | 2 | 2 | 0 | 1.24 | 2.11 | 182.86 | 147.60 | 314.85 | 149.38 |
| OLD | Mur_S3 | O160_Mur_S3 | 2011 | 1 | 1 | 0 | 1.01 | 1.76 | 150.54 | 148.69 | 265.66 | 151.17 |
| OLD | 14FG141 | O161_14FG141 | 2014 | 2 | 2 | 0 | 0.65 | 1.75 | 120.55 | 185.76 | 329.18 | 187.60 |
| Current | PN260 | C260_PN260 | 2019 | 0 | 0 | 0 | 0.06 | 0.60 | 5.41 | 93.05 | 56.75 | 93.87 |
| Current | PN261 | C261_PN261 | 2019 | 1 | 1 | 0 | 0.59 | 2.06 | 43.98 | 74.72 | 155.06 | 75.38 |
| Current | PN262 | C262_PN262 | 2019 | 2 | 1 | 1 | 1.49 | 2.11 | 126.55 | 84.67 | 180.59 | 85.69 |
| Current | PN263 | C263_PN263 | 2019 | 1 | 1 | 0 | 0.45 | 1.16 | 20.94 | 46.56 | 54.42 | 46.98 |
| Current | PN264 | C264_PN264 | 2019 | 1 | 1 | 0 | 0.03 | 0.49 | 3.26 | 110.80 | 54.74 | 111.79 |
| Current | PN265 | C265_PN265 | 2019 | 2 | 2 | 0 | 1.49 | 2.47 | 113.88 | 76.66 | 191.40 | 77.36 |
| Current | PN266 | C266_PN266 | 2019 | 1 | 1 | 0 | 0.17 | 0.96 | 16.30 | 93.73 | 93.81 | 97.33 |

| | | | | | | | | | | | | |
|---|---|---|---|---|---|---|---|---|---|---|---|---|
| Current | PN268 | C268_PN268 | 2019 | 2 | 2 | 0 | 1.26 | 2.09 | 134.39 | 106.88 | 225.40 | 108.10 |
| Current | PN269 | C269_PN269 | 2019 | 3 | 3 | 0 | 1.46 | 2.87 | 148.37 | 101.51 | 294.29 | 102.42 |
| Current | PN270 | C270_PN270 | 2019 | 0 | 0 | 0 | 0.05 | 0.57 | 4.60 | 90.87 | 52.01 | 91.69 |
| Current | PN271 | C271_PN271 | 2019 | 0 | 0 | 0 | 0.06 | 0.66 | 5.46 | 88.65 | 59.00 | 89.40 |
| Current | PN274 | C274_PN274 | 2019 | 0 | 0 | 0 | 0.07 | 0.62 | 6.89 | 93.56 | 58.68 | 94.53 |
| Current | PN275 | C275_PN275 | 2019 | 29 | 16 | 13 | 27.68 | 29.43 | 3371.32 | 121.78 | 3625.91 | 123.21 |
| Current | PN276 | C276_PN276 | 2019 | 1 | 1 | 0 | 0.08 | 0.87 | 9.15 | 107.98 | 94.94 | 109.08 |
| Current | PN277 | C277_PN277 | 2019 | 0 | 0 | 0 | 0.07 | 0.53 | 7.11 | 109.10 | 59.05 | 110.47 |
| Current | PN279 | C279_PN279 | 2019 | 0 | 0 | 0 | 0.09 | 0.75 | 12.00 | 136.84 | 104.10 | 138.07 |
| Current | PN284 | C284_PN284 | 2019 | 2 | 2 | 0 | 1.10 | 1.78 | 99.83 | 90.93 | 163.10 | 91.87 |
| Current | PN285 | C285_PN285 | 2019 | 0 | 0 | 0 | 0.10 | 0.68 | 7.73 | 79.36 | 54.50 | 80.12 |
| Current | PN286 | C286_PN286 | 2019 | 2 | 2 | 0 | 1.28 | 1.96 | 82.13 | 64.16 | 127.01 | 64.72 |
| Current | PN287 | C287_PN287 | 2019 | 0 | 0 | 0 | 0.21 | 1.23 | 14.78 | 71.96 | 89.63 | 72.60 |
| Current | PN288 | C288_PN288 | 2019 | 1 | 1 | 0 | 0.10 | 0.78 | 5.81 | 57.89 | 45.34 | 58.41 |
| Current | PN289 | C289_PN289 | 2019 | 1 | 1 | 0 | 0.07 | 0.74 | 5.15 | 74.74 | 55.71 | 75.42 |
| Current | PN290 | C290_PN290 | 2019 | 3 | 3 | 0 | 1.51 | 2.57 | 90.61 | 60.11 | 156.11 | 60.67 |
| Current | PN291 | C291_PN291 | 2019 | 1 | 1 | 0 | 0.23 | 1.25 | 15.46 | 67.36 | 87.66 | 69.93 |
| Current | PN292 | C292_PN292 | 2019 | 0 | 0 | 0 | 0.08 | 0.70 | 4.24 | 56.31 | 39.95 | 56.83 |
| Current | PN295 | C295_PN295 | 2019 | 30 | 3 | 27 | 30.55 | 31.73 | 2146.55 | 70.26 | 2254.74 | 71.06 |
| Current | PN297 | C297_PN297 | 2019 | 24 | 7 | 17 | 23.43 | 24.33 | 1426.87 | 60.90 | 1496.67 | 61.52 |
| Current | PN299 | C299_PN299 | 2019 | 1 | 1 | 0 | 0.44 | 1.62 | 30.14 | 68.83 | 112.22 | 69.45 |
| Current | PN300 | C300_PN300 | 2019 | 1 | 1 | 0 | 0.26 | 1.47 | 12.95 | 49.38 | 73.23 | 49.84 |
| Current | PN301 | C301_PN301 | 2019 | 1 | 1 | 0 | 0.40 | 1.70 | 24.84 | 61.34 | 105.39 | 61.91 |
| Current | PN302 | C302_PN302 | 2019 | 2 | 2 | 0 | 1.14 | 1.93 | 69.71 | 60.90 | 118.81 | 61.47 |
| Current | PN303 | C303_PN303 | 2019 | 1 | 1 | 0 | 0.08 | 1.00 | 6.69 | 82.58 | 83.28 | 83.36 |
| Current | PN304 | C304_PN304 | 2019 | 2 | 2 | 0 | 1.14 | 1.77 | 77.08 | 67.45 | 120.74 | 68.07 |
| Current | PN305 | C305_PN305 | 2019 | 0 | 0 | 0 | 0.09 | 0.70 | 6.88 | 78.37 | 55.63 | 79.12 |
| Current | PN308 | C308_PN308 | 2019 | 2 | 2 | 0 | 0.55 | 1.84 | 34.43 | 62.93 | 116.80 | 63.51 |
| Current | PN309 | C309_PN309 | 2019 | 28 | 8 | 20 | 26.93 | 27.83 | 1496.78 | 55.57 | 1561.77 | 56.11 |
| Current | PN312 | C312_PN312 | 2019 | 23 | 23 | 0 | 22.20 | 23.35 | 1473.70 | 66.38 | 1564.78 | 67.00 |
| Current | PN313 | C313_PN313 | 2019 | 25 | 9 | 16 | 23.96 | 25.22 | 1811.32 | 75.61 | 1925.23 | 76.34 |
| Current | PN314 | C314_PN314 | 2019 | 2 | 1 | 1 | 1.03 | 2.25 | 63.10 | 61.31 | 138.99 | 61.89 |
| Current | PN317 | C317_PN317 | 2019 | 1 | 1 | 0 | 0.62 | 1.45 | 41.10 | 66.42 | 97.19 | 67.18 |
| Current | PN318 | C318_PN318 | 2019 | 0 | 0 | 0 | 0.08 | 0.78 | 6.54 | 79.60 | 62.54 | 80.45 |
| Current | PN319 | C319_PN319 | 2019 | 1 | 1 | 0 | 0.28 | 1.06 | 23.67 | 83.62 | 89.95 | 84.54 |
| Current | PN320 | C320_PN320 | 2019 | 2 | 2 | 0 | 1.18 | 1.99 | 89.45 | 75.85 | 152.49 | 76.54 |
| Current | PN323 | C323_PN323 | 2019 | 2 | 2 | 0 | 0.11 | 1.00 | 7.26 | 65.64 | 66.21 | 66.25 |
| Current | PN324 | C324_PN324 | 2019 | 2 | 2 | 0 | 0.27 | 1.12 | 16.24 | 61.27 | 69.37 | 61.88 |
| Current | PN325 | C325_PN325 | 2019 | 1 | 1 | 0 | 1.14 | 1.56 | 77.19 | 67.46 | 106.27 | 68.05 |
| Current | PN326 | C326_PN326 | 2019 | 5 | 4 | 1 | 4.21 | 4.94 | 323.71 | 76.90 | 383.68 | 77.75 |
| Current | PN328 | C328_PN328 | 2019 | 26 | 26 | 0 | 25.13 | 26.18 | 1364.02 | 54.28 | 1434.63 | 54.80 |
| Current | PN329 | C329_PN329 | 2019 | 28 | 28 | 0 | 27.22 | 28.46 | 1903.26 | 69.91 | 2008.37 | 70.57 |
| Current | PN330 | C330_PN330 | 2019 | 0 | 0 | 0 | 0.10 | 0.62 | 6.66 | 66.47 | 41.48 | 67.14 |
| Current | PN331 | C331_PN331 | 2019 | 1 | 0 | 1 | 0.43 | 1.54 | 24.96 | 58.58 | 91.00 | 59.19 |
| Current | PN332 | C332_PN332 | 2019 | 18 | 5 | 13 | 15.97 | 18.10 | 961.96 | 60.22 | 1100.96 | 60.83 |
| Current | PN336 | C336_PN336 | 2019 | 1 | 1 | 0 | 1.23 | 1.83 | 73.41 | 59.58 | 110.29 | 60.15 |
| Current | PN342 | C342_PN342 | 2019 | 3 | 3 | 0 | 0.41 | 1.62 | 32.59 | 79.78 | 130.79 | 80.97 |
| Current | PN347 | C347_PN347 | 2019 | 0 | 0 | 0 | 0.16 | 1.33 | 10.39 | 64.78 | 86.82 | 65.42 |
| Current | PN348 | C348_PN348 | 2019 | 3 | 3 | 0 | 1.40 | 2.55 | 109.61 | 78.57 | 202.49 | 79.28 |

| | | | | | | | | | | | | | |
|---|---|---|---|---|---|---|---|---|---|---|---|---|---|
| | Current | PN349 | C349_PN349 | 2019 | 2 | 2 | 0 | 1.22 | 1.94 | 65.53 | 53.68 | 104.98 | 54.19 |
| | Current | PN351 | C351_PN351 | 2020 | 0 | 0 | 0 | 0.09 | 0.71 | 6.72 | 77.10 | 55.39 | 77.87 |
| | Current | PN352 | C352_PN352 | 2020 | 3 | 3 | 0 | 1.46 | 2.55 | 94.59 | 65.00 | 167.69 | 65.72 |
| | Current | PN353 | C353_PN353 | 2021 | 1 | 1 | 0 | 1.14 | 1.50 | 75.88 | 66.59 | 100.56 | 67.22 |
| | Current | PN354 | C354_PN354 | 2021 | 0 | 0 | 0 | 0.20 | 1.27 | 13.75 | 68.76 | 87.94 | 69.40 |
| | Current | PN357 | C357_PN357 | 2021 | 0 | 0 | 0 | 0.04 | 0.58 | 2.58 | 64.33 | 37.46 | 64.92 |
| | Current | PN358 | C358_PN358 | 2021 | 37 | 7 | 30 | 35.27 | 36.62 | 3350.52 | 95.00 | 3518.25 | 96.08 |
| | Current | PN360 | C360_PN360 | 2021 | 0 | 0 | 0 | 0.04 | 0.65 | 2.64 | 72.00 | 47.43 | 72.66 |
| | Current | PN361 | C361_PN361 | 2021 | 1 | 1 | 0 | 0.26 | 1.37 | 23.76 | 90.22 | 124.84 | 91.36 |
| | Current | PN362 | C362_PN362 | 2021 | 1 | 1 | 0 | 1.24 | 1.83 | 82.18 | 66.51 | 122.67 | 67.12 |
| | Current | PN363 | C363_PN363 | 2021 | 1 | 1 | 0 | 0.28 | 1.38 | 17.71 | 63.00 | 87.62 | 63.71 |
| | Current | PN364 | C364_PN364 | 2021 | 39 | 7 | 32 | 39.16 | 40.29 | 2222.86 | 56.76 | 2309.30 | 57.31 |
| | Current | PN366 | C366_PN366 | 2021 | 1 | 1 | 0 | 0.21 | 1.25 | 16.71 | 77.77 | 98.39 | 78.49 |
| | Current | PN367 | C367_PN367 | 2021 | 1 | 1 | 0 | 0.06 | 0.91 | 4.28 | 71.54 | 66.04 | 72.22 |
| | Current | PN369 | C369_PN369 | 2021 | 1 | 1 | 0 | 0.51 | 1.75 | 41.29 | 80.23 | 142.28 | 81.17 |
| | Current | PN371 | C371_PN371 | 2021 | 1 | 1 | 0 | 0.72 | 1.67 | 39.37 | 54.31 | 91.64 | 54.92 |
| | Current | PN372 | C372_PN372 | 2021 | 1 | 1 | 0 | 0.25 | 1.56 | 15.48 | 62.16 | 97.83 | 62.74 |
| | Current | PN375 | C375_PN375 | 2021 | 4 | 3 | 1 | 2.99 | 4.05 | 218.29 | 73.07 | 298.31 | 73.74 |
| | Current | PN376 | C376_PN376 | 2021 | 2 | 2 | 0 | 0.58 | 2.29 | 47.56 | 81.60 | 188.34 | 82.39 |
| | Current | PN377 | C377_PN377 | 2021 | 2 | 2 | 0 | 0.56 | 2.28 | 34.07 | 60.54 | 139.59 | 61.15 |
| | Current | PN380 | C380_PN380 | 2021 | 1 | 1 | 0 | 1.31 | 1.90 | 80.48 | 61.41 | 117.52 | 61.97 |
| Group 2 | DPIRD | WAC1141 | DIP1_WAC1141 | 1968 | 1 | 1 | 0 | 0.03 | 1.49 | 0.90 | 31.73 | 47.86 | 32.10 |
| | DPIRD | WAC1494 | DIP11_WAC1494 | 1969 | 0 | 0 | 0 | 0.06 | 0.74 | 15.39 | 245.00 | 184.18 | 249.45 |
| | DPIRD | WAC1495 | DIP12_WAC1495 | 1969 | 0 | 0 | 0 | 0.03 | 1.40 | 0.89 | 31.79 | 45.12 | 32.17 |
| | DPIRD | WAC1548 | DIP21_WAC1548 | 1969 | 0 | 0 | 1 | 0.04 | 1.52 | 1.38 | 32.40 | 49.79 | 32.77 |
| | DPIRD | WAC1564 | DIP27_WAC1564 | 1969 | 1 | 1 | 0 | 0.04 | 1.43 | 1.24 | 31.85 | 46.09 | 32.22 |
| | DPIRD | WAC2284 | DIP30_WAC2284 | 1973 | 0 | 0 | 0 | 0.04 | 0.57 | 10.94 | 243.27 | 140.43 | 248.06 |
| | DPIRD | WAC 2804 | DIP105_WAC2804 | 1980 | 0 | 0 | 0 | 0.04 | 1.34 | 1.52 | 37.98 | 51.50 | 38.51 |
| | DPIRD | WAC 2816 | DIP102_WAC2816 | 1980 | 0 | 0 | 0 | 0.04 | 1.46 | 1.42 | 31.83 | 46.89 | 32.21 |
| | DPIRD | WAC2285 | DIP31_WAC2285 | 1980 | 0 | 0 | 0 | 0.04 | 1.26 | 1.28 | 32.39 | 41.43 | 32.77 |
| | DPIRD | WAC2805 | DIP38_WAC2805 | 1980 | 1 | 1 | 0 | 0.05 | 1.41 | 1.45 | 32.16 | 45.98 | 32.54 |
| | DPIRD | WAC2817 | DIP42_WAC2817 | 1980 | 1 | 1 | 0 | 0.04 | 1.61 | 1.19 | 31.81 | 51.93 | 32.21 |
| | DPIRD | WAC4304 | DIP45_WAC4304 | 1985 | 0 | 0 | 0 | 0.03 | 0.96 | 1.06 | 31.91 | 31.14 | 32.30 |
| | DPIRD | WAC4316 | DIP49_WAC4316 | 1985 | 0 | 0 | 0 | 0.03 | 1.42 | 1.03 | 31.62 | 45.28 | 31.99 |
| | DPIRD | WAC13957 | DIP72_WAC13957 | 2016 | 0 | 0 | 0 | 0.04 | 1.57 | 1.42 | 32.04 | 50.91 | 32.45 |
| | OLD | WAC4319 | O7_WAC4319 | 1985 | 1 | 1 | 0 | 0.07 | 1.10 | 2.76 | 41.75 | 46.65 | 42.25 |
| | OLD | WAC4648 | O9_WAC4648 | 1986 | 1 | 1 | 0 | 0.05 | 0.90 | 2.75 | 59.33 | 54.19 | 60.04 |
| | OLD | WAC13068 | O17_WAC13068 | 2005 | 1 | 1 | 0 | 0.12 | 0.97 | 3.14 | 26.93 | 26.45 | 27.28 |
| | OLD | WAC13069 | O18_WAC13069 | 2005 | 1 | 1 | 0 | 0.07 | 0.80 | 3.13 | 46.79 | 38.06 | 47.33 |
| | OLD | WAC13523 | O46_WAC13523 | 2011 | 1 | 1 | 0 | 0.06 | 0.92 | 2.08 | 37.69 | 35.26 | 38.12 |
| | OLD | WAC13524 | O47_WAC13524 | 2011 | 1 | 1 | 0 | 0.06 | 0.76 | 2.29 | 39.83 | 32.42 | 42.70 |
| | OLD | WAC13527 | O50_WAC13527 | 2011 | 1 | 1 | 0 | 1.35 | 2.15 | 60.78 | 44.97 | 98.01 | 45.66 |
| | OLD | WAC13529 | O52_WAC13529 | 2011 | 1 | 1 | 0 | 0.06 | 0.93 | 3.92 | 60.98 | 57.22 | 61.72 |
| | OLD | WAC13530 | O53_WAC13530 | 2011 | 1 | 1 | 0 | 0.05 | 0.73 | 3.25 | 60.64 | 45.09 | 61.53 |
| | OLD | WAC13531 | O54_WAC13531 | 2011 | 1 | 1 | 0 | 0.07 | 0.87 | 3.38 | 49.27 | 43.63 | 49.89 |
| Group 3 | DPIRD | WAC1496 | DIP13_WAC1496 | 1969 | 4 | 4 | 0 | 3.82 | 5.01 | 153.30 | 40.18 | 203.64 | 40.65 |
| | DPIRD | WAC1547 | DIP20_WAC1547 | 1969 | 1 | 1 | 0 | 0.88 | 1.54 | 34.54 | 39.34 | 61.20 | 39.82 |
| | DPIRD | WAC2806 | DIP39_WAC2806 | 1980 | 18 | 0 | 18 | 17.35 | 18.13 | 587.00 | 33.83 | 621.06 | 34.25 |
| | DPIRD | WAC2808 | DIP40_WAC2808 | 1980 | 0 | 0 | 0 | 0.07 | 1.72 | 2.11 | 31.55 | 54.82 | 31.93 |

| | | | | | | | | | | | | | |
|---|---|---|---|---|---|---|---|---|---|---|---|---|---|
| | DPIRD | WAC4308 | DIP46_WAC4308 | 1985 | 2 | 1 | 1 | 0.27 | 2.22 | 8.44 | 31.28 | 70.36 | 31.71 |
| | OLD | WAC2810 | O3_WAC2810 | 1980 | 1 | 1 | 0 | 0.13 | 1.28 | 4.09 | 30.49 | 39.68 | 30.88 |
| | OLD | WAC2813 | O4_WAC2813 | 1980 | 1 | 1 | 0 | 0.16 | 1.26 | 7.06 | 44.49 | 56.81 | 45.08 |
| | OLD | WAC8635 | O13_WAC8635 | 1994 | 1 | 1 | 0 | 0.15 | 1.24 | 4.90 | 33.28 | 41.94 | 33.71 |
| | OLD | WAC9178 | O14_WAC9178 | 1996 | 1 | 1 | 0 | 0.17 | 1.24 | 5.66 | 33.49 | 41.94 | 33.91 |
| | OLD | WAC13418 | O15_WAC13418 | 1996 | 1 | 1 | 0 | 0.12 | 0.80 | 5.90 | 50.18 | 40.48 | 50.79 |
| | OLD | SN15 | O16_SN15 | 2001 | 1 | 1 | 0 | 0.11 | 0.99 | 3.94 | 34.59 | 34.70 | 34.97 |
| | OLD | WAC13405 | O42_WAC13405 | 2011 | 1 | 1 | 0 | 0.16 | 1.11 | 5.77 | 36.34 | 41.00 | 36.78 |
| | OLD | WAC13447 | O45_WAC13447 | 2011 | 1 | 1 | 0 | 0.14 | 1.33 | 8.19 | 56.64 | 76.17 | 57.39 |
| Group 4 | DPIRD | WAC2796 | DIP33_WAC2796 | 1980 | 8 | 7 | 1 | 6.21 | 8.97 | 198.60 | 31.97 | 290.19 | 32.34 |
| | DPIRD | WAC 4312 | DIP108_WAC4312 | 1985 | 9 | 9 | 0 | 6.88 | 9.95 | 218.91 | 31.80 | 320.53 | 32.22 |
| | DPIRD | WAC13958 | DIP73_WAC13958 | 2016 | 8 | 8 | 0 | 6.16 | 8.81 | 197.76 | 32.10 | 285.91 | 32.46 |
| | DPIRD | WAC13959 | DIP74_WAC13959 | 2016 | 9 | 9 | 0 | 7.43 | 8.45 | 2638.43 | 355.30 | 3055.81 | 361.77 |
| | DPIRD | WAC13960 | DIP75_WAC13960 | 2016 | 8 | 7 | 1 | 6.67 | 9.30 | 214.21 | 32.11 | 302.09 | 32.49 |
| | OLD | WAC13070 | O19_WAC13070 | 2005 | 10 | 10 | 0 | 9.11 | 10.96 | 463.05 | 50.82 | 564.01 | 51.44 |
| | OLD | WAC13074 | O23_WAC13074 | 2005 | 10 | 10 | 0 | 8.61 | 10.38 | 272.26 | 31.61 | 332.41 | 32.04 |
| | OLD | WAC13075 | O24_WAC13075 | 2005 | 10 | 10 | 0 | 8.80 | 10.31 | 498.09 | 56.58 | 590.60 | 57.31 |
| | OLD | WAC13076 | O25_WAC13076 | 2005 | 10 | 10 | 0 | 8.69 | 10.57 | 312.12 | 35.90 | 383.98 | 36.33 |
| | OLD | WAC13403 | O40_WAC13403 | 2011 | 10 | 10 | 0 | 8.57 | 10.29 | 286.98 | 33.50 | 348.75 | 33.89 |
| | OLD | WAC13404 | O41_WAC13404 | 2011 | 10 | 10 | 0 | 9.10 | 10.63 | 282.68 | 31.07 | 333.94 | 31.43 |
| | OLD | WAC13446 | O44_WAC13446 | 2011 | 10 | 10 | 0 | 8.18 | 9.96 | 268.80 | 32.87 | 331.39 | 33.27 |
| | OLD | WAC13525 | O48_WAC13525 | 2011 | 10 | 10 | 0 | 8.49 | 10.09 | 321.36 | 37.85 | 386.43 | 38.30 |
| | OLD | WAC13526 | O49_WAC13526 | 2011 | 10 | 10 | 0 | 7.75 | 9.79 | 539.85 | 69.69 | 690.67 | 70.55 |
| | OLD | WAC13528 | O51_WAC13528 | 2011 | 10 | 10 | 0 | 9.70 | 11.45 | 334.51 | 34.47 | 399.29 | 34.88 |
| | OLD | WAC13532 | O55_WAC13532 | 2011 | 10 | 10 | 0 | 8.27 | 10.01 | 419.22 | 50.67 | 514.05 | 51.33 |
| | OLD | WAC13630 | O59_WAC13630 | 2012 | 10 | 10 | 0 | 7.72 | 9.62 | 278.89 | 36.14 | 352.15 | 36.61 |
| | OLD | WAC13631 | O60_WAC13631 | 2012 | 10 | 10 | 0 | 8.55 | 10.30 | 233.68 | 27.32 | 284.71 | 27.65 |
| | OLD | WAC13632 | O61_WAC13632 | 2012 | 10 | 10 | 0 | 9.52 | 11.36 | 327.62 | 34.43 | 395.84 | 34.84 |
| Group 5 | OLD | 16FG160 | O146_16FG160 | 2016 | 44 | 26 | 18 | 42.13 | 43.72 | 5851.10 | 138.89 | 6132.86 | 140.28 |
| | OLD | 16FG161 | O147_16FG161 | 2016 | 45 | 27 | 18 | 42.99 | 44.74 | 7368.13 | 171.39 | 7747.97 | 173.19 |
| | OLD | 16FG162 | O148_16FG162 | 2016 | 45 | 26 | 19 | 43.41 | 45.01 | 5950.50 | 137.09 | 6232.78 | 138.48 |
| | OLD | 16FG163_2 | O150_16FG163_2 | 2016 | 43 | 26 | 17 | 42.02 | 43.45 | 6272.76 | 149.27 | 6557.85 | 150.92 |
| | OLD | 16FG164 | O151_16FG164 | 2016 | 43 | 26 | 17 | 41.99 | 43.51 | 6096.14 | 145.17 | 6381.01 | 146.67 |
| | OLD | 16FG165 | O152_16FG165 | 2016 | 44 | 26 | 18 | 42.82 | 44.27 | 5868.37 | 137.06 | 6131.18 | 138.49 |
| | OLD | 16FG166 | O153_16FG166 | 2016 | 41 | 25 | 16 | 39.30 | 40.78 | 5511.54 | 140.24 | 5774.35 | 141.59 |
| | OLD | 16FG167 | O154_16FG167 | 2016 | 44 | 25 | 19 | 42.78 | 44.27 | 6061.58 | 141.69 | 6336.79 | 143.14 |
| | OLD | 16FG168 | O155_16FG168 | 2016 | 44 | 26 | 18 | 43.33 | 44.75 | 7060.10 | 162.94 | 7370.69 | 164.71 |
| | OLD | 16FG169 | O156_16FG169 | 2016 | 42 | 25 | 18 | 41.30 | 42.82 | 6033.59 | 146.10 | 6340.24 | 148.07 |
| | OLD | 16FG170 | O158_16FG170 | 2016 | 44 | 25 | 19 | 42.44 | 43.95 | 5672.28 | 133.65 | 5956.86 | 135.53 |
| | OLD | 16FG171 | O159_16FG171 | 2016 | 45 | 26 | 19 | 43.77 | 45.11 | 6877.00 | 157.11 | 7194.20 | 159.49 |
| Group 6 | DPIRD | WAC1201 | DIP10_WAC1201 | 1968 | 0 | 0 | 0 | 0.05 | 1.45 | 1.49 | 32.19 | 47.05 | 32.55 |
| | DPIRD | WAC 1198 | DIP109_WAC1198 | 1968 | 0 | 0 | 0 | 0.02 | 1.50 | 0.61 | 31.83 | 48.45 | 32.21 |
| | DPIRD | WAC 1194 | DIP111_WAC1194 | 1968 | 0 | 0 | 0 | 0.01 | 1.43 | 0.37 | 37.64 | 54.77 | 38.19 |
| | DPIRD | WAC1188 | DIP4_WAC1188 | 1968 | 0 | 0 | 0 | 0.02 | 1.67 | 0.61 | 31.82 | 53.64 | 32.20 |
| | DPIRD | WAC1190 | DIP5_WAC1190 | 1968 | 0 | 0 | 0 | 0.02 | 1.56 | 0.53 | 31.74 | 50.11 | 32.14 |
| | DPIRD | WAC1193 | DIP6_WAC1193 | 1968 | 0 | 0 | 0 | 0.02 | 1.28 | 0.70 | 32.27 | 41.69 | 32.65 |
| | DPIRD | WAC1195 | DIP7_WAC1195 | 1968 | 0 | 0 | 0 | 0.02 | 1.98 | 0.61 | 31.70 | 63.72 | 32.11 |
| | DPIRD | WAC1197 | DIP8_WAC1197 | 1968 | 0 | 0 | 0 | 0.01 | 1.68 | 0.41 | 31.89 | 54.29 | 32.28 |
| | DPIRD | WAC1199 | DIP9_WAC1199 | 1968 | 0 | 0 | 0 | 0.01 | 1.47 | 0.38 | 38.10 | 56.65 | 38.56 |

| | | | | | | | | | | | | | |
|---|---|---|---|---|---|---|---|---|---|---|---|---|---|
| | DPIRD | WAC1497 | DIP14_WAC1497 | 1969 | 0 | 0 | 0 | 0.01 | 1.62 | 0.48 | 31.77 | 52.25 | 32.16 |
| | DPIRD | WAC1565 | DIP28_WAC1565 | 1969 | 0 | 0 | 0 | 0.02 | 1.48 | 0.53 | 31.81 | 47.59 | 32.20 |
| Group 7 | Current | PN280 | C280_PN280 | 2019 | 38 | 10 | 28 | 37.40 | 38.83 | 4269.65 | 114.16 | 4482.41 | 115.45 |
| | Current | PN282 | C282_PN282 | 2019 | 38 | 12 | 26 | 36.81 | 38.40 | 4214.16 | 114.49 | 4434.49 | 115.48 |
| | Current | PN283 | C283_PN283 | 2019 | 41 | 11 | 30 | 40.52 | 42.18 | 4429.17 | 109.32 | 4650.94 | 110.27 |
| | Current | PN306 | C306_PN306 | 2019 | 39 | 8 | 31 | 37.50 | 39.27 | 2476.36 | 66.04 | 2617.44 | 66.66 |
| | Current | PN307 | C307_PN307 | 2019 | 40 | 9 | 31 | 38.17 | 39.84 | 3118.87 | 81.71 | 3287.03 | 82.50 |
| | Current | PN315 | C315_PN315 | 2019 | 37 | 11 | 27 | 35.50 | 37.07 | 2175.55 | 61.28 | 2293.29 | 61.86 |
| | Current | PN316 | C316_PN316 | 2019 | 39 | 15 | 24 | 37.23 | 39.06 | 2408.55 | 64.70 | 2551.09 | 65.32 |
| | Current | PN334 | C334_PN334 | 2019 | 41 | 11 | 30 | 39.36 | 41.08 | 3013.03 | 76.55 | 3175.94 | 77.31 |
| | Current | PN335 | C335_PN335 | 2019 | 39 | 12 | 27 | 37.74 | 39.47 | 2611.47 | 69.19 | 2757.63 | 69.86 |
| | Current | PN337 | C337_PN337 | 2019 | 38 | 12 | 26 | 37.26 | 39.04 | 2877.58 | 77.22 | 3042.41 | 77.94 |
| | Current | PN338 | C338_PN338 | 2019 | 36 | 12 | 24 | 36.35 | 38.05 | 2042.83 | 56.20 | 2159.92 | 56.77 |
| | Current | PN340 | C340_PN340 | 2019 | 37 | 12 | 25 | 36.15 | 37.81 | 2523.50 | 69.81 | 2665.59 | 70.49 |
| | Current | PN343 | C343_PN343 | 2019 | 34 | 11 | 23 | 33.59 | 35.19 | 2883.05 | 85.82 | 3049.41 | 86.65 |
| | Current | PN345 | C345_PN345 | 2019 | 42 | 13 | 29 | 40.91 | 42.73 | 2621.11 | 64.06 | 2763.96 | 64.68 |
| | Current | PN346 | C346_PN346 | 2019 | 37 | 12 | 25 | 37.40 | 38.98 | 2397.48 | 64.10 | 2522.14 | 64.71 |
| Group 8 | Current | PN296 | C296_PN296 | 2019 | 2 | 2 | 0 | 0.45 | 1.89 | 31.29 | 68.89 | 131.59 | 69.54 |
| | Current | PN298 | C298_PN298 | 2019 | 2 | 2 | 0 | 0.48 | 1.98 | 33.16 | 69.62 | 138.91 | 70.28 |
| | Current | PN321 | C321_PN321 | 2019 | 2 | 2 | 0 | 0.55 | 1.96 | 36.73 | 66.94 | 132.24 | 67.57 |
| | Current | PN322 | C322_PN322 | 2019 | 2 | 2 | 0 | 0.52 | 1.90 | 27.98 | 54.16 | 104.10 | 54.68 |
| | Current | PN333 | C333_PN333 | 2019 | 1 | 1 | 0 | 0.40 | 1.50 | 17.28 | 43.34 | 65.68 | 43.78 |
| | Current | PN341 | C341_PN341 | 2019 | 1 | 1 | 0 | 0.22 | 1.00 | 14.73 | 67.35 | 67.93 | 68.04 |
| | Current | PN344 | C344_PN344 | 2019 | 2 | 2 | 0 | 0.55 | 1.64 | 28.17 | 51.33 | 85.05 | 51.82 |
| | Current | PN350 | C350_PN350 | 2019 | 2 | 2 | 0 | 0.52 | 1.70 | 40.82 | 78.15 | 134.16 | 78.88 |
| | Current | PN355 | C355_PN355 | 2021 | 1 | 1 | 0 | 0.52 | 1.67 | 36.11 | 69.93 | 117.93 | 70.58 |
| | Current | PN356 | C356_PN356 | 2021 | 2 | 2 | 0 | 0.50 | 1.83 | 27.78 | 55.33 | 102.25 | 55.86 |
| | Current | PN359 | C359_PN359 | 2021 | 2 | 2 | 0 | 0.52 | 1.89 | 36.18 | 69.02 | 131.67 | 69.63 |
| | Current | PN365 | C365_PN365 | 2021 | 2 | 2 | 0 | 0.52 | 1.75 | 31.31 | 60.23 | 106.25 | 60.82 |
| | Current | PN368 | C368_PN368 | 2021 | 2 | 2 | 0 | 0.51 | 1.83 | 30.81 | 60.09 | 111.01 | 60.63 |
| | Current | PN370 | C370_PN370 | 2021 | 2 | 2 | 0 | 0.55 | 1.81 | 37.68 | 68.29 | 124.76 | 68.94 |
| | Current | PN373 | C373_PN373 | 2021 | 2 | 2 | 0 | 0.56 | 1.80 | 33.73 | 60.02 | 109.08 | 60.59 |
| | Current | PN374 | C374_PN374 | 2021 | 2 | 2 | 0 | 0.59 | 1.81 | 36.97 | 62.35 | 113.67 | 62.94 |
| | Current | PN378 | C378_PN378 | 2021 | 2 | 2 | 0 | 0.51 | 2.07 | 33.71 | 65.47 | 136.82 | 66.06 |
| | Current | PN379 | C379_PN379 | 2021 | 2 | 2 | 0 | 0.52 | 1.77 | 27.01 | 51.82 | 92.44 | 52.36 |

**STable 3**: Effectors sensitivity profiles of 7 wheat lines

| Wheat | ToxA | Tox1 | Tox3 | Tox5 | Tox267 | Released time* | SNB disease mean |
|---|---|---|---|---|---|---|---|
| 1. Halberd | Sensitive | Sensitive | Sensitive | Sensitive | Sensitive | 1969 | 6.24 |
| 2. Eradu | Sensitive | Insensitive | Sensitive | NA | NA | 1981 | 3.73 |
| 3. Calingiri | Insensitive | Sensitive | Insensitive | Sensitive | Sensitive | 1997 | 3.97 |
| 4. Wyalkatchem | Insensitive | Weak sensitive | Sensitive | Insensitive | Sensitive | 2001 | 3.13 |
| 5. Mace | Insensitive | Insensitive | Sensitive | Sensitive | Weak sensitive | 2008 | 3.01 |
| 6. Calibre | Insensitive | Insensitive | Sensitive | Insensitive | Intermediate | 2021 | 2.82 |
| 7. Scepter | Insensitive | Insensitive | Sensitive | Insensitive | Intermediate | 2015 | 2.75 |

*Source: https://library.dpird.wa.gov.au/cgi/viewcontent.cgi?article=1171&context=bulletins; https://www.grainland.com.au/slides/slide/lancer-wheat-

#### STable 4: QC genome assemblies using Quast

| Assembly | # contigs (>= 0 bp) | # contigs (>= 1000 bp) | # contigs (>= 5000 bp) | # contigs (>= 10000 bp) | # contigs (>= 25000 bp) | # contigs (>= 50000 bp) | Total length (>= 0 bp) | Total length (>= 1000 bp) | Total length (>= 5000 bp) | Total length (>= 10000 bp) | Total length (>= 25000 bp) | Total length (>= 50000 bp) | # contigs | Largest contig | Total length | Reference length | GC (%) | Reference GC (%) | N50 | NG50 | N90 | NG90 | auN | auNG | L50 | LG50 | L90 | LG90 | # mis-assemblies | [illegible] contigs |
|---|---|---|---|---|---|---|---|---|---|---|---|---|---|---|---|---|---|---|---|---|---|---|---|---|---|---|---|---|---|---|
| DPIRD1 | 3583 | 476 | 253 | 192 | 141 | 122 | 4E+07 | 4E+07 | 4E+07 | 4E+07 | 3E+07 | 3E+07 | 590 | 1E+06 | 3.7E+07 | 3.7E+07 | 50.42 | 50.13 | 340374 | 339717 | 82714 | 65884 | 414504 | 407406 | 33 | 34 | 105 | 113 | 213 | 111 |
| DPIRD10 | 3039 | 506 | 226 | 149 | 102 | 93 | 4E+07 | 4E+07 | 4E+07 | 4E+07 | 3E+07 | 3E+07 | 645 | 2E+06 | 3.7E+07 | 3.7E+07 | 50.51 | 50.13 | 557439 | 557439 | 108195 | 89669 | 606682 | 595782 | 22 | 22 | 77 | 83 | 199 | 100 |
| DPIRD100 | 3038 | 397 | 211 | 148 | 93 | 77 | 4E+07 | 4E+07 | 4E+07 | 4E+07 | 4E+07 | 3E+07 | 551 | 2E+06 | 3.7E+07 | 3.7E+07 | 50.42 | 50.13 | 641956 | 641956 | 119211 | 102783 | 706520 | 698454 | 18 | 18 | 64 | 68 | 206 | 89 |
| DPIRD102 | 3427 | 458 | 259 | 196 | 148 | 125 | 4E+07 | 4E+07 | 4E+07 | 4E+07 | 3E+07 | 3E+07 | 552 | 967760 | 3.7E+07 | 3.7E+07 | 50.41 | 50.13 | 331157 | 327737 | 86036 | 64464 | 393127 | 386417 | 35 | 36 | 110 | 118 | 216 | 105 |
| DPIRD103 | 3419 | 536 | 289 | 207 | 150 | 126 | 4E+07 | 4E+07 | 4E+07 | 4E+07 | 4E+07 | 3E+07 | 645 | 1E+06 | 3.7E+07 | 3.7E+07 | 50.3 | 50.13 | 333969 | 330786 | 71654 | 66953 | 400669 | 398214 | 34 | 35 | 113 | 116 | 202 | 106 |
| DPIRD104 | 2526 | 365 | 224 | 189 | 136 | 117 | 4E+07 | 4E+07 | 4E+07 | 4E+07 | 4E+07 | 4E+07 | 414 | 2E+06 | 3.7E+07 | 3.7E+07 | 50.25 | 50.13 | 386358 | 386358 | 95585 | 93714 | 487575 | 485722 | 30 | 30 | 97 | 98 | 204 | 98 |
| DPIRD105 | 2589 | 382 | 243 | 194 | 154 | 132 | 4E+07 | 4E+07 | 4E+07 | 4E+07 | 4E+07 | 3E+07 | 444 | 950583 | 3.6E+07 | 3.7E+07 | 50.52 | 50.13 | 330704 | 324970 | 86604 | 64297 | 381928 | 372804 | 35 | 36 | 110 | 120 | 204 | 107 |
| DPIRD106 | 3460 | 517 | 268 | 220 | 154 | 136 | 4E+07 | 4E+07 | 4E+07 | 4E+07 | 4E+07 | 3E+07 | 622 | 958037 | 3.7E+07 | 3.7E+07 | 50.05 | 50.13 | 325598 | 325598 | 78571 | 79813 | 361989 | 362962 | 38 | 38 | 120 | 119 | 222 | 116 |
| DPIRD107 | 4817 | 591 | 248 | 184 | 129 | 112 | 4E+07 | 4E+07 | 4E+07 | 4E+07 | 4E+07 | 3E+07 | ### | 1E+06 | 3.7E+07 | 3.7E+07 | 50.43 | 50.13 | 390927 | 390927 | 89091 | 88020 | 459726 | 458978 | 29 | 29 | 101 | 102 | 203 | 109 |
| DPIRD108 | 2462 | 364 | 223 | 176 | 132 | 112 | 4E+07 | 4E+07 | 4E+07 | 4E+07 | 4E+07 | 3E+07 | 417 | 2E+06 | 3.7E+07 | 3.7E+07 | 50.36 | 50.13 | 403322 | 401312 | 114233 | 95538 | 475230 | 468817 | 29 | 30 | 93 | 97 | 229 | 105 |
| DPIRD109 | 2755 | 483 | 292 | 235 | 177 | 148 | 4E+07 | 4E+07 | 4E+07 | 4E+07 | 4E+07 | 3E+07 | 561 | 1E+06 | 3.7E+07 | 3.7E+07 | 50.45 | 50.13 | 305269 | 295476 | 67613 | 63118 | 344734 | 339313 | 38 | 39 | 135 | 143 | 195 | 110 |
| DPIRD11 | 318 | 63 | 53 | 48 | 45 | 42 | 4E+07 | 4E+07 | 4E+07 | 4E+07 | 4E+07 | 4E+07 | 84 | 2E+06 | 3.7E+07 | 3.7E+07 | 50.21 | 50.13 | 1E+06 | 1E+06 | 383045 | 365514 | 1E+06 | 1E+06 | 12 | 12 | 28 | 29 | 317 | 43 |
| DPIRD110 | 3093 | 483 | 298 | 226 | 167 | 138 | 4E+07 | 4E+07 | 4E+07 | 4E+07 | 4E+07 | 3E+07 | 563 | 943175 | 3.8E+07 | 3.7E+07 | 50.33 | 50.13 | 327653 | 329503 | 69843 | 78767 | 360485 | 363937 | 38 | 37 | 126 | 121 | 195 | 98 |
| DPIRD111 | 2620 | 384 | 243 | 188 | 140 | 122 | 4E+07 | 4E+07 | 4E+07 | 4E+07 | 4E+07 | 3E+07 | 451 | 2E+06 | 3.7E+07 | 3.7E+07 | 50.44 | 50.13 | 349643 | 349031 | 102448 | 73198 | 449091 | 442178 | 32 | 33 | 101 | 107 | 195 | 88 |
| DPIRD112 | 3343 | 465 | 259 | 194 | 137 | 118 | 4E+07 | 4E+07 | 4E+07 | 4E+07 | 4E+07 | 3E+07 | 547 | 1E+06 | 3.7E+07 | 3.7E+07 | 50.31 | 50.13 | 372011 | 372011 | 89325 | 87487 | 465208 | 462068 | 29 | 29 | 102 | 104 | 213 | 101 |
| DPIRD12 | 3378 | 512 | 283 | 214 | 157 | 131 | 4E+07 | 4E+07 | 4E+07 | 4E+07 | 3E+07 | 3E+07 | 624 | 1E+06 | 3.7E+07 | 3.7E+07 | 50.4 | 50.13 | 340905 | 336183 | 69019 | 59627 | 376916 | 370706 | 35 | 36 | 120 | 128 | 207 | 104 |
| DPIRD13 | 1436 | 330 | 152 | 131 | 120 | 107 | 4E+07 | 3E+07 | 3E+07 | 3E+07 | 3E+07 | 3E+07 | 548 | 1E+06 | 3.5E+07 | 3.7E+07 | 51.72 | 50.13 | 414392 | 373910 | 125851 | 52372 | 512920 | 480311 | 25 | 28 | 82 | 106 | 151 | 89 |
| DPIRD14 | 2874 | 482 | 279 | 206 | 157 | 135 | 4E+07 | 4E+07 | 4E+07 | 4E+07 | 4E+07 | 3E+07 | 583 | 1E+06 | 3.7E+07 | 3.7E+07 | 50.46 | 50.13 | 334851 | 330726 | 70595 | 63850 | 378021 | 371854 | 37 | 38 | 117 | 125 | 192 | 101 |
| DPIRD15 | 12314 | ### | 671 | 526 | 362 | 244 | 4E+07 | 4E+07 | 4E+07 | 3E+07 | 3E+07 | 3E+07 | ### | 818438 | 3.8E+07 | 3.7E+07 | 50.09 | 50.13 | 93356 | 95926 | 11348 | 16947 | 123006 | 126208 | 124 | 118 | 504 | 441 | 168 | 132 |
| DPIRD16 | 562 | 79 | 71 | 60 | 53 | 47 | 4E+07 | 4E+07 | 4E+07 | 4E+07 | 4E+07 | 4E+07 | 134 | 3E+06 | 3.7E+07 | 3.7E+07 | 50.19 | 50.13 | 1E+06 | 1E+06 | 356156 | 326111 | 1E+06 | 1E+06 | 11 | 11 | 29 | 30 | 308 | 49 |
| DPIRD17 | 2539 | 455 | 256 | 198 | 155 | 133 | 4E+07 | 4E+07 | 4E+07 | 4E+07 | 3E+07 | 3E+07 | 526 | 1E+06 | 3.7E+07 | 3.7E+07 | 50.43 | 50.13 | 306460 | 302989 | 82018 | 63139 | 382363 | 374520 | 35 | 36 | 114 | 124 | 224 | 104 |
| DPIRD18 | 2040 | 309 | 176 | 143 | 104 | 93 | 4E+07 | 4E+07 | 4E+07 | 4E+07 | 4E+07 | 4E+07 | 347 | 2E+06 | 3.7E+07 | 3.7E+07 | 50.45 | 50.13 | 494262 | 493513 | 162600 | 116770 | 608347 | 598400 | 22 | 23 | 69 | 73 | 201 | 78 |
| DPIRD19 | 3896 | 547 | 280 | 214 | 152 | 130 | 4E+07 | 4E+07 | 4E+07 | 4E+07 | 4E+07 | 3E+07 | 655 | 937970 | 3.7E+07 | 3.7E+07 | 50.28 | 50.13 | 349732 | 344000 | 67604 | 66449 | 388866 | 387326 | 34 | 35 | 117 | 119 | 190 | 106 |
| DPIRD2 | 3470 | 418 | 201 | 146 | 101 | 86 | 4E+07 | 4E+07 | 4E+07 | 4E+07 | 4E+07 | 3E+07 | 561 | 2E+06 | 3.7E+07 | 3.7E+07 | 50.62 | 50.13 | 557688 | 556375 | 117330 | 115597 | 697539 | 689929 | 20 | 21 | 70 | 73 | 207 | 81 |
| DPIRD20 | 2347 | 446 | 160 | 116 | 104 | 93 | 4E+07 | 4E+07 | 3E+07 | 3E+07 | 3E+07 | 3E+07 | 777 | 2E+06 | 3.6E+07 | 3.7E+07 | 51.38 | 50.13 | 503627 | 443715 | 119871 | 64797 | 638242 | 606438 | 20 | 22 | 73 | 90 | 184 | 97 |
| DPIRD21 | 2764 | 323 | 183 | 142 | 101 | 90 | 4E+07 | 4E+07 | 4E+07 | 4E+07 | 4E+07 | 3E+07 | 393 | 2E+06 | 3.6E+07 | 3.7E+07 | 50.54 | 50.13 | 490259 | 489699 | 158859 | 133847 | 587581 | 572951 | 23 | 24 | 71 | 76 | 195 | 89 |

| | | | | | | | | | | | | | | | | | | | | | | | | | | | | | | |
|---|---|---|---|---|---|---|---|---|---|---|---|---|---|---|---|---|---|---|---|---|---|---|---|---|---|---|---|---|---|---|
| DPIRD22 | 3109 | 418 | 223 | 153 | 109 | 99 | 4E+07 | 4E+07 | 4E+07 | 4E+07 | 4E+07 | 4E+07 | 508 | 2E+06 | 3.7E+07 | 3.7E+07 | 50.3 | 50.13 | 423336 | 423336 | 137628 | 125968 | 575317 | 570462 | 26 | 26 | 82 | 84 | 230 | 100 |
| DPIRD23 | 3043 | 584 | 197 | 146 | 115 | 101 | 4E+07 | 4E+07 | 3E+07 | 3E+07 | 3E+07 | 3E+07 | 909 | 2E+06 | 3.6E+07 | 3.7E+07 | 50.76 | 50.13 | 426778 | 415670 | 113888 | 49689 | 535259 | 515708 | 26 | 27 | 86 | 103 | 178 | 102 |
| DPIRD24 | 3627 | 498 | 235 | 163 | 106 | 90 | 4E+07 | 4E+07 | 4E+07 | 4E+07 | 3E+07 | 3E+07 | 650 | 2E+06 | 3.7E+07 | 3.7E+07 | 50.36 | 50.13 | 460150 | 460150 | 133857 | 104900 | 598506 | 592180 | 24 | 24 | 77 | 80 | 207 | 90 |
| DPIRD25 | 3044 | 459 | 188 | 131 | 95 | 87 | 4E+07 | 4E+07 | 4E+07 | 4E+07 | 4E+07 | 3E+07 | 581 | 1E+06 | 3.7E+07 | 3.7E+07 | 50.38 | 50.13 | 553398 | 549085 | 132658 | 125630 | 642860 | 634091 | 20 | 21 | 69 | 72 | 207 | 89 |
| DPIRD27 | 3534 | 494 | 279 | 213 | 158 | 134 | 4E+07 | 4E+07 | 4E+07 | 4E+07 | 3E+07 | 3E+07 | 600 | 1E+06 | 3.7E+07 | 3.7E+07 | 50.41 | 50.13 | 328709 | 319551 | 68328 | 61118 | 393559 | 386996 | 34 | 35 | 120 | 128 | 206 | 99 |
| DPIRD28 | 3003 | 507 | 270 | 203 | 148 | 128 | 4E+07 | 4E+07 | 4E+07 | 4E+07 | 3E+07 | 3E+07 | 620 | 2E+06 | 3.7E+07 | 3.7E+07 | 50.47 | 50.13 | 330126 | 330126 | 85910 | 64694 | 427787 | 420727 | 34 | 34 | 114 | 121 | 189 | 92 |
| DPIRD29 | 2837 | 333 | 192 | 147 | 111 | 94 | 4E+07 | 4E+07 | 4E+07 | 4E+07 | 4E+07 | 3E+07 | 384 | 2E+06 | 3.7E+07 | 3.7E+07 | 50.36 | 50.13 | 480246 | 475972 | 132827 | 108026 | 637261 | 626500 | 23 | 24 | 72 | 77 | 231 | 95 |
| DPIRD3 | 3565 | 475 | 240 | 144 | 99 | 83 | 4E+07 | 4E+07 | 4E+07 | 4E+07 | 4E+07 | 3E+07 | 594 | 1E+06 | 3.7E+07 | 3.7E+07 | 50.29 | 50.13 | 558120 | 558120 | 145404 | 117467 | 609194 | 604902 | 22 | 22 | 71 | 73 | 216 | 93 |
| DPIRD30 | 388 | 62 | 52 | 49 | 47 | 43 | 4E+07 | 4E+07 | 4E+07 | 4E+07 | 4E+07 | 4E+07 | 87 | 4E+06 | 3.7E+07 | 3.7E+07 | 50.22 | 50.13 | 1E+06 | 1E+06 | 422830 | 417007 | 1E+06 | 1E+06 | 11 | 12 | 29 | 30 | 316 | 43 |
| DPIRD31 | 2445 | 494 | 178 | 135 | 103 | 95 | 4E+07 | 4E+07 | 4E+07 | 4E+07 | 3E+07 | 3E+07 | 659 | 2E+06 | 3.6E+07 | 3.7E+07 | 50.62 | 50.13 | 496709 | 493698 | 121650 | 102820 | 597017 | 580496 | 22 | 23 | 76 | 84 | 202 | 99 |
| DPIRD32 | 2917 | 606 | 224 | 136 | 105 | 94 | 4E+07 | 4E+07 | 4E+07 | 3E+07 | 3E+07 | 3E+07 | 819 | 2E+06 | 3.7E+07 | 3.7E+07 | 50.6 | 50.13 | 516488 | 478678 | 105391 | 74170 | 569199 | 555806 | 24 | 25 | 80 | 88 | 226 | 118 |
| DPIRD33 | 2222 | 353 | 199 | 140 | 101 | 86 | 4E+07 | 4E+07 | 4E+07 | 4E+07 | 4E+07 | 3E+07 | 407 | 2E+06 | 3.7E+07 | 3.7E+07 | 50.35 | 50.13 | 546779 | 532564 | 137830 | 133413 | 656115 | 647610 | 20 | 21 | 69 | 72 | 235 | 87 |
| DPIRD34 | 509 | 112 | 86 | 71 | 62 | 52 | 4E+07 | 4E+07 | 4E+07 | 4E+07 | 4E+07 | 4E+07 | 163 | 2E+06 | 3.8E+07 | 3.7E+07 | 50.25 | 50.13 | 1E+06 | 1E+06 | 361690 | 365056 | 1E+06 | 1E+06 | 14 | 14 | 37 | 36 | 288 | 51 |
| DPIRD35 | 2899 | 504 | 294 | 218 | 165 | 141 | 4E+07 | 4E+07 | 4E+07 | 4E+07 | 4E+07 | 3E+07 | 580 | 897737 | 3.7E+07 | 3.7E+07 | 50.32 | 50.13 | 298557 | 295148 | 66570 | 62580 | 339257 | 336397 | 39 | 40 | 129 | 133 | 213 | 126 |
| DPIRD36 | 2967 | 499 | 294 | 224 | 158 | 135 | 4E+07 | 4E+07 | 4E+07 | 4E+07 | 4E+07 | 3E+07 | 573 | 961846 | 3.7E+07 | 3.7E+07 | 50.2 | 50.13 | 334025 | 331750 | 71821 | 68407 | 345265 | 343045 | 39 | 40 | 122 | 125 | 222 | 101 |
| DPIRD37 | 3450 | 419 | 228 | 163 | 119 | 104 | 4E+07 | 4E+07 | 4E+07 | 4E+07 | 4E+07 | 3E+07 | 518 | 1E+06 | 3.7E+07 | 3.7E+07 | 50.47 | 50.13 | 432210 | 431597 | 119706 | 100908 | 469660 | 461136 | 28 | 29 | 88 | 93 | 208 | 106 |
| DPIRD38 | 2636 | 357 | 190 | 139 | 109 | 96 | 4E+07 | 4E+07 | 4E+07 | 4E+07 | 4E+07 | 3E+07 | 428 | 1E+06 | 3.6E+07 | 3.7E+07 | 50.53 | 50.13 | 478907 | 472602 | 124686 | 105215 | 558635 | 544943 | 24 | 25 | 75 | 83 | 199 | 88 |
| DPIRD39 | 1333 | 337 | 157 | 128 | 116 | 105 | 4E+07 | 3E+07 | 3E+07 | 3E+07 | 3E+07 | 3E+07 | 609 | 2E+06 | 3.5E+07 | 3.7E+07 | 51.67 | 50.13 | 472914 | 456570 | 109456 | 58399 | 592649 | 554956 | 22 | 24 | 78 | 104 | 153 | 78 |
| DPIRD4 | 2715 | 461 | 268 | 199 | 149 | 130 | 4E+07 | 4E+07 | 4E+07 | 4E+07 | 4E+07 | 3E+07 | 564 | 1E+06 | 3.7E+07 | 3.7E+07 | 50.45 | 50.13 | 342814 | 342743 | 71501 | 65735 | 429495 | 422682 | 32 | 33 | 111 | 119 | 192 | 97 |
| DPIRD40 | 3914 | 455 | 236 | 147 | 105 | 97 | 4E+07 | 4E+07 | 4E+07 | 4E+07 | 4E+07 | 3E+07 | 590 | 1E+06 | 3.7E+07 | 3.7E+07 | 50.28 | 50.13 | 524870 | 502579 | 104424 | 95498 | 536948 | 532377 | 25 | 26 | 78 | 81 | 15 | 13 |
| DPIRD41 | 2844 | 470 | 285 | 201 | 142 | 127 | 4E+07 | 4E+07 | 4E+07 | 4E+07 | 4E+07 | 3E+07 | 539 | 1E+06 | 3.7E+07 | 3.7E+07 | 50.35 | 50.13 | 329277 | 323510 | 88549 | 85948 | 415102 | 411471 | 34 | 35 | 112 | 115 | 198 | 101 |
| DPIRD42 | 2818 | 451 | 289 | 243 | 188 | 149 | 4E+07 | 4E+07 | 4E+07 | 4E+07 | 4E+07 | 3E+07 | 506 | 1E+06 | 3.7E+07 | 3.7E+07 | 50.37 | 50.13 | 285381 | 284966 | 62616 | 53541 | 349929 | 344635 | 39 | 40 | 138 | 147 | 231 | 105 |
| DPIRD43 | 1806 | 491 | 182 | 139 | 121 | 107 | 4E+07 | 4E+07 | 3E+07 | 3E+07 | 3E+07 | 3E+07 | 874 | 1E+06 | 3.6E+07 | 3.7E+07 | 51.34 | 50.13 | 421602 | 383137 | 106411 | 49431 | 512307 | 488436 | 25 | 27 | 88 | 108 | 134 | 67 |
| DPIRD44 | 3375 | 371 | 202 | 150 | 100 | 90 | 4E+07 | 4E+07 | 4E+07 | 4E+07 | 4E+07 | 3E+07 | 443 | 2E+06 | 3.7E+07 | 3.7E+07 | 50.47 | 50.13 | 682718 | 682718 | 131809 | 113111 | 701534 | 688775 | 19 | 19 | 72 | 77 | 212 | 96 |
| DPIRD45 | 3638 | 473 | 185 | 143 | 119 | 106 | 4E+07 | 4E+07 | 3E+07 | 3E+07 | 3E+07 | 3E+07 | 984 | 1E+06 | 3.6E+07 | 3.7E+07 | 51.27 | 50.13 | 445376 | 438716 | 92807 | 56429 | 547408 | 523987 | 23 | 25 | 85 | 104 | 177 | 92 |
| DPIRD46 | 3747 | 501 | 308 | 226 | 171 | 146 | 4E+07 | 4E+07 | 4E+07 | 4E+07 | 4E+07 | 3E+07 | 613 | 880926 | 3.7E+07 | 3.7E+07 | 50.23 | 50.13 | 300480 | 293630 | 66987 | 66094 | 332464 | 330496 | 40 | 41 | 134 | 137 | 15 | 13 |
| DPIRD47 | 2965 | 634 | 213 | 150 | 108 | 99 | 4E+07 | 4E+07 | 4E+07 | 4E+07 | 3E+07 | 3E+07 | 900 | 1E+06 | 3.7E+07 | 3.7E+07 | 50.51 | 50.13 | 476209 | 471157 | 102063 | 82823 | 556958 | 546091 | 24 | 25 | 84 | 91 | 212 | 109 |
| DPIRD48 | 1561 | 457 | 184 | 137 | 115 | 101 | 4E+07 | 4E+07 | 3E+07 | 3E+07 | 3E+07 | 3E+07 | 817 | 2E+06 | 3.6E+07 | 3.7E+07 | 51.29 | 50.13 | 450015 | 424868 | 114102 | 51160 | 560710 | 534391 | 23 | 25 | 82 | 101 | 166 | 90 |
| DPIRD49 | 3817 | 616 | 231 | 157 | 111 | 95 | 4E+07 | 4E+07 | 4E+07 | 4E+07 | 3E+07 | 3E+07 | 838 | 1E+06 | 3.7E+07 | 3.7E+07 | 50.44 | 50.13 | 463860 | 463860 | 100117 | 85791 | 544423 | 537279 | 25 | 25 | 85 | 89 | 228 | 102 |
| DPIRD5 | 2904 | 432 | 260 | 192 | 139 | 122 | 4E+07 | 4E+07 | 4E+07 | 4E+07 | 4E+07 | 3E+07 | 525 | 1E+06 | 3.7E+07 | 3.7E+07 | 50.46 | 50.13 | 372542 | 372542 | 93830 | 71477 | 413939 | 407190 | 32 | 32 | 104 | 110 | 198 | 101 |
| DPIRD50 | 2932 | 509 | 273 | 197 | 156 | 134 | 4E+07 | 4E+07 | 4E+07 | 4E+07 | 3E+07 | 3E+07 | 620 | 2E+06 | 3.7E+07 | 3.7E+07 | 50.49 | 50.13 | 302455 | 301853 | 78610 | 61232 | 404935 | 397797 | 35 | 36 | 117 | 126 | 214 | 119 |
| DPIRD51 | 2456 | 423 | 257 | 181 | 128 | 113 | 4E+07 | 4E+07 | 4E+07 | 4E+07 | 4E+07 | 4E+07 | 497 | 1E+06 | 3.8E+07 | 3.7E+07 | 50.36 | 50.13 | 428770 | 428770 | 101319 | 102671 | 479441 | 481508 | 29 | 29 | 93 | 92 | 208 | 100 |

| | | | | | | | | | | | | | | | | | | | | | | | | | | | | | | |
|---|---|---|---|---|---|---|---|---|---|---|---|---|---|---|---|---|---|---|---|---|---|---|---|---|---|---|---|---|---|---|
| DPIRD55 | 4405 | 653 | 305 | 220 | 154 | 130 | 4E+07 | 4E+07 | 4E+07 | 4E+07 | 4E+07 | 3E+07 | 803 | 1E+06 | 3.8E+07 | 3.7E+07 | 50.05 | 50.13 | 329958 | 329958 | 68702 | 77994 | 380140 | 382541 | 36 | 36 | 125 | 122 | 216 | 112 |
| DPIRD56 | 4563 | 694 | 392 | 269 | 174 | 143 | 4E+07 | 4E+07 | 4E+07 | 4E+07 | 4E+07 | 3E+07 | 902 | 990172 | 3.9E+07 | 3.7E+07 | 50.16 | 50.13 | 300620 | 308689 | 42341 | 67447 | 354489 | 365531 | 39 | 37 | 152 | 133 | 233 | 133 |
| DPIRD57 | 5323 | 802 | 373 | 242 | 184 | 157 | 4E+07 | 4E+07 | 4E+07 | 4E+07 | 4E+07 | 3E+07 | ### | 1E+06 | 3.8E+07 | 3.7E+07 | 50.29 | 50.13 | 271523 | 272332 | 47751 | 60337 | 326135 | 334182 | 44 | 42 | 162 | 146 | 200 | 122 |
| DPIRD58 | 3913 | 691 | 386 | 274 | 201 | 164 | 4E+07 | 4E+07 | 4E+07 | 4E+07 | 4E+07 | 3E+07 | 880 | 1E+06 | 3.8E+07 | 3.7E+07 | 50.36 | 50.13 | 241372 | 244130 | 46061 | 53464 | 314165 | 319899 | 46 | 44 | 173 | 161 | 218 | 108 |
| DPIRD6 | 2837 | 516 | 215 | 133 | 100 | 87 | 4E+07 | 4E+07 | 4E+07 | 4E+07 | 3E+07 | 3E+07 | 696 | 2E+06 | 3.7E+07 | 3.7E+07 | 50.54 | 50.13 | 605328 | 569105 | 114874 | 89690 | 661579 | 648788 | 20 | 21 | 72 | 78 | 183 | 83 |
| DPIRD60 | 8647 | 949 | 362 | 254 | 173 | 147 | 4E+07 | 4E+07 | 4E+07 | 4E+07 | 4E+07 | 3E+07 | ### | 1E+06 | 3.9E+07 | 3.7E+07 | 50.2 | 50.13 | 284377 | 291643 | 32059 | 65943 | 315693 | 328623 | 45 | 43 | 166 | 137 | 202 | 113 |
| DPIRD62 | 2877 | 339 | 185 | 130 | 94 | 76 | 4E+07 | 4E+07 | 4E+07 | 4E+07 | 4E+07 | 3E+07 | 416 | 2E+06 | 3.6E+07 | 3.7E+07 | 50.56 | 50.13 | 594692 | 593868 | 169043 | 110611 | 662382 | 645882 | 20 | 21 | 60 | 67 | 190 | 78 |
| DPIRD63 | 8438 | ### | 431 | 283 | 195 | 163 | 4E+07 | 4E+07 | 4E+07 | 4E+07 | 3E+07 | 3E+07 | ### | 965132 | 3.9E+07 | 3.7E+07 | 50.33 | 50.13 | 252308 | 264900 | 16077 | 40632 | 274275 | 286073 | 50 | 47 | 230 | 173 | 204 | 131 |
| DPIRD64 | 3106 | 495 | 241 | 180 | 134 | 116 | 4E+07 | 4E+07 | 4E+07 | 4E+07 | 3E+07 | 3E+07 | 658 | 2E+06 | 3.7E+07 | 3.7E+07 | 50.49 | 50.13 | 402198 | 384504 | 83126 | 66551 | 468560 | 460885 | 29 | 30 | 98 | 106 | 205 | 105 |
| DPIRD66 | 2439 | 338 | 176 | 134 | 93 | 82 | 4E+07 | 4E+07 | 4E+07 | 4E+07 | 4E+07 | 3E+07 | 416 | 1E+06 | 3.6E+07 | 3.7E+07 | 50.49 | 50.13 | 563072 | 560224 | 164213 | 128019 | 616849 | 601552 | 22 | 23 | 65 | 71 | 211 | 92 |
| DPIRD68 | 2657 | 362 | 207 | 144 | 106 | 92 | 4E+07 | 4E+07 | 4E+07 | 4E+07 | 4E+07 | 3E+07 | 419 | 1E+06 | 3.7E+07 | 3.7E+07 | 50.46 | 50.13 | 524561 | 492911 | 143534 | 112260 | 593419 | 583648 | 22 | 23 | 73 | 77 | 193 | 79 |
| DPIRD69 | 2529 | 377 | 214 | 145 | 105 | 93 | 4E+07 | 4E+07 | 4E+07 | 4E+07 | 4E+07 | 3E+07 | 447 | 2E+06 | 3.7E+07 | 3.7E+07 | 50.44 | 50.13 | 482638 | 464907 | 133533 | 110599 | 604673 | 595225 | 23 | 24 | 76 | 81 | 196 | 90 |
| DPIRD7 | 2292 | 395 | 277 | 212 | 163 | 134 | 4E+07 | 4E+07 | 4E+07 | 4E+07 | 4E+07 | 3E+07 | 455 | 1E+06 | 3.7E+07 | 3.7E+07 | 50.41 | 50.13 | 311973 | 301340 | 84094 | 69444 | 374979 | 369600 | 36 | 37 | 118 | 125 | 202 | 94 |
| DPIRD70 | 2767 | 485 | 231 | 150 | 111 | 101 | 4E+07 | 4E+07 | 4E+07 | 4E+07 | 3E+07 | 3E+07 | 607 | 2E+06 | 3.7E+07 | 3.7E+07 | 50.51 | 50.13 | 465355 | 464141 | 102851 | 91070 | 534911 | 524918 | 26 | 27 | 84 | 90 | 197 | 90 |
| DPIRD71 | 2449 | 433 | 273 | 206 | 157 | 133 | 4E+07 | 4E+07 | 4E+07 | 4E+07 | 4E+07 | 3E+07 | 481 | 964533 | 3.7E+07 | 3.7E+07 | 50.45 | 50.13 | 304506 | 303541 | 86145 | 67379 | 360048 | 354354 | 38 | 39 | 118 | 125 | 196 | 96 |
| DPIRD72 | 2689 | 383 | 225 | 175 | 141 | 120 | 4E+07 | 4E+07 | 4E+07 | 4E+07 | 4E+07 | 3E+07 | 461 | 1E+06 | 3.6E+07 | 3.7E+07 | 50.54 | 50.13 | 397362 | 382750 | 101443 | 73439 | 447602 | 436397 | 29 | 30 | 99 | 109 | 193 | 97 |
| DPIRD73 | 2572 | 382 | 194 | 147 | 105 | 92 | 4E+07 | 4E+07 | 4E+07 | 4E+07 | 4E+07 | 3E+07 | 448 | 2E+06 | 3.7E+07 | 3.7E+07 | 50.38 | 50.13 | 548016 | 548016 | 116401 | 102413 | 610025 | 601121 | 22 | 22 | 71 | 75 | 232 | 98 |
| DPIRD74 | 264 | 58 | 49 | 41 | 39 | 36 | 4E+07 | 4E+07 | 4E+07 | 4E+07 | 4E+07 | 4E+07 | 70 | 4E+06 | 3.7E+07 | 3.7E+07 | 50.23 | 50.13 | 1E+06 | 1E+06 | 513885 | 482239 | 1E+06 | 1E+06 | 11 | 11 | 26 | 27 | 323 | 38 |
| DPIRD75 | 2450 | 362 | 204 | 152 | 115 | 98 | 4E+07 | 4E+07 | 4E+07 | 4E+07 | 4E+07 | 3E+07 | 424 | 2E+06 | 3.7E+07 | 3.7E+07 | 50.36 | 50.13 | 481259 | 479653 | 125592 | 110986 | 597449 | 589382 | 23 | 24 | 78 | 82 | 227 | 97 |
| DPIRD76 | 3001 | 405 | 225 | 176 | 132 | 118 | 4E+07 | 4E+07 | 4E+07 | 4E+07 | 4E+07 | 3E+07 | 487 | 2E+06 | 3.7E+07 | 3.7E+07 | 50.33 | 50.13 | 398871 | 398871 | 93546 | 79804 | 516190 | 508300 | 27 | 27 | 96 | 101 | 231 | 102 |
| DPIRD77 | 3344 | 354 | 192 | 147 | 104 | 92 | 4E+07 | 4E+07 | 4E+07 | 4E+07 | 4E+07 | 4E+07 | 459 | 1E+06 | 3.7E+07 | 3.7E+07 | 50.29 | 50.13 | 579839 | 578431 | 132701 | 116620 | 619520 | 611828 | 21 | 22 | 71 | 74 | 224 | 99 |
| DPIRD79 | 3974 | 512 | 253 | 204 | 156 | 138 | 4E+07 | 4E+07 | 4E+07 | 4E+07 | 4E+07 | 3E+07 | 852 | 1E+06 | 3.7E+07 | 3.7E+07 | 50.32 | 50.13 | 330303 | 330303 | 79707 | 73935 | 372334 | 370254 | 36 | 36 | 121 | 123 | 221 | 108 |
| DPIRD8 | 2514 | 397 | 259 | 200 | 144 | 130 | 4E+07 | 4E+07 | 4E+07 | 4E+07 | 4E+07 | 3E+07 | 461 | 1E+06 | 3.7E+07 | 3.7E+07 | 50.43 | 50.13 | 337563 | 323178 | 101483 | 87082 | 410267 | 404124 | 33 | 34 | 110 | 116 | 201 | 93 |
| DPIRD82 | 5729 | 881 | 314 | 212 | 137 | 108 | 4E+07 | 4E+07 | 4E+07 | 4E+07 | 4E+07 | 3E+07 | ### | 1E+06 | 3.9E+07 | 3.7E+07 | 50.48 | 50.13 | 416346 | 419112 | 34383 | 58887 | 466227 | 483038 | 30 | 29 | 126 | 100 | 190 | 93 |
| DPIRD83 | 7080 | ### | 324 | 215 | 159 | 142 | 4E+07 | 4E+07 | 4E+07 | 4E+07 | 3E+07 | 3E+07 | ### | 805296 | 3.8E+07 | 3.7E+07 | 50.46 | 50.13 | 272596 | 288688 | 32473 | 63770 | 326281 | 333871 | 41 | 40 | 155 | 138 | 202 | 123 |
| DPIRD84 | 3942 | 425 | 216 | 161 | 120 | 104 | 4E+07 | 4E+07 | 4E+07 | 4E+07 | 3E+07 | 3E+07 | 651 | 1E+06 | 3.7E+07 | 3.7E+07 | 50.53 | 50.13 | 432761 | 426083 | 109902 | 88513 | 466526 | 456477 | 29 | 30 | 87 | 95 | 208 | 107 |
| DPIRD85 | 3001 | 368 | 215 | 168 | 133 | 106 | 4E+07 | 4E+07 | 4E+07 | 4E+07 | 4E+07 | 3E+07 | 440 | 1E+06 | 3.7E+07 | 3.7E+07 | 50.51 | 50.13 | 425257 | 422099 | 101152 | 94936 | 475941 | 468916 | 28 | 29 | 90 | 95 | 224 | 98 |
| DPIRD86 | 600 | 88 | 74 | 59 | 50 | 40 | 4E+07 | 4E+07 | 4E+07 | 4E+07 | 4E+07 | 4E+07 | 136 | 2E+06 | 3.7E+07 | 3.7E+07 | 50.37 | 50.13 | 1E+06 | 1E+06 | 455629 | 455629 | 1E+06 | 1E+06 | 11 | 12 | 28 | 28 | 303 | 47 |
| DPIRD87 | 3027 | 402 | 235 | 167 | 132 | 108 | 4E+07 | 4E+07 | 4E+07 | 4E+07 | 4E+07 | 3E+07 | 491 | 1E+06 | 3.7E+07 | 3.7E+07 | 50.53 | 50.13 | 444054 | 425506 | 98370 | 77343 | 471479 | 464306 | 28 | 29 | 92 | 98 | 216 | 100 |
| DPIRD88 | 3047 | 489 | 258 | 180 | 132 | 111 | 4E+07 | 4E+07 | 4E+07 | 4E+07 | 4E+07 | 3E+07 | 629 | 1E+06 | 3.7E+07 | 3.7E+07 | 50.53 | 50.13 | 383758 | 383758 | 88435 | 73783 | 474558 | 468777 | 30 | 30 | 98 | 103 | 215 | 93 |
| DPIRD89 | 3684 | 623 | 235 | 151 | 107 | 93 | 4E+07 | 4E+07 | 4E+07 | 4E+07 | 3E+07 | 3E+07 | 897 | 2E+06 | 3.7E+07 | 3.7E+07 | 50.52 | 50.13 | 518210 | 518210 | 82711 | 65561 | 607928 | 603631 | 22 | 22 | 79 | 83 | 200 | 87 |
| DPIRD9 | 2699 | 381 | 221 | 162 | 125 | 106 | 4E+07 | 4E+07 | 4E+07 | 4E+07 | 4E+07 | 3E+07 | 448 | 1E+06 | 3.7E+07 | 3.7E+07 | 50.45 | 50.13 | 444105 | 440553 | 107147 | 92892 | 485246 | 477630 | 27 | 28 | 86 | 91 | 191 | 89 |

| | | | | | | | | | | | | | | | | | | | | | | | | | | | | | | |
|---|---|---|---|---|---|---|---|---|---|---|---|---|---|---|---|---|---|---|---|---|---|---|---|---|---|---|---|---|---|---|
| DPIRD90 | 3226 | 463 | 252 | 191 | 140 | 122 | 4E+07 | 4E+07 | 4E+07 | 4E+07 | 3E+07 | 3E+07 | 567 | 2E+06 | 3.7E+07 | 3.7E+07 | 50.41 | 50.13 | 362479 | 350675 | 78560 | 68684 | 449737 | 442781 | 32 | 33 | 105 | 112 | 212 | 107 |
| DPIRD91 | 3105 | 381 | 207 | 155 | 118 | 101 | 4E+07 | 4E+07 | 4E+07 | 4E+07 | 4E+07 | 3E+07 | 484 | 2E+06 | 3.7E+07 | 3.7E+07 | 50.47 | 50.13 | 438430 | 437976 | 123323 | 112133 | 531963 | 521628 | 26 | 27 | 83 | 88 | 216 | 103 |
| DPIRD92 | 2161 | 342 | 213 | 173 | 132 | 113 | 4E+07 | 4E+07 | 4E+07 | 4E+07 | 4E+07 | 3E+07 | 409 | 1E+06 | 3.7E+07 | 3.7E+07 | 50.35 | 50.13 | 357079 | 351737 | 110624 | 100425 | 444748 | 436886 | 31 | 32 | 94 | 100 | 222 | 95 |
| DPIRD93 | 2910 | 455 | 249 | 190 | 151 | 127 | 4E+07 | 4E+07 | 4E+07 | 4E+07 | 4E+07 | 3E+07 | 540 | 1E+06 | 3.7E+07 | 3.7E+07 | 50.52 | 50.13 | 335176 | 331147 | 81951 | 65017 | 385688 | 377895 | 36 | 37 | 111 | 120 | 215 | 110 |
| DPIRD94 | 3159 | 579 | 336 | 274 | 206 | 169 | 4E+07 | 4E+07 | 4E+07 | 4E+07 | 4E+07 | 3E+07 | 736 | 1E+06 | 3.7E+07 | 3.7E+07 | 50.42 | 50.13 | 233897 | 231271 | 50980 | 49854 | 302667 | 301638 | 48 | 49 | 167 | 170 | 206 | 111 |
| DPIRD96 | 2438 | 336 | 197 | 149 | 101 | 91 | 4E+07 | 4E+07 | 4E+07 | 4E+07 | 4E+07 | 3E+07 | 381 | 2E+06 | 3.7E+07 | 3.7E+07 | 50.5 | 50.13 | 493036 | 493036 | 132823 | 113858 | 622242 | 610529 | 23 | 23 | 73 | 78 | 221 | 99 |
| DPIRD97 | 2760 | 375 | 203 | 149 | 105 | 100 | 4E+07 | 4E+07 | 4E+07 | 4E+07 | 4E+07 | 3E+07 | 441 | 2E+06 | 3.7E+07 | 3.7E+07 | 50.51 | 50.13 | 493324 | 482887 | 133824 | 110559 | 578051 | 566868 | 25 | 26 | 79 | 84 | 215 | 102 |
| PN260 | 491 | 100 | 87 | 79 | 68 | 62 | 4E+07 | 4E+07 | 4E+07 | 4E+07 | 4E+07 | 4E+07 | 138 | 2E+06 | 3.7E+07 | 3.7E+07 | 50.39 | 50.13 | 955514 | 951458 | 298977 | 278363 | 856530 | 840693 | 16 | 17 | 44 | 46 | 276 | 65 |
| PN261 | 496 | 129 | 97 | 85 | 69 | 61 | 4E+07 | 4E+07 | 4E+07 | 4E+07 | 4E+07 | 4E+07 | 173 | 2E+06 | 3.7E+07 | 3.7E+07 | 50.29 | 50.13 | 1E+06 | 1E+06 | 278940 | 249133 | 956262 | 941051 | 15 | 15 | 39 | 41 | 270 | 67 |
| PN262 | 26514 | ### | 140 | 74 | 58 | 53 | 5E+07 | 4E+07 | 4E+07 | 4E+07 | 4E+07 | 4E+07 | ### | 2E+06 | 4.4E+07 | 3.7E+07 | 47.53 | 50.13 | 862020 | 968044 | 1047 | 312593 | 831110 | 980095 | 18 | 15 | ### | 41 | 275 | 52 |
| PN263 | 947 | 245 | 134 | 99 | 69 | 54 | 4E+07 | 4E+07 | 4E+07 | 4E+07 | 4E+07 | 4E+07 | 378 | 2E+06 | 3.7E+07 | 3.7E+07 | 50.34 | 50.13 | 1E+06 | 1E+06 | 258234 | 258234 | 1E+06 | 1E+06 | 13 | 12 | 40 | 40 | 260 | 58 |
| PN264 | 407 | 94 | 83 | 69 | 59 | 53 | 4E+07 | 4E+07 | 4E+07 | 4E+07 | 4E+07 | 4E+07 | 126 | 4E+06 | 3.7E+07 | 3.7E+07 | 50.35 | 50.13 | 1E+06 | 1E+06 | 310074 | 300564 | 1E+06 | 1E+06 | 13 | 13 | 37 | 39 | 257 | 57 |
| PN265 | 515 | 138 | 115 | 91 | 79 | 61 | 4E+07 | 4E+07 | 4E+07 | 4E+07 | 4E+07 | 4E+07 | 175 | 2E+06 | 3.7E+07 | 3.7E+07 | 50.22 | 50.13 | 963151 | 963151 | 282787 | 277508 | 906842 | 899373 | 15 | 15 | 45 | 46 | 297 | 61 |
| PN266 | 639 | 111 | 83 | 73 | 59 | 52 | 4E+07 | 4E+07 | 4E+07 | 4E+07 | 4E+07 | 4E+07 | 150 | 2E+06 | 3.7E+07 | 3.7E+07 | 50.36 | 50.13 | 996241 | 996241 | 356165 | 330927 | 1E+06 | 1E+06 | 14 | 14 | 37 | 39 | 289 | 53 |
| PN268 | 594 | 116 | 91 | 75 | 63 | 55 | 4E+07 | 4E+07 | 4E+07 | 4E+07 | 4E+07 | 4E+07 | 157 | 2E+06 | 3.7E+07 | 3.7E+07 | 50.23 | 50.13 | 1E+06 | 1E+06 | 338675 | 321164 | 1E+06 | 1E+06 | 13 | 13 | 38 | 39 | 287 | 56 |
| PN269 | 524 | 141 | 102 | 97 | 88 | 83 | 4E+07 | 4E+07 | 4E+07 | 4E+07 | 4E+07 | 4E+07 | 190 | 2E+06 | 4.2E+07 | 3.7E+07 | 50.7 | 50.13 | 840597 | 945867 | 227339 | 351567 | 905469 | 1E+06 | 18 | 15 | 54 | 39 | 289 | 52 |
| PN270 | 428 | 108 | 82 | 76 | 65 | 55 | 4E+07 | 4E+07 | 4E+07 | 4E+07 | 4E+07 | 4E+07 | 144 | 3E+06 | 3.7E+07 | 3.7E+07 | 50.34 | 50.13 | 1E+06 | 1E+06 | 313136 | 307477 | 1E+06 | 1E+06 | 13 | 13 | 37 | 39 | 294 | 63 |
| PN271 | 492 | 102 | 80 | 71 | 62 | 55 | 4E+07 | 4E+07 | 4E+07 | 4E+07 | 4E+07 | 4E+07 | 137 | 3E+06 | 3.7E+07 | 3.7E+07 | 50.28 | 50.13 | 1E+06 | 1E+06 | 353403 | 340576 | 1E+06 | 1E+06 | 13 | 13 | 35 | 36 | 291 | 62 |
| PN274 | 2935 | 131 | 89 | 75 | 65 | 56 | 4E+07 | 4E+07 | 4E+07 | 4E+07 | 4E+07 | 4E+07 | 328 | 2E+06 | 3.7E+07 | 3.7E+07 | 50.32 | 50.13 | 1E+06 | 1E+06 | 359371 | 333182 | 1E+06 | 1E+06 | 14 | 14 | 39 | 40 | 294 | 64 |
| PN275 | 469 | 134 | 102 | 86 | 75 | 65 | 4E+07 | 4E+07 | 4E+07 | 4E+07 | 4E+07 | 4E+07 | 165 | 3E+06 | 3.7E+07 | 3.7E+07 | 50.33 | 50.13 | 858511 | 858511 | 268090 | 232699 | 1E+06 | 1E+06 | 14 | 14 | 41 | 43 | 266 | 61 |
| PN276 | 782 | 159 | 60 | 54 | 50 | 46 | 4E+07 | 4E+07 | 4E+07 | 4E+07 | 4E+07 | 4E+07 | 313 | 3E+06 | 3.7E+07 | 3.7E+07 | 50.2 | 50.13 | 1E+06 | 1E+06 | 394873 | 394873 | 1E+06 | 1E+06 | 13 | 13 | 32 | 32 | 304 | 43 |
| PN277 | 446 | 107 | 91 | 68 | 56 | 51 | 4E+07 | 4E+07 | 4E+07 | 4E+07 | 4E+07 | 4E+07 | 151 | 3E+06 | 3.7E+07 | 3.7E+07 | 50.26 | 50.13 | 1E+06 | 1E+06 | 343272 | 338031 | 1E+06 | 1E+06 | 12 | 12 | 35 | 36 | 292 | 50 |
| PN279 | 413 | 106 | 85 | 68 | 57 | 49 | 4E+07 | 4E+07 | 4E+07 | 4E+07 | 4E+07 | 4E+07 | 135 | 2E+06 | 3.7E+07 | 3.7E+07 | 50.29 | 50.13 | 1E+06 | 1E+06 | 351544 | 338634 | 1E+06 | 1E+06 | 13 | 13 | 34 | 36 | 276 | 47 |
| PN280 | 477 | 155 | 120 | 98 | 83 | 73 | 4E+07 | 4E+07 | 4E+07 | 4E+07 | 4E+07 | 4E+07 | 188 | 3E+06 | 3.7E+07 | 3.7E+07 | 50.33 | 50.13 | 710082 | 702205 | 229237 | 215592 | 946215 | 939235 | 16 | 17 | 51 | 52 | 308 | 75 |
| PN282 | 498 | 144 | 116 | 92 | 78 | 66 | 4E+07 | 4E+07 | 4E+07 | 4E+07 | 4E+07 | 4E+07 | 174 | 3E+06 | 3.7E+07 | 3.7E+07 | 50.33 | 50.13 | 809429 | 809429 | 268572 | 247221 | 905521 | 898970 | 16 | 16 | 48 | 49 | 310 | 69 |
| PN283 | 465 | 143 | 119 | 98 | 86 | 69 | 4E+07 | 4E+07 | 4E+07 | 4E+07 | 4E+07 | 4E+07 | 167 | 3E+06 | 3.7E+07 | 3.7E+07 | 50.33 | 50.13 | 728004 | 728004 | 235753 | 230183 | 970629 | 963172 | 16 | 16 | 50 | 51 | 308 | 72 |
| PN284 | 266 | 97 | 80 | 70 | 60 | 57 | 4E+07 | 4E+07 | 4E+07 | 4E+07 | 4E+07 | 4E+07 | 119 | 2E+06 | 3.7E+07 | 3.7E+07 | 50.25 | 50.13 | 1E+06 | 1E+06 | 297432 | 281618 | 1E+06 | 1E+06 | 13 | 13 | 40 | 41 | 271 | 53 |
| PN285 | 447 | 116 | 95 | 76 | 71 | 69 | 4E+07 | 4E+07 | 4E+07 | 4E+07 | 4E+07 | 4E+07 | 151 | 2E+06 | 3.7E+07 | 3.7E+07 | 50.37 | 50.13 | 892301 | 810998 | 270475 | 248865 | 837424 | 825316 | 16 | 17 | 47 | 49 | 237 | 65 |
| PN286 | 389 | 121 | 93 | 86 | 70 | 66 | 4E+07 | 4E+07 | 4E+07 | 4E+07 | 4E+07 | 4E+07 | 160 | 2E+06 | 3.7E+07 | 3.7E+07 | 50.5 | 50.13 | 878290 | 853928 | 261150 | 245069 | 841817 | 822623 | 16 | 17 | 48 | 51 | 263 | 64 |
| PN287 | 416 | 127 | 103 | 87 | 73 | 65 | 4E+07 | 4E+07 | 4E+07 | 4E+07 | 4E+07 | 4E+07 | 161 | 2E+06 | 3.7E+07 | 3.7E+07 | 50.33 | 50.13 | 824607 | 824607 | 259848 | 235074 | 933459 | 920412 | 15 | 15 | 46 | 48 | 288 | 66 |
| PN288 | 454 | 137 | 111 | 100 | 84 | 77 | 4E+07 | 4E+07 | 4E+07 | 4E+07 | 4E+07 | 4E+07 | 174 | 2E+06 | 3.7E+07 | 3.7E+07 | 50.36 | 50.13 | 743938 | 681426 | 233280 | 201278 | 831340 | 817121 | 16 | 17 | 54 | 56 | 259 | 72 |
| PN289 | 422 | 118 | 98 | 82 | 66 | 59 | 4E+07 | 4E+07 | 4E+07 | 4E+07 | 4E+07 | 4E+07 | 152 | 2E+06 | 3.7E+07 | 3.7E+07 | 50.35 | 50.13 | 955768 | 955768 | 271819 | 257019 | 923285 | 907570 | 15 | 15 | 43 | 45 | 253 | 63 |

| | | | | | | | | | | | | | | | | | | | | | | | | | | | | | | |
|---|---|---|---|---|---|---|---|---|---|---|---|---|---|---|---|---|---|---|---|---|---|---|---|---|---|---|---|---|---|---|
| PN290 | 470 | 167 | 132 | 99 | 85 | 76 | 4E+07 | 4E+07 | 4E+07 | 4E+07 | 4E+07 | 4E+07 | 198 | 2E+06 | 4.1E+07 | 3.7E+07 | 49.41 | 50.13 | 886018 | 909018 | 246608 | 323719 | 877797 | 957628 | 17 | 16 | 52 | 41 | 312 | 67 |
| PN291 | 574 | 107 | 81 | 73 | 65 | 57 | 4E+07 | 4E+07 | 4E+07 | 4E+07 | 4E+07 | 4E+07 | 142 | 2E+06 | 3.8E+07 | 3.7E+07 | 50.41 | 50.13 | 1E+06 | 1E+06 | 327698 | 348877 | 1E+06 | 1E+06 | 14 | 14 | 39 | 38 | 295 | 59 |
| PN292 | 475 | 166 | 133 | 112 | 93 | 86 | 4E+07 | 4E+07 | 4E+07 | 4E+07 | 4E+07 | 4E+07 | 198 | 2E+06 | 3.7E+07 | 3.7E+07 | 50.28 | 50.13 | 652090 | 645295 | 176460 | 166053 | 731767 | 720225 | 19 | 20 | 59 | 62 | 278 | 78 |
| PN295 | 928 | 286 | 181 | 144 | 111 | 92 | 4E+07 | 4E+07 | 4E+07 | 4E+07 | 4E+07 | 4E+07 | 399 | 2E+06 | 3.8E+07 | 3.7E+07 | 50.19 | 50.13 | 550715 | 552517 | 152087 | 184162 | 624318 | 640262 | 23 | 22 | 70 | 65 | 266 | 79 |
| PN296 | 354 | 100 | 76 | 68 | 60 | 56 | 4E+07 | 4E+07 | 4E+07 | 4E+07 | 4E+07 | 4E+07 | 124 | 2E+06 | 3.7E+07 | 3.7E+07 | 50.47 | 50.13 | 1E+06 | 1E+06 | 332672 | 307071 | 1E+06 | 1E+06 | 14 | 14 | 35 | 38 | 281 | 58 |
| PN297 | 3242 | 264 | 177 | 131 | 97 | 81 | 4E+07 | 4E+07 | 4E+07 | 4E+07 | 4E+07 | 4E+07 | 754 | 2E+06 | 3.8E+07 | 3.7E+07 | 49.96 | 50.13 | 590021 | 614403 | 150846 | 170329 | 775730 | 787060 | 19 | 18 | 61 | 58 | 314 | 100 |
| PN298 | 347 | 96 | 76 | 70 | 62 | 54 | 4E+07 | 4E+07 | 4E+07 | 4E+07 | 4E+07 | 4E+07 | 130 | 3E+06 | 3.7E+07 | 3.7E+07 | 50.46 | 50.13 | 980039 | 946190 | 320367 | 312970 | 1E+06 | 1E+06 | 13 | 14 | 39 | 41 | 274 | 60 |
| PN299 | 380 | 119 | 95 | 83 | 71 | 61 | 4E+07 | 4E+07 | 4E+07 | 4E+07 | 4E+07 | 4E+07 | 142 | 2E+06 | 3.7E+07 | 3.7E+07 | 50.44 | 50.13 | 807044 | 807044 | 298829 | 261065 | 952615 | 936110 | 15 | 15 | 42 | 44 | 301 | 62 |
| PN300 | 630 | 226 | 167 | 143 | 114 | 98 | 4E+07 | 4E+07 | 4E+07 | 4E+07 | 4E+07 | 4E+07 | 293 | 2E+06 | 4.3E+07 | 3.7E+07 | 49.71 | 50.13 | 806934 | 896753 | 153482 | 325370 | 840139 | 972043 | 18 | 15 | 67 | 43 | 264 | 73 |
| PN301 | 370 | 127 | 102 | 91 | 71 | 66 | 4E+07 | 4E+07 | 4E+07 | 4E+07 | 4E+07 | 4E+07 | 167 | 2E+06 | 3.7E+07 | 3.7E+07 | 50.43 | 50.13 | 981010 | 981010 | 267339 | 210716 | 922302 | 906416 | 15 | 15 | 42 | 45 | 283 | 67 |
| PN302 | 324 | 111 | 88 | 74 | 56 | 52 | 4E+07 | 4E+07 | 4E+07 | 4E+07 | 4E+07 | 4E+07 | 140 | 3E+06 | 3.7E+07 | 3.7E+07 | 50.39 | 50.13 | 1E+06 | 1E+06 | 328623 | 271642 | 1E+06 | 1E+06 | 13 | 14 | 35 | 37 | 318 | 62 |
| PN303 | 419 | 101 | 84 | 70 | 61 | 52 | 4E+07 | 4E+07 | 4E+07 | 4E+07 | 4E+07 | 4E+07 | 127 | 2E+06 | 3.7E+07 | 3.7E+07 | 50.38 | 50.13 | 1E+06 | 1E+06 | 344969 | 304599 | 1E+06 | 1E+06 | 14 | 14 | 37 | 39 | 294 | 60 |
| PN304 | 478 | 120 | 97 | 84 | 73 | 69 | 4E+07 | 4E+07 | 4E+07 | 4E+07 | 4E+07 | 4E+07 | 159 | 3E+06 | 3.7E+07 | 3.7E+07 | 50.35 | 50.13 | 744301 | 744301 | 269738 | 248699 | 920123 | 907112 | 16 | 16 | 47 | 49 | 288 | 68 |
| PN305 | 376 | 104 | 82 | 71 | 61 | 52 | 4E+07 | 4E+07 | 4E+07 | 4E+07 | 4E+07 | 4E+07 | 134 | 2E+06 | 3.7E+07 | 3.7E+07 | 50.34 | 50.13 | 1E+06 | 1E+06 | 328080 | 313506 | 1E+06 | 1E+06 | 13 | 13 | 35 | 37 | 310 | 58 |
| PN306 | 532 | 178 | 135 | 112 | 91 | 79 | 4E+07 | 4E+07 | 4E+07 | 4E+07 | 4E+07 | 4E+07 | 210 | 3E+06 | 3.7E+07 | 3.7E+07 | 50.33 | 50.13 | 629527 | 629527 | 210163 | 206751 | 878770 | 872102 | 18 | 18 | 57 | 58 | 294 | 79 |
| PN307 | 509 | 172 | 134 | 113 | 88 | 76 | 4E+07 | 4E+07 | 4E+07 | 4E+07 | 4E+07 | 4E+07 | 201 | 3E+06 | 3.7E+07 | 3.7E+07 | 50.33 | 50.13 | 639152 | 639152 | 210463 | 194214 | 847746 | 841356 | 17 | 17 | 55 | 56 | 301 | 78 |
| PN308 | 451 | 127 | 97 | 80 | 65 | 59 | 4E+07 | 4E+07 | 4E+07 | 4E+07 | 4E+07 | 4E+07 | 155 | 2E+06 | 3.7E+07 | 3.7E+07 | 50.44 | 50.13 | 1E+06 | 1E+06 | 256502 | 210826 | 1E+06 | 999189 | 13 | 14 | 39 | 42 | 292 | 62 |
| PN309 | 643 | 219 | 162 | 131 | 100 | 86 | 4E+07 | 4E+07 | 4E+07 | 4E+07 | 4E+07 | 4E+07 | 282 | 2E+06 | 3.8E+07 | 3.7E+07 | 50.06 | 50.13 | 626692 | 626692 | 155049 | 155516 | 799302 | 803073 | 17 | 17 | 61 | 60 | 305 | 92 |
| PN312 | 565 | 143 | 112 | 95 | 77 | 65 | 4E+07 | 4E+07 | 4E+07 | 4E+07 | 4E+07 | 4E+07 | 174 | 2E+06 | 3.7E+07 | 3.7E+07 | 50.31 | 50.13 | 1E+06 | 1E+06 | 225969 | 203601 | 963475 | 953003 | 14 | 14 | 45 | 47 | 275 | 69 |
| PN313 | 444 | 125 | 100 | 86 | 73 | 60 | 4E+07 | 4E+07 | 4E+07 | 4E+07 | 4E+07 | 4E+07 | 148 | 2E+06 | 3.7E+07 | 3.7E+07 | 50.3 | 50.13 | 975392 | 975392 | 253568 | 212483 | 989246 | 978649 | 14 | 14 | 42 | 43 | 284 | 66 |
| PN314 | 809 | 134 | 97 | 80 | 65 | 56 | 4E+07 | 4E+07 | 4E+07 | 4E+07 | 4E+07 | 4E+07 | 191 | 2E+06 | 3.7E+07 | 3.7E+07 | 50.44 | 50.13 | 1E+06 | 1E+06 | 338838 | 267966 | 1E+06 | 985229 | 14 | 14 | 37 | 40 | 288 | 63 |
| PN315 | 561 | 190 | 139 | 116 | 95 | 82 | 4E+07 | 4E+07 | 4E+07 | 4E+07 | 4E+07 | 4E+07 | 226 | 2E+06 | 3.7E+07 | 3.7E+07 | 50.33 | 50.13 | 620021 | 614523 | 194159 | 191532 | 728353 | 723183 | 20 | 21 | 59 | 60 | 296 | 84 |
| PN316 | 650 | 245 | 182 | 148 | 106 | 89 | 4E+07 | 4E+07 | 4E+07 | 4E+07 | 4E+07 | 4E+07 | 281 | 1E+06 | 3.8E+07 | 3.7E+07 | 50.32 | 50.13 | 634289 | 653202 | 160835 | 191170 | 653535 | 667858 | 21 | 20 | 66 | 62 | 287 | 78 |
| PN317 | 1652 | 345 | 167 | 121 | 83 | 70 | 4E+07 | 4E+07 | 4E+07 | 4E+07 | 4E+07 | 4E+07 | 573 | 2E+06 | 3.9E+07 | 3.7E+07 | 50.34 | 50.13 | 855508 | 873346 | 175947 | 268238 | 917214 | 946641 | 15 | 14 | 51 | 46 | 266 | 58 |
| PN318 | 373 | 115 | 92 | 74 | 65 | 56 | 4E+07 | 4E+07 | 4E+07 | 4E+07 | 4E+07 | 4E+07 | 141 | 3E+06 | 3.7E+07 | 3.7E+07 | 50.39 | 50.13 | 1E+06 | 1E+06 | 311908 | 298394 | 1E+06 | 1E+06 | 13 | 13 | 37 | 38 | 279 | 52 |
| PN319 | 1151 | 232 | 137 | 107 | 83 | 68 | 4E+07 | 4E+07 | 4E+07 | 4E+07 | 4E+07 | 4E+07 | 369 | 2E+06 | 3.8E+07 | 3.7E+07 | 50.39 | 50.13 | 878765 | 968873 | 224252 | 233699 | 890261 | 898863 | 16 | 15 | 48 | 46 | 302 | 67 |
| PN320 | 252 | 93 | 73 | 64 | 58 | 54 | 4E+07 | 4E+07 | 4E+07 | 4E+07 | 4E+07 | 4E+07 | 106 | 2E+06 | 3.7E+07 | 3.7E+07 | 50.25 | 50.13 | 1E+06 | 1E+06 | 346639 | 343533 | 1E+06 | 1E+06 | 12 | 13 | 37 | 38 | 280 | 49 |
| PN321 | 447 | 139 | 103 | 86 | 75 | 67 | 4E+07 | 4E+07 | 4E+07 | 4E+07 | 4E+07 | 4E+07 | 180 | 2E+06 | 3.7E+07 | 3.7E+07 | 50.28 | 50.13 | 833993 | 833993 | 304301 | 284260 | 824095 | 812732 | 17 | 17 | 46 | 48 | 267 | 66 |
| PN322 | 511 | 173 | 131 | 104 | 87 | 73 | 4E+07 | 4E+07 | 4E+07 | 4E+07 | 4E+07 | 4E+07 | 216 | 1E+06 | 3.7E+07 | 3.7E+07 | 50.32 | 50.13 | 716681 | 712946 | 216484 | 200245 | 706429 | 695564 | 19 | 20 | 54 | 57 | 257 | 73 |
| PN323 | 329 | 119 | 95 | 84 | 72 | 65 | 4E+07 | 4E+07 | 4E+07 | 4E+07 | 4E+07 | 4E+07 | 151 | 2E+06 | 3.7E+07 | 3.7E+07 | 50.18 | 50.13 | 898961 | 898961 | 249592 | 249592 | 937028 | 933348 | 15 | 15 | 44 | 44 | 279 | 69 |
| PN324 | 2826 | 224 | 103 | 88 | 73 | 65 | 4E+07 | 4E+07 | 4E+07 | 4E+07 | 4E+07 | 4E+07 | 765 | 2E+06 | 3.8E+07 | 3.7E+07 | 50.04 | 50.13 | 857736 | 857736 | 243757 | 275572 | 812683 | 819702 | 17 | 17 | 47 | 46 | 288 | 77 |
| PN325 | 302 | 101 | 76 | 66 | 57 | 52 | 4E+07 | 4E+07 | 4E+07 | 4E+07 | 4E+07 | 4E+07 | 124 | 3E+06 | 3.6E+07 | 3.7E+07 | 50.55 | 50.13 | 1E+06 | 1E+06 | 311107 | 300271 | 1E+06 | 1E+06 | 12 | 13 | 37 | 39 | 274 | 54 |

| | | | | | | | | | | | | | | | | | | | | | | | | | | | | | | |
|---|---|---|---|---|---|---|---|---|---|---|---|---|---|---|---|---|---|---|---|---|---|---|---|---|---|---|---|---|---|---|
| PN326 | 403 | 130 | 107 | 82 | 72 | 66 | 4E+07 | 4E+07 | 4E+07 | 4E+07 | 4E+07 | 4E+07 | 162 | 3E+06 | 3.7E+07 | 3.7E+07 | 50.21 | 50.13 | 895052 | 895052 | 249534 | 249534 | 1E+06 | 1E+06 | 14 | 14 | 44 | 44 | 276 | 66 |
| PN328 | 701 | 228 | 152 | 125 | 91 | 78 | 4E+07 | 4E+07 | 4E+07 | 4E+07 | 4E+07 | 4E+07 | 282 | 2E+06 | 3.7E+07 | 3.7E+07 | 50.16 | 50.13 | 614600 | 614600 | 211031 | 211031 | 713140 | 710964 | 20 | 20 | 57 | 57 | 261 | 74 |
| PN329 | 590 | 198 | 147 | 122 | 95 | 76 | 4E+07 | 4E+07 | 4E+07 | 4E+07 | 4E+07 | 4E+07 | 238 | 2E+06 | 3.7E+07 | 3.7E+07 | 50.13 | 50.13 | 720951 | 720951 | 177339 | 177280 | 767536 | 765982 | 18 | 18 | 53 | 54 | 258 | 66 |
| PN330 | 425 | 110 | 93 | 77 | 65 | 62 | 4E+07 | 4E+07 | 4E+07 | 4E+07 | 4E+07 | 4E+07 | 146 | 3E+06 | 3.7E+07 | 3.7E+07 | 50.42 | 50.13 | 776096 | 776096 | 317770 | 286135 | 936506 | 922166 | 16 | 16 | 43 | 45 | 271 | 62 |
| PN331 | 460 | 111 | 89 | 82 | 70 | 64 | 4E+07 | 4E+07 | 4E+07 | 4E+07 | 4E+07 | 4E+07 | 140 | 2E+06 | 3.7E+07 | 3.7E+07 | 50.37 | 50.13 | 1E+06 | 1E+06 | 256202 | 215265 | 954310 | 936855 | 15 | 15 | 42 | 44 | 289 | 62 |
| PN332 | 552 | 160 | 119 | 100 | 82 | 69 | 4E+07 | 4E+07 | 4E+07 | 4E+07 | 4E+07 | 4E+07 | 204 | 2E+06 | 3.7E+07 | 3.7E+07 | 50.35 | 50.13 | 778054 | 778054 | 219213 | 218656 | 832699 | 824042 | 17 | 17 | 50 | 51 | 274 | 67 |
| PN333 | 10627 | 214 | 115 | 102 | 81 | 71 | 4E+07 | 4E+07 | 4E+07 | 4E+07 | 4E+07 | 4E+07 | ### | 2E+06 | 3.8E+07 | 3.7E+07 | 50.26 | 50.13 | 724441 | 724441 | 201773 | 211583 | 720165 | 731662 | 19 | 19 | 56 | 54 | 252 | 66 |
| PN334 | 516 | 160 | 126 | 102 | 88 | 75 | 4E+07 | 4E+07 | 4E+07 | 4E+07 | 4E+07 | 4E+07 | 195 | 3E+06 | 3.7E+07 | 3.7E+07 | 50.33 | 50.13 | 686278 | 686278 | 197398 | 194145 | 947474 | 940674 | 17 | 17 | 54 | 55 | 297 | 72 |
| PN335 | 481 | 169 | 133 | 108 | 95 | 79 | 4E+07 | 4E+07 | 4E+07 | 4E+07 | 4E+07 | 4E+07 | 205 | 1E+06 | 3.7E+07 | 3.7E+07 | 50.33 | 50.13 | 674976 | 674976 | 194667 | 194198 | 677901 | 672956 | 20 | 20 | 58 | 59 | 298 | 83 |
| PN336 | 464 | 113 | 85 | 75 | 62 | 59 | 4E+07 | 4E+07 | 4E+07 | 4E+07 | 4E+07 | 4E+07 | 138 | 2E+06 | 3.7E+07 | 3.7E+07 | 50.3 | 50.13 | 1E+06 | 990268 | 318564 | 289478 | 1E+06 | 1E+06 | 13 | 14 | 39 | 40 | 302 | 57 |
| PN337 | 456 | 195 | 157 | 130 | 112 | 94 | 4E+07 | 4E+07 | 4E+07 | 4E+07 | 4E+07 | 4E+07 | 220 | 2E+06 | 3.7E+07 | 3.7E+07 | 50.33 | 50.13 | 510465 | 510465 | 152796 | 145267 | 660650 | 655787 | 23 | 23 | 68 | 69 | 286 | 91 |
| PN338 | 591 | 189 | 148 | 117 | 95 | 82 | 4E+07 | 4E+07 | 4E+07 | 4E+07 | 4E+07 | 4E+07 | 230 | 2E+06 | 3.7E+07 | 3.7E+07 | 50.34 | 50.13 | 603189 | 603189 | 168721 | 167167 | 756823 | 750979 | 19 | 19 | 59 | 60 | 280 | 79 |
| PN340 | 494 | 167 | 133 | 108 | 91 | 80 | 4E+07 | 4E+07 | 4E+07 | 4E+07 | 4E+07 | 4E+07 | 195 | 3E+06 | 3.7E+07 | 3.7E+07 | 50.33 | 50.13 | 623064 | 623064 | 205929 | 194645 | 907313 | 900673 | 18 | 18 | 57 | 58 | 293 | 75 |
| PN341 | 457 | 131 | 104 | 85 | 74 | 62 | 4E+07 | 4E+07 | 4E+07 | 4E+07 | 4E+07 | 4E+07 | 162 | 2E+06 | 3.7E+07 | 3.7E+07 | 50.34 | 50.13 | 994136 | 953865 | 328424 | 309336 | 877382 | 863928 | 15 | 16 | 44 | 46 | 265 | 69 |
| PN342 | 474 | 122 | 90 | 72 | 64 | 58 | 4E+07 | 4E+07 | 4E+07 | 4E+07 | 4E+07 | 4E+07 | 153 | 2E+06 | 3.7E+07 | 3.7E+07 | 50.38 | 50.13 | 1E+06 | 946057 | 311640 | 298395 | 934923 | 922933 | 15 | 16 | 39 | 41 | 249 | 56 |
| PN343 | 440 | 150 | 120 | 105 | 89 | 76 | 4E+07 | 4E+07 | 4E+07 | 4E+07 | 4E+07 | 4E+07 | 178 | 3E+06 | 3.7E+07 | 3.7E+07 | 50.33 | 50.13 | 688433 | 656831 | 212999 | 205866 | 913614 | 906743 | 17 | 18 | 54 | 55 | 310 | 70 |
| PN344 | 436 | 141 | 106 | 92 | 74 | 64 | 4E+07 | 4E+07 | 4E+07 | 4E+07 | 4E+07 | 4E+07 | 197 | 2E+06 | 3.7E+07 | 3.7E+07 | 50.35 | 50.13 | 1E+06 | 1E+06 | 279585 | 217193 | 945115 | 927834 | 14 | 15 | 42 | 44 | 261 | 67 |
| PN345 | 467 | 176 | 143 | 117 | 97 | 81 | 4E+07 | 4E+07 | 4E+07 | 4E+07 | 4E+07 | 4E+07 | 203 | 2E+06 | 3.7E+07 | 3.7E+07 | 50.33 | 50.13 | 623254 | 623254 | 190021 | 173051 | 720153 | 714886 | 19 | 19 | 59 | 60 | 297 | 78 |
| PN346 | 538 | 200 | 157 | 122 | 105 | 86 | 4E+07 | 4E+07 | 4E+07 | 4E+07 | 4E+07 | 4E+07 | 234 | 2E+06 | 3.7E+07 | 3.7E+07 | 50.33 | 50.13 | 621392 | 621392 | 165811 | 160832 | 704842 | 699602 | 20 | 20 | 64 | 66 | 283 | 88 |
| PN347 | 412 | 112 | 86 | 75 | 61 | 55 | 4E+07 | 4E+07 | 4E+07 | 4E+07 | 4E+07 | 4E+07 | 141 | 2E+06 | 3.7E+07 | 3.7E+07 | 50.53 | 50.13 | 1E+06 | 1E+06 | 275261 | 256725 | 1E+06 | 1E+06 | 14 | 14 | 39 | 41 | 271 | 60 |
| PN348 | 421 | 108 | 82 | 73 | 67 | 61 | 4E+07 | 4E+07 | 4E+07 | 4E+07 | 4E+07 | 4E+07 | 134 | 3E+06 | 3.7E+07 | 3.7E+07 | 50.4 | 50.13 | 1E+06 | 1E+06 | 302164 | 243564 | 1E+06 | 1E+06 | 13 | 13 | 40 | 42 | 280 | 63 |
| PN349 | 447 | 138 | 106 | 90 | 82 | 73 | 4E+07 | 4E+07 | 4E+07 | 4E+07 | 4E+07 | 4E+07 | 170 | 2E+06 | 3.7E+07 | 3.7E+07 | 50.32 | 50.13 | 777804 | 777804 | 221242 | 218686 | 852532 | 840368 | 16 | 16 | 50 | 52 | 255 | 69 |
| PN350 | 408 | 112 | 87 | 76 | 67 | 61 | 4E+07 | 4E+07 | 4E+07 | 4E+07 | 4E+07 | 4E+07 | 144 | 2E+06 | 3.7E+07 | 3.7E+07 | 50.35 | 50.13 | 1E+06 | 1E+06 | 308620 | 254300 | 1E+06 | 988802 | 14 | 14 | 40 | 42 | 268 | 56 |
| PN351 | 597 | 94 | 70 | 59 | 56 | 54 | 4E+07 | 4E+07 | 4E+07 | 4E+07 | 4E+07 | 4E+07 | 131 | 2E+06 | 3.7E+07 | 3.7E+07 | 50.44 | 50.13 | 1E+06 | 970192 | 327236 | 315658 | 1E+06 | 1E+06 | 13 | 14 | 38 | 39 | 306 | 58 |
| PN352 | 374 | 124 | 100 | 86 | 69 | 59 | 4E+07 | 4E+07 | 4E+07 | 4E+07 | 4E+07 | 4E+07 | 159 | 2E+06 | 3.7E+07 | 3.7E+07 | 50.21 | 50.13 | 996198 | 996198 | 271191 | 250858 | 1E+06 | 1E+06 | 13 | 13 | 39 | 40 | 279 | 66 |
| PN353 | 521 | 136 | 108 | 77 | 61 | 55 | 4E+07 | 4E+07 | 4E+07 | 4E+07 | 4E+07 | 4E+07 | 161 | 2E+06 | 3.7E+07 | 3.7E+07 | 50.24 | 50.13 | 1E+06 | 1E+06 | 276715 | 253710 | 1E+06 | 1E+06 | 13 | 13 | 36 | 37 | 308 | 57 |
| PN354 | 353 | 106 | 94 | 84 | 69 | 64 | 4E+07 | 4E+07 | 4E+07 | 4E+07 | 4E+07 | 4E+07 | 125 | 2E+06 | 3.6E+07 | 3.7E+07 | 50.63 | 50.13 | 999530 | 999530 | 258111 | 232502 | 974611 | 948122 | 14 | 14 | 44 | 47 | 273 | 63 |
| PN355 | 425 | 118 | 89 | 78 | 66 | 56 | 4E+07 | 4E+07 | 4E+07 | 4E+07 | 4E+07 | 4E+07 | 153 | 2E+06 | 3.7E+07 | 3.7E+07 | 50.34 | 50.13 | 982563 | 982563 | 332794 | 321320 | 912903 | 896653 | 16 | 16 | 40 | 42 | 258 | 56 |
| PN356 | 463 | 140 | 108 | 93 | 74 | 59 | 4E+07 | 4E+07 | 4E+07 | 4E+07 | 4E+07 | 4E+07 | 178 | 2E+06 | 3.7E+07 | 3.7E+07 | 50.34 | 50.13 | 942150 | 942150 | 274622 | 263258 | 930385 | 913865 | 15 | 15 | 44 | 46 | 258 | 65 |
| PN357 | 471 | 136 | 99 | 82 | 70 | 61 | 4E+07 | 4E+07 | 4E+07 | 4E+07 | 4E+07 | 4E+07 | 184 | 3E+06 | 3.7E+07 | 3.7E+07 | 50.45 | 50.13 | 891052 | 891052 | 282321 | 256991 | 981547 | 962495 | 15 | 15 | 43 | 46 | 272 | 68 |
| PN358 | 1143 | 267 | 163 | 122 | 93 | 75 | 4E+07 | 4E+07 | 4E+07 | 4E+07 | 4E+07 | 4E+07 | 433 | 2E+06 | 3.8E+07 | 3.7E+07 | 50.35 | 50.13 | 643644 | 670395 | 193742 | 210701 | 741681 | 754514 | 19 | 18 | 57 | 55 | 288 | 74 |
| PN359 | 348 | 115 | 88 | 78 | 65 | 58 | 4E+07 | 4E+07 | 4E+07 | 4E+07 | 4E+07 | 4E+07 | 144 | 3E+06 | 3.7E+07 | 3.7E+07 | 50.3 | 50.13 | 1E+06 | 987273 | 309491 | 300822 | 1E+06 | 1E+06 | 13 | 14 | 38 | 39 | 291 | 56 |

| | | | | | | | | | | | | | | | | | | | | | | | | | | | | | | |
|---|---|---|---|---|---|---|---|---|---|---|---|---|---|---|---|---|---|---|---|---|---|---|---|---|---|---|---|---|---|---|
| PN360 | 358 | 100 | 80 | 69 | 62 | 58 | 4E+07 | 4E+07 | 4E+07 | 4E+07 | 4E+07 | 4E+07 | 136 | 2E+06 | 3.7E+07 | 3.7E+07 | 50.44 | 50.13 | 983220 | 983220 | 309070 | 280812 | 1E+06 | 981829 | 14 | 14 | 40 | 42 | 280 | 61 |
| PN361 | 549 | 128 | 102 | 88 | 78 | 66 | 4E+07 | 4E+07 | 4E+07 | 4E+07 | 4E+07 | 4E+07 | 152 | 2E+06 | 3.7E+07 | 3.7E+07 | 50.24 | 50.13 | 892694 | 886330 | 241836 | 222594 | 859351 | 851640 | 16 | 17 | 46 | 47 | 277 | 67 |
| PN362 | 412 | 111 | 85 | 74 | 62 | 56 | 4E+07 | 4E+07 | 4E+07 | 4E+07 | 4E+07 | 4E+07 | 137 | 2E+06 | 3.7E+07 | 3.7E+07 | 50.32 | 50.13 | 1E+06 | 1E+06 | 294887 | 294605 | 1E+06 | 1E+06 | 14 | 14 | 38 | 39 | 300 | 53 |
| PN363 | 583 | 138 | 101 | 88 | 77 | 69 | 4E+07 | 4E+07 | 4E+07 | 4E+07 | 4E+07 | 4E+07 | 180 | 2E+06 | 3.7E+07 | 3.7E+07 | 50.24 | 50.13 | 931751 | 909454 | 264701 | 239125 | 880613 | 872728 | 15 | 16 | 45 | 46 | 270 | 70 |
| PN364 | 639 | 195 | 138 | 113 | 97 | 82 | 4E+07 | 4E+07 | 4E+07 | 4E+07 | 4E+07 | 4E+07 | 250 | 1E+06 | 3.7E+07 | 3.7E+07 | 50.31 | 50.13 | 559002 | 550028 | 188851 | 173178 | 651624 | 643790 | 21 | 22 | 61 | 63 | 264 | 83 |
| PN365 | 475 | 126 | 97 | 85 | 73 | 65 | 4E+07 | 4E+07 | 4E+07 | 4E+07 | 4E+07 | 4E+07 | 162 | 2E+06 | 3.7E+07 | 3.7E+07 | 50.34 | 50.13 | 905862 | 905862 | 275794 | 274621 | 931552 | 915056 | 15 | 15 | 44 | 46 | 251 | 61 |
| PN366 | 391 | 95 | 77 | 70 | 63 | 57 | 4E+07 | 4E+07 | 4E+07 | 4E+07 | 4E+07 | 4E+07 | 121 | 2E+06 | 3.7E+07 | 3.7E+07 | 50.49 | 50.13 | 997422 | 997422 | 339779 | 321692 | 970154 | 949861 | 15 | 15 | 37 | 39 | 299 | 58 |
| PN367 | 471 | 94 | 79 | 66 | 62 | 51 | 4E+07 | 4E+07 | 4E+07 | 4E+07 | 4E+07 | 4E+07 | 121 | 2E+06 | 3.7E+07 | 3.7E+07 | 50.28 | 50.13 | 1E+06 | 1E+06 | 366202 | 366202 | 1E+06 | 1E+06 | 14 | 14 | 37 | 37 | 298 | 55 |
| PN368 | 433 | 125 | 97 | 83 | 70 | 62 | 4E+07 | 4E+07 | 4E+07 | 4E+07 | 4E+07 | 4E+07 | 153 | 2E+06 | 3.7E+07 | 3.7E+07 | 50.34 | 50.13 | 1E+06 | 1E+06 | 307830 | 219872 | 953976 | 937165 | 15 | 15 | 41 | 44 | 256 | 61 |
| PN369 | 567 | 111 | 92 | 81 | 71 | 64 | 4E+07 | 4E+07 | 4E+07 | 4E+07 | 4E+07 | 4E+07 | 163 | 2E+06 | 3.7E+07 | 3.7E+07 | 50.24 | 50.13 | 1E+06 | 1E+06 | 249129 | 249129 | 1E+06 | 1E+06 | 14 | 14 | 45 | 45 | 302 | 65 |
| PN370 | 407 | 106 | 86 | 76 | 59 | 53 | 4E+07 | 4E+07 | 4E+07 | 4E+07 | 4E+07 | 4E+07 | 135 | 2E+06 | 3.7E+07 | 3.7E+07 | 50.34 | 50.13 | 1E+06 | 1E+06 | 327764 | 293081 | 1E+06 | 1E+06 | 13 | 13 | 36 | 38 | 262 | 54 |
| PN371 | 1591 | 372 | 187 | 134 | 89 | 68 | 4E+07 | 4E+07 | 4E+07 | 4E+07 | 4E+07 | 4E+07 | 599 | 3E+06 | 3.8E+07 | 3.7E+07 | 50.38 | 50.13 | 788040 | 921364 | 166305 | 190327 | 906177 | 930193 | 16 | 15 | 54 | 49 | 291 | 73 |
| PN372 | 432 | 163 | 144 | 122 | 100 | 86 | 4E+07 | 4E+07 | 4E+07 | 4E+07 | 4E+07 | 4E+07 | 194 | 3E+06 | 4.2E+07 | 3.7E+07 | 49.08 | 50.13 | 1E+06 | 1E+06 | 169877 | 335908 | 1E+06 | 1E+06 | 14 | 12 | 53 | 36 | 278 | 57 |
| PN373 | 415 | 121 | 99 | 86 | 68 | 58 | 4E+07 | 4E+07 | 4E+07 | 4E+07 | 4E+07 | 4E+07 | 151 | 2E+06 | 3.7E+07 | 3.7E+07 | 50.34 | 50.13 | 1E+06 | 1E+06 | 307821 | 286443 | 900912 | 884894 | 16 | 16 | 41 | 43 | 262 | 60 |
| PN374 | 408 | 122 | 99 | 85 | 70 | 63 | 4E+07 | 4E+07 | 4E+07 | 4E+07 | 4E+07 | 4E+07 | 148 | 3E+06 | 3.7E+07 | 3.7E+07 | 50.34 | 50.13 | 983141 | 983141 | 285065 | 278277 | 1E+06 | 1E+06 | 14 | 14 | 42 | 44 | 245 | 58 |
| PN375 | 377 | 87 | 69 | 61 | 57 | 50 | 4E+07 | 4E+07 | 4E+07 | 4E+07 | 4E+07 | 4E+07 | 116 | 2E+06 | 3.7E+07 | 3.7E+07 | 50.39 | 50.13 | 1E+06 | 1E+06 | 385329 | 347674 | 1E+06 | 1E+06 | 14 | 14 | 34 | 35 | 293 | 51 |
| PN376 | 306 | 84 | 71 | 64 | 58 | 51 | 4E+07 | 4E+07 | 4E+07 | 4E+07 | 4E+07 | 4E+07 | 114 | 2E+06 | 3.7E+07 | 3.7E+07 | 50.21 | 50.13 | 1E+06 | 1E+06 | 340948 | 335427 | 1E+06 | 1E+06 | 13 | 13 | 33 | 34 | 327 | 51 |
| PN377 | 405 | 123 | 99 | 83 | 71 | 62 | 4E+07 | 4E+07 | 4E+07 | 4E+07 | 4E+07 | 4E+07 | 163 | 2E+06 | 3.7E+07 | 3.7E+07 | 50.22 | 50.13 | 932432 | 932432 | 275334 | 240539 | 909271 | 900250 | 15 | 15 | 44 | 45 | 303 | 63 |
| PN378 | 342 | 99 | 74 | 64 | 59 | 53 | 4E+07 | 4E+07 | 4E+07 | 4E+07 | 4E+07 | 4E+07 | 131 | 3E+06 | 3.7E+07 | 3.7E+07 | 50.47 | 50.13 | 1E+06 | 1E+06 | 333319 | 307069 | 1E+06 | 1E+06 | 14 | 14 | 36 | 39 | 284 | 61 |
| PN379 | 487 | 121 | 97 | 86 | 74 | 59 | 4E+07 | 4E+07 | 4E+07 | 4E+07 | 4E+07 | 4E+07 | 158 | 1E+06 | 3.7E+07 | 3.7E+07 | 50.37 | 50.13 | 1E+06 | 1E+06 | 283041 | 240585 | 882280 | 866753 | 16 | 16 | 42 | 44 | 270 | 64 |
| PN380 | 443 | 111 | 90 | 80 | 64 | 58 | 4E+07 | 4E+07 | 4E+07 | 4E+07 | 4E+07 | 4E+07 | 132 | 2E+06 | 3.7E+07 | 3.7E+07 | 50.31 | 50.13 | 877333 | 851968 | 321503 | 282668 | 983457 | 973715 | 14 | 15 | 41 | 43 | 307 | 59 |
| mean | 1949 | 305 | 168 | 130 | 100 | 87 | 4E+07 | 4E+07 | 4E+07 | 4E+07 | 4E+07 | 4E+07 | 446 | 2E+06 | 3.7E+07 | 3.7E+07 | 50.37 | 50.13 | 698005 | 695548 | 195615 | 185106 | 750476 | 744962 | 22 | 22 | 79 | 73 | 247 | 79 |
| STD | 2501 | 244 | 88 | 61 | 41 | 32 | 1E+06 | 911154 | 935917 | 1E+06 | 1E+06 | 1E+06 | 656 | 676673 | 1024689 | 0 | 0.32 | 1E-13 | 298357 | 300025 | 105428 | 104251 | 276382 | 276000 | 11 | 11 | 122 | 43 | 49 | 22 |

**STable 4 (cont.): QC genome assemblies using Quast**

| Assembly | Misassembled contigs length | # local misassemblies | # scaffold gap ext. mis. | # scaffold gap loc. mis. | # un-aligned mis. contigs | # unaligned contigs | Unaligned length | Genome fraction (%) | Duplication ratio | # N's per 100 kbp | # mismatches per 100 kbp | # indels per 100 kbp | # genomic features | Largest alignment | Total aligned length | NA50 | NGA50 | NA90 | NGA90 | auNA | auNGA | LA50 | LGA50 | LA90 | LGA90 |
|---|---|---|---|---|---|---|---|---|---|---|---|---|---|---|---|---|---|---|---|---|---|---|---|---|---|
| DPIRD1 | 26730249 | 212 | 2 | 0 | 18 | 166 + 222 pa | 2350222 | 91.90 | 1.001 | 3.96 | 472.73 | 26.49 | 120581 + 10 | 745400 | 34383588 | 205102 | 204931 | 23055 | 10392 | 231654.1 | 227687.4 | 58 | 59 | 222 | 258 |
| DPIRD10 | 27724687 | 196 | 0 | 1 | 15 | 156 + 221 pa | 2077949 | 92.53 | 1.001 | 4.8 | 454.37 | 25.99 | 121777 + 88 | 1268762 | 34623514 | 238973 | 238479 | 32479 | 12693 | 306818.1 | 301305.3 | 44 | 46 | 173 | 202 |
| DPIRD100 | 30952880 | 203 | 1 | 3 | 14 | 254 + 171 pa | 2364868 | 92.47 | 1 | 3.93 | 471.49 | 26.39 | 121159 + 82 | 799235 | 34588701 | 226748 | 219795 | 27753 | 18929 | 267733.4 | 264677.1 | 50 | 51 | 184 | 201 |
| DPIRD102 | 26021205 | 209 | 2 | 0 | 23 | 135 + 220 pa | 2342973 | 91.94 | 1.001 | 3.71 | 472.63 | 26.47 | 120558 + 10 | 707857 | 34394834 | 204330 | 200718 | 20181 | 10080 | 220412.3 | 216650.8 | 60 | 61 | 233 | 273 |
| DPIRD103 | 25689161 | 193 | 2 | 4 | 17 | 203 + 240 pa | 2668410 | 92.14 | 1.001 | 5.17 | 484.14 | 26.24 | 120951 + 96 | 759425 | 34476903 | 185696 | 185696 | 19867 | 13679 | 224158.7 | 222785.5 | 59 | 59 | 238 | 251 |
| DPIRD104 | 28039180 | 200 | 0 | 1 | 18 | 94 + 207 par | 2773946 | 92.10 | 1.001 | 6.45 | 475.54 | 25.87 | 121086 + 93 | 1278026 | 34461881 | 219205 | 219205 | 18372 | 15392 | 263332.9 | 262332.2 | 53 | 53 | 217 | 224 |
| DPIRD105 | 27301057 | 187 | 2 | 0 | 10 | 58 + 190 par | 1924299 | 92.36 | 1.001 | 4.2 | 466.32 | 26.44 | 120884 + 98 | 705446 | 34559414 | 190943 | 188817 | 29378 | 14912 | 222351.3 | 217039.7 | 60 | 62 | 218 | 254 |
| DPIRD106 | 26759813 | 188 | 1 | 0 | 15 | 150 + 230 pa | 2785372 | 92.72 | 1.001 | 3.96 | 464.62 | 26.4 | 121465 + 10 | 814432 | 34691460 | 182968 | 182968 | 12437 | 14351 | 217076.4 | 217660 | 61 | 61 | 272 | 266 |
| DPIRD107 | 25987461 | 196 | 1 | 2 | 19 | 480 + 234 pa | 2669249 | 92.58 | 1.001 | 5.82 | 463.05 | 25.43 | 121441 + 11 | 804825 | 34644846 | 204983 | 204983 | 14484 | 13299 | 259360.3 | 258938.1 | 53 | 53 | 220 | 224 |
| DPIRD108 | 29061240 | 190 | 1 | 2 | 21 | 100 + 206 pa | 2501491 | 91.90 | 1 | 3.68 | 438.98 | 25 | 120988 + 87 | 779562 | 34374983 | 203719 | 201616 | 25852 | 14385 | 238878.4 | 235654.7 | 55 | 56 | 215 | 237 |
| DPIRD109 | 24829036 | 199 | 1 | 0 | 13 | 96 + 225 par | 2127663 | 92.64 | 1.001 | 4.85 | 453.54 | 25.79 | 121581 + 10 | 606182 | 34663569 | 173082 | 170537 | 26848 | 14791 | 196322.7 | 193235.8 | 67 | 69 | 248 | 275 |
| DPIRD11 | 36694501 | 212 | 2 | 3 | 4 | 11 + 49 part | 2523535 | 92.23 | 1.001 | 4.11 | 476.63 | 27.21 | 120770 + 82 | 839259 | 34529777 | 268502 | 267904 | 34104 | 24245 | 305730.1 | 303068.2 | 43 | 44 | 168 | 178 |
| DPIRD110 | 25378284 | 187 | 1 | 2 | 18 | 122 + 248 pa | 2965614 | 92.91 | 1.001 | 5.86 | 480.2 | 26.61 | 121193 + 10 | 739414 | 34771925 | 190569 | 190569 | 10391 | 14441 | 212786.5 | 214824.5 | 62 | 62 | 277 | 251 |
| DPIRD111 | 25532553 | 201 | 1 | 0 | 19 | 74 + 192 par | 2145890 | 92.66 | 1 | 3.89 | 450.97 | 25.86 | 121563 + 10 | 864252 | 34657224 | 199015 | 192772 | 29483 | 15155 | 243872 | 240117.9 | 55 | 57 | 208 | 232 |
| DPIRD112 | 27656906 | 191 | 1 | 3 | 15 | 163 + 211 pa | 2660024 | 92.12 | 1.001 | 3.97 | 482.68 | 26.33 | 120991 + 93 | 975020 | 34464318 | 191486 | 189965 | 20569 | 14791 | 248777.2 | 247098.4 | 56 | 57 | 230 | 243 |
| DPIRD12 | 24648320 | 214 | 1 | 0 | 18 | 151 + 232 pa | 2343781 | 91.95 | 1.001 | 3.96 | 473.85 | 26.71 | 120466 + 11 | 738471 | 34410511 | 188071 | 183462 | 20848 | 8366 | 225582.6 | 221865.7 | 58 | 60 | 244 | 284 |
| DPIRD13 | 24509834 | 182 | 0 | 1 | 8 | 217 + 147 pa | 1203931 | 90.35 | 1 | 5.85 | 464.14 | 26.13 | 121228 + 81 | 739207 | 33783656 | 258911 | 246045 | 54705 | 1928 | 286323.5 | 268120.5 | 44 | 49 | 147 | 251 |
| DPIRD14 | 23722504 | 206 | 1 | 0 | 15 | 120 + 232 pa | 2105320 | 92.63 | 1.001 | 4.11 | 453.32 | 26.07 | 121602 + 10 | 864358 | 34660646 | 196593 | 192672 | 28743 | 14466 | 243798 | 239820.1 | 55 | 57 | 219 | 246 |
| DPIRD15 | 10052023 | 173 | 5 | 94 | 32 | 794 + 457 pa | 3066193 | 93.21 | 1.011 | 252.98 | 449.82 | 26.98 | 120686 + 20 | 326922 | 35235223 | 79174 | 82362 | 1788 | 6792 | 94175.4 | 96627.1 | 144 | 138 | 864 | 610 |
| DPIRD16 | 36144638 | 182 | 0 | 3 | 6 | 23 + 62 part | 2369792 | 92.80 | 1.002 | 3.54 | 460.81 | 26.55 | 121865 + 73 | 841552 | 34754578 | 299516 | 299516 | 34926 | 30549 | 324595.6 | 322344.2 | 40 | 40 | 158 | 165 |
| DPIRD17 | 24530226 | 187 | 2 | 0 | 15 | 113 + 188 pa | 2017993 | 92.45 | 1.001 | 3.82 | 483.09 | 26.62 | 121058 + 10 | 687799 | 34592771 | 183619 | 174825 | 27558 | 13336 | 214278.6 | 209883.1 | 62 | 64 | 234 | 269 |
| DPIRD18 | 28071862 | 208 | 1 | 2 | 13 | 70 + 156 par | 2317552 | 92.12 | 1 | 3.2 | 473.28 | 26.78 | 120761 + 88 | 705293 | 34452397 | 253453 | 253149 | 30549 | 14537 | 265298.4 | 260960.9 | 48 | 49 | 179 | 203 |
| DPIRD19 | 25361597 | 201 | 1 | 1 | 19 | 189 + 246 pa | 2720636 | 92.12 | 1.002 | 4.18 | 463.92 | 25.3 | 121265 + 10 | 807868 | 34499586 | 202735 | 201914 | 15839 | 13336 | 246063.2 | 245088.8 | 53 | 54 | 234 | 243 |
| DPIRD2 | 30478452 | 207 | 1 | 0 | 20 | 266 + 169 pa | 2476596 | 92.15 | 1.001 | 5.11 | 465.11 | 26.1 | 121071 + 81 | 747918 | 34496845 | 246973 | 246864 | 28204 | 18191 | 275433.2 | 272428.4 | 47 | 48 | 183 | 199 |
| DPIRD20 | 24917862 | 169 | 1 | 0 | 12 | 308 + 162 pa | 1464743 | 90.97 | 1 | 5.31 | 459.76 | 25.51 | 121299 + 71 | 990276 | 34026786 | 270416 | 247418 | 50449 | 5784 | 296272.3 | 281509 | 43 | 47 | 153 | 221 |
| DPIRD21 | 29408230 | 190 | 2 | 0 | 16 | 73 + 168 par | 1909304 | 92.33 | 1.001 | 4.2 | 465.3 | 26.5 | 121078 + 75 | 747886 | 34540803 | 246436 | 242802 | 42638 | 18929 | 291753.2 | 284489 | 45 | 47 | 163 | 193 |

| | | | | | | | | | | | | | | | | | | | | | | | | | |
|---|---|---|---|---|---|---|---|---|---|---|---|---|---|---|---|---|---|---|---|---|---|---|---|---|---|
| DPIRD22 | 28104988 | 181 | 4 | 2 | 10 | 149 + 207 pa | 2501783 | 92.35 | 1.001 | 5.06 | 469.24 | 25.8 | 121572 + 79 | 977510 | 34560366 | 239882 | 239882 | 25489 | 14956 | 287095.4 | 284672.7 | 47 | 47 | 187 | 200 |
| DPIRD23 | 23458649 | 163 | 1 | 0 | 18 | 259 + 206 pa | 1833531 | 91.30 | 1.001 | 5.48 | 456.92 | 25.23 | 120410 + 82 | 800171 | 34159164 | 245424 | 243141 | 34395 | 3137 | 271446.8 | 261531.6 | 48 | 50 | 174 | 278 |
| DPIRD24 | 27515422 | 201 | 0 | 2 | 21 | 175 + 211 pa | 2382223 | 92.42 | 1.001 | 3.91 | 458.18 | 25.21 | 121175 + 93 | 779509 | 34595150 | 238186 | 236851 | 19515 | 12154 | 261624.6 | 258859.2 | 49 | 50 | 198 | 220 |
| DPIRD25 | 29941146 | 191 | 0 | 0 | 15 | 189 + 194 pa | 2394584 | 92.13 | 1.001 | 3.81 | 466.27 | 26.41 | 121336 + 86 | 753270 | 34474083 | 272369 | 263975 | 28764 | 13406 | 274284.8 | 270543.5 | 48 | 49 | 181 | 203 |
| DPIRD27 | 24560642 | 209 | 2 | 1 | 20 | 152 + 220 pa | 2365465 | 91.88 | 1.001 | 4.02 | 471.33 | 26.46 | 120415 + 11 | 715222 | 34382282 | 178547 | 177147 | 20848 | 9958 | 220108.4 | 216438 | 62 | 63 | 247 | 286 |
| DPIRD28 | 24308090 | 195 | 2 | 1 | 17 | 132 + 229 pa | 2120972 | 92.58 | 1.001 | 5.3 | 453.19 | 25.74 | 121574 + 10 | 788745 | 34638139 | 189159 | 185818 | 24376 | 14545 | 225474.3 | 221753.1 | 59 | 61 | 219 | 249 |
| DPIRD29 | 31351511 | 178 | 1 | 0 | 19 | 81 + 196 par | 2418357 | 91.79 | 1 | 2.22 | 486.58 | 27.08 | 120770 + 76 | 738882 | 34333035 | 247733 | 245399 | 26927 | 11838 | 258496.5 | 254131.5 | 50 | 51 | 187 | 218 |
| DPIRD3 | 29413351 | 203 | 1 | 2 | 20 | 236 + 225 pa | 2653922 | 92.11 | 1.001 | 7.14 | 477.59 | 26.52 | 121409 + 88 | 680798 | 34458886 | 258958 | 250383 | 19021 | 13177 | 285043.4 | 283035.2 | 45 | 46 | 193 | 207 |
| DPIRD30 | 36585464 | 212 | 2 | 0 | 4 | 10 + 49 part | 2515106 | 92.23 | 1.001 | 3.17 | 477.53 | 27.31 | 120776 + 80 | 839266 | 34511221 | 267904 | 263692 | 34104 | 23057 | 305324.5 | 302558.2 | 43 | 44 | 167 | 178 |
| DPIRD31 | 28213931 | 188 | 2 | 0 | 18 | 153 + 198 pa | 1813541 | 92.20 | 1.001 | 4.68 | 471.26 | 26.85 | 121058 + 79 | 747897 | 34524027 | 244493 | 240351 | 37618 | 11243 | 276182 | 268539.2 | 47 | 49 | 170 | 209 |
| DPIRD32 | 25788547 | 206 | 1 | 4 | 24 | 253 + 236 pa | 2041266 | 91.98 | 1.002 | 5.58 | 464.8 | 25.22 | 121089 + 90 | 744166 | 34449070 | 235468 | 233953 | 30549 | 8392 | 276464.9 | 269959.9 | 46 | 48 | 177 | 223 |
| DPIRD33 | 30742335 | 187 | 1 | 2 | 26 | 112 + 196 pa | 2505901 | 91.92 | 1.001 | 4.63 | 441.72 | 24.92 | 121120 + 76 | 761590 | 34388658 | 246997 | 246340 | 27921 | 14367 | 285337.2 | 281638.5 | 45 | 46 | 187 | 207 |
| DPIRD34 | 36096216 | 185 | 0 | 0 | 10 | 45 + 76 part | 3174573 | 92.37 | 1.001 | 3.08 | 464.21 | 25.22 | 121526 + 73 | 789592 | 34579865 | 291777 | 292148 | 14035 | 27837 | 299096.1 | 302058.7 | 44 | 43 | 186 | 170 |
| DPIRD35 | 26600145 | 179 | 1 | 1 | 25 | 141 + 258 pa | 2553000 | 92.24 | 1.001 | 5.23 | 479.36 | 26.69 | 120861 + 98 | 876350 | 34508564 | 185773 | 184184 | 16218 | 11251 | 217845.5 | 216008.9 | 63 | 64 | 256 | 277 |
| DPIRD36 | 24204044 | 205 | 1 | 1 | 16 | 164 + 239 pa | 2735109 | 91.93 | 1.001 | 4.25 | 477.66 | 26.37 | 120846 + 11 | 485515 | 34388167 | 202176 | 200943 | 14912 | 11838 | 206961.4 | 205630.7 | 62 | 63 | 254 | 270 |
| DPIRD37 | 28277479 | 195 | 2 | 1 | 19 | 132 + 214 pa | 2182187 | 92.26 | 1.001 | 5.69 | 465.06 | 25.62 | 121270 + 83 | 804845 | 34518264 | 231566 | 224800 | 31379 | 13555 | 278804.9 | 273745 | 46 | 47 | 186 | 215 |
| DPIRD38 | 29310345 | 191 | 2 | 0 | 14 | 73 + 182 par | 1908050 | 92.32 | 1.001 | 4.41 | 467.36 | 26.54 | 120980 + 81 | 748138 | 34550822 | 234424 | 227436 | 37036 | 18456 | 276197.2 | 269427.6 | 47 | 49 | 174 | 204 |
| DPIRD39 | 26039530 | 169 | 1 | 1 | 7 | 292 + 133 pa | 1230200 | 90.27 | 1 | 5.32 | 461.67 | 25.77 | 121066 + 77 | 980099 | 33758552 | 242294 | 234634 | 54577 | 1492 | 293777.8 | 275093.5 | 44 | 48 | 152 | 267 |
| DPIRD4 | 24305439 | 199 | 1 | 0 | 16 | 114 + 219 pa | 2118342 | 92.66 | 1.001 | 4.41 | 453.16 | 26.01 | 121604 + 96 | 788427 | 34666277 | 199026 | 198038 | 28919 | 15592 | 234811.5 | 231086.7 | 57 | 59 | 213 | 238 |
| DPIRD40 | 914744 | 10 | 0 | 1 | 2 | 34 + 31 part | 288629 | 98.15 | 1.001 | 4.12 | 18.21 | 5.66 | 126707 + 39 | 1231823 | 36745604 | 524870 | 502570 | 103618 | 93246 | 534458.7 | 529909.2 | 25 | 26 | 79 | 82 |
| DPIRD41 | 24456583 | 190 | 1 | 2 | 18 | 142 + 224 pa | 2360636 | 92.68 | 1.001 | 4.66 | 455.47 | 25.18 | 121357 + 10 | 643330 | 34689687 | 206026 | 203661 | 27778 | 16692 | 218288 | 216378.7 | 60 | 61 | 224 | 237 |
| DPIRD42 | 23030971 | 208 | 1 | 0 | 17 | 100 + 207 pa | 2393753 | 91.97 | 1.001 | 3.29 | 471.94 | 26.35 | 120342 + 12 | 624668 | 34416388 | 161817 | 161439 | 18455 | 11033 | 197519.5 | 194531.1 | 66 | 67 | 278 | 315 |
| DPIRD43 | 23229493 | 192 | 2 | 0 | 8 | 412 + 177 pa | 1676076 | 90.78 | 1 | 3.3 | 477.66 | 26.24 | 121125 + 83 | 881617 | 33950151 | 244630 | 234725 | 48388 | 5052 | 274719.2 | 261918.2 | 47 | 51 | 167 | 242 |
| DPIRD44 | 29362268 | 199 | 1 | 0 | 13 | 104 + 202 pa | 2169473 | 92.30 | 1.001 | 3.17 | 464.74 | 25.52 | 121359 + 79 | 877412 | 34532134 | 238605 | 237328 | 32039 | 16401 | 302012.2 | 296519.3 | 44 | 45 | 168 | 194 |
| DPIRD45 | 23611077 | 180 | 2 | 1 | 4 | 468 + 169 pa | 1769587 | 90.86 | 1.001 | 5.58 | 463.16 | 26.46 | 120779 + 85 | 747982 | 33993916 | 240195 | 234457 | 44841 | 3926 | 272839.9 | 261166.2 | 47 | 51 | 170 | 253 |
| DPIRD46 | 1394472 | 11 | 0 | 3 | 1 | 45 + 28 part | 306893 | 98.46 | 1.001 | 5.72 | 12.84 | 5.85 | 126641 + 64 | 880926 | 36847350 | 300436 | 293630 | 62652 | 59336 | 330102.8 | 328148.8 | 40 | 41 | 136 | 140 |
| DPIRD47 | 25867836 | 173 | 1 | 2 | 33 | 312 + 274 pa | 2206843 | 91.98 | 1.001 | 6.12 | 476.1 | 26.47 | 121348 + 79 | 979855 | 34433942 | 241367 | 238950 | 26399 | 9576 | 275278.3 | 269907 | 48 | 49 | 184 | 226 |
| DPIRD48 | 23953707 | 166 | 0 | 1 | 12 | 302 + 173 pa | 1582557 | 90.94 | 1.001 | 4.26 | 455.75 | 25.38 | 120842 + 79 | 1266510 | 34030235 | 247576 | 238986 | 46066 | 4209 | 303078.6 | 288852.7 | 44 | 48 | 163 | 245 |
| DPIRD49 | 27670447 | 209 | 1 | 0 | 21 | 321 + 235 pa | 2536293 | 91.76 | 1.001 | 3.89 | 475.64 | 26.71 | 120580 + 95 | 745324 | 34341151 | 242169 | 239071 | 18332 | 6724 | 264278.9 | 260811.1 | 49 | 50 | 208 | 250 |
| DPIRD5 | 25232628 | 198 | 0 | 0 | 17 | 117 + 203 pa | 2110630 | 92.64 | 1.001 | 3.43 | 452.08 | 25.89 | 121629 + 99 | 864390 | 34654437 | 197717 | 196679 | 29483 | 15397 | 238330.3 | 234444.1 | 57 | 59 | 210 | 235 |
| DPIRD50 | 26440278 | 190 | 1 | 0 | 17 | 126 + 242 pa | 2069051 | 92.56 | 1.001 | 2.79 | 463.29 | 25.98 | 121293 + 11 | 746999 | 34646506 | 191403 | 182602 | 25851 | 11842 | 226955.2 | 222954.4 | 57 | 59 | 229 | 261 |
| DPIRD51 | 29468558 | 183 | 0 | 0 | 23 | 142 + 213 pa | 3035786 | 92.21 | 1.001 | 3.54 | 465.33 | 25.09 | 121317 + 93 | 657665 | 34506119 | 252916 | 252916 | 10766 | 13929 | 248430.5 | 249501.7 | 51 | 51 | 237 | 226 |

| | | | | | | | | | | | | | | | | | | | | | | | | | |
|---|---|---|---|---|---|---|---|---|---|---|---|---|---|---|---|---|---|---|---|---|---|---|---|---|---|
| DPIRD55 | 22646761 | 198 | 2 | 2 | 22 | 276 + 292 pa | 3002227 | 92.46 | 1.001 | 4.06 | 459.83 | 25.2 | 120974 + 10 | 744459 | 34605797 | 193780 | 195684 | 9576 | 14386 | 216785.9 | 218155.2 | 62 | 61 | 268 | 250 |
| DPIRD56 | 27499453 | 187 | 1 | 3 | 27 | 371 + 287 pa | 3839241 | 92.63 | 1.002 | 11.85 | 496.43 | 26.79 | 121398 + 10 | 816458 | 34700291 | 181654 | 185573 | 84 | 13548 | 212980.6 | 219614.9 | 64 | 61 | 764 | 267 |
| DPIRD57 | 23825749 | 188 | 1 | 2 | 31 | 492 + 283 pa | 3705207 | 92.36 | 1.002 | 11.89 | 472.94 | 26.03 | 121178 + 11 | 753659 | 34589624 | 161296 | 165709 | 965 | 10294 | 201048.5 | 206009.4 | 68 | 65 | 529 | 284 |
| DPIRD58 | 21961733 | 196 | 3 | 1 | 22 | 369 + 271 pa | 3640316 | 91.94 | 1.001 | 10.61 | 472.58 | 26.48 | 120906 + 13 | 716487 | 34410599 | 145679 | 153840 | 1412 | 10713 | 181589.4 | 184903.8 | 76 | 73 | 459 | 312 |
| DPIRD6 | 26205783 | 200 | 0 | 0 | 21 | 172 + 224 pa | 2055504 | 92.43 | 1.001 | 3.22 | 455.46 | 25.99 | 121741 + 85 | 1268873 | 34596190 | 248131 | 244510 | 36306 | 14791 | 307869.8 | 301917.3 | 43 | 45 | 167 | 194 |
| DPIRD60 | 22236712 | 171 | 2 | 2 | 24 | 866 + 308 pa | 4197107 | 92.67 | 1.002 | 14.94 | 472.51 | 26.41 | 121629 + 12 | 589934 | 34699537 | 175052 | 181020 | - | 10294 | 193080.1 | 200987.9 | 69 | 65 | - | 273 |
| DPIRD62 | 29419371 | 187 | 3 | 3 | 23 | 100 + 207 pa | 2156889 | 91.66 | 1.001 | 4.9 | 463.19 | 26.74 | 121131 + 76 | 1021488 | 34291906 | 264525 | 262132 | 37744 | 13701 | 312738 | 304947.7 | 42 | 43 | 163 | 198 |
| DPIRD63 | 19621369 | 198 | 4 | 3 | 21 | 943 + 353 pa | 4169711 | 92.85 | 1.002 | 19.09 | 471.59 | 26.11 | 121284 + 14 | 705864 | 34788448 | 143921 | 151357 | - | 9937 | 184010.3 | 191925.7 | 75 | 70 | - | 314 |
| DPIRD64 | 27337957 | 170 | 2 | 0 | 19 | 198 + 252 pa | 2289001 | 92.08 | 1.001 | 3.34 | 481.23 | 26.89 | 121339 + 94 | 674586 | 34471750 | 199074 | 195450 | 25350 | 13336 | 223586 | 219924.1 | 59 | 61 | 212 | 242 |
| DPIRD66 | 31134428 | 186 | 1 | 1 | 17 | 105 + 182 pa | 1999837 | 92.10 | 1 | 3.3 | 455.72 | 25.45 | 120999 + 81 | 882678 | 34449667 | 270801 | 266599 | 37235 | 13781 | 294128.7 | 286834.8 | 44 | 46 | 162 | 194 |
| DPIRD68 | 29351085 | 185 | 1 | 0 | 20 | 78 + 197 par | 2228399 | 92.30 | 1.001 | 3.79 | 456.05 | 25.99 | 121264 + 81 | 671943 | 34534245 | 256858 | 247799 | 34811 | 15357 | 267373 | 262970.4 | 48 | 49 | 173 | 196 |
| DPIRD69 | 28846581 | 188 | 1 | 0 | 23 | 102 + 199 pa | 2249916 | 92.32 | 1.001 | 3.49 | 457.97 | 26.02 | 121230 + 81 | 672028 | 34545156 | 251564 | 247800 | 33755 | 15459 | 269206.9 | 265000.8 | 48 | 49 | 171 | 194 |
| DPIRD7 | 24802581 | 202 | 1 | 0 | 13 | 76 + 207 par | 2151318 | 92.72 | 1.001 | 4.24 | 453.51 | 26.08 | 121614 + 10 | 606274 | 34687853 | 186642 | 185736 | 25240 | 14791 | 204979.2 | 202039.1 | 64 | 66 | 231 | 256 |
| DPIRD70 | 27118186 | 194 | 1 | 0 | 20 | 162 + 218 pa | 2165827 | 92.21 | 1.001 | 3.45 | 457.96 | 25.96 | 121183 + 90 | 646169 | 34509436 | 246862 | 239130 | 33137 | 13071 | 253394.2 | 248660.1 | 51 | 52 | 186 | 216 |
| DPIRD71 | 24421869 | 189 | 1 | 1 | 17 | 102 + 212 pa | 2260658 | 92.28 | 1.001 | 3.28 | 455.55 | 26.02 | 121000 + 10 | 601853 | 34527651 | 192737 | 187216 | 29348 | 13408 | 201329.9 | 198146.3 | 65 | 67 | 227 | 254 |
| DPIRD72 | 27059003 | 190 | 3 | 0 | 14 | 74 + 189 par | 1871643 | 92.36 | 1.001 | 4.08 | 470.11 | 26.7 | 120905 + 92 | 748137 | 34566691 | 208415 | 205272 | 34284 | 15884 | 245638.6 | 239489.4 | 54 | 56 | 200 | 236 |
| DPIRD73 | 30845650 | 187 | 1 | 3 | 22 | 137 + 205 pa | 2445767 | 91.91 | 1.001 | 4.39 | 440.96 | 24.99 | 121135 + 75 | 779550 | 34386977 | 251820 | 246768 | 29486 | 14389 | 279565.9 | 275485.4 | 46 | 47 | 185 | 206 |
| DPIRD74 | 36838469 | 197 | 0 | 6 | 6 | 12 + 45 part | 2618658 | 92.02 | 1.001 | 2.7 | 442.65 | 25.26 | 121198 + 66 | 779638 | 34447381 | 302423 | 299991 | 33940 | 26897 | 311570.8 | 308980.3 | 41 | 42 | 162 | 172 |
| DPIRD75 | 29473487 | 189 | 1 | 5 | 19 | 112 + 202 pa | 2498784 | 91.90 | 1 | 3.9 | 439.36 | 24.89 | 121102 + 77 | 779534 | 34377144 | 246338 | 244289 | 28154 | 14385 | 275314 | 271596.5 | 47 | 48 | 192 | 212 |
| DPIRD76 | 27653483 | 174 | 1 | 0 | 11 | 110 + 185 pa | 2180448 | 92.53 | 1.001 | 3.7 | 469.24 | 26.36 | 121131 + 94 | 761233 | 34627257 | 184574 | 179421 | 27622 | 14860 | 231117 | 227584.6 | 58 | 60 | 225 | 250 |
| DPIRD77 | 29725876 | 173 | 1 | 1 | 18 | 115 + 171 pa | 2299821 | 92.53 | 1.001 | 2.91 | 467.48 | 26.36 | 121237 + 83 | 788934 | 34617664 | 239333 | 234671 | 29587 | 18207 | 279305.7 | 275838.1 | 46 | 47 | 191 | 208 |
| DPIRD79 | 25118759 | 172 | 2 | 0 | 10 | 442 + 208 pa | 2544583 | 92.54 | 1.001 | 5.41 | 468.77 | 26.3 | 121051 + 10 | 583064 | 34620478 | 169468 | 168979 | 19887 | 14992 | 201828.3 | 200700.6 | 66 | 67 | 255 | 266 |
| DPIRD8 | 25423830 | 202 | 1 | 0 | 15 | 73 + 203 par | 2148534 | 92.67 | 1.001 | 4.35 | 451.74 | 25.66 | 121615 + 10 | 864510 | 34669923 | 190802 | 185737 | 28330 | 15065 | 228698.8 | 225274 | 61 | 63 | 218 | 243 |
| DPIRD82 | 24270365 | 160 | 3 | 13 | 11 | 713 + 265 pa | 3787330 | 93.16 | 1.003 | 19.61 | 468.13 | 25.97 | 122206 + 10 | 661872 | 34918901 | 236807 | 242235 | 539 | 13408 | 251956.1 | 261041.1 | 52 | 49 | 729 | 229 |
| DPIRD83 | 24099777 | 172 | 1 | 11 | 24 | 901 + 320 pa | 3454586 | 92.72 | 1.003 | 68.02 | 448.63 | 25.87 | 121719 + 11 | 706030 | 34771629 | 181063 | 186445 | 1419 | 11009 | 222759.3 | 227941.2 | 63 | 60 | 493 | 262 |
| DPIRD84 | 28852957 | 192 | 1 | 3 | 18 | 191 + 198 pa | 2113333 | 92.11 | 1.001 | 4.83 | 459.58 | 25.84 | 121477 + 98 | 643628 | 34454371 | 238371 | 233353 | 34988 | 13690 | 258048.8 | 252490.4 | 50 | 51 | 193 | 224 |
| DPIRD85 | 26956414 | 190 | 0 | 4 | 13 | 117 + 190 pa | 2275883 | 92.36 | 1.001 | 4.64 | 469.95 | 26.3 | 121227 + 83 | 771223 | 34551245 | 202756 | 201352 | 28839 | 16757 | 241610.2 | 238044.1 | 54 | 56 | 207 | 227 |
| DPIRD86 | 35532870 | 189 | 1 | 4 | 6 | 45 + 60 part | 2370258 | 92.64 | 1.001 | 3.81 | 478.38 | 27.02 | 121416 + 67 | 1279055 | 34685616 | 287772 | 287772 | 40167 | 34149 | 355701 | 352639.6 | 39 | 39 | 158 | 165 |
| DPIRD87 | 27633860 | 187 | 1 | 2 | 15 | 129 + 192 pa | 2288365 | 92.25 | 1.001 | 4.94 | 467.42 | 26.21 | 121067 + 86 | 738819 | 34518546 | 203780 | 201890 | 28389 | 14187 | 248426 | 244646 | 54 | 55 | 213 | 238 |
| DPIRD88 | 27214623 | 175 | 2 | 0 | 17 | 262 + 206 pa | 2428073 | 92.15 | 1.001 | 5.72 | 483.53 | 27.63 | 120975 + 85 | 705735 | 34493328 | 205544 | 198987 | 27558 | 14640 | 227770.8 | 224996.4 | 58 | 59 | 222 | 242 |
| DPIRD89 | 26992369 | 159 | 2 | 0 | 12 | 397 + 209 pa | 2498562 | 92.47 | 1.001 | 6.7 | 469.14 | 26.17 | 121723 + 77 | 666776 | 34610293 | 233723 | 233555 | 18191 | 12259 | 257867.2 | 256044.5 | 51 | 52 | 200 | 216 |
| DPIRD9 | 28081648 | 198 | 1 | 0 | 15 | 86 + 195 par | 2134506 | 92.64 | 1.001 | 4.11 | 451.64 | 25.99 | 121746 + 90 | 864509 | 34658410 | 229226 | 215165 | 31918 | 18527 | 266916.5 | 262727.4 | 49 | 51 | 187 | 208 |

| | | | | | | | | | | | | | | | | | | | | | | | | | |
|---|---|---|---|---|---|---|---|---|---|---|---|---|---|---|---|---|---|---|---|---|---|---|---|---|---|
| DPIRD90 | 26139577 | 173 | 1 | 1 | 20 | 109 + 246 pa | 2251199 | 92.31 | 1 | 3.39 | 451.73 | 25.7 | 121131 + 83 | 738942 | 34527157 | 199001 | 197732 | 28192 | 12943 | 231795.7 | 228211.1 | 57 | 58 | 214 | 243 |
| DPIRD91 | 30146986 | 184 | 1 | 3 | 12 | 149 + 187 pa | 2207785 | 92.06 | 1.001 | 4.04 | 459.2 | 25.98 | 121406 + 87 | 642758 | 34445436 | 233226 | 223702 | 33182 | 17617 | 246172.7 | 241389.8 | 53 | 55 | 194 | 220 |
| DPIRD92 | 27298114 | 192 | 0 | 1 | 18 | 78 + 180 par | 2336410 | 91.88 | 1.001 | 2.55 | 464.48 | 26.5 | 120651 + 90 | 739039 | 34377666 | 186552 | 180438 | 28549 | 12593 | 222501.4 | 218568.1 | 60 | 62 | 216 | 247 |
| DPIRD93 | 25799306 | 195 | 0 | 0 | 10 | 108 + 205 pa | 2037701 | 92.42 | 1.001 | 5.12 | 472.97 | 26.62 | 120931 + 10 | 759031 | 34585076 | 185695 | 180791 | 27824 | 11658 | 222659 | 218159.8 | 60 | 62 | 230 | 269 |
| DPIRD94 | 23338004 | 162 | 1 | 0 | 14 | 283 + 225 pa | 2601596 | 92.59 | 1.001 | 4.79 | 466.3 | 25.72 | 121510 + 10 | 557867 | 34647930 | 144987 | 144235 | 14936 | 14379 | 171495.1 | 170912.2 | 76 | 77 | 306 | 313 |
| DPIRD96 | 29577165 | 191 | 3 | 1 | 21 | 74 + 199 par | 2252206 | 92.00 | 1.001 | 4.51 | 470.77 | 25.96 | 121338 + 82 | 1104043 | 34420411 | 277790 | 267355 | 32883 | 13495 | 312877.3 | 306987.5 | 43 | 44 | 164 | 193 |
| DPIRD97 | 29329750 | 190 | 3 | 1 | 22 | 111 + 203 pa | 2247927 | 91.94 | 1.001 | 4.38 | 469.55 | 25.89 | 121303 + 85 | 1104002 | 34401055 | 239080 | 238804 | 29520 | 13408 | 291329.6 | 285693.5 | 46 | 47 | 173 | 205 |
| PN260 | 34878596 | 183 | 1 | 7 | 3 | 9 + 77 part | 1955382 | 92.81 | 1.001 | 3.56 | 488.38 | 28.59 | 121762 + 69 | 1246880 | 34726108 | 271407 | 260332 | 45392 | 30659 | 333158.1 | 326998.1 | 41 | 42 | 162 | 179 |
| PN261 | 35758475 | 189 | 0 | 3 | 11 | 10 + 89 part | 2104339 | 92.62 | 1.002 | 2.74 | 488.38 | 27.44 | 121738 + 68 | 808581 | 34684997 | 315931 | 312253 | 47848 | 26938 | 324882 | 319714.1 | 39 | 40 | 152 | 167 |
| PN262 | 34707421 | 208 | 0 | 4 | 10 | 8227 + 73 pa | 9723334 | 91.81 | 1.001 | 17.22 | 508.96 | 28.19 | 120994 + 83 | 870656 | 34357622 | 231809 | 275058 | - | 22814 | 244455.6 | 288276.7 | 60 | 47 | - | 178 |
| PN263 | 34889126 | 163 | 1 | 8 | 12 | 222 + 100 pa | 2704699 | 92.83 | 1.001 | 3.16 | 473.64 | 26.52 | 122053 + 63 | 1020342 | 34737317 | 257998 | 257998 | 28748 | 30345 | 296601.3 | 297115.3 | 45 | 45 | 173 | 171 |
| PN264 | 35030639 | 208 | 1 | 3 | 7 | 8 + 67 part | 2083459 | 92.62 | 1.001 | 2.92 | 473.6 | 27.11 | 121402 + 79 | 857082 | 34647842 | 286795 | 286410 | 46789 | 29587 | 324740.4 | 319220.8 | 41 | 42 | 150 | 165 |
| PN265 | 35053393 | 220 | 1 | 8 | 18 | 25 + 104 par | 2868564 | 91.38 | 1.001 | 2.53 | 546.04 | 30.07 | 120836 + 88 | 828771 | 34209403 | 250974 | 250974 | 20844 | 10564 | 298048.1 | 295593.1 | 44 | 44 | 193 | 212 |
| PN266 | 34953952 | 178 | 0 | 4 | 11 | 6 + 74 part | 1968037 | 92.84 | 1.002 | 3.5 | 467.39 | 26.69 | 121095 + 69 | 726121 | 34777286 | 281418 | 274025 | 46194 | 28204 | 299336.2 | 294214 | 43 | 44 | 156 | 172 |
| PN268 | 34357627 | 199 | 0 | 10 | 12 | 21 + 92 part | 2475658 | 92.49 | 1.001 | 3.21 | 474.85 | 27.38 | 121775 + 79 | 818757 | 34624587 | 244039 | 242340 | 29587 | 25531 | 293676.1 | 291444.6 | 44 | 45 | 175 | 184 |
| PN269 | 35008254 | 196 | 1 | 5 | 4 | 67 + 65 part | 7920737 | 92.09 | 1.001 | 4.09 | 465.89 | 26.39 | 121352 + 78 | 1022213 | 34479714 | 240491 | 289295 | - | 22427 | 264375.1 | 299837.7 | 54 | 45 | - | 172 |
| PN270 | 35146550 | 230 | 1 | 3 | 8 | 17 + 79 part | 2338742 | 91.87 | 1.001 | 3.38 | 542.19 | 30.53 | 120623 + 84 | 1264140 | 34395601 | 261346 | 255094 | 31925 | 15564 | 324413.5 | 318798.2 | 41 | 42 | 178 | 202 |
| PN271 | 34009294 | 206 | 0 | 7 | 6 | 7 + 72 part | 2404141 | 92.47 | 1.001 | 3.47 | 458.22 | 26.41 | 121417 + 77 | 716401 | 34609402 | 268365 | 261493 | 37628 | 28735 | 287858.6 | 285028.1 | 45 | 46 | 163 | 173 |
| PN274 | 35688220 | 233 | 1 | 8 | 8 | 197 + 76 par | 2577806 | 91.86 | 1.001 | 2.58 | 539.51 | 30.42 | 120630 + 84 | 1263819 | 34380460 | 247951 | 244001 | 28734 | 18929 | 319255.1 | 315646.6 | 42 | 43 | 183 | 199 |
| PN275 | 33388483 | 197 | 0 | 4 | 10 | 19 + 90 part | 2328726 | 92.66 | 1.002 | 4.09 | 474.45 | 26.33 | 121454 + 72 | 730362 | 34709513 | 294765 | 294446 | 45758 | 27821 | 292071.5 | 289472.5 | 44 | 45 | 158 | 168 |
| PN276 | 36153327 | 200 | 0 | 1 | 4 | 214 + 51 par | 2711103 | 92.15 | 1.001 | 2.82 | 469.81 | 27.09 | 120952 + 88 | 910840 | 34482609 | 248358 | 248358 | 28549 | 24515 | 284302.3 | 282946.2 | 46 | 46 | 178 | 184 |
| PN277 | 34937714 | 193 | 0 | 4 | 9 | 20 + 68 part | 2460037 | 92.51 | 1.002 | 3.1 | 479.79 | 26.85 | 121531 + 73 | 744322 | 34653408 | 302473 | 302473 | 32735 | 28736 | 310938.4 | 308650.5 | 42 | 42 | 163 | 171 |
| PN279 | 35272411 | 205 | 1 | 3 | 11 | 22 + 75 part | 2390479 | 91.97 | 1.001 | 4.55 | 485.36 | 27.85 | 120852 + 70 | 1246902 | 34432519 | 290228 | 278840 | 36305 | 20836 | 349601.6 | 344351.3 | 40 | 41 | 153 | 171 |
| PN280 | 34275673 | 197 | 1 | 5 | 10 | 16 + 106 par | 2662004 | 91.99 | 1.002 | 0.83 | 501.06 | 28.76 | 121488 + 78 | 741910 | 34446930 | 244443 | 244443 | 22698 | 16275 | 284274.2 | 282176.9 | 46 | 46 | 185 | 198 |
| PN282 | 34147910 | 201 | 0 | 1 | 8 | 17 + 106 par | 2665925 | 91.99 | 1.001 | 0.98 | 501.34 | 28.77 | 121471 + 78 | 704437 | 34429447 | 244443 | 244443 | 22940 | 17281 | 281118.7 | 279085.1 | 47 | 47 | 182 | 195 |
| PN283 | 34097862 | 198 | 0 | 4 | 10 | 11 + 107 par | 2665697 | 91.95 | 1.002 | 1.42 | 500.39 | 28.8 | 121458 + 79 | 741874 | 34431980 | 252635 | 249097 | 22940 | 17307 | 284162.9 | 281979.7 | 46 | 47 | 182 | 195 |
| PN284 | 34210739 | 190 | 2 | 3 | 5 | 8 + 73 part | 2200825 | 92.76 | 1.001 | 3.49 | 481.55 | 27.5 | 121404 + 65 | 718124 | 34711185 | 267606 | 264660 | 44163 | 33097 | 278303.5 | 274932.7 | 47 | 48 | 166 | 177 |
| PN285 | 34405303 | 194 | 1 | 4 | 6 | 12 + 77 part | 2077406 | 92.83 | 1.002 | 2.87 | 458.68 | 25.48 | 121837 + 63 | 1039327 | 34762785 | 303682 | 301914 | 45204 | 30131 | 334139.4 | 329308.2 | 40 | 41 | 149 | 162 |
| PN286 | 34552897 | 205 | 0 | 4 | 11 | 12 + 87 part | 2174782 | 91.81 | 1.001 | 2.46 | 508.82 | 28.15 | 120978 + 86 | 870655 | 34356754 | 268479 | 266788 | 43514 | 18606 | 283921.7 | 277448 | 47 | 48 | 161 | 188 |
| PN287 | 35018667 | 212 | 1 | 6 | 11 | 20 + 98 part | 2415594 | 92.03 | 1.001 | 2.99 | 482.98 | 27.33 | 121066 + 80 | 703397 | 34442141 | 246731 | 244864 | 29591 | 18929 | 269862 | 266090.3 | 48 | 49 | 181 | 200 |
| PN288 | 32716563 | 204 | 1 | 3 | 8 | 14 + 93 part | 2103950 | 92.52 | 1.001 | 3.37 | 472.31 | 27.05 | 121306 + 80 | 857076 | 34629915 | 247450 | 244588 | 43049 | 27944 | 270844.8 | 266212.2 | 49 | 51 | 173 | 190 |
| PN289 | 34504798 | 203 | 1 | 4 | 7 | 12 + 83 part | 2094090 | 92.56 | 1.001 | 4.56 | 473.14 | 27.08 | 121319 + 80 | 857082 | 34640801 | 268518 | 264060 | 44903 | 29430 | 311587.5 | 306284.2 | 43 | 44 | 160 | 175 |

| | | | | | | | | | | | | | | | | | | | | | | | | | |
|---|---|---|---|---|---|---|---|---|---|---|---|---|---|---|---|---|---|---|---|---|---|---|---|---|---|
| PN290 | 34890976 | 215 | 1 | 0 | 14 | 42 + 108 par | 6570839 | 91.37 | 1.001 | 2.36 | 546.06 | 30.03 | 120877 + 88 | 829206 | 34210798 | 207630 | 238778 | - | 10766 | 251452.6 | 274320.7 | 55 | 48 | - | 222 |
| PN291 | 36471571 | 182 | 1 | 6 | 11 | 8 + 72 part | 2002693 | 92.83 | 1.03 | 3.54 | 461.6 | 26.57 | 121087 + 70 | 715041 | 35733718 | 289341 | 289341 | 44458 | 50201 | 296470.4 | 299289.7 | 44 | 44 | 163 | 156 |
| PN292 | 32579463 | 198 | 1 | 3 | 11 | 15 + 109 par | 2159929 | 92.49 | 1.001 | 1.81 | 470.55 | 26.82 | 121402 + 79 | 738397 | 34628200 | 262127 | 259714 | 33100 | 20509 | 287845.4 | 283305.2 | 44 | 45 | 175 | 195 |
| PN295 | 32307039 | 187 | 1 | 3 | 17 | 181 + 148 pa | 3652956 | 92.61 | 1.002 | 7.61 | 492.91 | 27.8 | 121775 + 80 | 744574 | 34682424 | 243080 | 250023 | 3252 | 24725 | 247642.5 | 253967 | 53 | 51 | 294 | 201 |
| PN296 | 34343610 | 207 | 0 | 2 | 3 | 12 + 64 part | 1994842 | 92.30 | 1.001 | 2.78 | 473.12 | 26.29 | 121079 + 72 | 964706 | 34546166 | 294198 | 281768 | 49906 | 26311 | 313787.5 | 306695.5 | 42 | 44 | 145 | 165 |
| PN297 | 34642433 | 228 | 0 | 6 | 16 | 292 + 164 pa | 3595915 | 91.58 | 1.002 | 3.91 | 558.99 | 30.83 | 120594 + 11 | 690826 | 34315215 | 202865 | 223289 | 802 | 9937 | 268157.3 | 272073.6 | 48 | 46 | 427 | 253 |
| PN298 | 34472668 | 204 | 0 | 1 | 5 | 9 + 68 part | 1992874 | 92.30 | 1.001 | 2.58 | 474.46 | 26.57 | 121063 + 73 | 1267082 | 34547454 | 294298 | 281768 | 47675 | 27643 | 330505.7 | 323067.2 | 42 | 43 | 143 | 163 |
| PN299 | 33580433 | 187 | 0 | 3 | 3 | 17 + 77 part | 2074041 | 92.42 | 1.003 | 2.09 | 473.66 | 26.14 | 121753 + 70 | 1225240 | 34655770 | 246717 | 245930 | 41891 | 23374 | 316159.5 | 310681.7 | 45 | 46 | 171 | 191 |
| PN300 | 32075072 | 183 | 1 | 5 | 15 | 85 + 114 par | 8577788 | 92.42 | 1.004 | 3.37 | 474.26 | 26.01 | 121808 + 71 | 978393 | 34676141 | 189783 | 246717 | - | 22503 | 245352.3 | 283873.3 | 61 | 48 | - | 190 |
| PN301 | 34295219 | 187 | 0 | 4 | 6 | 16 + 88 part | 2087380 | 92.41 | 1.003 | 1.82 | 470.58 | 25.79 | 121762 + 69 | 978915 | 34653126 | 254639 | 248521 | 34203 | 19892 | 297333.2 | 292211.6 | 45 | 46 | 168 | 189 |
| PN302 | 35912305 | 209 | 0 | 1 | 7 | 14 + 81 part | 2188044 | 92.42 | 1.001 | 2.5 | 485.02 | 27.36 | 121147 + 76 | 1082967 | 34597250 | 266813 | 265390 | 33759 | 20745 | 304102.9 | 299200.8 | 43 | 44 | 167 | 188 |
| PN303 | 35318549 | 195 | 0 | 2 | 8 | 5 + 77 part | 1954366 | 92.63 | 1.001 | 2.42 | 476.9 | 26.96 | 121450 + 67 | 1279128 | 34647932 | 285248 | 280820 | 42036 | 29587 | 355459.2 | 348284.2 | 39 | 40 | 155 | 174 |
| PN304 | 34974934 | 208 | 0 | 2 | 7 | 10 + 86 part | 2069067 | 92.89 | 1.002 | 3.19 | 467.93 | 27.22 | 121650 + 77 | 971026 | 34785252 | 246085 | 245638 | 42928 | 28334 | 287450.2 | 283385.6 | 46 | 47 | 166 | 180 |
| PN305 | 35155946 | 229 | 0 | 5 | 4 | 17 + 76 part | 2353646 | 91.82 | 1.001 | 3.15 | 538.81 | 30.42 | 120611 + 84 | 1263974 | 34372271 | 247726 | 240489 | 29999 | 15716 | 316415.9 | 310878.9 | 42 | 44 | 181 | 206 |
| PN306 | 32570054 | 193 | 1 | 3 | 14 | 15 + 126 par | 2657158 | 91.98 | 1.001 | 0.78 | 500.74 | 28.8 | 121480 + 77 | 741985 | 34423492 | 244443 | 244443 | 22708 | 15592 | 274875.8 | 272790.3 | 48 | 48 | 190 | 203 |
| PN307 | 34181407 | 193 | 1 | 3 | 12 | 16 + 123 par | 2656015 | 91.97 | 1.001 | 1.51 | 501.96 | 29.13 | 121443 + 81 | 741883 | 34433058 | 238347 | 238347 | 19920 | 15095 | 277317.4 | 275227.1 | 48 | 48 | 191 | 205 |
| PN308 | 35374482 | 203 | 0 | 6 | 8 | 7 + 96 part | 1911319 | 92.39 | 1.001 | 3.69 | 483.11 | 28.91 | 121117 + 67 | 904399 | 34585373 | 267878 | 264733 | 44458 | 25558 | 308372.1 | 301172 | 42 | 44 | 161 | 183 |
| PN309 | 34597665 | 227 | 0 | 9 | 19 | 50 + 162 par | 3368308 | 91.32 | 1.001 | 4.04 | 559.28 | 30.96 | 120640 + 93 | 737278 | 34188754 | 207462 | 207462 | 6837 | 9931 | 272378.6 | 273663.6 | 47 | 47 | 269 | 250 |
| PN312 | 34901797 | 191 | 0 | 8 | 18 | 10 + 109 par | 2388232 | 92.38 | 1.002 | 3.67 | 465.85 | 27.33 | 121154 + 65 | 890761 | 34591015 | 247071 | 241166 | 31989 | 22960 | 290284.1 | 287129.1 | 46 | 47 | 172 | 185 |
| PN313 | 35228806 | 193 | 0 | 0 | 15 | 9 + 99 part | 2393131 | 92.36 | 1.002 | 2.9 | 465.66 | 27.09 | 121155 + 64 | 890379 | 34590134 | 244076 | 244076 | 32176 | 24148 | 297633.1 | 294444.7 | 45 | 45 | 170 | 183 |
| PN314 | 35216684 | 199 | 0 | 5 | 15 | 22 + 94 part | 1931401 | 92.44 | 1.001 | 2.76 | 481.83 | 28.59 | 121147 + 68 | 904110 | 34608328 | 267878 | 267577 | 42579 | 23245 | 320033.9 | 312835.7 | 41 | 42 | 159 | 182 |
| PN315 | 33673433 | 193 | 1 | 4 | 16 | 31 + 133 par | 2665082 | 92.00 | 1.002 | 1.77 | 500.55 | 28.88 | 121466 + 79 | 741874 | 34449214 | 239823 | 238353 | 22459 | 16275 | 279509.5 | 277525.6 | 47 | 48 | 188 | 201 |
| PN316 | 33307352 | 192 | 1 | 1 | 24 | 58 + 156 par | 3737155 | 91.98 | 1.002 | 1.83 | 502.6 | 28.92 | 121480 + 80 | 741883 | 34461234 | 225682 | 227386 | 881 | 15817 | 263990.2 | 269775.9 | 51 | 50 | 357 | 205 |
| PN317 | 33058060 | 170 | 1 | 0 | 14 | 420 + 104 pa | 3817879 | 92.85 | 1.001 | 9.25 | 476.3 | 26.78 | 121862 + 67 | 846327 | 34748612 | 239155 | 244002 | 535 | 30529 | 274343.1 | 283144.8 | 50 | 47 | 363 | 175 |
| PN318 | 33138648 | 200 | 0 | 3 | 5 | 15 + 77 part | 2121272 | 92.89 | 1.002 | 3.53 | 474.01 | 26.51 | 121907 + 74 | 1269432 | 34785804 | 303454 | 288828 | 38295 | 28423 | 350370.2 | 346004.6 | 40 | 41 | 158 | 171 |
| PN319 | 35037845 | 185 | 0 | 3 | 16 | 202 + 112 pa | 3153240 | 92.34 | 1.002 | 9.33 | 483.72 | 27.61 | 121603 + 74 | 973570 | 34580889 | 245782 | 247171 | 14035 | 25565 | 288149.9 | 290933.9 | 47 | 46 | 207 | 189 |
| PN320 | 34191152 | 188 | 2 | 2 | 6 | 10 + 68 part | 2204225 | 92.74 | 1.001 | 3.17 | 481.83 | 27.7 | 121403 + 64 | 718071 | 34722499 | 289870 | 281777 | 44425 | 34939 | 285804 | 282306.9 | 45 | 46 | 162 | 172 |
| PN321 | 33220467 | 206 | 0 | 1 | 13 | 16 + 97 part | 2119989 | 92.77 | 1.002 | 4.02 | 455.35 | 26.48 | 121699 + 71 | 1031337 | 34735640 | 309629 | 309629 | 39687 | 28192 | 330318.7 | 325764.1 | 40 | 40 | 149 | 163 |
| PN322 | 31218727 | 205 | 0 | 1 | 17 | 18 + 110 par | 2059646 | 92.78 | 1.002 | 2.61 | 456.43 | 26.28 | 121689 + 73 | 1031531 | 34742830 | 284755 | 281647 | 37912 | 25489 | 319013.8 | 314107.3 | 41 | 42 | 156 | 172 |
| PN323 | 35614249 | 205 | 1 | 6 | 7 | 3 + 79 part | 2369717 | 93.14 | 1.001 | 2.19 | 468.21 | 26.49 | 121900 + 78 | 817027 | 34845769 | 273458 | 273458 | 29081 | 28033 | 304798.5 | 303601.7 | 43 | 43 | 165 | 170 |
| PN324 | 34816892 | 212 | 1 | 2 | 6 | 334 + 110 pa | 2642974 | 93.51 | 1.002 | 2.94 | 472.8 | 26.6 | 121980 + 10 | 816689 | 35034902 | 267354 | 281925 | 20848 | 28541 | 295711.8 | 298265.6 | 44 | 43 | 185 | 174 |
| PN325 | 34411184 | 195 | 0 | 0 | 6 | 10 + 70 part | 1815500 | 92.39 | 1 | 3.01 | 480.95 | 28.42 | 121202 + 69 | 968701 | 34555922 | 294554 | 274087 | 48873 | 24598 | 324214.5 | 315612.7 | 41 | 42 | 146 | 173 |

| | | | | | | | | | | | | | | | | | | | | | | | | | |
|---|---|---|---|---|---|---|---|---|---|---|---|---|---|---|---|---|---|---|---|---|---|---|---|---|---|
| PN326 | 34712306 | 174 | 0 | 2 | 9 | 20 + 87 part | 2593467 | 92.67 | 1.002 | 2.46 | 471.97 | 26.27 | 121711 + 61 | 847182 | 34717272 | 276764 | 276764 | 25489 | 24226 | 298452.1 | 297857.2 | 44 | 44 | 173 | 175 |
| PN328 | 31623631 | 189 | 0 | 7 | 21 | 42 + 138 par | 2875217 | 91.83 | 1.002 | 4.95 | 463.15 | 27.22 | 120618 + 78 | 1246669 | 34385386 | 267721 | 267721 | 15572 | 13358 | 293855.2 | 292958.6 | 46 | 46 | 190 | 197 |
| PN329 | 32283154 | 197 | 1 | 2 | 24 | 39 + 126 par | 2912943 | 91.87 | 1.001 | 4.29 | 461.56 | 26.92 | 120624 + 76 | 1232624 | 34380681 | 282145 | 278818 | 17018 | 15202 | 319367.6 | 318720.9 | 43 | 44 | 189 | 193 |
| PN330 | 35158111 | 191 | 2 | 1 | 10 | 14 + 85 part | 2141839 | 92.64 | 1.001 | 1.96 | 475.95 | 27.52 | 121341 + 64 | 783407 | 34665780 | 243272 | 242259 | 44605 | 30345 | 277245.9 | 273000.5 | 48 | 49 | 165 | 179 |
| PN331 | 33568614 | 176 | 0 | 1 | 6 | 6 + 82 part | 2138845 | 92.32 | 1.001 | 1.79 | 471.73 | 26.97 | 121066 + 68 | 1146582 | 34543906 | 269701 | 267552 | 39787 | 25336 | 346736.6 | 340394.7 | 40 | 41 | 156 | 175 |
| PN332 | 33225103 | 190 | 0 | 2 | 11 | 22 + 94 part | 2393081 | 92.39 | 1.001 | 2.09 | 472.63 | 26.63 | 121264 + 80 | 728936 | 34580440 | 298394 | 298394 | 29987 | 20830 | 289675.4 | 286663.9 | 44 | 44 | 175 | 189 |
| PN333 | 31119912 | 201 | 0 | 3 | 9 | 1943 + 98 pa | 3229563 | 92.83 | 1.001 | 3.45 | 463.37 | 27.12 | 121694 + 80 | 1032263 | 34749916 | 259632 | 260163 | 11134 | 25558 | 307622.6 | 312533.2 | 44 | 43 | 205 | 175 |
| PN334 | 33943440 | 197 | 2 | 2 | 11 | 19 + 115 par | 2670284 | 91.99 | 1.002 | 2.29 | 499.83 | 28.77 | 121526 + 76 | 742063 | 34445046 | 244443 | 239823 | 22940 | 16275 | 288944.3 | 286870.5 | 45 | 46 | 182 | 194 |
| PN335 | 33194204 | 197 | 2 | 3 | 7 | 18 + 116 par | 2664106 | 91.99 | 1.001 | 1.97 | 500.97 | 28.65 | 121474 + 81 | 741883 | 34441525 | 240099 | 239823 | 22940 | 15713 | 285288.1 | 283207.1 | 46 | 47 | 187 | 200 |
| PN336 | 34909221 | 201 | 0 | 3 | 9 | 8 + 79 part | 2464244 | 92.34 | 1 | 1.64 | 486.93 | 27.95 | 121013 + 75 | 936678 | 34537249 | 259740 | 255617 | 31916 | 20282 | 303484 | 300619.7 | 43 | 44 | 173 | 186 |
| PN337 | 32474933 | 197 | 2 | 0 | 13 | 16 + 140 par | 2648066 | 92.01 | 1.002 | 0.95 | 502.34 | 28.79 | 121325 + 81 | 741881 | 34459055 | 230340 | 225682 | 19357 | 15592 | 268005.3 | 266032.5 | 49 | 50 | 205 | 219 |
| PN338 | 32953219 | 195 | 2 | 7 | 16 | 26 + 132 par | 2642563 | 92.00 | 1.001 | 2.02 | 502.27 | 28.88 | 121492 + 79 | 741874 | 34423751 | 249097 | 239823 | 22940 | 16114 | 279821.9 | 277661.1 | 46 | 47 | 189 | 202 |
| PN340 | 33539083 | 197 | 1 | 4 | 12 | 17 + 114 par | 2669800 | 91.97 | 1.001 | 1.6 | 500.15 | 28.69 | 121436 + 79 | 741874 | 34428328 | 238048 | 236599 | 21548 | 15592 | 271689.7 | 269701.3 | 49 | 50 | 190 | 203 |
| PN341 | 34449704 | 197 | 0 | 2 | 6 | 13 + 85 part | 2023146 | 92.92 | 1.001 | 3.1 | 472.42 | 27.28 | 121605 + 78 | 1020560 | 34777519 | 285207 | 282058 | 42994 | 27924 | 334805.7 | 329672 | 39 | 40 | 152 | 168 |
| PN342 | 35731752 | 179 | 2 | 2 | 10 | 9 + 76 part | 2041706 | 92.95 | 1.003 | 4.16 | 480.65 | 26.47 | 121780 + 67 | 1019945 | 34850844 | 313329 | 308520 | 42346 | 31302 | 343801.7 | 339392.5 | 38 | 39 | 140 | 151 |
| PN343 | 34061885 | 193 | 1 | 3 | 10 | 17 + 109 par | 2652617 | 91.99 | 1.002 | 1.86 | 501.52 | 28.94 | 121363 + 77 | 741883 | 34442868 | 244443 | 244443 | 22940 | 18061 | 280586.1 | 278475.7 | 47 | 47 | 187 | 199 |
| PN344 | 33137631 | 199 | 0 | 3 | 8 | 15 + 90 part | 1942097 | 92.85 | 1.001 | 3.09 | 461.08 | 26.62 | 121724 + 80 | 1032232 | 34751068 | 262182 | 260163 | 46066 | 25558 | 318261.5 | 312442.2 | 42 | 43 | 156 | 173 |
| PN345 | 33447150 | 200 | 0 | 2 | 11 | 15 + 126 par | 2659771 | 91.98 | 1.002 | 1.79 | 500.69 | 28.73 | 121492 + 78 | 741874 | 34447167 | 230854 | 230854 | 21548 | 15713 | 280409.7 | 278358.9 | 47 | 47 | 194 | 207 |
| PN346 | 31969638 | 196 | 0 | 4 | 16 | 25 + 139 par | 2640754 | 92.01 | 1.002 | 1.17 | 503.68 | 29.06 | 121484 + 80 | 702637 | 34463585 | 238048 | 236802 | 21304 | 15098 | 265889.9 | 263913.2 | 49 | 50 | 194 | 207 |
| PN347 | 34881472 | 200 | 1 | 5 | 3 | 6 + 78 part | 1941055 | 92.59 | 1.002 | 3.26 | 477.21 | 27.69 | 121675 + 75 | 1012195 | 34672029 | 277345 | 265712 | 47562 | 28330 | 332357.7 | 325556.9 | 41 | 42 | 146 | 164 |
| PN348 | 35550507 | 200 | 1 | 6 | 7 | 5 + 78 part | 2152190 | 92.40 | 1.001 | 1.78 | 457.55 | 26.4 | 121167 + 71 | 872244 | 34589357 | 301238 | 300218 | 40111 | 27575 | 323385.9 | 317792.5 | 40 | 41 | 158 | 175 |
| PN349 | 33896062 | 194 | 0 | 7 | 14 | 10 + 94 part | 2304155 | 92.27 | 1.001 | 1.84 | 478.3 | 27.52 | 121064 + 74 | 1005321 | 34546369 | 256338 | 247606 | 40151 | 25030 | 290344.2 | 286201.6 | 46 | 47 | 172 | 187 |
| PN350 | 33885850 | 205 | 0 | 5 | 7 | 12 + 72 part | 1956957 | 92.85 | 1.001 | 3.41 | 462.77 | 27.02 | 121727 + 78 | 849354 | 34762672 | 274603 | 262182 | 47549 | 32877 | 319568.8 | 313885.4 | 41 | 42 | 151 | 166 |
| PN351 | 35341973 | 200 | 0 | 5 | 5 | 21 + 67 part | 2132360 | 92.37 | 1.001 | 2.74 | 492.29 | 27.18 | 121492 + 73 | 1246678 | 34579086 | 302634 | 291893 | 36689 | 25352 | 350277.2 | 343960.4 | 38 | 39 | 157 | 176 |
| PN352 | 34093157 | 196 | 0 | 3 | 10 | 12 + 81 part | 2360265 | 92.80 | 1.001 | 2.33 | 483.28 | 27.28 | 121563 + 73 | 1029896 | 34741993 | 295986 | 295986 | 33792 | 28747 | 310338.5 | 307995.9 | 43 | 43 | 169 | 177 |
| PN353 | 35867682 | 212 | 0 | 3 | 4 | 24 + 83 part | 2718152 | 91.99 | 1 | 2.28 | 502.45 | 28.95 | 121425 + 76 | 937335 | 34404727 | 258266 | 258266 | 20848 | 15380 | 299404.8 | 297555.4 | 44 | 44 | 185 | 197 |
| PN354 | 34324324 | 182 | 0 | 4 | 13 | 3 + 83 part | 1724801 | 92.42 | 1.003 | 2.28 | 471.9 | 26.82 | 121658 + 68 | 929226 | 34641505 | 260798 | 245396 | 48185 | 26925 | 308335.9 | 299955.7 | 42 | 44 | 155 | 180 |
| PN355 | 33654922 | 205 | 0 | 3 | 9 | 8 + 79 part | 1958954 | 92.87 | 1.001 | 3.45 | 461.84 | 27.01 | 121727 + 78 | 1032469 | 34758489 | 285087 | 281768 | 47434 | 27519 | 335962.8 | 329982.4 | 40 | 41 | 146 | 162 |
| PN356 | 33025461 | 197 | 1 | 5 | 9 | 16 + 91 part | 1947986 | 92.87 | 1.001 | 3.7 | 462.85 | 26.69 | 121718 + 77 | 849119 | 34762548 | 262182 | 260163 | 46066 | 27944 | 317377 | 311741.6 | 42 | 43 | 153 | 169 |
| PN357 | 35353236 | 202 | 1 | 5 | 9 | 18 + 85 part | 2049434 | 92.46 | 1.001 | 2.78 | 465.57 | 25.66 | 121379 + 79 | 744669 | 34613549 | 292996 | 282771 | 37802 | 22833 | 296810.8 | 291049.6 | 43 | 44 | 163 | 184 |
| PN358 | 33347437 | 220 | 1 | 2 | 18 | 239 + 125 pa | 3391021 | 92.47 | 1.002 | 8.83 | 482.12 | 27.14 | 121392 + 85 | 706115 | 34633562 | 259318 | 262490 | 7628 | 21448 | 266371.9 | 270980.9 | 50 | 48 | 237 | 194 |
| PN359 | 35056211 | 196 | 0 | 5 | 12 | 8 + 82 part | 2384175 | 92.44 | 1.001 | 3.11 | 483.07 | 27.16 | 121575 + 78 | 901773 | 34599268 | 259632 | 259632 | 30345 | 20836 | 313145.6 | 309775.8 | 43 | 43 | 169 | 183 |

| | | | | | | | | | | | | | | | | | | | | | | | | | |
|---|---|---|---|---|---|---|---|---|---|---|---|---|---|---|---|---|---|---|---|---|---|---|---|---|---|
| PN360 | 36202479 | 203 | 2 | 3 | 3 | 12 + 73 part | 2090207 | 92.38 | 1.002 | 3.01 | 464.11 | 25.64 | 121401 + 77 | 746763 | 34592294 | 306617 | 305836 | 38045 | 26453 | 312547 | 306634.3 | 41 | 42 | 155 | 176 |
| PN361 | 34610615 | 208 | 0 | 3 | 10 | 10 + 90 part | 2292766 | 92.80 | 1.002 | 3.2 | 481.58 | 27.03 | 121629 + 71 | 1224363 | 34756160 | 268502 | 268502 | 36593 | 28453 | 325645.1 | 322723.2 | 42 | 42 | 159 | 169 |
| PN362 | 35537458 | 196 | 0 | 2 | 15 | 3 + 85 part | 2429896 | 92.34 | 1.001 | 1.29 | 483.81 | 28.03 | 121023 + 71 | 937224 | 34567979 | 255617 | 251274 | 33611 | 21414 | 304533.1 | 301362 | 43 | 44 | 171 | 184 |
| PN363 | 34093518 | 206 | 0 | 2 | 13 | 13 + 102 par | 2291456 | 92.81 | 1.001 | 2.9 | 482.44 | 27.14 | 121707 + 71 | 754641 | 34747036 | 290904 | 274169 | 34197 | 28984 | 309039.5 | 306272.3 | 42 | 43 | 158 | 167 |
| PN364 | 32546590 | 193 | 1 | 5 | 14 | 41 + 128 par | 2315075 | 92.48 | 1.001 | 3.73 | 501.31 | 28.86 | 121396 + 79 | 865751 | 34615922 | 245796 | 244763 | 30459 | 20483 | 291125.9 | 287625.8 | 46 | 47 | 185 | 201 |
| PN365 | 31896377 | 201 | 0 | 3 | 10 | 15 + 84 part | 1963374 | 92.86 | 1.001 | 2.92 | 461.77 | 26.85 | 121708 + 78 | 1032069 | 34754876 | 274603 | 260160 | 46119 | 27944 | 327469.8 | 321671 | 40 | 42 | 153 | 169 |
| PN366 | 35138262 | 192 | 1 | 4 | 10 | 6 + 76 part | 2048340 | 92.28 | 1.002 | 3.64 | 480.24 | 26.7 | 121457 + 76 | 1079503 | 34554949 | 306340 | 294450 | 40111 | 23899 | 324934.8 | 318138 | 40 | 41 | 160 | 183 |
| PN367 | 35581372 | 190 | 0 | 6 | 12 | 4 + 78 part | 2016990 | 92.69 | 1.009 | 2.19 | 480.19 | 27.42 | 121448 + 68 | 1101165 | 34959351 | 276986 | 274802 | 42036 | 37110 | 326839.6 | 323548.4 | 41 | 42 | 159 | 167 |
| PN368 | 32140749 | 209 | 0 | 3 | 9 | 14 + 82 part | 1959877 | 92.85 | 1.001 | 2.93 | 461.46 | 26.81 | 121717 + 80 | 1031824 | 34750906 | 274603 | 260163 | 46066 | 27088 | 325201.5 | 319471 | 40 | 42 | 154 | 170 |
| PN369 | 35505627 | 199 | 1 | 2 | 10 | 8 + 88 part | 2552470 | 92.67 | 1.001 | 3.26 | 487.57 | 27.63 | 121728 + 73 | 992833 | 34694494 | 261848 | 261848 | 30366 | 28204 | 297300.2 | 296347 | 45 | 45 | 178 | 182 |
| PN370 | 33927991 | 206 | 1 | 5 | 8 | 9 + 71 part | 1961608 | 92.87 | 1.001 | 3.19 | 461.71 | 26.87 | 121727 + 79 | 1032113 | 34748457 | 285197 | 281768 | 47434 | 27944 | 342244.9 | 336173.3 | 39 | 40 | 146 | 161 |
| PN371 | 35470577 | 186 | 1 | 3 | 20 | 388 + 123 pa | 3743785 | 92.37 | 1.003 | 10.48 | 514.61 | 28.92 | 121312 + 80 | 883028 | 34624141 | 260364 | 272731 | 1028 | 18170 | 321612.8 | 330136.1 | 42 | 40 | 351 | 191 |
| PN372 | 34523505 | 180 | 1 | 3 | 12 | 74 + 83 part | 7271151 | 92.42 | 1.002 | 2.62 | 472.67 | 26.99 | 121681 + 68 | 929226 | 34638831 | 235760 | 261999 | - | 26468 | 277241.4 | 310858.5 | 51 | 42 | - | 179 |
| PN373 | 33725048 | 203 | 0 | 6 | 5 | 11 + 80 part | 1959928 | 92.87 | 1.001 | 3.48 | 461.28 | 26.9 | 121729 + 78 | 1032260 | 34757568 | 281768 | 274603 | 45984 | 28235 | 338011.1 | 332001.6 | 39 | 40 | 149 | 164 |
| PN374 | 32760217 | 204 | 1 | 5 | 9 | 9 + 82 part | 1958882 | 92.87 | 1.001 | 3.98 | 460.94 | 26.89 | 121725 + 78 | 1032077 | 34749438 | 274603 | 263811 | 46285 | 27944 | 324347.9 | 318608.3 | 41 | 42 | 151 | 167 |
| PN375 | 35851407 | 174 | 0 | 1 | 6 | 8 + 66 part | 2166824 | 92.44 | 1.002 | 2.35 | 470.14 | 26.83 | 121077 + 69 | 972889 | 34619794 | 278669 | 268071 | 38295 | 25852 | 331415.6 | 326079.6 | 40 | 42 | 158 | 175 |
| PN376 | 35532618 | 193 | 0 | 2 | 12 | 2 + 65 part | 2296535 | 92.73 | 1.001 | 2.63 | 497.68 | 27.77 | 121730 + 74 | 910867 | 34715864 | 256114 | 253561 | 34554 | 27912 | 282382.4 | 279645.4 | 47 | 48 | 179 | 189 |
| PN377 | 35578305 | 189 | 0 | 0 | 15 | 5 + 95 part | 2302938 | 92.72 | 1.001 | 2.6 | 496.26 | 27.63 | 121712 + 75 | 910924 | 34683061 | 251255 | 247405 | 31628 | 27311 | 277934.7 | 275177.3 | 48 | 49 | 186 | 197 |
| PN378 | 34673012 | 204 | 0 | 1 | 3 | 12 + 66 part | 1995190 | 92.27 | 1.001 | 2.82 | 472.87 | 26.25 | 121036 + 75 | 964505 | 34541770 | 294847 | 281768 | 47434 | 25489 | 315992.6 | 308830.5 | 41 | 43 | 148 | 169 |
| PN379 | 34285040 | 195 | 1 | 4 | 13 | 9 + 84 part | 2047806 | 92.61 | 1.002 | 2.24 | 466.23 | 26.86 | 121345 + 73 | 849283 | 34678724 | 312257 | 312060 | 40211 | 25558 | 318706.4 | 313097.4 | 40 | 41 | 153 | 171 |
| PN380 | 34618494 | 197 | 0 | 3 | 8 | 9 + 87 part | 2424512 | 92.37 | 1.001 | 1.2 | 487.63 | 28.11 | 121037 + 74 | 801915 | 34558659 | 247131 | 246121 | 33611 | 21414 | 295577 | 292649 | 45 | 46 | 175 | 187 |
| mean | 30654151 | 192.5 | 0.88 | 3 | 13.2 | 163.40 + 14 | 2508764 | 92.37 | 1.0014 | 5.5315 | 470.69164 | 26.68 | 121351.26 + | 859445.74 | 34583401 | 245263 | 243106 | 30622.6 | 19745 | 279997.31 | 277602.76 | 48.78 | 49.31 | 202.4 | 207.8 |
| STD | 5342199 | 22.19 | 0.89 | 6.78 | 6.2 | 599.08 + 71. | 1027969.4 | 0.75 | 0.0023 | 18.034 | 49.279834 | 2.3869 | 638.71 + 16 | 179365.63 | 298913.01 | 43929.33 | 42030.2 | 13523.2 | 9627.28 | 45096.914 | 43782.235 | 10.47 | 9.956 | 91.91 | 46.72 |

**STable 5: Group asignment of 360 P. nodorum isolates by Discriminant Analysis of Principal Components**

| Isolate | grp_1 | grp_2 | grp_3 | grp_4 | grp_5 | grp_6 | grp_7 | grp_8 |
|---|---|---|---|---|---|---|---|---|
| DIP1_WAC1141 | 0.00 | 1.00 | 0.00 | 0.00 | 0.00 | 0.00 | 0.00 | 0.00 |
| DIP10_WAC1201 | 0.00 | 0.00 | 0.00 | 0.00 | 0.00 | 1.00 | 0.00 | 0.00 |
| DIP100_WAC14144 | 1.00 | 0.00 | 0.00 | 0.00 | 0.00 | 0.00 | 0.00 | 0.00 |
| DIP102_WAC2816 | 0.00 | 1.00 | 0.00 | 0.00 | 0.00 | 0.00 | 0.00 | 0.00 |
| DIP103_WAC2812 | 1.00 | 0.00 | 0.00 | 0.00 | 0.00 | 0.00 | 0.00 | 0.00 |
| DIP104_WAC8397 | 1.00 | 0.00 | 0.00 | 0.00 | 0.00 | 0.00 | 0.00 | 0.00 |
| DIP105_WAC2804 | 0.00 | 1.00 | 0.00 | 0.00 | 0.00 | 0.00 | 0.00 | 0.00 |
| DIP106_WAC1544 | 1.00 | 0.00 | 0.00 | 0.00 | 0.00 | 0.00 | 0.00 | 0.00 |
| DIP107_WAC2799 | 1.00 | 0.00 | 0.00 | 0.00 | 0.00 | 0.00 | 0.00 | 0.00 |
| DIP108_WAC4312 | 0.00 | 0.00 | 0.00 | 1.00 | 0.00 | 0.00 | 0.00 | 0.00 |
| DIP109_WAC1198 | 0.00 | 0.00 | 0.00 | 0.00 | 0.00 | 1.00 | 0.00 | 0.00 |
| DIP11_WAC1494 | 0.00 | 1.00 | 0.00 | 0.00 | 0.00 | 0.00 | 0.00 | 0.00 |
| DIP110_WAC8380 | 1.00 | 0.00 | 0.00 | 0.00 | 0.00 | 0.00 | 0.00 | 0.00 |
| DIP111_WAC1194 | 0.00 | 0.00 | 0.00 | 0.00 | 0.00 | 1.00 | 0.00 | 0.00 |
| DIP112_WAC4307 | 1.00 | 0.00 | 0.00 | 0.00 | 0.00 | 0.00 | 0.00 | 0.00 |
| DIP12_WAC1495 | 0.00 | 1.00 | 0.00 | 0.00 | 0.00 | 0.00 | 0.00 | 0.00 |
| DIP13_WAC1496 | 0.00 | 0.00 | 1.00 | 0.00 | 0.00 | 0.00 | 0.00 | 0.00 |
| DIP14_WAC1497 | 0.00 | 0.00 | 0.00 | 0.00 | 0.00 | 1.00 | 0.00 | 0.00 |
| DIP15_WAC1498 | 1.00 | 0.00 | 0.00 | 0.00 | 0.00 | 0.00 | 0.00 | 0.00 |
| DIP16_WAC1500 | 1.00 | 0.00 | 0.00 | 0.00 | 0.00 | 0.00 | 0.00 | 0.00 |
| DIP17_WAC1508 | 1.00 | 0.00 | 0.00 | 0.00 | 0.00 | 0.00 | 0.00 | 0.00 |
| DIP18_WAC1545 | 1.00 | 0.00 | 0.00 | 0.00 | 0.00 | 0.00 | 0.00 | 0.00 |
| DIP19_WAC1546 | 1.00 | 0.00 | 0.00 | 0.00 | 0.00 | 0.00 | 0.00 | 0.00 |
| DIP2_WAC1178 | 0.93 | 0.00 | 0.07 | 0.00 | 0.00 | 0.00 | 0.00 | 0.00 |
| DIP20_WAC1547 | 0.00 | 0.00 | 1.00 | 0.00 | 0.00 | 0.00 | 0.00 | 0.00 |
| DIP21_WAC1548 | 0.00 | 1.00 | 0.00 | 0.00 | 0.00 | 0.00 | 0.00 | 0.00 |
| DIP22_WAC1549 | 1.00 | 0.00 | 0.00 | 0.00 | 0.00 | 0.00 | 0.00 | 0.00 |
| DIP23_WAC1550 | 1.00 | 0.00 | 0.00 | 0.00 | 0.00 | 0.00 | 0.00 | 0.00 |
| DIP24_WAC1551 | 1.00 | 0.00 | 0.00 | 0.00 | 0.00 | 0.00 | 0.00 | 0.00 |
| DIP25_WAC1552 | 1.00 | 0.00 | 0.00 | 0.00 | 0.00 | 0.00 | 0.00 | 0.00 |
| DIP27_WAC1564 | 0.00 | 1.00 | 0.00 | 0.00 | 0.00 | 0.00 | 0.00 | 0.00 |
| DIP28_WAC1565 | 0.00 | 0.00 | 0.00 | 0.00 | 0.00 | 1.00 | 0.00 | 0.00 |
| DIP29_WAC2217 | 1.00 | 0.00 | 0.00 | 0.00 | 0.00 | 0.00 | 0.00 | 0.00 |
| DIP3_WAC1179 | 1.00 | 0.00 | 0.00 | 0.00 | 0.00 | 0.00 | 0.00 | 0.00 |
| DIP30_WAC2284 | 0.00 | 1.00 | 0.00 | 0.00 | 0.00 | 0.00 | 0.00 | 0.00 |
| DIP31_WAC2286 | 0.00 | 1.00 | 0.00 | 0.00 | 0.00 | 0.00 | 0.00 | 0.00 |
| DIP32_WAC2286 | 1.00 | 0.00 | 0.00 | 0.00 | 0.00 | 0.00 | 0.00 | 0.00 |
| DIP33_WAC2796 | 0.00 | 0.00 | 0.00 | 1.00 | 0.00 | 0.00 | 0.00 | 0.00 |
| DIP34_WAC2797 | 1.00 | 0.00 | 0.00 | 0.00 | 0.00 | 0.00 | 0.00 | 0.00 |
| DIP35_WAC2798 | 1.00 | 0.00 | 0.00 | 0.00 | 0.00 | 0.00 | 0.00 | 0.00 |
| DIP36_WAC2802 | 1.00 | 0.00 | 0.00 | 0.00 | 0.00 | 0.00 | 0.00 | 0.00 |
| DIP37_WAC2803 | 1.00 | 0.00 | 0.00 | 0.00 | 0.00 | 0.00 | 0.00 | 0.00 |
| DIP38_WAC2805 | 0.00 | 1.00 | 0.00 | 0.00 | 0.00 | 0.00 | 0.00 | 0.00 |
| DIP39_WAC2806 | 0.46 | 0.00 | 0.54 | 0.00 | 0.00 | 0.00 | 0.00 | 0.00 |
| DIP4_WAC1188 | 0.00 | 0.00 | 0.00 | 0.00 | 0.00 | 1.00 | 0.00 | 0.00 |
| DIP40_WAC2808 | 0.00 | 0.00 | 1.00 | 0.00 | 0.00 | 0.00 | 0.00 | 0.00 |
| DIP41_WAC2809 | 1.00 | 0.00 | 0.00 | 0.00 | 0.00 | 0.00 | 0.00 | 0.00 |

| | | | | | | | | |
|---|---|---|---|---|---|---|---|---|
| DIP42_WAC2817 | 0.00 | 1.00 | 0.00 | 0.00 | 0.00 | 0.00 | 0.00 | 0.00 |
| DIP43_WAC4292 | 1.00 | 0.00 | 0.00 | 0.00 | 0.00 | 0.00 | 0.00 | 0.00 |
| DIP44_WAC4301 | 1.00 | 0.00 | 0.00 | 0.00 | 0.00 | 0.00 | 0.00 | 0.00 |
| DIP45_WAC4304 | 0.00 | 1.00 | 0.00 | 0.00 | 0.00 | 0.00 | 0.00 | 0.00 |
| DIP46_WAC4308 | 0.00 | 0.00 | 1.00 | 0.00 | 0.00 | 0.00 | 0.00 | 0.00 |
| DIP47_WAC4310 | 1.00 | 0.00 | 0.00 | 0.00 | 0.00 | 0.00 | 0.00 | 0.00 |
| DIP48_WAC4311 | 1.00 | 0.00 | 0.00 | 0.00 | 0.00 | 0.00 | 0.00 | 0.00 |
| DIP49_WAC4316 | 0.00 | 1.00 | 0.00 | 0.00 | 0.00 | 0.00 | 0.00 | 0.00 |
| DIP5_WAC1190 | 0.00 | 0.00 | 0.00 | 0.00 | 0.00 | 1.00 | 0.00 | 0.00 |
| DIP50_WAC4317 | 1.00 | 0.00 | 0.00 | 0.00 | 0.00 | 0.00 | 0.00 | 0.00 |
| DIP51_WAC4318 | 1.00 | 0.00 | 0.00 | 0.00 | 0.00 | 0.00 | 0.00 | 0.00 |
| DIP55_WAC8383 | 1.00 | 0.00 | 0.00 | 0.00 | 0.00 | 0.00 | 0.00 | 0.00 |
| DIP56_WAC8387 | 1.00 | 0.00 | 0.00 | 0.00 | 0.00 | 0.00 | 0.00 | 0.00 |
| DIP57_WAC8391 | 1.00 | 0.00 | 0.00 | 0.00 | 0.00 | 0.00 | 0.00 | 0.00 |
| DIP58_WAC8399 | 1.00 | 0.00 | 0.00 | 0.00 | 0.00 | 0.00 | 0.00 | 0.00 |
| DIP6_WAC1193 | 0.00 | 0.00 | 0.00 | 0.00 | 0.00 | 1.00 | 0.00 | 0.00 |
| DIP60_WAC8417 | 1.00 | 0.00 | 0.00 | 0.00 | 0.00 | 0.00 | 0.00 | 0.00 |
| DIP62_WAC13864 | 1.00 | 0.00 | 0.00 | 0.00 | 0.00 | 0.00 | 0.00 | 0.00 |
| DIP63_WAC13865 | 1.00 | 0.00 | 0.00 | 0.00 | 0.00 | 0.00 | 0.00 | 0.00 |
| DIP64_WAC13866 | 1.00 | 0.00 | 0.00 | 0.00 | 0.00 | 0.00 | 0.00 | 0.00 |
| DIP66_WAC13868 | 1.00 | 0.00 | 0.00 | 0.00 | 0.00 | 0.00 | 0.00 | 0.00 |
| DIP68_WAC13870 | 1.00 | 0.00 | 0.00 | 0.00 | 0.00 | 0.00 | 0.00 | 0.00 |
| DIP69_WAC13871 | 1.00 | 0.00 | 0.00 | 0.00 | 0.00 | 0.00 | 0.00 | 0.00 |
| DIP7_WAC1195 | 0.00 | 0.00 | 0.00 | 0.00 | 0.00 | 1.00 | 0.00 | 0.00 |
| DIP70_WAC13872 | 1.00 | 0.00 | 0.00 | 0.00 | 0.00 | 0.00 | 0.00 | 0.00 |
| DIP71_WAC13873 | 1.00 | 0.00 | 0.00 | 0.00 | 0.00 | 0.00 | 0.00 | 0.00 |
| DIP72_WAC13957 | 0.00 | 1.00 | 0.00 | 0.00 | 0.00 | 0.00 | 0.00 | 0.00 |
| DIP73_WAC13958 | 0.00 | 0.00 | 0.00 | 1.00 | 0.00 | 0.00 | 0.00 | 0.00 |
| DIP74_WAC13959 | 0.00 | 0.00 | 0.00 | 1.00 | 0.00 | 0.00 | 0.00 | 0.00 |
| DIP75_WAC13960 | 0.00 | 0.00 | 0.00 | 1.00 | 0.00 | 0.00 | 0.00 | 0.00 |
| DIP76_WAC13966 | 1.00 | 0.00 | 0.00 | 0.00 | 0.00 | 0.00 | 0.00 | 0.00 |
| DIP77_WAC13967 | 1.00 | 0.00 | 0.00 | 0.00 | 0.00 | 0.00 | 0.00 | 0.00 |
| DIP79_WAC13969 | 1.00 | 0.00 | 0.00 | 0.00 | 0.00 | 0.00 | 0.00 | 0.00 |
| DIP8_WAC1197 | 0.00 | 0.00 | 0.00 | 0.00 | 0.00 | 1.00 | 0.00 | 0.00 |
| DIP82_WAC13979 | 0.96 | 0.00 | 0.04 | 0.00 | 0.00 | 0.00 | 0.00 | 0.00 |
| DIP83_WAC14056 | 1.00 | 0.00 | 0.00 | 0.00 | 0.00 | 0.00 | 0.00 | 0.00 |
| DIP84_WAC14057 | 1.00 | 0.00 | 0.00 | 0.00 | 0.00 | 0.00 | 0.00 | 0.00 |
| DIP85_WAC14058 | 1.00 | 0.00 | 0.00 | 0.00 | 0.00 | 0.00 | 0.00 | 0.00 |
| DIP86_WAC14059 | 1.00 | 0.00 | 0.00 | 0.00 | 0.00 | 0.00 | 0.00 | 0.00 |
| DIP87_WAC14060 | 1.00 | 0.00 | 0.00 | 0.00 | 0.00 | 0.00 | 0.00 | 0.00 |
| DIP88_WAC14061 | 1.00 | 0.00 | 0.00 | 0.00 | 0.00 | 0.00 | 0.00 | 0.00 |
| DIP89_WAC14062 | 1.00 | 0.00 | 0.00 | 0.00 | 0.00 | 0.00 | 0.00 | 0.00 |
| DIP9_WAC1199 | 0.00 | 0.00 | 0.00 | 0.00 | 0.00 | 1.00 | 0.00 | 0.00 |
| DIP90_WAC14066 | 1.00 | 0.00 | 0.00 | 0.00 | 0.00 | 0.00 | 0.00 | 0.00 |
| DIP91_WAC14067 | 1.00 | 0.00 | 0.00 | 0.00 | 0.00 | 0.00 | 0.00 | 0.00 |
| DIP92_WAC14068 | 1.00 | 0.00 | 0.00 | 0.00 | 0.00 | 0.00 | 0.00 | 0.00 |
| DIP93_WAC14137 | 0.89 | 0.00 | 0.11 | 0.00 | 0.00 | 0.00 | 0.00 | 0.00 |
| DIP94_WAC14138 | 1.00 | 0.00 | 0.00 | 0.00 | 0.00 | 0.00 | 0.00 | 0.00 |
| DIP96_WAC14140 | 1.00 | 0.00 | 0.00 | 0.00 | 0.00 | 0.00 | 0.00 | 0.00 |

| | | | | | | | | |
|---|---|---|---|---|---|---|---|---|
| DIP97_WAC14141 | 1.00 | 0.00 | 0.00 | 0.00 | 0.00 | 0.00 | 0.00 | 0.00 |
| O1_WAC2216 | 1.00 | 0.00 | 0.00 | 0.00 | 0.00 | 0.00 | 0.00 | 0.00 |
| O10_WAC8384 | 1.00 | 0.00 | 0.00 | 0.00 | 0.00 | 0.00 | 0.00 | 0.00 |
| O100_15FG102 | 1.00 | 0.00 | 0.00 | 0.00 | 0.00 | 0.00 | 0.00 | 0.00 |
| O101_15FG103 | 1.00 | 0.00 | 0.00 | 0.00 | 0.00 | 0.00 | 0.00 | 0.00 |
| O102_15FG104 | 1.00 | 0.00 | 0.00 | 0.00 | 0.00 | 0.00 | 0.00 | 0.00 |
| O103_15FG105 | 1.00 | 0.00 | 0.00 | 0.00 | 0.00 | 0.00 | 0.00 | 0.00 |
| O104_FG106 | 1.00 | 0.00 | 0.00 | 0.00 | 0.00 | 0.00 | 0.00 | 0.00 |
| O105_FG107 | 1.00 | 0.00 | 0.00 | 0.00 | 0.00 | 0.00 | 0.00 | 0.00 |
| O106_FG108 | 1.00 | 0.00 | 0.00 | 0.00 | 0.00 | 0.00 | 0.00 | 0.00 |
| O107_15FG107 | 1.00 | 0.00 | 0.00 | 0.00 | 0.00 | 0.00 | 0.00 | 0.00 |
| O108_15FG109 | 1.00 | 0.00 | 0.00 | 0.00 | 0.00 | 0.00 | 0.00 | 0.00 |
| O109_15FG110 | 1.00 | 0.00 | 0.00 | 0.00 | 0.00 | 0.00 | 0.00 | 0.00 |
| O11_WAC8390 | 1.00 | 0.00 | 0.00 | 0.00 | 0.00 | 0.00 | 0.00 | 0.00 |
| O110_15FG112 | 1.00 | 0.00 | 0.00 | 0.00 | 0.00 | 0.00 | 0.00 | 0.00 |
| O111_15FG113 | 1.00 | 0.00 | 0.00 | 0.00 | 0.00 | 0.00 | 0.00 | 0.00 |
| O112_15FG114 | 1.00 | 0.00 | 0.00 | 0.00 | 0.00 | 0.00 | 0.00 | 0.00 |
| O113_15FG115 | 1.00 | 0.00 | 0.00 | 0.00 | 0.00 | 0.00 | 0.00 | 0.00 |
| O114_15FG116 | 1.00 | 0.00 | 0.00 | 0.00 | 0.00 | 0.00 | 0.00 | 0.00 |
| O115_15FG117 | 1.00 | 0.00 | 0.00 | 0.00 | 0.00 | 0.00 | 0.00 | 0.00 |
| O116_15FG118 | 1.00 | 0.00 | 0.00 | 0.00 | 0.00 | 0.00 | 0.00 | 0.00 |
| O117_15FG119 | 1.00 | 0.00 | 0.00 | 0.00 | 0.00 | 0.00 | 0.00 | 0.00 |
| O118_15FG120 | 0.85 | 0.00 | 0.15 | 0.00 | 0.00 | 0.00 | 0.00 | 0.00 |
| O119_15FG04 | 1.00 | 0.00 | 0.00 | 0.00 | 0.00 | 0.00 | 0.00 | 0.00 |
| O12_WAC8410 | 1.00 | 0.00 | 0.00 | 0.00 | 0.00 | 0.00 | 0.00 | 0.00 |
| O120_15FG28 | 1.00 | 0.00 | 0.00 | 0.00 | 0.00 | 0.00 | 0.00 | 0.00 |
| O122_15FG33 | 1.00 | 0.00 | 0.00 | 0.00 | 0.00 | 0.00 | 0.00 | 0.00 |
| O123_15FG37 | 1.00 | 0.00 | 0.00 | 0.00 | 0.00 | 0.00 | 0.00 | 0.00 |
| O124_15FG38 | 1.00 | 0.00 | 0.00 | 0.00 | 0.00 | 0.00 | 0.00 | 0.00 |
| O127_15FG47 | 1.00 | 0.00 | 0.00 | 0.00 | 0.00 | 0.00 | 0.00 | 0.00 |
| O128_15FG49 | 1.00 | 0.00 | 0.00 | 0.00 | 0.00 | 0.00 | 0.00 | 0.00 |
| O129_15FG226 | 1.00 | 0.00 | 0.00 | 0.00 | 0.00 | 0.00 | 0.00 | 0.00 |
| O13_WAC8635 | 0.00 | 0.00 | 1.00 | 0.00 | 0.00 | 0.00 | 0.00 | 0.00 |
| O130_15FG229 | 1.00 | 0.00 | 0.00 | 0.00 | 0.00 | 0.00 | 0.00 | 0.00 |
| O131_15FG237 | 1.00 | 0.00 | 0.00 | 0.00 | 0.00 | 0.00 | 0.00 | 0.00 |
| O132_Northam_Mace | 1.00 | 0.00 | 0.00 | 0.00 | 0.00 | 0.00 | 0.00 | 0.00 |
| O133_Northam_Mage | 1.00 | 0.00 | 0.00 | 0.00 | 0.00 | 0.00 | 0.00 | 0.00 |
| O134_Northam_Emu1 | 1.00 | 0.00 | 0.00 | 0.00 | 0.00 | 0.00 | 0.00 | 0.00 |
| O135_Northam_Emu2 | 1.00 | 0.00 | 0.00 | 0.00 | 0.00 | 0.00 | 0.00 | 0.00 |
| O136_Northam_Mace | 1.00 | 0.00 | 0.00 | 0.00 | 0.00 | 0.00 | 0.00 | 0.00 |
| O14_WAC9178 | 0.00 | 0.00 | 1.00 | 0.00 | 0.00 | 0.00 | 0.00 | 0.00 |
| O140_FG_W003_5 | 1.00 | 0.00 | 0.00 | 0.00 | 0.00 | 0.00 | 0.00 | 0.00 |
| O141_Nor_RAC2182_ | 1.00 | 0.00 | 0.00 | 0.00 | 0.00 | 0.00 | 0.00 | 0.00 |
| O142_Northam_WGT | 1.00 | 0.00 | 0.00 | 0.00 | 0.00 | 0.00 | 0.00 | 0.00 |
| O143_Nor_RAC2182_ | 1.00 | 0.00 | 0.00 | 0.00 | 0.00 | 0.00 | 0.00 | 0.00 |
| O15_WAC13418 | 0.00 | 0.00 | 1.00 | 0.00 | 0.00 | 0.00 | 0.00 | 0.00 |
| O16_SN15 | 0.00 | 0.00 | 1.00 | 0.00 | 0.00 | 0.00 | 0.00 | 0.00 |
| O17_WAC13068 | 0.00 | 1.00 | 0.00 | 0.00 | 0.00 | 0.00 | 0.00 | 0.00 |
| O18_WAC13069 | 0.00 | 1.00 | 0.00 | 0.00 | 0.00 | 0.00 | 0.00 | 0.00 |

| | | | | | | | | |
|---|---|---|---|---|---|---|---|---|
| O19_WAC13070 | 0.00 | 0.00 | 0.00 | 1.00 | 0.00 | 0.00 | 0.00 | 0.00 |
| O2_WAC2285 | 1.00 | 0.00 | 0.00 | 0.00 | 0.00 | 0.00 | 0.00 | 0.00 |
| O20_WAC13071 | 1.00 | 0.00 | 0.00 | 0.00 | 0.00 | 0.00 | 0.00 | 0.00 |
| O21_WAC13072 | 1.00 | 0.00 | 0.00 | 0.00 | 0.00 | 0.00 | 0.00 | 0.00 |
| O22_WAC13073 | 1.00 | 0.00 | 0.00 | 0.00 | 0.00 | 0.00 | 0.00 | 0.00 |
| O23_WAC13074 | 0.00 | 0.00 | 0.00 | 1.00 | 0.00 | 0.00 | 0.00 | 0.00 |
| O24_WAC13075 | 0.00 | 0.00 | 0.00 | 1.00 | 0.00 | 0.00 | 0.00 | 0.00 |
| O25_WAC13076 | 0.00 | 0.00 | 0.00 | 1.00 | 0.00 | 0.00 | 0.00 | 0.00 |
| O26_WAC13077 | 1.00 | 0.00 | 0.00 | 0.00 | 0.00 | 0.00 | 0.00 | 0.00 |
| O27_Meck1 | 1.00 | 0.00 | 0.00 | 0.00 | 0.00 | 0.00 | 0.00 | 0.00 |
| O28_Meck3 | 1.00 | 0.00 | 0.00 | 0.00 | 0.00 | 0.00 | 0.00 | 0.00 |
| O29_Meck5 | 1.00 | 0.00 | 0.00 | 0.00 | 0.00 | 0.00 | 0.00 | 0.00 |
| O3_WAC2810 | 0.00 | 0.00 | 1.00 | 0.00 | 0.00 | 0.00 | 0.00 | 0.00 |
| O30_Meck6 | 0.98 | 0.00 | 0.02 | 0.00 | 0.00 | 0.00 | 0.00 | 0.00 |
| O31_Meck8 | 1.00 | 0.00 | 0.00 | 0.00 | 0.00 | 0.00 | 0.00 | 0.00 |
| O32_S1FT3A | 1.00 | 0.00 | 0.00 | 0.00 | 0.00 | 0.00 | 0.00 | 0.00 |
| O33_S4FT3A | 1.00 | 0.00 | 0.00 | 0.00 | 0.00 | 0.00 | 0.00 | 0.00 |
| O34_S1FT3B | 1.00 | 0.00 | 0.00 | 0.00 | 0.00 | 0.00 | 0.00 | 0.00 |
| O35_Gerald1 | 1.00 | 0.00 | 0.00 | 0.00 | 0.00 | 0.00 | 0.00 | 0.00 |
| O36_Gerald4 | 1.00 | 0.00 | 0.00 | 0.00 | 0.00 | 0.00 | 0.00 | 0.00 |
| O37_Mur_51 | 1.00 | 0.00 | 0.00 | 0.00 | 0.00 | 0.00 | 0.00 | 0.00 |
| O39_WAC13402 | 1.00 | 0.00 | 0.00 | 0.00 | 0.00 | 0.00 | 0.00 | 0.00 |
| O4_WAC2813 | 0.00 | 0.00 | 1.00 | 0.00 | 0.00 | 0.00 | 0.00 | 0.00 |
| O40_WAC13403 | 0.00 | 0.00 | 0.00 | 1.00 | 0.00 | 0.00 | 0.00 | 0.00 |
| O41_WAC13404 | 0.00 | 0.00 | 0.00 | 1.00 | 0.00 | 0.00 | 0.00 | 0.00 |
| O42_WAC13405 | 0.00 | 0.00 | 1.00 | 0.00 | 0.00 | 0.00 | 0.00 | 0.00 |
| O43_WAC13443 | 1.00 | 0.00 | 0.00 | 0.00 | 0.00 | 0.00 | 0.00 | 0.00 |
| O44_WAC13446 | 0.00 | 0.00 | 0.00 | 1.00 | 0.00 | 0.00 | 0.00 | 0.00 |
| O45_WAC13447 | 0.00 | 0.00 | 1.00 | 0.00 | 0.00 | 0.00 | 0.00 | 0.00 |
| O46_WAC13523 | 0.00 | 1.00 | 0.00 | 0.00 | 0.00 | 0.00 | 0.00 | 0.00 |
| O47_WAC13524 | 0.00 | 1.00 | 0.00 | 0.00 | 0.00 | 0.00 | 0.00 | 0.00 |
| O48_WAC13525 | 0.00 | 0.00 | 0.00 | 1.00 | 0.00 | 0.00 | 0.00 | 0.00 |
| O49_WAC13526 | 0.00 | 0.00 | 0.00 | 1.00 | 0.00 | 0.00 | 0.00 | 0.00 |
| O5_WAC4303 | 1.00 | 0.00 | 0.00 | 0.00 | 0.00 | 0.00 | 0.00 | 0.00 |
| O50_WAC13527 | 0.00 | 1.00 | 0.00 | 0.00 | 0.00 | 0.00 | 0.00 | 0.00 |
| O51_WAC13528 | 0.00 | 0.00 | 0.00 | 1.00 | 0.00 | 0.00 | 0.00 | 0.00 |
| O52_WAC13529 | 0.00 | 1.00 | 0.00 | 0.00 | 0.00 | 0.00 | 0.00 | 0.00 |
| O53_WAC13530 | 0.00 | 1.00 | 0.00 | 0.00 | 0.00 | 0.00 | 0.00 | 0.00 |
| O54_WAC13531 | 0.00 | 1.00 | 0.00 | 0.00 | 0.00 | 0.00 | 0.00 | 0.00 |
| O55_WAC13532 | 0.00 | 0.00 | 0.00 | 1.00 | 0.00 | 0.00 | 0.00 | 0.00 |
| O56_WAC13615 | 1.00 | 0.00 | 0.00 | 0.00 | 0.00 | 0.00 | 0.00 | 0.00 |
| O57_WAC13616 | 1.00 | 0.00 | 0.00 | 0.00 | 0.00 | 0.00 | 0.00 | 0.00 |
| O58_WAC13617 | 1.00 | 0.00 | 0.00 | 0.00 | 0.00 | 0.00 | 0.00 | 0.00 |
| O59_WAC13630 | 0.00 | 0.00 | 0.00 | 1.00 | 0.00 | 0.00 | 0.00 | 0.00 |
| O6_WAC4321 | 1.00 | 0.00 | 0.00 | 0.00 | 0.00 | 0.00 | 0.00 | 0.00 |
| O60_WAC13631 | 0.00 | 0.00 | 0.00 | 1.00 | 0.00 | 0.00 | 0.00 | 0.00 |
| O61_WAC13632 | 0.00 | 0.00 | 0.00 | 1.00 | 0.00 | 0.00 | 0.00 | 0.00 |
| O62_WAC13666 | 1.00 | 0.00 | 0.00 | 0.00 | 0.00 | 0.00 | 0.00 | 0.00 |
| O63_WAC13667 | 1.00 | 0.00 | 0.00 | 0.00 | 0.00 | 0.00 | 0.00 | 0.00 |

| | | | | | | | | |
|---|---|---|---|---|---|---|---|---|
| O64_WAC13690 | 1.00 | 0.00 | 0.00 | 0.00 | 0.00 | 0.00 | 0.00 | 0.00 |
| O65_WAC13691 | 1.00 | 0.00 | 0.00 | 0.00 | 0.00 | 0.00 | 0.00 | 0.00 |
| O66_206FG226 | 1.00 | 0.00 | 0.00 | 0.00 | 0.00 | 0.00 | 0.00 | 0.00 |
| O67_205FG215_1 | 1.00 | 0.00 | 0.00 | 0.00 | 0.00 | 0.00 | 0.00 | 0.00 |
| O68_201FG209 | 1.00 | 0.00 | 0.00 | 0.00 | 0.00 | 0.00 | 0.00 | 0.00 |
| O69_201FG211 | 1.00 | 0.00 | 0.00 | 0.00 | 0.00 | 0.00 | 0.00 | 0.00 |
| O7_WAC4319 | 0.00 | 1.00 | 0.00 | 0.00 | 0.00 | 0.00 | 0.00 | 0.00 |
| O70_202FG212 | 1.00 | 0.00 | 0.00 | 0.00 | 0.00 | 0.00 | 0.00 | 0.00 |
| O71_204FG221 | 1.00 | 0.00 | 0.00 | 0.00 | 0.00 | 0.00 | 0.00 | 0.00 |
| O72_204FG223 | 1.00 | 0.00 | 0.00 | 0.00 | 0.00 | 0.00 | 0.00 | 0.00 |
| O73_205FG225 | 1.00 | 0.00 | 0.00 | 0.00 | 0.00 | 0.00 | 0.00 | 0.00 |
| O74_206FG227 | 1.00 | 0.00 | 0.00 | 0.00 | 0.00 | 0.00 | 0.00 | 0.00 |
| O75_205FG216 | 1.00 | 0.00 | 0.00 | 0.00 | 0.00 | 0.00 | 0.00 | 0.00 |
| O76_201FG219 | 1.00 | 0.00 | 0.00 | 0.00 | 0.00 | 0.00 | 0.00 | 0.00 |
| O77_201FG218 | 1.00 | 0.00 | 0.00 | 0.00 | 0.00 | 0.00 | 0.00 | 0.00 |
| O78_204FG214 | 1.00 | 0.00 | 0.00 | 0.00 | 0.00 | 0.00 | 0.00 | 0.00 |
| O79_903FG214 | 1.00 | 0.00 | 0.00 | 0.00 | 0.00 | 0.00 | 0.00 | 0.00 |
| O8_WAC4808 | 1.00 | 0.00 | 0.00 | 0.00 | 0.00 | 0.00 | 0.00 | 0.00 |
| O80_201FG49 | 1.00 | 0.00 | 0.00 | 0.00 | 0.00 | 0.00 | 0.00 | 0.00 |
| O81_203FG58 | 1.00 | 0.00 | 0.00 | 0.00 | 0.00 | 0.00 | 0.00 | 0.00 |
| O82_205FG63 | 1.00 | 0.00 | 0.00 | 0.00 | 0.00 | 0.00 | 0.00 | 0.00 |
| O83_206FG66 | 1.00 | 0.00 | 0.00 | 0.00 | 0.00 | 0.00 | 0.00 | 0.00 |
| O84_206FG67 | 1.00 | 0.00 | 0.00 | 0.00 | 0.00 | 0.00 | 0.00 | 0.00 |
| O85_206FG68 | 1.00 | 0.00 | 0.00 | 0.00 | 0.00 | 0.00 | 0.00 | 0.00 |
| O86_205FG142 | 1.00 | 0.00 | 0.00 | 0.00 | 0.00 | 0.00 | 0.00 | 0.00 |
| O87_53FG143_1 | 1.00 | 0.00 | 0.00 | 0.00 | 0.00 | 0.00 | 0.00 | 0.00 |
| O88_201FG208 | 1.00 | 0.00 | 0.00 | 0.00 | 0.00 | 0.00 | 0.00 | 0.00 |
| O89_203FG213 | 1.00 | 0.00 | 0.00 | 0.00 | 0.00 | 0.00 | 0.00 | 0.00 |
| O9_WAC4648 | 0.00 | 1.00 | 0.00 | 0.00 | 0.00 | 0.00 | 0.00 | 0.00 |
| O91_204FG222 | 1.00 | 0.00 | 0.00 | 0.00 | 0.00 | 0.00 | 0.00 | 0.00 |
| O92_205FG410 | 1.00 | 0.00 | 0.00 | 0.00 | 0.00 | 0.00 | 0.00 | 0.00 |
| O93_202FG414 | 1.00 | 0.00 | 0.00 | 0.00 | 0.00 | 0.00 | 0.00 | 0.00 |
| O94_WAC739 | 1.00 | 0.00 | 0.00 | 0.00 | 0.00 | 0.00 | 0.00 | 0.00 |
| O95_WAC740 | 1.00 | 0.00 | 0.00 | 0.00 | 0.00 | 0.00 | 0.00 | 0.00 |
| O96_WAC741 | 1.00 | 0.00 | 0.00 | 0.00 | 0.00 | 0.00 | 0.00 | 0.00 |
| O97_15FG99 | 1.00 | 0.00 | 0.00 | 0.00 | 0.00 | 0.00 | 0.00 | 0.00 |
| O98_15FG100 | 1.00 | 0.00 | 0.00 | 0.00 | 0.00 | 0.00 | 0.00 | 0.00 |
| O99_15FG101 | 1.00 | 0.00 | 0.00 | 0.00 | 0.00 | 0.00 | 0.00 | 0.00 |
| O143_16FG06 | 1.00 | 0.00 | 0.00 | 0.00 | 0.00 | 0.00 | 0.00 | 0.00 |
| O144_16FG158 | 1.00 | 0.00 | 0.00 | 0.00 | 0.00 | 0.00 | 0.00 | 0.00 |
| O145_16FG159 | 1.00 | 0.00 | 0.00 | 0.00 | 0.00 | 0.00 | 0.00 | 0.00 |
| O146_16FG160 | 0.00 | 0.00 | 0.00 | 0.00 | 1.00 | 0.00 | 0.00 | 0.00 |
| O147_16FG161 | 0.00 | 0.00 | 0.00 | 0.00 | 1.00 | 0.00 | 0.00 | 0.00 |
| O148_16FG162 | 0.00 | 0.00 | 0.00 | 0.00 | 1.00 | 0.00 | 0.00 | 0.00 |
| O150_16FG163_2 | 0.00 | 0.00 | 0.00 | 0.00 | 1.00 | 0.00 | 0.00 | 0.00 |
| O151_16FG164 | 0.00 | 0.00 | 0.00 | 0.00 | 1.00 | 0.00 | 0.00 | 0.00 |
| O152_16FG165 | 0.00 | 0.00 | 0.00 | 0.00 | 1.00 | 0.00 | 0.00 | 0.00 |
| O153_16FG166 | 0.00 | 0.00 | 0.00 | 0.00 | 1.00 | 0.00 | 0.00 | 0.00 |
| O154_16FG167 | 0.00 | 0.00 | 0.00 | 0.00 | 1.00 | 0.00 | 0.00 | 0.00 |

| | | | | | | | | |
|---|---|---|---|---|---|---|---|---|
| O155_16FG168 | 0.00 | 0.00 | 0.00 | 0.00 | 1.00 | 0.00 | 0.00 | 0.00 |
| O156_16FG169 | 0.00 | 0.00 | 0.00 | 0.00 | 1.00 | 0.00 | 0.00 | 0.00 |
| O157_WAC13955 | 1.00 | 0.00 | 0.00 | 0.00 | 0.00 | 0.00 | 0.00 | 0.00 |
| O158_16FG170 | 0.00 | 0.00 | 0.00 | 0.00 | 1.00 | 0.00 | 0.00 | 0.00 |
| O159_16FG171 | 0.00 | 0.00 | 0.00 | 0.00 | 1.00 | 0.00 | 0.00 | 0.00 |
| C260_PN260 | 1.00 | 0.00 | 0.00 | 0.00 | 0.00 | 0.00 | 0.00 | 0.00 |
| C261_PN261 | 1.00 | 0.00 | 0.00 | 0.00 | 0.00 | 0.00 | 0.00 | 0.00 |
| C262_PN262 | 1.00 | 0.00 | 0.00 | 0.00 | 0.00 | 0.00 | 0.00 | 0.00 |
| C263_PN263 | 1.00 | 0.00 | 0.00 | 0.00 | 0.00 | 0.00 | 0.00 | 0.00 |
| C264_PN264 | 1.00 | 0.00 | 0.00 | 0.00 | 0.00 | 0.00 | 0.00 | 0.00 |
| C265_PN265 | 1.00 | 0.00 | 0.00 | 0.00 | 0.00 | 0.00 | 0.00 | 0.00 |
| C266_PN266 | 1.00 | 0.00 | 0.00 | 0.00 | 0.00 | 0.00 | 0.00 | 0.00 |
| C268_PN268 | 1.00 | 0.00 | 0.00 | 0.00 | 0.00 | 0.00 | 0.00 | 0.00 |
| C269_PN269 | 1.00 | 0.00 | 0.00 | 0.00 | 0.00 | 0.00 | 0.00 | 0.00 |
| C270_PN270 | 1.00 | 0.00 | 0.00 | 0.00 | 0.00 | 0.00 | 0.00 | 0.00 |
| C271_PN271 | 1.00 | 0.00 | 0.00 | 0.00 | 0.00 | 0.00 | 0.00 | 0.00 |
| C274_PN274 | 1.00 | 0.00 | 0.00 | 0.00 | 0.00 | 0.00 | 0.00 | 0.00 |
| C275_PN275 | 0.96 | 0.00 | 0.04 | 0.00 | 0.00 | 0.00 | 0.00 | 0.00 |
| C276_PN276 | 1.00 | 0.00 | 0.00 | 0.00 | 0.00 | 0.00 | 0.00 | 0.00 |
| C277_PN277 | 1.00 | 0.00 | 0.00 | 0.00 | 0.00 | 0.00 | 0.00 | 0.00 |
| C279_PN279 | 1.00 | 0.00 | 0.00 | 0.00 | 0.00 | 0.00 | 0.00 | 0.00 |
| C280_PN280 | 0.00 | 0.00 | 0.00 | 0.00 | 0.00 | 0.00 | 1.00 | 0.00 |
| C282_PN282 | 0.00 | 0.00 | 0.00 | 0.00 | 0.00 | 0.00 | 1.00 | 0.00 |
| C283_PN283 | 0.00 | 0.00 | 0.00 | 0.00 | 0.00 | 0.00 | 1.00 | 0.00 |
| C284_PN284 | 1.00 | 0.00 | 0.00 | 0.00 | 0.00 | 0.00 | 0.00 | 0.00 |
| C285_PN285 | 1.00 | 0.00 | 0.00 | 0.00 | 0.00 | 0.00 | 0.00 | 0.00 |
| C286_PN286 | 1.00 | 0.00 | 0.00 | 0.00 | 0.00 | 0.00 | 0.00 | 0.00 |
| C287_PN287 | 0.93 | 0.00 | 0.07 | 0.00 | 0.00 | 0.00 | 0.00 | 0.00 |
| C288_PN288 | 1.00 | 0.00 | 0.00 | 0.00 | 0.00 | 0.00 | 0.00 | 0.00 |
| C289_PN289 | 1.00 | 0.00 | 0.00 | 0.00 | 0.00 | 0.00 | 0.00 | 0.00 |
| C290_PN290 | 1.00 | 0.00 | 0.00 | 0.00 | 0.00 | 0.00 | 0.00 | 0.00 |
| C291_PN291 | 1.00 | 0.00 | 0.00 | 0.00 | 0.00 | 0.00 | 0.00 | 0.00 |
| C292_PN292 | 1.00 | 0.00 | 0.00 | 0.00 | 0.00 | 0.00 | 0.00 | 0.00 |
| C295_PN295 | 1.00 | 0.00 | 0.00 | 0.00 | 0.00 | 0.00 | 0.00 | 0.00 |
| C296_PN296 | 0.00 | 0.00 | 0.00 | 0.00 | 0.00 | 0.00 | 0.00 | 1.00 |
| C297_PN297 | 1.00 | 0.00 | 0.00 | 0.00 | 0.00 | 0.00 | 0.00 | 0.00 |
| C298_PN298 | 0.00 | 0.00 | 0.00 | 0.00 | 0.00 | 0.00 | 0.00 | 1.00 |
| C299_PN299 | 1.00 | 0.00 | 0.00 | 0.00 | 0.00 | 0.00 | 0.00 | 0.00 |
| C300_PN300 | 1.00 | 0.00 | 0.00 | 0.00 | 0.00 | 0.00 | 0.00 | 0.00 |
| C301_PN301 | 1.00 | 0.00 | 0.00 | 0.00 | 0.00 | 0.00 | 0.00 | 0.00 |
| C302_PN302 | 1.00 | 0.00 | 0.00 | 0.00 | 0.00 | 0.00 | 0.00 | 0.00 |
| C303_PN303 | 1.00 | 0.00 | 0.00 | 0.00 | 0.00 | 0.00 | 0.00 | 0.00 |
| C304_PN304 | 1.00 | 0.00 | 0.00 | 0.00 | 0.00 | 0.00 | 0.00 | 0.00 |
| C305_PN305 | 1.00 | 0.00 | 0.00 | 0.00 | 0.00 | 0.00 | 0.00 | 0.00 |
| C306_PN306 | 0.00 | 0.00 | 0.00 | 0.00 | 0.00 | 0.00 | 1.00 | 0.00 |
| C307_PN307 | 0.00 | 0.00 | 0.00 | 0.00 | 0.00 | 0.00 | 1.00 | 0.00 |
| C308_PN308 | 1.00 | 0.00 | 0.00 | 0.00 | 0.00 | 0.00 | 0.00 | 0.00 |
| C309_PN309 | 1.00 | 0.00 | 0.00 | 0.00 | 0.00 | 0.00 | 0.00 | 0.00 |
| C312_PN312 | 1.00 | 0.00 | 0.00 | 0.00 | 0.00 | 0.00 | 0.00 | 0.00 |

| | | | | | | | | |
|---|---|---|---|---|---|---|---|---|
| C313_PN313 | 1.00 | 0.00 | 0.00 | 0.00 | 0.00 | 0.00 | 0.00 | 0.00 |
| C314_PN314 | 1.00 | 0.00 | 0.00 | 0.00 | 0.00 | 0.00 | 0.00 | 0.00 |
| C315_PN315 | 0.00 | 0.00 | 0.00 | 0.00 | 0.00 | 0.00 | 1.00 | 0.00 |
| C316_PN316 | 0.00 | 0.00 | 0.00 | 0.00 | 0.00 | 0.00 | 1.00 | 0.00 |
| C317_PN317 | 1.00 | 0.00 | 0.00 | 0.00 | 0.00 | 0.00 | 0.00 | 0.00 |
| C318_PN318 | 1.00 | 0.00 | 0.00 | 0.00 | 0.00 | 0.00 | 0.00 | 0.00 |
| C319_PN319 | 0.20 | 0.00 | 0.80 | 0.00 | 0.00 | 0.00 | 0.00 | 0.00 |
| C320_PN320 | 1.00 | 0.00 | 0.00 | 0.00 | 0.00 | 0.00 | 0.00 | 0.00 |
| C321_PN321 | 0.00 | 0.00 | 0.00 | 0.00 | 0.00 | 0.00 | 0.00 | 1.00 |
| C322_PN322 | 0.00 | 0.00 | 0.00 | 0.00 | 0.00 | 0.00 | 0.00 | 1.00 |
| C323_PN323 | 1.00 | 0.00 | 0.00 | 0.00 | 0.00 | 0.00 | 0.00 | 0.00 |
| C324_PN324 | 1.00 | 0.00 | 0.00 | 0.00 | 0.00 | 0.00 | 0.00 | 0.00 |
| C325_PN325 | 1.00 | 0.00 | 0.00 | 0.00 | 0.00 | 0.00 | 0.00 | 0.00 |
| C326_PN326 | 1.00 | 0.00 | 0.00 | 0.00 | 0.00 | 0.00 | 0.00 | 0.00 |
| C328_PN328 | 1.00 | 0.00 | 0.00 | 0.00 | 0.00 | 0.00 | 0.00 | 0.00 |
| C329_PN329 | 1.00 | 0.00 | 0.00 | 0.00 | 0.00 | 0.00 | 0.00 | 0.00 |
| C330_PN330 | 1.00 | 0.00 | 0.00 | 0.00 | 0.00 | 0.00 | 0.00 | 0.00 |
| C331_PN331 | 0.99 | 0.00 | 0.01 | 0.00 | 0.00 | 0.00 | 0.00 | 0.00 |
| C332_PN332 | 1.00 | 0.00 | 0.00 | 0.00 | 0.00 | 0.00 | 0.00 | 0.00 |
| C333_PN333 | 0.00 | 0.00 | 0.00 | 0.00 | 0.00 | 0.00 | 0.00 | 1.00 |
| C334_PN334 | 0.00 | 0.00 | 0.00 | 0.00 | 0.00 | 0.00 | 1.00 | 0.00 |
| C335_PN335 | 0.00 | 0.00 | 0.00 | 0.00 | 0.00 | 0.00 | 1.00 | 0.00 |
| C336_PN336 | 1.00 | 0.00 | 0.00 | 0.00 | 0.00 | 0.00 | 0.00 | 0.00 |
| C337_PN337 | 0.00 | 0.00 | 0.00 | 0.00 | 0.00 | 0.00 | 1.00 | 0.00 |
| C338_PN338 | 0.00 | 0.00 | 0.00 | 0.00 | 0.00 | 0.00 | 1.00 | 0.00 |
| C340_PN340 | 0.00 | 0.00 | 0.00 | 0.00 | 0.00 | 0.00 | 1.00 | 0.00 |
| C341_PN341 | 0.00 | 0.00 | 0.00 | 0.00 | 0.00 | 0.00 | 0.00 | 1.00 |
| C342_PN342 | 1.00 | 0.00 | 0.00 | 0.00 | 0.00 | 0.00 | 0.00 | 0.00 |
| C343_PN343 | 0.00 | 0.00 | 0.00 | 0.00 | 0.00 | 0.00 | 1.00 | 0.00 |
| C344_PN344 | 0.00 | 0.00 | 0.00 | 0.00 | 0.00 | 0.00 | 0.00 | 1.00 |
| C345_PN345 | 0.00 | 0.00 | 0.00 | 0.00 | 0.00 | 0.00 | 1.00 | 0.00 |
| C346_PN346 | 0.00 | 0.00 | 0.00 | 0.00 | 0.00 | 0.00 | 1.00 | 0.00 |
| C347_PN347 | 1.00 | 0.00 | 0.00 | 0.00 | 0.00 | 0.00 | 0.00 | 0.00 |
| C348_PN348 | 1.00 | 0.00 | 0.00 | 0.00 | 0.00 | 0.00 | 0.00 | 0.00 |
| C349_PN349 | 1.00 | 0.00 | 0.00 | 0.00 | 0.00 | 0.00 | 0.00 | 0.00 |
| C350_PN350 | 0.00 | 0.00 | 0.00 | 0.00 | 0.00 | 0.00 | 0.00 | 1.00 |
| C351_PN351 | 1.00 | 0.00 | 0.00 | 0.00 | 0.00 | 0.00 | 0.00 | 0.00 |
| C352_PN352 | 1.00 | 0.00 | 0.00 | 0.00 | 0.00 | 0.00 | 0.00 | 0.00 |
| C353_PN353 | 0.00 | 0.00 | 0.00 | 0.00 | 0.00 | 0.00 | 0.00 | 1.00 |
| C354_PN354 | 1.00 | 0.00 | 0.00 | 0.00 | 0.00 | 0.00 | 0.00 | 0.00 |
| C355_PN355 | 0.00 | 0.00 | 0.00 | 0.00 | 0.00 | 0.00 | 0.00 | 1.00 |
| C356_PN356 | 0.00 | 0.00 | 0.00 | 0.00 | 0.00 | 0.00 | 0.00 | 1.00 |
| C357_PN357 | 1.00 | 0.00 | 0.00 | 0.00 | 0.00 | 0.00 | 0.00 | 0.00 |
| C358_PN358 | 1.00 | 0.00 | 0.00 | 0.00 | 0.00 | 0.00 | 0.00 | 0.00 |
| C359_PN359 | 0.00 | 0.00 | 0.00 | 0.00 | 0.00 | 0.00 | 0.00 | 1.00 |
| C360_PN360 | 1.00 | 0.00 | 0.00 | 0.00 | 0.00 | 0.00 | 0.00 | 0.00 |
| C361_PN361 | 1.00 | 0.00 | 0.00 | 0.00 | 0.00 | 0.00 | 0.00 | 0.00 |
| C362_PN362 | 1.00 | 0.00 | 0.00 | 0.00 | 0.00 | 0.00 | 0.00 | 0.00 |
| C363_PN363 | 1.00 | 0.00 | 0.00 | 0.00 | 0.00 | 0.00 | 0.00 | 0.00 |

| | | | | | | | | |
|---|---|---|---|---|---|---|---|---|
| C364_PN364 | 1.00 | 0.00 | 0.00 | 0.00 | 0.00 | 0.00 | 0.00 | 0.00 |
| C365_PN365 | 0.00 | 0.00 | 0.00 | 0.00 | 0.00 | 0.00 | 0.00 | 1.00 |
| C366_PN366 | 1.00 | 0.00 | 0.00 | 0.00 | 0.00 | 0.00 | 0.00 | 0.00 |
| C367_PN367 | 1.00 | 0.00 | 0.00 | 0.00 | 0.00 | 0.00 | 0.00 | 0.00 |
| C368_PN368 | 0.00 | 0.00 | 0.00 | 0.00 | 0.00 | 0.00 | 0.00 | 1.00 |
| C369_PN369 | 1.00 | 0.00 | 0.00 | 0.00 | 0.00 | 0.00 | 0.00 | 0.00 |
| C370_PN370 | 0.00 | 0.00 | 0.00 | 0.00 | 0.00 | 0.00 | 0.00 | 1.00 |
| C371_PN371 | 1.00 | 0.00 | 0.00 | 0.00 | 0.00 | 0.00 | 0.00 | 0.00 |
| C372_PN372 | 1.00 | 0.00 | 0.00 | 0.00 | 0.00 | 0.00 | 0.00 | 0.00 |
| C373_PN373 | 0.00 | 0.00 | 0.00 | 0.00 | 0.00 | 0.00 | 0.00 | 1.00 |
| C374_PN374 | 0.00 | 0.00 | 0.00 | 0.00 | 0.00 | 0.00 | 0.00 | 1.00 |
| C375_PN375 | 1.00 | 0.00 | 0.00 | 0.00 | 0.00 | 0.00 | 0.00 | 0.00 |
| C376_PN376 | 1.00 | 0.00 | 0.00 | 0.00 | 0.00 | 0.00 | 0.00 | 0.00 |
| C377_PN377 | 1.00 | 0.00 | 0.00 | 0.00 | 0.00 | 0.00 | 0.00 | 0.00 |
| C378_PN378 | 0.00 | 0.00 | 0.00 | 0.00 | 0.00 | 0.00 | 0.00 | 1.00 |
| C379_PN379 | 0.00 | 0.00 | 0.00 | 0.00 | 0.00 | 0.00 | 0.00 | 1.00 |
| C380_PN380 | 1.00 | 0.00 | 0.00 | 0.00 | 0.00 | 0.00 | 0.00 | 0.00 |
| O55_Mur_S3 | 1.00 | 0.00 | 0.00 | 0.00 | 0.00 | 0.00 | 0.00 | 0.00 |
| O66_14FG141 | 1.00 | 0.00 | 0.00 | 0.00 | 0.00 | 0.00 | 0.00 | 0.00 |

**STable 6**: Nanopore reference genomes statistics

| Attribute | 15FG38 | 16FG168 | PN315 |
|---|---|---|---|
| Total sequenced bases | 2,183,637,067 | 5,646,704,864 | 4,204,825,646 |
| Nuclear scaffolds | 23 | 24 | 24 |
| Nuclear scaffolds with both telomeres | 17 | 19 | 14 |
| Nuclear scafflods with one telomeres | 6 | 5 | 10 |
| Nuclear genome (Mbp) | 37.7 | 37.6 | 37.8 |
| Mitochondrial genome (bp) | 108,867 | ND[c] | 103,075 |
| L50 (kbp)[a] | 1.79E+03 | 1.68E+03 | 1.70E+03 |
| N50 (contigs)b | 9 | 9 | 9 |
| Genome BUSCOs (%) | 99 | 98.9 | 99 |
| Gene number | 18,728 | 18,953 | 18,747 |
| Percentage RIP affected area | 7.99 | 7.35 | 7.01 |
| Region R0 (0 - 30% GC content) | | | |
| Proportion_of_genome: | 6.98 | 5.95 | 5.96 |
| Number_of_regions | 175 | 188 | 163 |
| Average_length_kbp | 15 | 11.9 | 13.8 |
| St_dev_length_kbp | 15.9 | 13.3 | 16.1 |
| Number_of_genes | 285 | 158 | 229 |
| Gene_density_(genes_per_Mbp) | 108 | 70.5 | 102 |
| Region R1 (30 - 50% GC content) | | | |
| Proportion_of_genome: | 17.4 | 19.1 | 17.9 |
| Number_of_regions | 854 | 908 | 878 |
| Average_length_kbp | 7.7 | 7.9 | 7.71 |
| St_dev_length_kbp | 7.67 | 7.79 | 8.78 |
| Number_of_genes | 3,096 | 3,330 | 3,039 |
| Gene_density_(genes_per_Mbp) | 471 | 464 | 449 |
| Region R2 (50 - 70% GC content) | | | |
| Proportion_of_genome: | 75.6 | 75 | 76.1 |
| Number_of_regions | 813 | 832 | 834 |
| Average_length_kbp | 35 | 33.9 | 34.5 |
| St_dev_length_kbp | 68.3 | 55.9 | 66.6 |
| Number_of_genes | 15,218 | 15,201 | 15,333 |
| Gene_density_(genes_per_Mbp) | 534 | 539 | 532 |

[a]Length of the smallest contig in an ordered set of contigs corresponding to 50% of the assembly length.

[b]Smallest number of contigs whose length equals 50% of the genome assembly.

[c]ND = Not Detected

**STable 7**: Molly meta data in three reference genomes

| Isolate | # | Chromosome/contig | Modification | size (bp) | LRAR | Start | End | Gene predicted in Molly | Distance_to_the_closest_gene | Gene_id | Promoter | Gene annotation |
|---|---|---|---|---|---|---|---|---|---|---|---|---|
| 15FG38 | 1 | Chromosome_02\|15FG38_ctg_02C | intact/weak | 1866 | y | 164,659 | 166,524 | Putative transposase | 1534 | SNOG_021890 | na | dipeptidase |
| 15FG38 | 2 | Chromosome_02\|15FG38_ctg_02C | intact/weak | 1866 | y | 176,703 | 174,838 | Putative transposase | 471 | SNOG_019210 | y | hypothetical protein |
| 15FG38 | 3 | Chromosome_02\|15FG38_ctg_02C | intact/weak | 1866 | n | 220,143 | 218,278 | Putative transposase | 6261 | SNOG_432590 | na | hypothetical protein |
| 15FG38 | 4 | Chromosome_02\|15FG38_ctg_02C | intact/weak | 1866 | y | 818,767 | 820,632 | Putative transposase | 5516 | SNOG_019120 | na | hypothetical protein |
| 15FG38 | 5 | Chromosome_08\|15FG38_ctg_05C | strong | 1866 | n | 630,774 | 628,909 | Putative transposase | 96 | SNOG_041800 | y | fungal-specific transcription factor domain-containing protein |
| 15FG38 | 6 | Chromosome_09\|15FG38_ctg_08C | strong | 1866 | y | 1,288,742 | 1,290,607 | hypothetical protein HBI01_066130 | 5329 | SNOG_432530 | na | Retrovirushypothetical proteinrelated Pol polyprotein from transposon TNT 1hypothetical protein94 |
| 15FG38 | 7 | Chromosome_09\|15FG38_ctg_08C | strong | 1866 | y | 1,306,180 | 1,304,315 | Putative transposase | 130 | SNOG_441370 | y | hypothetical protein |
| 15FG38 | 8 | Chromosome_09\|15FG38_ctg_08C | strong | 1866 | y | 1,327,503 | 1,325,638 | no | 0 | KAH6023318 | y | hypothetical protein |
| 15FG38 | 9 | Chromosome_11\|15FG38_ctg_10C | strong | 1866 | y | 1,088,107 | 1,086,242 | Putative transposase | 2818 | SNOG_164900 | na | MYND-type zinc finger protein samB |
| 15FG38 | 10 | Chromosome_12\|15FG38_ctg_11C | intact/weak | 1866 | y | 331,100 | 332,965 | Putative transposase | 990 | SNOG_439000 | y | reverse transcriptase domain protein, partial |
| 15FG38 | 11 | Chromosome_12\|15FG38_ctg_11C | intact/weak | 1866 | y | 341,196 | 343,061 | Putative transposase | 799 | SNOG_448380 | y | hypothetical protein |
| 15FG38 | 12 | Chromosome_18\|15FG38_ctg_13+ | strong | 1866 | y | 895,343 | 897,208 | Putative transposase | 173 | SNOG_429580 | y | putative transposase |
| 15FG38 | 13 | Chromosome_18\|15FG38_ctg_13+ | strong | 1866 | y | 956,723 | 954,858 | Putative transposase | 8 | SNOG_429580 | y | hypothetical protein |
| 15FG38 | 14 | Chromosome_18\|15FG38_ctg_13+ | intact/weak | 1866 | y | 1,008,877 | 1,007,012 | Putative transposase | 3378 | SNOG_429580 | na | putative transposase |
| 15FG38 | 15 | Chromosome_18\|15FG38_ctg_13+ | intact/weak | 1866 | y | 1,299,001 | 1,297,136 | Putative transposase | 8677 | SNOG_425430 | na | putative transposase |
| 15FG38 | 16 | Chromosome_18\|15FG38_ctg_13+ | intact/weak | 1866 | y | 1,382,558 | 1,384,423 | Putative transposase | 2841 | SNOG_309300 | na | chitinase |
| 15FG38 | 17 | Chromosome_03\|15FG38_ctg_14C | strong | 1866 | n | 571,175 | 569,310 | Putative transposase | 0 | SNOG_036030 | y | YT521-B splicing factor |
| 15FG38 | 18 | Chromosome_15\|15FG38_ctg_15+ | intact/weak | 1866 | n | 1,336,389 | 1,334,524 | Putative transposase | 1442 | KAH4395491 | n | hypothetical protein |
| 15FG38 | 19 | Chromosome_22\|15FG38_ctg_20C | strong | 1866 | y | 109,076 | 107,211 | Putative transposase | 4676 | SNOG_123090 | na | Piwi-domain-containing protein |
| 16FG168 | 1 | Chromosome_01\|16FG168_ctg_01C | strong | 1866 | n | 6,883 | 8,748 | no | 1718 | SNOG_401220 | na | hypothetical protein |
| 16FG168 | 2 | Chromosome_01\|16FG168_ctg_01C | intact/weak | 1866 | n | 1,107,529 | 1,109,394 | Putative transposase | 263 | SNOG_401210 | n | hypothetical protein |
| 16FG168 | 3 | Chromosome_01\|16FG168_ctg_01C | strong | 1866 | n | 2,841,995 | 2,843,860 | Putative transposase | 1405 | SNOG_000020 | y | Six-hairpin glycosidase-like protein |
| 16FG168 | 4 | Chromosome_01\|16FG168_ctg_01C | intact/weak | 1866 | n | 2,844,181 | 2,846,046 | Putative transposase | 1445 | SNOG_427210 | y | hypothetical protein |
| 16FG168 | 5 | Chromosome_02\|16FG168_ctg_02+ | intact/weak | 1866 | n | 865,604 | 867,469 | Putative transposase | 151 | SNOG_022140 | y | glycoside hydrolase family 18 protein, partial |
| 16FG168 | 6 | Chromosome_02\|16FG168_ctg_02+ | strong | 1866 | y | 1,084,946 | 1,083,081 | Putative transposase | 987 | SNOG_023250 | y | AP2 domain-containing protein |
| 16FG168 | 7 | Chromosome_02\|16FG168_ctg_02+ | strong | 1866 | y | 1,176,186 | 1,178,051 | no | 547 | SNOG_427900 | n | hypothetical protein |
| 16FG168 | 8 | Chromosome_06\|16FG168_ctg_03C | strong | 1866 | y | 13,054 | 11,189 | no | 989 | SNOG_425430 | n | putative transposase |
| 16FG168 | 9 | Chromosome_06\|16FG168_ctg_03C | strong | 1866 | y | 409,028 | 407,163 | Putative transposase | 648 | SNOG_056370 | y | hypothetical protein |
| 16FG168 | 10 | Chromosome_06\|16FG168_ctg_03C | strong | 1866 | y | 771,877 | 773,742 | Putative transposase | 130 | SNOG_425430 | na | putative transposase |
| 16FG168 | 11 | Chromosome_05\|16FG168_ctg_04C | intact/weak | 1866 | y | 366,053 | 367,918 | Putative transposase | 905 | SNOG_407900 | n | HSP20-like chaperone |
| 16FG168 | 12 | Chromosome_05\|16FG168_ctg_04C | intact/weak | 1866 | y | 2,218,882 | 2,217,017 | Putative transposase | 532 | SNOG_442950 | y | hypothetical protein |
| 16FG168 | 13 | Chromosome_08\|16FG168_ctg_05+ | intact/weak | 1866 | n | 844,815 | 842,950 | Putative transposase | 1643 | SNOG_042790 | na | hypothetical protein |
| 16FG168 | 14 | Chromosome_04\|16FG168_ctg_06C | strong | 1866 | y | 376,055 | 374,190 | no | 174 | SNOG_075920 | y | DUF262 multi-domain protein |
| 16FG168 | 15 | Chromosome_04\|16FG168_ctg_06C | strong | 1866 | y | 537,215 | 539,080 | hypothetical protein HBI01_066130 | 1295 | SNOG_439000 | y | reverse transcriptase domain protein, partial |
| 16FG168 | 16 | Chromosome_04\|16FG168_ctg_06C | strong | 1866 | y | 1,331,855 | 1,329,990 | Putative transposase | 236 | SNOG_445480 | n | hypothetical protein |
| 16FG168 | 17 | Chromosome_09\|16FG168_ctg_07C | intact/weak | 1866 | y | 1,586,672 | 1,588,537 | Putative transposase | 602 | SNOG_428620 | n | hypothetical protein |
| 16FG168 | 18 | Chromosome_07\|16FG168_ctg_08C | intact/weak | 1866 | y | 661,800 | 663,664 | Putative transposase | 544 | SNOG_448320 | y | hypothetical protein |
| 16FG168 | 19 | Chromosome_12\|16FG168_ctg_10C | strong | 1866 | y | 265,811 | 263,946 | no | 6 | SNOG_443270 | y | no hit |
| 16FG168 | 20 | Chromosome_12\|16FG168_ctg_10C | strong | 1866 | y | 302,268 | 300,403 | Putative transposase | 456 | SNOG_425420 | y | hypothetical protein |
| 16FG168 | 21 | Chromosome_12\|16FG168_ctg_10C | strong | 1866 | y | 312,059 | 310,194 | no | 1554 | SNOG_442450 | na | hypothetical protein |
| 16FG168 | 22 | Chromosome_12\|16FG168_ctg_10C | strong | 1866 | y | 313,943 | 312,078 | no | 524 | SNOG_439000 | n | reverse transcriptase domain protein, partial |
| 16FG168 | 23 | Chromosome_12\|16FG168_ctg_10C | strong | 1866 | y | 338,324 | 340,189 | Putative transposase | 1233 | SNOG_439040 | n | hypothetical protein |
| 16FG168 | 24 | Chromosome_12\|16FG168_ctg_10C | strong | 1866 | y | 1,100,765 | 1,098,900 | Putative transposase | 2349 | SNOG_439030 | y | hypothetical protein |
| 16FG168 | 25 | Chromosome_11\|16FG168_ctg_11+ | strong | 1866 | y | 1,170,408 | 1,168,543 | Putative transposase | 2825 | SNOG_445480 | na | hypothetical protein |
| 16FG168 | 26 | Chromosome_13\|16FG168_ctg_12+ | strong | 1867 | y | 2,014 | 3,879 | Putative transposase | 13177 | SNOG_446090 | na | toll-like receptor 4 |
| 16FG168 | 27 | Chromosome_13\|16FG168_ctg_12+ | strong | 1866 | y | 232,455 | 234,320 | Putative transposase | 12681 | SNOG_440380 | na | hypothetical protein |
| 16FG168 | 28 | Chromosome_13\|16FG168_ctg_12+ | strong | 1866 | y | 252,072 | 250,207 | Putative transposase | 683 | SNOG_303430 | y | carbohydrate-binding module family 18 protein |
| 16FG168 | 29 | Chromosome_03\|16FG168_ctg_13C | intact/weak | 1866 | n | 1,372,321 | 1,374,186 | Putative transposase | 34499 | KAH5799522 | n | hypothetical protein |
| 16FG168 | 30 | Chromosome_15\|16FG168_ctg_14C | strong | 1866 | y | 1,108,164 | 1,106,299 | Putative transposase | 3884 | KAH5799522 | n | hypothetical protein |
| 16FG168 | 31 | Chromosome_14\|16FG168_ctg_15C | intact/weak | 1866 | y | 21,034 | 22,899 | Putative transposase | 5293 | HBI93_253790A | n | no hit |
| 16FG168 | 32 | Chromosome_14\|16FG168_ctg_15C | intact/weak | 1866 | y | 889,301 | 887,436 | Putative transposase | 1179 | SNOG_419200 | y | hypothetical protein |
| 16FG168 | 33 | Chromosome_14\|16FG168_ctg_15C | intact/weak | 1866 | n | 1,303,573 | 1,301,709 | Putative transposase | 4249 | SNOG_142850 | na | Atrophin-1 multi-domain protein |
| 16FG168 | 34 | Chromosome_14\|16FG168_ctg_15C | intact/weak | 1866 | y | 1,305,823 | 1,307,688 | Putative transposase | 1298 | SNOG_439000 | y | reverse transcriptase domain protein, partial |
| 16FG168 | 35 | Chromosome_17\|16FG168_ctg_16+ | strong | 1866 | y | 613,569 | 611,704 | no | 4531 | SNOG_431380 | na | hypothetical protein |
| 16FG168 | 36 | Chromosome_17\|16FG168_ctg_16+ | intact/weak | 1866 | y | 1,301,316 | 1,303,180 | Putative transposase | 1860 | SNOG_443930 | na | hypothetical protein |
| 16FG168 | 37 | Chromosome_18\|16FG168_ctg_17C | strong | 1866 | y | 996,785 | 998,650 | Putative transposase | 3546 | SNOG_429590 | na | putative transposase |
| 16FG168 | 38 | Chromosome_16\|16FG168_ctg_18+ | strong | 1866 | n | 550,325 | 552,190 | hypothetical protein HBI01_066130 | 69 | SNOG_308880 | y | hypothetical protein |
| 16FG168 | 39 | Chromosome_16\|16FG168_ctg_18+ | strong | 1866 | y | 1,214,384 | 1,212,519 | no | 4234 | SNOG_443930 | na | hypothetical protein |
| 16FG168 | 40 | Chromosome_21\|16FG168_ctg_20C | strong | 1866 | n | 875,473 | 877,338 | Putative transposase | 1 | SNOG_125450 | n | S-adenosyl-L-methionine-dependent methyltransferase |
| 16FG168 | 41 | Chromosome_21\|16FG168_ctg_20C | intact/weak | 1866 | n | 1,107,456 | 1,105,591 | Putative transposase | 494 | SNOG_126260 | y | hypothetical protein |
| 16FG168 | 42 | Chromosome_20\|16FG168_ctg_21C | strong | 1866 | y | 421,178 | 423,043 | no | 537 | SNOG_117240 | y | HET-domain-containing protein |
| 16FG168 | 43 | Chromosome_20\|16FG168_ctg_21C | intact/weak | 1866 | n | 617,113 | 618,978 | Putative transposase | 593 | SNOG_117230 | n | HET-domain-containing protein |
| 16FG168 | 44 | Chromosome_20\|16FG168_ctg_21C | intact/weak | 1866 | n | 622,343 | 620,478 | Putative transposase | 1180 | SNOG_425430 | y | putative transposase |
| PN315 | 1 | Chromosome_02\|PN315_nd_ctg_02C | intact/weak | 1866 | y | 193,849 | 191,984 | Putative transposase | 1300 | SNOG_446100 | n | hypothetical protein |
| PN315 | 2 | Chromosome_02\|PN315_nd_ctg_02C | intact/weak | 1866 | n | 2,937,245 | 2,935,380 | Putative transposase | 1544 | SNOG_01915 | n | hypothetical protein |
| PN315 | 3 | Chromosome_06\|PN315_nd_ctg_03+ | strong | 1866 | y | 373,706 | 375,571 | Putative transposase | 260 | SNOG_303920 | n | para-aminobenzoate synthase component 1 |
| PN315 | 4 | Chromosome_06\|PN315_nd_ctg_03+ | strong | 1866 | y | 535,506 | 537,371 | hypothetical protein HBI01_066130 | 3903 | SNOG_056930 | na | hypothetical protein |
| PN315 | 5 | Chromosome_06\|PN315_nd_ctg_03+ | intact/weak | 1866 | y | 735,166 | 737,031 | Putative transposase | 2345 | SNOG_445220 | na | CENP-B protein, partial |
| PN315 | 6 | Chromosome_06\|PN315_nd_ctg_03+ | intact/weak | 1866 | n | 1,253,423 | 1,255,288 | Putative transposase | 586 | SNOG_433610 | n | hypothetical protein |
| PN315 | 7 | Chromosome_05\|PN315_nd_ctg_04C | strong | 1866 | y | 876,266 | 874,401 | no | 58 | SNOG_158330 | y | Extradiol aromatic ring-opening dioxygenase |
| PN315 | 8 | Chromosome_05\|PN315_nd_ctg_04C | strong | 1866 | y | 1,775,007 | 1,776,872 | no | 304 | SNOG_158290 | y | non-reducing polyketide synthase PKS19-like protein |
| PN315 | 9 | Chromosome_05\|PN315_nd_ctg_04C | intact/weak | 1866 | y | 1,781,320 | 1,779,455 | Putative transposase | 756 | SNOG_443240 | n | hypothetical protein |
| PN315 | 10 | Chromosome_05\|PN315_nd_ctg_04C | intact/weak | 1866 | y | 1,788,403 | 1,786,538 | Putative transposase | 1428 | SNOG_425430 | y | putative transposase |
| PN315 | 11 | Chromosome_05\|PN315_nd_ctg_04C | intact/weak | 1866 | n | 1,833,751 | 1,835,616 | Putative transposase | 873 | SNOG_445480 | n | hypothetical protein |
| PN315 | 12 | Chromosome_05\|PN315_nd_ctg_04C | intact/weak | 1866 | n | 1,846,571 | 1,844,706 | Putative transposase | 979 | SNOG_445480 | y | hypothetical protein |
| PN315 | 13 | Chromosome_05\|PN315_nd_ctg_04C | intact/weak | 1866 | n | 1,862,333 | 1,860,468 | Putative transposase | 4321 | SNOG_029010 | na | MFS general substrate transporter |
| PN315 | 14 | Chromosome_08\|PN315_nd_ctg_05+ | intact/weak | 1866 | n | 1,564,274 | 1,562,409 | Putative transposase | 1139 | SNOG_045850 | n | Putative ribonuclease Z/Hydroxyacylglutathione hydrolase |
| PN315 | 15 | Chromosome_08\|PN315_nd_ctg_05+ | intact/weak | 1866 | y | 1,963,283 | 1,961,418 | Putative transposase | 498 | SNOG_047620 | y | hypothetical protein |
| PN315 | 16 | Chromosome_04\|PN315_nd_ctg_06C | intact/weak | 1866 | n | 1,493,920 | 1,492,055 | Putative transposase | 402 | SNOG_076960 | y | cytochrome P450 monooxygenase-like protein |
| PN315 | 17 | Chromosome_09\|PN315_nd_ctg_07C | intact/weak | 1866 | n | 411,069 | 412,934 | Putative transposase | 9334 | SNOG_436920 | na | hypothetical protein |
| PN315 | 18 | Chromosome_09\|PN315_nd_ctg_07C | intact/weak | 1866 | y | 1,223,981 | 1,222,116 | Putative transposase | 7104 | SNOG_143720 | na | major facilitator superfamily-domain-containing protein |
| PN315 | 19 | Chromosome_09\|PN315_nd_ctg_07C | intact/weak | 1866 | n | 1,225,901 | 1,224,036 | Putative transposase | 773 | SNOG_066400 | y | aminotransferase |
| PN315 | 20 | Chromosome_09\|PN315_nd_ctg_07C | intact/weak | 1866 | y | 1,544,686 | 1,542,821 | Putative transposase | 773 | SNOG_066400 | n | aminotransferase |
| PN315 | 21 | Chromosome_09\|PN315_nd_ctg_07C | strong | 1866 | y | 1,620,605 | 1,618,740 | Putative transposase | 88 | SNOG_062780 | n | heterokaryon incompatibility protein |
| PN315 | 22 | Chromosome_10\|PN315_nd_ctg_08C | intact/weak | 1866 | y | 643,076 | 641,211 | Putative transposase | 704 | SNOG_437630 | n | hypothetical protein |
| PN315 | 23 | Chromosome_13\|PN315_nd_ctg_10C | strong | 1866 | y | 251,086 | 249,221 | Putative transposase | 1633 | SNOG_303430 | na | carbohydrate-binding module family 18 protein |
| PN315 | 24 | Chromosome_13\|PN315_nd_ctg_10C | strong | 1866 | y | 266,206 | 268,071 | Putative transposase | 2348 | SNOG_442460 | na | hypothetical protein |
| PN315 | 25 | Chromosome_03\|PN315_nd_ctg_12C | intact/weak | 1866 | y | 1,294,564 | 1,296,429 | Putative transposase | 452 | SNOG_429580 | n | putative transposase |
| PN315 | 26 | Chromosome_11\|PN315_nd_ctg_13C | intact/weak | 1866 | n | 690,178 | 688,313 | Putative transposase | 313 | SNOG_093130 | n | amidase |
| PN315 | 27 | Chromosome_15\|PN315_nd_ctg_14C | strong | 1866 | y | 441,736 | 439,871 | hypothetical protein HBI01_066130 | 2723 | SNOG_103750 | na | protein kinase |
| PN315 | 28 | Chromosome_15\|PN315_nd_ctg_14C | strong | 1866 | y | 1,099,202 | 1,097,337 | hypothetical protein HBI01_066130 | 8555 | KAH4807197 | na | hypothetical protein |
| PN315 | 29 | Chromosome_14\|PN315_nd_ctg_16C | intact/weak | 1866 | n | 417,729 | 415,864 | Putative transposase | 1314 | SNOG_419200 | y | hypothetical protein |
| PN315 | 30 | Chromosome_14\|PN315_nd_ctg_16C | intact/weak | 1866 | y | 841,447 | 843,312 | Putative transposase | 5179 | SNOG_432530 | na | Retrovirushypothetical proteinrelated Pol polyprotein from transposon TNT 1hypothetical protein94 |
| PN315 | 31 | Chromosome_14\|PN315_nd_ctg_16C | intact/weak | 1866 | n | 1,258,530 | 1,260,395 | Putative transposase | 626 | SNOG_434920 | y | Membrane-bound metallopeptidase |
| PN315 | 32 | Chromosome_19\|PN315_nd_ctg_18C | intact/weak | 1866 | y | 724,670 | 722,805 | Putative transposase | 1298 | SNOG_436920 | n | hypothetical protein |
| PN315 | 33 | Chromosome_22\|PN315_nd_ctg_19C | intact/weak | 1866 | y | 621,751 | 623,616 | Putative transposase | 11421 | SNOG_439850 | na | hypothetical protein |
| PN315 | 34 | Chromosome_23\|PN315_nd_ctg_23C | intact/weak | 1866 | y | 250,174 | 252,039 | Putative transposase | 2052 | SNOG_439520 | na | hypothetical protein |
| PN315 | 35 | Chromosome_05\|PN315_nd_ctg_24+ | intact/weak | 1866 | y | 102,319 | 104,184 | Putative transposase | 138 | SNOG_448570 | y | hypothetical protein |
| PN315 | 36 | Chromosome_05\|PN315_nd_ctg_24+ | intact/weak | 1866 | y | 112,693 | 114,558 | Putative transposase | 1236 | SNOG_441370 | n | hypothetical protein |

### **STable 8**: GC-content and SNP found in Molly

| Collection | Isolate | Isolate_tree_name | Year of collection | Copy # | Modification | | | | Genome-GC-content | | Molly-GC-content | | Molly-SNP-strong | | | | Molly-SNP-weak | | | |
|---|---|---|---|---|---|---|---|---|---|---|---|---|---|---|---|---|---|---|---|---|
| | | | | | Strong (>20 SNP) | Weak/intact (0-20 SNP) | Percent strong | Percent weak | LRAR | non-LRAR | Intact/weak | Strong | Total-SNP | RIP-like-SNP | %-RIP-like-SNP | %-RIP-effected | Total-SNP | RIP-like-SNP | %-RIP-like-SNP | %-RIP-effected |
| OLD_group 1 | 15FG38_nanopore | O124_15FG38 | 2015 | 19 | 9 | 10 | 47 | 53 | 27.50 | 51.92 | 0.52 | 0.45 | 363 | 363 | 100.00 | 19.45 | 24 | 24 | 100.00 | 1.29 |
| OLD_group 5 | 16FG168_nanopore | O155_16FG168 | 2016 | 44 | 27 | 17 | 61 | 39 | 27.90 | 51.71 | 0.52 | 0.47 | 471 | 463 | 98.30 | 24.81 | 21 | 4 | 19.05 | 0.21 |
| Current_group 7 | PN315_nanopore | C315_PN315 | 2019 | 36 | 9 | 27 | 25 | 75 | 27.49 | 52.22 | 0.52 | 0.48 | 264 | 264 | 100.00 | 14.15 | 11 | 11 | 100.00 | 0.59 |
| OLD_group 1 | WAC8384 | O10_WAC8384 | 1990 | 34 | 31 | 3 | 91 | 9 | 28.06 | 51.56 | 0.52 | 0.44 | 565 | 531 | 93.98 | 28.46 | 106 | 104 | 98.11 | 5.57 |
| OLD_group 1 | Mur_51 | O37_Mur_51 | 2011 | 13 | 13 | 0 | 100 | 0 | 27.51 | 51.63 | na | 0.43 | 563 | 530 | 94.14 | 28.40 | na | na | na | na |
| Current_group 1 | PN275 | C275_PN275 | 2019 | 29 | 16 | 13 | 55 | 45 | 27.73 | 51.60 | 0.52 | 0.49 | 557 | 524 | 94.08 | 28.08 | 34 | 31 | 91.18 | 1.66 |
| Current_group 1 | PN295 | C275_PN295 | 2019 | 30 | 3 | 27 | 10 | 90 | 28.25 | 52.04 | 0.52 | 0.44 | 250 | 250 | 100.00 | 13.40 | 16 | 15 | 93.75 | 0.80 |
| Current_group 1 | PN312 | C312_PN312 | 2019 | 23 | 23 | 0 | 100 | 0 | 27.92 | 51.52 | na | 0.47 | 267 | 267 | 100.00 | 14.31 | na | na | na | na |
| Current_group 1 | PN328 | C328_PN328 | 2019 | 26 | 26 | 0 | 100 | 0 | 27.97 | 51.87 | na | 0.43 | 467 | 466 | 99.79 | 24.97 | na | na | na | na |
| Current_group 1 | PN329 | C329_PN329 | 2019 | 28 | 28 | 0 | 100 | 0 | 27.46 | 52.04 | na | 0.43 | 466 | 465 | 99.79 | 24.92 | na | na | na | na |
